# Equine herpesvirus 4 infected domestic horses associated with Sintashta spoke-wheeled chariots around 4,000 years ago

**DOI:** 10.1101/2023.09.08.556822

**Authors:** Ophélie Lebrasseur, Kuldeep Dilip More, Ludovic Orlando

**Author notes:** Corresponding authors (OL) (KM).

## Abstract

Equine viral outbreaks have disrupted the socio-economic life of past human societies up until the late 19th century, and continue to be of major concern to the horse industry today. With a seroprevalence of 60-80%, equine herpesvirus 4 (EHV-4) is the most common horse pathogen on the planet. Yet, its evolutionary history remains understudied. Here, we screen the sequenced data of 264 archaeological horse remains to detect the presence of EHV-4. We recover the first ancient EHV-4 genome with 4.2X average depth-of-coverage from a specimen excavated in the Southeastern Urals and dated to the Early Bronze Age period, approximately 3,900 years ago. The recovery of an EHV-4 virus outside of the upper respiratory tract not only points to an animal particularly infected, but also highlights the importance of post-cranial bones in pathogen characterisation. Bayesian phylogenetic reconstruction provides a minimal time estimate for EHV-4 diversification to around 4,000 years ago, a time when modern domestic horses spread across the Central Asian steppes together with spoke-wheeled Sintashta chariots, or earlier. The analyses also considerably revise the diversification time of the two EHV-4 subclades from the 16th century based solely on modern data to nearly a thousand years ago. Our study paves the way for a robust reconstruction of the history of non-human pathogens and their impact on animal health.

## Introduction

Equine herpesviruses are pathogens of major economic, clinical and epidemiological importance to the horse industry worldwide (Patel and Heldens 2005; Lunn *et al*. 2009; Ma, Azab and Osterrieder 2013; EFSA Panel on Animal Health and Welfare (AHAW) *et al*. 2022). The equine alphaherpesvirus 1 (EHV-1) outbreak of early 2021 in Europe was the most severe in decades, which led the European Food Safety Authority to implement disease prevention and control measures in accordance with the Animal Health Law (Patel and Heldens 2005; EFSA Panel on Animal Health and Welfare (AHAW) *et al*. 2022). Following infection by EHV-1, horses develop respiratory syndromes that can not only lead to abortions in pregnant mares and increase mortality rates in neonatal foals, but can also evolve into myeloencephalitis (Telford *et al*. 1992; Studdert *et al*. 2003; Pavulraj *et al*. 2021). Other equine herpesviruses, such as equine gammaherpesviruses 2 and 5, are ubiquitous in horse populations (70-90% seroprevalence) but rarely lead to symptomatic infections, while the EHV-3 alphaherpesvirus causes Equine Coital Exanthema, a venereal mucocutaneous disease (Stasiak, Dunowska and Rola 2018; Vissani, Damiani and Barrandeguy 2021). The EHV-4 alphaherpesvirus, on the contrary, primarily causes upper respiratory tract disease, but only occasionally advances into neurological disorders and abortions (Telford *et al*. 1998; Khattab *et al*. 2022). Yet, EHV-4 remains a major concern due to its higher seroprevalence across the world (60-80% amongst horses, donkeys and their mule hybrids (Mekonnen, Eshetu and Gizaw 2017; Pavulraj *et al*. 2021)), which may derive from its capacity to develop year-round infections as opposed to winter for EHV-1 (Matsumura *et al*. 1992; Ma, Azab and Osterrieder 2013; Vaz *et al*. 2016; Pavulraj *et al*. 2021). It is also known to establish life-long latency in the trigeminal ganglion and the submandibular lymph nodes following primary infection, leading to likely periodical reactivation (Matsumura *et al*. 1992; Borchers, Wolfinger and Ludwig 1999; Patel and Heldens 2005; Osterrieder and Van de Walle 2010; Ma, Azab and Osterrieder 2013; Vargas-Bermudez *et al*. 2018; Pavulraj *et al*. 2021). Owing to the considerable economic threat EHV-4 poses, much attention has been given to its pathology. Yet, its evolutionary history in the context of horse domestication remains uninvestigated.

Textual sources provide evidence for respiratory tract disease in horses as early as 412 BCE in Sicily (Williams 1924). Whether EHV-4 or any other virus causing influenza-type diseases was responsible for these outbreaks cannot, however, be determined from historical accounts nor from archaeological bone assemblages in the absence of obvious pathological lesions. As a consequence, the earliest confirmed record of an equine herpesvirus-like disease dates back to 1936, when the first EHV infection was described in the United States by Dimock and Edwards (Dimock and Edwards 1933, 1936; Bryans and Allen 1989). Further reports followed shortly after, for example in Hungary in 1941 (Manninger and Csontos 1941; Bryans and Allen 1989). EHV-1 and EHV-4 were, however, only recognised as genetically closely-related yet distinct viruses in 1981 (Telford *et al*. 1998). Therefore, determining which of these two virus types was responsible for the pre-1981 outbreaks remains difficult until serological tests became available from 1985. (Duxbury and Oxer 1968; Burrows and Goodridge 1974; Yeargan, Allen and Bryans 1985; Allen and Bryans 1986; Matsumura *et al*. 1992; Crabb and Studdert 1995; Telford *et al*. 1998; Gilkerson *et al*. 1999; Patel and Heldens 2005; Vaz *et al*. 2016). Since such retrospective genetic analyses have been limited to samples collected in the second half of the 20^th^ century, the potential importance of EHV-4 outbreaks in deeper historical times remains overlooked.

Over the last decade, methodological advances in ancient DNA research have improved access to genetic information of past populations (Orlando *et al*. 2021). This has revolutionised our understanding of human history, not only revealing changing patterns of mobility, admixture and selection through space and time (Nielsen *et al*. 2017), but also the causative agents of past epizootics (Spyrou *et al*. 2019a). Ancient DNA analyses of human bone assemblages have, thus far, led to the successful genome characterisation of a diversity of pathogens, including both bacterial (e.g. *Yersinia pestis* (Spyrou *et al*. 2022), *Mycobacterium leprae* (Schuenemann *et al*. 2013), *Salmonella enterica* (Key *et al*. 2020)) and viral (e.g. HBV (Kocher *et al*. 2021), smallpox (Duggan *et al*. 2016), HSV (Guellil *et al*. 2022)). Faunal archaeological remains have, however, received considerably less attention with respect to pathogen DNA characterisation, despite the reported impact of animal disease outbreaks throughout history (Roman-Binois 2017; Frantz *et al*. 2020). For example, glanders, another established respiratory tract disease, and its impact on cavalries may have been crucial in deciding the outcome of battles, which is a subject of repeated debate among historians (Chandler 1963; Sharrer 1995; Dvorak and Spickler 2008). The extensive sequencing of horse archaeological remains dating to the last 50,000 years provide a unique opportunity to start mapping infectious diseases prior to, during, and following domestication around 4,200 years ago in the Western Eurasian steppes (Librado *et al*. 2021; Orlando *et al*. 2021), with the potential to solve long-standing historical debates.

In this study, we screened the sequencing data of 264 ancient horses first presented by Librado and colleagues (Librado *et al*. 2021) for the presence of EHV-1 and EHV-4. We identify one Bronze Age horse as positive for EHV-4. The underlying archaeological bone was radiocarbon dated to 1,853 cal BCE and found in association with Sintashta material culture, which developed the spoked wheel technology and spread horse-drawn chariots across the Asian steppes between ca. 3,800 and 4,100 years ago (Anthony 2007). The available sequence data led to the characterisation of a partial EHV-4 genome sequence, with a 4.2X average depth-of-coverage. Phylogenetic modelling reveals that EHV-4 was pervasive in domestic horse populations for at least 4,000 years. We identify two amino acid changes that have been positively selected following the emergence of modern subclades around 1,000 years ago in ORF48 which codes for protein UL14 supposedly important in virion morphogenesis. This work, and the exponentially increasing sequencing data made available from faunal archaeological remains (Frantz *et al*. 2020), opens for a deeper understanding of the past history of infectious diseases in domestic animals, and their impact on health, economy and societies.

## Results

### EHV-4 Identification and Authentication

Raw FastQ sequence files underlying the comprehensive dataset of ancient horse genomes published in (Librado *et al*. 2021) were mapped against de-novo assemblies of EHV-1 and EHV-4 genomes that were used as reference genomes (Telford *et al*. 1992, 1998). The sequence data from the five libraries constructed from the DNA extracts of the tibia of a mare labelled UR17x29 (BioSample: SAMEA9533417) and recovered on the burial site of Kamennyi Ambar 5 (kurgan 8, grave 2), Kartaly District, south of the Chelyabinsk Oblast, Russia (Fig 1A; see (Librado *et al*. 2021) for complete archaeological details), showed a minute fraction of high-quality alignments against the EHV-4 genome (isolate 306-74, Genbank Acc. KT324743, (Vaz *et al*. 2016)). This specimen was previously radiocarbon dated to 1,936-1,770 cal BCE (95.4% probability, InCal2020 (Reimer *et al*. 2020); 3,530±30 uncal. BP, UCIAMS-199248), providing a median age estimate of 1,853 cal BCE. In order to confirm the above results as true-positives for EHV-4, we carried out competitive mapping against candidate reference genomes in HAYSTAC (Dimopoulos *et al*. 2022), and alignment-free taxonomic assignment in Kraken2 (Wood, Lu and Langmead 2019). The potential candidates were selected to represent the default list of pathogens from HOPS (n= 220) which we complemented with 27 horse-specific pathogens (see Table S1 for details). Both approaches returned consistent results across all libraries (ERR6466108-ERR6466112), supporting the presence of opportunistic pathogens abundant in the environment (e.g. *Mycobacterium* complex), as well as EHV-4 and another equine-specific pathogen, namely *Prescottella equi* (Tables S2 and S3). While *P. equi* represents a well-known opportunistic pathogen in immuno-compromised horses, we decided to focus the downstream analyses on EHV-4, as the only one candidate supported with a sufficient number of reads to assess post-mortem DNA damage profile and associated with significant genome coverage (>0.05x). We therefore considered EHV-4 BAM alignment that we filtered for PCR duplicates and alignments showing mapping quality scores inferior to 37. This left a total of 13,019 sequence alignments, corresponding to an average depth-of-coverage of 4.24-fold, with 68.57% of the sites covered ≥3X (Fig 2A). Local coverage dropped within GC-rich and/or duplicated repeat regions including ORF64-ORF66, as expected (Fig 2A).

**Figure 1.**
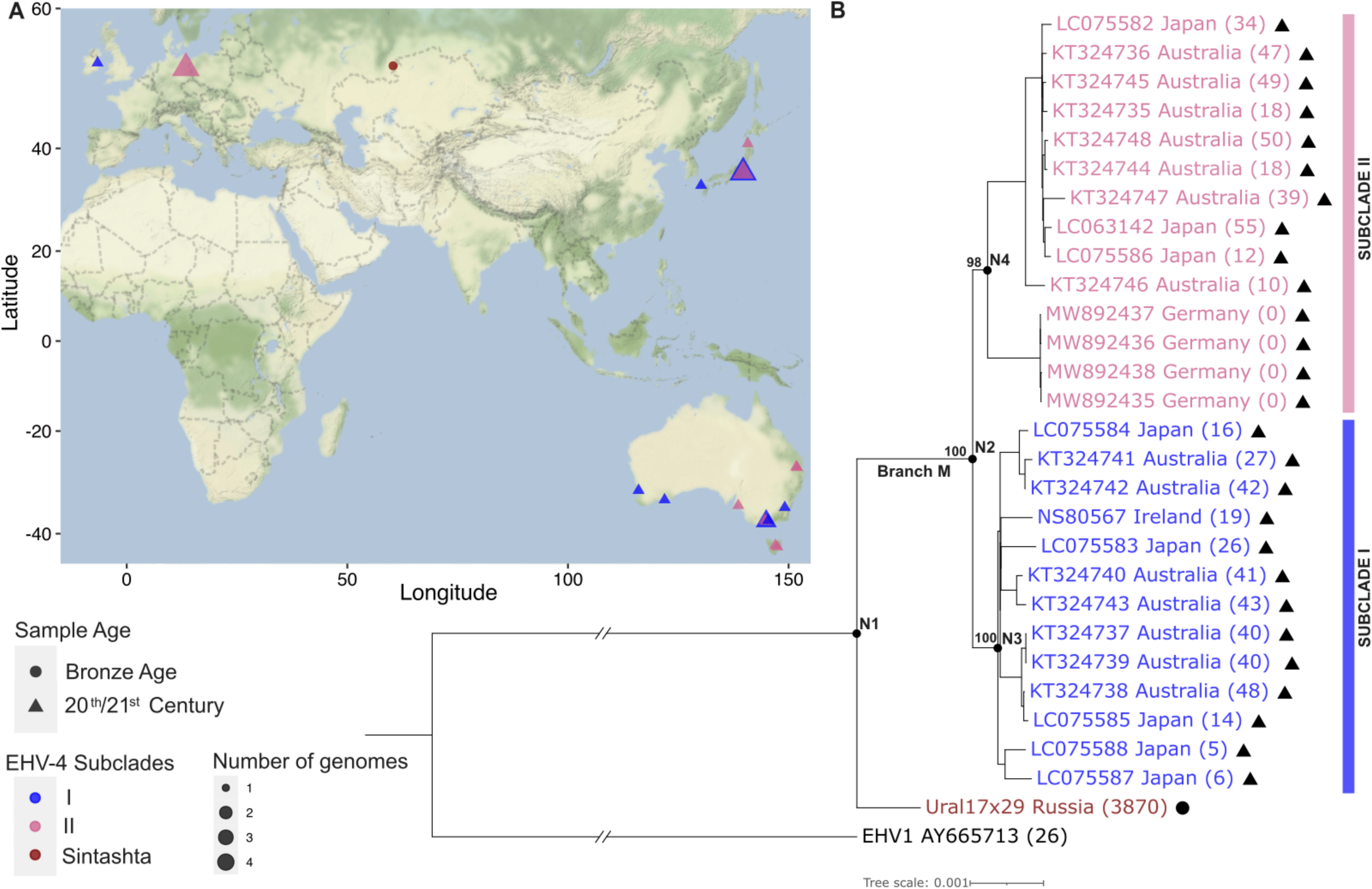
Sample map and phylogenetic relationships. (A) Location of the samples used in this study, including subclades, sample dates and numbers of genomes. For samples with geographic information at country-level, the coordinates of the regional or country capital were used. (B) Maximum-Likelihood tree and node bootstrap support. Sample names comprise Genbank accession numbers, sampling locations and, in parentheses, sampling dates in years Before Present (BP) with ‘2017’ set as ‘Present’ based on the most recently sampled isolates. The symbols refer to Figure 1A. EHV-1 was used as an outgroup.

**Figure 2.**
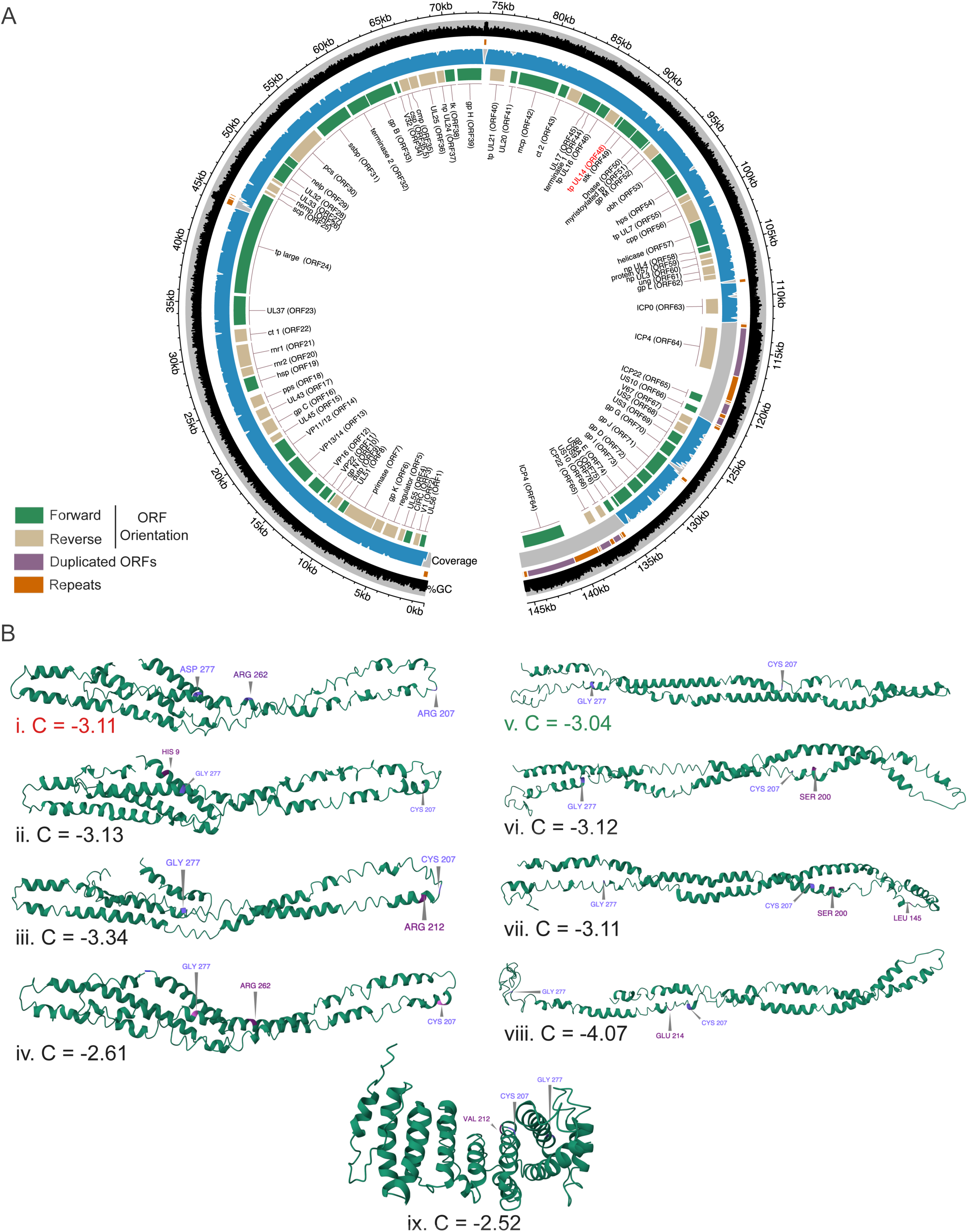
EHV-4 genome organisation and protein structure predictions. (A) %GC and coverage was calculated within non-overlapping 50bp and 100bp sliding windows, respectively. Coverage is given as a percentage based on a maximum of 7.3x. The Open Reading Frame encoding for the tegument protein (tp) UL14 (highlighted in red) was found to have been positively selected along the branch ancestral to subclades I and II (Branch M, Figure 1B). The grey area on the coverage track shows the drop in coverage represented by the duplicated repeat regions including ORF64-ORF66. cmp: capsid maturation protein; cpp: capsid portal protein; csp: capsid scaffold protein; ct. capsid triplex; dutp: deoxiuridine triphosphatase; gp: glycoprotein; hps: helicase primase subunit; hsp: host shutoff protein; mcp: major capsid protein; nelp: nuclear egress lamina protein; nemp: nuclear egress membrane protein; np: nuclear protein; pcs: DNA polymerase catalytic subunit; obh: origin binding helicase; pps: DNA polymerase processivity subunit; rnr: ribonucleotide reductase; scp: small capsid protein; ssbp: single-stranded DNA binding protein; stk: serine/threonine protein kinase; tk: thymidine kinase; tp: tegument protein; ung: uracil DNA glycosylase. (B) Secondary structures predicted by I-TASSER for UL14 proteins. Amino acid substitutions are shown in light purple (substitutions differing between the ancient and modern versions) and in dark purple (substitutions differing from the most commonly observed version). The confidence score C denotes the quality of predicted models ranging from -5 to 2. The predictions for the most common version and UR17x29 are shown in green and red c-scores, respectively.

All the aligned sequences consisted of read pairs showing sufficient overlap to be collapsed as part of a single template, due to the extreme short size of the DNA molecules recovered (average = 46.9 bp; median= 42 bp; 95% confidence interval = 27-84 bp). Additionally, the genomic positions immediately preceding alignment starts were enriched in cytosine residues, while those immediately following were enriched in guanine residues. This was true for both horse and EHV-4 read alignments (Figure S1A and S1B, respectively), and reflects the enzymatic treatment of DNA extracts with the USER mix (Rohland *et al*. 2015), which is known to cleave DNA strands at those cytosines deaminated into uracils post-mortem, leaving only moderate C→T nucleotide mis-incorporations characteristic of ancient DNA damage (Orlando *et al*. 2021) (0.706% at alignment starts *vs* 0.211% on average across the following 24 positions). Complementary G→A mis-incorporation rates were inflated at alignment stops (0.468% *vs* 0.211% on average across the preceding 24 positions). The observed sequence complementarity of mis-incorporation signatures at read termini is typical of ancient DNA data, once incorporated into double stranded DNA libraries following USER enzymatic treatment (Orlando *et al*. 2021). Together with the short size of the DNA templates characterised and the stringent quality filters used in sequence alignments, these findings suggest that, despite representing only a limited fraction of the metagenomic content of the archaeological bone analysed, authentic ancient EHV-4 DNA molecules were sequenced together with the DNA molecules of their host, which covered the horse reference genome at an average of 2.08-fold depth-of-coverage (Librado *et al*. 2021).

### Phylogenetic Analyses

At the time of writing, a total of 27 published EHV-4 genomes are available from previous studies. These were generated from fresh biological tissues that were collected in Australia, Germany, Ireland and Japan between 1962 and 2017 (Telford *et al*. 1998; Vaz *et al*. 2016; Izume *et al*. 2017; Pavulraj *et al*. 2021) (Table S4). However, using MAFFT v7.453 (Katoh and Standley 2013), these sequences and the ancient UR17x29 genome characterised here for the first time proved difficult to align against the EHV-1 reference sequence (strain Ab4p; AY665713). The inclusion of EHV-1 as an outgroup is, however, required to polarise phylogenetic reconstructions. We, thus, aligned each ORF independently and concatenated all the resulting alignments together before undertaking phylogenetic reconstruction, with the exception of ORFs 24, 47/44 and 71 which include numerous tandem repeats, duplicated ORFs (corresponding to ORF64-ORF67 for EHV-1 but to ORF64-ORF66 for EHV-4; Fig 2A) and previously identified tandem repeat regions (Izume *et al*. 2017). The concatenation of ORF sequences provided a final alignment comprising 104,418 orthologous sites. Maximum-likelihood phylogenetic analysis reiterated the previously reported subdivision of EHV-4 viruses within two main subclades, referred to as subclades I and II (Fig 1B). While these subclades are not defined by geography and/or pathogenicity, subclade I is known to almost exclusively consist of abortion isolates (Izume *et al*. 2017; Kolb *et al*. 2017). The phylogenetic reconstruction also returned strong support for the existence of a subgroup within subclade II defined by a single T-to-C point mutation affecting position 624 of ORF33 (61,770 on EHV-4 genome reference AF030027) (Telford *et al*. 1998; Pavulraj *et al*. 2021). Our maximum-likelihood phylogenetic analysis and pairwise genetic distance matrix revealed that the UR17x29 ancient sequence was considerably distant to the EHV-1 outgroup (0.32739), but represented a close relative to all modern EHV-4 sequences hitherto sequenced, showing pairwise genetic distances no larger than 0.00369 (Fig 1B, Table S5). The UR17x29 genome was, however, found to fall outside the monophyletic group of modern EHV-4 strains including both subclades. This supports the UR17x29 genome as deriving from an ancient, divergent EHV-4 strain not segregating today, ruling out modern contamination.

### Recombination

The extensive recombination previously reported in EHV-4 (Vaz *et al*. 2016; Kolb *et al*. 2017) may have introduced bias in the phylogenetic reconstructions above, as these were based on genome-wide ORF alignments. The multiple reticulate and pairwise homoplasy (Phi) test analyses implemented in SplitsTree4 (Huson and Bryant 2006) indeed provided significant statistical support for the presence of recombination in our dataset (*p*-value = 2.628E-8). To further address the extent to which such recombination may have affected our phylogenetic findings, we mapped the recombination breakpoints using the most-commonly used detection methods (RDP (Martin and Rybicki 2000), GENECONV (Padidam, Sawyer and Fauquet 1999), Chimaera (Posada and Crandall 2001), MaxChi (Maynard Smith 1992), BootScan (Martin *et al*. 2005), SiScan (Gibbs, Armstrong and Gibbs 2000) and 3Seq (Lam, Ratmann and Boni 2018)) implemented in RDP5 (Martin *et al*. 2021) and relying on two successive analytical steps. Considering all methods together, as well as conservative *p*-values and recombination scores, these analyses predicted the presence of 12 recombination events throughout the EHV-4 genome (Table S6). The largest genomic block predicted to not be affected by recombination spanned approximately 85kb (nucleotide coordinates 4,254-90,128 of the EHV-4 reference genome AF030027 (Telford *et al*. 1998) with tandem repeats excluded), which was further confirmed to be devoid of recombination through the Phi test implemented in SplitsTree4 (*p*-value = 0.1121). Phylogenetic reconstruction through IQ-TREE (Minh *et al*. 2020) (Fig S2) returned the same topology as that obtained on the genome-wide ORF alignments, ruling out any significant impact of recombination in the evolutionary history reconstructed above.

### Divergence Times

The characterization of a deeply divergent ancient genome sequence provided the first opportunity to calibrate the timing of major phylogenetic divergence events underlying the EHV-4 evolutionary tree. To achieve this, we applied the Bayesian framework implemented in BEAST v2.6.7 (Bouckaert *et al*. 2019) to the 85kb recombination-free alignment. Bayesian phylogeny was reconstructed under the birth-death skyline serial model with tip-dating, and assuming either a strict, or a relaxed lognormal clock. Model selection through Nested Sampling analyses (Skilling 2006; Maturana Russel *et al*. 2019) as implemented in the Beast2 NS package v.1.1.0. provided support for the strict clock model, which was therefore used for assessing divergence times. The estimated age of the EHV-4 tMRCA (time to the Most Recent Common Ancestor) was dated to 1,905 BCE (median; 95% confidence interval, CI = 2,090-1,853 BCE; node N1, Figs 3A and 3B). This date estimate is contemporary to the spread of the Sintashta culture, which is also associated with the archaeological context into which sample UR17x29 was found. Therefore, our work provides definitive proof that EHV-4 infected horses as far back as the Bronze Age. We caution that the 1,905 BCE age estimate should not be confused with the earliest horse infection by EHV-4. In fact, it is reasonable that EHV-4-like alphaherpesviruses already infected equine ancestors as they diversified in modern species 4 to 4.5 million years ago (Orlando *et al*. 2013), given that the alphaherpesvirus evolutionary tree mirrors that of their mammalian hosts, in line with a co-divergence history of viruses with their hosts. The Bronze Age virus characterised in this study also significantly revised the divergence time between subclades I and II (node N2) to around 1,071 CE (Fig 3A, median; 95% CI = 899-1,214 CE), versus 1,531 CE (Fig S3, 95% CI = 1,361-1,669 CE). Therefore, the two main phylogenetic subclades structuring modern EHV-4 diversity already emerged a thousand years ago.

**Figure 3.**
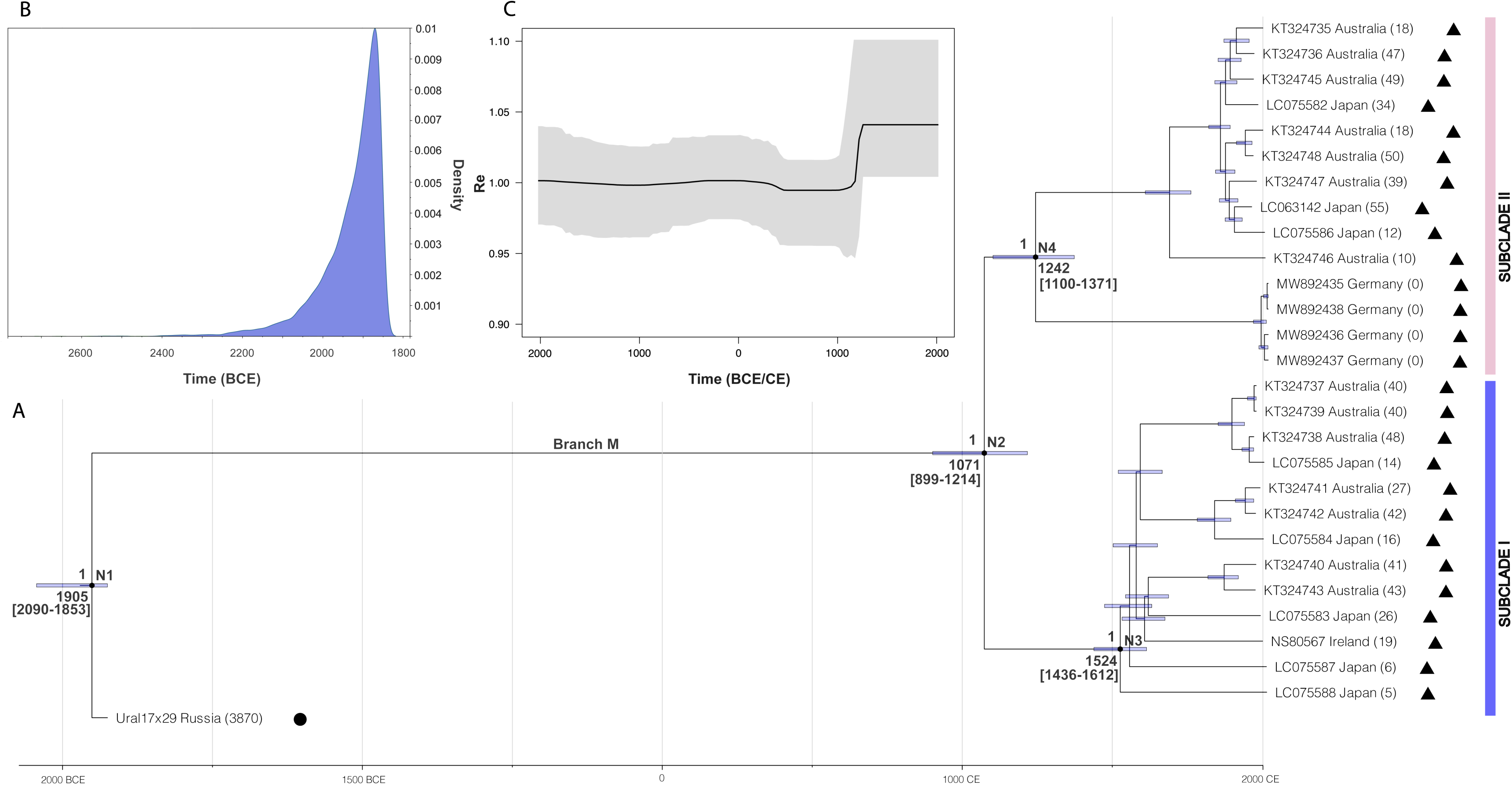
Time-scaled EHV-4 evolutionary history. (A) Bayesian phylogeny under the strict clock and birth-death skyline serial models based on the ancient and modern dataset. Node posterior supports are provided above nodes to the left, whilst median estimates of divergence times are provided below nodes together with the 95% confidence interval in square brackets [Height_95%_HPD]. Sequence labels refer to Genbank accession numbers, sampling locations and, in parentheses, sampling dates formatted as years Before Present (BP) with ‘Present’ set to ‘2017’. These dates were used for tip-dating calibration. The symbols refer to Figure 1A. (B) Posterior distribution of the time to the Most Recent Common Ancestor of all EHV-4 strains considered in this study (N1). Time is indicated in calendar years (BCE: Before Common Era, CE: Common Era). (C) Temporal trajectory for the reproduction number R_e_ of the EHV-4 virus.

### Positive Selection

We next leveraged the robust phylogeny reconstructed above to test for the presence of sites that were positively selected along the branch leading to modern subclades I and II (branch M, Figs 1B and 3A). The branch-site model implemented in PAML v.4.9 (Yang 2007) revealed ORF48 as the only positively selected ORF (p<0.05), with two of the five amino acid substitutions suggested as potentially adaptive appearing fixed in all modern strains (C619T and A830G changing Cysteine into Arginine and Aspartate into Glutamate, respectively). The prediction of three other polymorphic sites as potentially adaptive along branch M may result from a combination of a backward mutation in the German samples clustering within subclade II (G785A changing Arginine into Glutamine), and confounding consequences of recombination as ORF48 includes one predicted recombination breakpoint. Similar selection scans failed at detecting selection candidates between subclades I and II.

ORF48 encodes for the UL14 tegument protein, which is hypothesised to be involved in virion morphogenesis, by transporting other tegument proteins from the nucleus into the cytoplasm (Telford *et al*. 1998). The best secondary structures predicted by I-TASSER (Yang *et al*. 2015) showed two different prominent models comprising eight versions of UL14 proteins found amongst modern strains and ancient UR17x29 (Fig 2B) with an exception of the Irish strain NS80567 (Fig 2B, ix). No structural difference was observed between UL14 protein structures of modern strains and of UR17x29 as shown in Fig 2B, suggesting that the two point mutations observed above may not play a decisive role in the secondary structure of UL14 protein. However, due to the lack of confirmed structure/s of homologous protein/s, our analyses cannot be considered as conclusive but rather remain predictive in nature. UL14 has been shown to possess anti-apoptotic and heat shock protein-like activities while playing a pivotal role in protein localisation in various herpesviruses increasing longevity of an infected cell (Cunningham *et al*. 2000; Yamauchi *et al*. 2002, 2008; De Martino *et al*. 2007; Oda *et al*. 2016). Therefore, the strong signature of positive selection detected for UL14 suggests that key evolutionary changes may have contributed to post-replication viral sustenance in infected cells. *In-vitro* study of UL14 deletion mutants and X-ray crystallography-based structure reconstructions are, however, needed to fully understand the role and the exact functional consequences of the two point mutations identified here.

## Discussion

This study demonstrates that EHV-4 was present in the southeastern Urals 1,853 BCE. This pushes back the age of the oldest EHV-4 virus characterised so far by nearly 3,800 years. The viral DNA material was preserved in a tibia that was found in a cemetery associated with the Sintashta culture. This culture is well-known for triggering important socio-economic changes in the Bronze Age Eurasian steppes. These range from proto-urban developments in settlement organisation, metallurgy (Hanks *et al*. 2018; Librado *et al*. 2021), and the invention of spoke-wheeled chariotry (Hanks *et al*. 2018; Klecel and Martyniuk 2021; Librado *et al*. 2021), which not only revolutionised transport but also accompanied the rise of innovative warfare technologies (Drews 1993; Anthony 2009; Klecel and Martyniuk 2021). The spread of Sintashta-associated populations together with their horse-driven spoke-wheeled chariots fuelled the expansion of Indo-Iranian languages across Central Asia and beyond (Allentoft *et al*. 2015; Narasimhan *et al*. 2019; Lazaridis *et al*. 2022). Interestingly, patterns of mitochondrial, Y-chromosomal and nuclear variation in ancient horses (Gaunitz *et al*. 2018; Fages *et al*. 2019; Librado *et al*. 2021) recently showed that the genetic lineage that gave rise to modern domestic horses (DOM2) expanded both demographically and geographically in the late 3rd millennium BCE. The period spanning 2,090-1,853 BCE, which we estimated for the EHV-4 tMRCA, not only coincides with the rapid increase of the DOM2 effective size by at least an order of magnitude, but also their expansion outside the domestication homeland, from the lower Volga-Don basin into Moldova, Anatolia and Central Asia (Librado *et al*. 2021). In fact, this period marks the time when horses became paramount to human societies (Kelekna 2009). The increasingly global demand for horses in relation to chariotry, warfare and mobility, as well as the underlying changes in management practices, would, thus, have expedited the spread of EHV-4. This historical turning point may have also facilitated the dispersal of other equine diseases across long distances, and human diseases, as recently suggested for the plague (Rascovan *et al*. 2019; Valtueña *et al*. 2022). The tMRCA of all EHV-4 strains including our ancient genome slightly postdates the rise of DOM2 horse domestic bloodlines. This may indicate that domestication resulted in considerable subsampling of EHV-4 diversity, as a result of a demographic bottleneck within host populations, and/or that the rapid demographic growth and expansion of DOM2 horses has favoured the surfing of only a subset of the EHV-4 diversity then present. The latter has been previously reported for parasites in which diversity is lost as the host population expands away from its original environment (White and Perkins 2012). Disentangling both scenarios will require mapping out the EHV-4 diversity in pre-DOM2 lineages. Given the relatively limited EHV-4 prevalence rate measured through shotgun sequencing in our ancient horse panel, this will likely require dedicated target-enrichment approaches. Furthermore, our knowledge of modern EHV-4 diversity is currently limited to a few countries only and should be characterised globally before the true impact of EHV-4 infection during domestication can be measured.

In this study, only a single EHV-4 genome could be characterised amongst 135 ancient DOM2 individuals (0.7%). This is in striking contrast with the high seroprevalence observed in unvaccinated equids today (60-80% (Mekonnen, Eshetu and Gizaw 2017; Pavulraj *et al*. 2021)). At first glance, this would suggest a low seroprevalence rate in ancient populations. However, EHV-4 infections generally start through the inhalation of aerosols containing viral particles (Ma, Azab and Osterrieder 2013; Khusro *et al*. 2020), and viral replication remains primarily restricted to the epithelial cells of the upper respiratory tract (Osterrieder and Van de Walle 2010; Vandekerckhove *et al*. 2011; Ma, Azab and Osterrieder 2013). Such soft tissues are generally not available in the archaeological record, which considerably limits our detection capacity. Regardless, the detection rate of viral material in extent animals is known to not necessarily reflect seroprevalence, as reported in other herpesviruses, such as the human Varicella zoster virus (Moustafa *et al*. 2017). Therefore, the single positive case detected in our study does not inform on past seroprevalence rates but only reflects a horse experiencing high viral replication with substantial viral particles in the systemic blood circulation (Pusterla *et al*. 2005; Vandekerckhove *et al*. 2011).

We observe that our estimates of R_0_ remain close to 1 throughout the whole timeline investigated here, including following the diversification of subclades I and II (Fig 3C), which suggests that the epidemiological spread was from one individual to another on average rather than exponential. However, we caution that our estimates for the viral basic reproduction number are based on a single ancient sample which considerably limits our resolution in the past. More ancient EHV-4 genomes sequences are needed before the true dynamics of past EHV-4 outbreaks can be elucidated.

Our work, although limited to only a partial sequence of the EHV-4 genome, provides the first temporal calibration for the evolutionary tree of EHV-4 viruses. In contrast to horses for which an abundant palaeontological record provides fine-grained resolution into macroevolutionary changes (Macfadden 2005), equine viruses and pathogens hardly leave any obvious fossil remains. Therefore, the molecular traces left by the viral pathogens once infecting ancient equine populations offer the unique opportunity to access and chart past epidemiological outbreaks through space and time. The cultural and osteological material composing the archaeological context also adds crucial information to refine the social, societal, and environmental conditions driving the spread of past infectious diseases. The immense potential of ancient DNA for advancing understanding of the evolutionary history of pathogens is now fully established for humans (Spyrou *et al*. 2019a), following pioneering work characterising ancient plague (Spyrou *et al*. 2019b; Valtueña *et al*. 2022), tuberculosis (Bos *et al*. 2014; Kay *et al*. 2015; Kerner *et al*. 2021), leprosy (Schuenemann *et al*. 2013, 2018; Pfrengle *et al*. 2021) and more. Beyond bacterial pathogen vectors, this approach has also started to illuminate the past history of both DNA (e.g. HBV (Kocher *et al*. 2021), HSV (Guellil *et al*. 2022)) and RNA (e.g. Influenza (Taubenberger *et al*. 2005), Measles (Düx *et al*. 2020)) viruses. Despite such potential and the growing interest for animal domestication in ancient DNA research (Frantz *et al*. 2020), this approach remains overlooked for non-human animals. Yet, the domestication process itself, as well as the mass production sustaining an ever-increasing human population, are known to have increasingly exposed animals to infectious diseases, with particularly detrimental consequences on their health (Morand, McIntyre and Baylis 2014). Therefore, the methodology developed in our study, and more generally the tools supporting ancient DNA research, provide an effective framework to both map out animal pathogens in the past, and to assess the impact of domestication and changing management practices on animal health.

## Materials and Methods

### Sample and sequence alignment

We mapped the shotgun sequence data underlying the comprehensive dataset of ancient horse genomes published in (Librado *et al*. 2021) against the complete reference genomes of EHV-1 (AY665713 (Telford *et al*. 1992)) and EHV-4 (KT324743 (Vaz *et al*. 2016)). Complete archaeological details for all samples pertaining to this comprehensive dataset can be found in the Supplementary Methods of (Librado *et al*. 2021). The sequence data was processed using PALEOMIX (Schubert *et al*. 2014) and the BWA aligner (Li and Durbin 2009), disregarding seed (-l 1024) and using the stringent alignment filters of (Spyrou *et al*. 2019b) (i.e. -n 0.1, quality scores greater or equal to 37). The UR17x29 sample (BioSample: SAMEA9533417) returned substantial high-quality alignments for EHV-4. To confirm the reads were true positives, we ran HAYSTAC (Dimopoulos *et al*. 2022) and Kraken2 (Wood, Lu and Langmead 2019) on all 5 libraries (ERR6466108-ERR6466112). For both HAYSTAC and Kraken2, we referred to the HOPS default pathogen list (https://github.com/rhuebler/HOPS/blob/external/Resources/default_list.txt) which comprised 356 pathogens though two entries were duplicates, hence bringing the total of pathogens to 354. We removed pathogens listed as Family or Genus level as no specific reference genome could be given. We also excluded substrains and incomplete genomes. We complemented this dataset by adding 27 horse-specific pathogens, bringing our final reference database to 247 pathogens (see Table S1 for a detailed list of pathogens considered including NCBI accession numbers and reasons for exclusion). We custom-built the database using HAYSTAC (Dimopoulos *et al*. 2022) by providing the NCBI Accession numbers, and ran the analysis using the abundance mode (--mode abundances) and a posterior probability for taxon assignment ≥0.75 (default parameter). Kraken2 was run with default parameters (Wood, Lu and Langmead 2019). We selected EHV-4 for downstream analyses based on the following HAYSTAC thresholds: Dirichlet read number >1000 which is required for reliable mapDamage analysis, an evenness of coverage <10, and a coverage of >0.05x. Fragmentation and nucleotide mis-incorporation plots were generated using mapDamage2 v2.0.8 (Jónsson *et al*. 2013), using default parameters (Fig S1A and S1B). The genome sequence was reconstructed using Bcftools (Danecek *et al*. 2021) mpileup, considering base phred quality scores of at least 30. The pileup sequence format (mpileup) provided the input for calling genotypes, assuming haploidy. Following a normalization step (norm -c x -d all), sites showing coverage below 3X or genotype phred qualities below 30 were disregarded. The resulting VCF file was indexed using tabix v1.7-41-g816a220 (Li *et al*. 2009) and the genome sequence was converted into fasta format using the vcf_to_fasta command from the PALEOMIX pipeline (Schubert *et al*. 2014).

### Phylogenetic Analyses

To investigate the phylogenetic placement of our ancient sequence, we first used NCBImeta v0.8.3 (Eaton 2020) to identify all complete EHV-4 genomes previously published in NCBI. It returned 141 entries of which 27 were complete unique genomes from four countries only, namely Australia, Germany, Ireland and Japan. We hence compiled a dataset of these 27 published EHV-4 genomes comprising the Irish strain NS80567 (Telford *et al*. 1998), 14 isolates from Australia (Vaz *et al*. 2016), eight isolates from Japan (Izume *et al*. 2017) and four isolates from Germany (Pavulraj *et al*. 2021) (Table S4). The sequence of the EHV-1 reference strain Ab4p (AY665713) was added as an outgroup (Telford *et al*. 1992). We conducted a multiple sequence alignment using MAFFT v7.453 (Katoh and Standley 2013) but the EHV-4 sequences were difficult to align against the EHV-1 reference sequence. Therefore, with EHV-1 and EHV-4 genomes both comprising 76 orthologous genes, we proceeded to the independent alignment of each ORF using MAFFT v7.453, followed by a manual verification on AliView v1.28 (Larsson 2014) and Seaview v4.7 (Gouy, Guindon and Gascuel 2010). We observed considerable disparity significantly caused by direct tandem repeats in the alignments of ORF 24, 47/44 and 71 (previously observed in (Izume *et al*. 2017)); these ORFs were thus removed from the alignment. The duplicated genes ORF64-ORF67 for EHV-1 and ORF64-ORF66 for EHV-4 were also discarded, as were the direct tandem repeat regions previously reported in the literature (Izume *et al*. 2017). Our procedure resulted in a multiple alignment of 104,418 orthologous sites comprising the sequences of 28 individual EHV-4 strains including our ancient sample, and one EHV-1 outgroup. We then generated a maximum likelihood tree using IQ-Tree v2.0.3 (Minh *et al*. 2020), with 1,000 bootstrap pseudoreplicates (option -b 1000) and the EHV-1 reference strain Ab4p as the outgroup (option -o). The integrated program ModelFinder (Kalyaanamoorthy *et al*. 2017) identified the best substitution model according to BIC as K3Pu+F+G4 (option -m TEST) and the EHV-1 reference strain Ab4p was rooted as an outgroup when visualising the resulting tree topology in iTOL v6.7 (Letunic and Bork 2021). Pairwise genetic distances were computed between pairs at tips from the maximum likelihood phylogenetic tree using its branch lengths through the cophenetic.phylo function in the R package ape (Paradis and Schliep 2019) (Table S5).

### Recombination analyses

Prior to conducting Bayesian phylogenetic analyses, we first tested for recombination in our dataset. We generated a multiple sequence alignment comprising our ancient and all 27 modern EHV-4 full genomes, using MAFFT v7.453 (Katoh and Standley 2013). We manually verified the alignment and removed the 17 direct tandem repeat regions identified in (Izume *et al*. 2017) using Seaview v4.7 (Gouy, Guindon and Gascuel 2010), which provided a final alignment of 28 genomes along 142,260 sites. Recombination was investigated using SplitsTree4 v4.18.3 (Huson and Bryant 2006) and RDP5 (Martin *et al*. 2021). Phylogenetic recombination networks were generated in SplitsTree4 using default parameters, and the statistical analyses of the recombination networks were performed using the pairwise homoplasy (Phi) test (Bruen, Philippe and Bryant 2006) as implemented in the software. Breakpoint identification was conducted using the seven most commonly used recombination detection methods implemented in RDP5 (v5.29), comprising RDP (Martin and Rybicki 2000), GENECONV (Padidam, Sawyer and Fauquet 1999), Chimaera (Posada and Crandall 2001), MaxChi (Maynard Smith 1992), BootScan (Martin *et al*. 2005), SiScan (Gibbs, Armstrong and Gibbs 2000) and 3Seq (Lam, Ratmann and Boni 2018), and with the following parameters: linear sequences, Bonferroni correction and highest acceptable *p*-value of 0.05. Recombination events identified by at least two of the seven methods, with *p*-values < 0.05 and good recombination scores (>0.4, (Martin *et al*. 2021)) were kept for downstream analyses to ensure as conservative an approach as possible. This resulted in 12 recombination events (Table S6) leading to a conserved region of 84,900 sites common to all sequences (positions 4,254 to 90,128 of the EHV-4 reference genome AF030027 (Telford *et al*. 1998) with direct tandem repeat regions removed (Izume *et al*. 2017)).

### Bayesian phylogenetics

To investigate the time to the Most Recent Common Ancestor (tMRCA) of all EHV-4 sequences, and the divergence time of phylogenetic subclades I and II, we constructed a time-calibrated Bayesian phylogeny based on the modern sequences and our ancient radiocarbon dated sample using BEAST v2.6.7 (Bouckaert *et al*. 2019), employing a birth-death skyline serial model and tip-dating. Tip dates were specified numerically as years Before Present (BP) with ‘Present’ set as ‘2017’ based on the most recently sampled isolates (Pavulraj *et al*. 2021). To ascertain the best-fitting clock prior between a strict clock and a relaxed clock log normal, we performed a model selection test using the Nested Sampling (NS) approach (Skilling 2006; Maturana Russel *et al*. 2019) available through the NS package (v.1.1.0) and based upon established guidelines from the dedicated Taming the Beast (Barido-Sottani et al., 2018) tutorial (https://taming-the-beast.org/tutorials/NS-tutorial/NS-tutorial.pdf) and the GitHub Nested Sampling FAQ (https://github.com/BEAST2-Dev/nested-sampling/wiki/FAQ). The Nested Sampling analysis was run with a Chain Length of 1E+12, a subchain length of 800,000 and 50 particles, which ensured the points generated at each iteration were independent and the marginal likelihoods were comparable between models (Strict clock: SD=1.76; Relaxed clock log normal: SD=1.83). The resulting log BF (ML Strict - ML Relaxed) is 4.28, which according to (Kass and Raftery 1995) provides overwhelming support for the Strict clock model.

For our non-recombined EHV4 dataset including UR17x29, K3Pu+F+I was identified as the best substitution model by ModelFinder (Kalyaanamoorthy *et al*. 2017; Minh *et al*. 2020) according to BIC. As this model was not available in BEAST v2.6.7, the next best model identified and available was GTR+F+I. All rates were set to the default value of 1.0 and estimated relative to the rate parameter of CT. Frequencies were set to Empirical. The Gamma Category Count was set to 0 and the Proportion of Invariant was estimated with a starting value of 0.9 based upon the ModelFinder estimate of 0.913. To reflect the epidemiology of EHV-4, we set the substitution rate to the default value of 1.0, and the clock rate to be estimated with a starting value of 1.0E-6 substitutions per site per year, which corresponds to the average overall substitution rate for double-stranded DNA viruses (Sanjuán *et al*. 2010). The lower and upper substitution rate boundaries were set to 1.0E-8 and 1.0E-5 respectively to reflect current knowledge on herpesviruses. For the effective reproductive number, Re, we used five equidistant intervals between the root and the last sample with a Log Normal (0,1.25) prior distribution resulting in a median of 1 and most weight being placed below 7.82 but still encompassing the upper boundary. Re was set to be estimated with a starting value of 3.0 and a lower boundary of 0 and an upper boundary of 15 based on known Re infection rate of EHV-1 considering they belong to the same subtype (Meade 2012). The infection period of EHV4 usually lasts 14 days but can extend up to 60 days (Pavulraj *et al*. 2021), meaning the become uninfectious rate falls between 6 and 26 per year. We set the value for M of the log normal distribution in real space to 26, with a standard deviation of 0.75. This renders our prior on the period of infectiousness primarily between 5.4 and 81 days. The sampling proportion was set to a beta distribution with A=1.0 and B=999.0, reflecting the very low sampling proportion. We specified the origin of our tree prior to follow a uniform distribution, and to be estimated with a starting value of 4,000 years with lower and upper boundaries of 3,871 and Infinity respectively based on the age of our oldest sample. Our ucldMean prior followed a uniform distribution and was based on the clock rate input. Finally, the MCMC chain length was run for a total of 200 million iterations for the relaxed clock log normal model, and for a total of 100 million iterations for the strict clock model. Convergence was visually confirmed by ensuring the effective sample sizes (ESS) estimation for all parameters was >200, disregarding the first 10% iterations as burn-in, using Tracer v.1.7.2 (Rambaut *et al*. 2018). Maximum clade credibility trees with Median node heights and following a 10% burn-in were generated using Tree Annotator v2.6.6 as implemented in BEAST v2.6.7 (Bouckaert *et al*. 2019) and visualised with FigTree v1.4.4. (http://tree.bio.ed.ac.uk/software/figtree/). R_0_ estimates (R_e_) were plotted using R-package *bdskytools* (https://github.com/laduplessis/bdskytools).

For our non-recombined modern dataset, TIM+F+I was identified as the best substitution model by ModelFinder (Kalyaanamoorthy *et al*. 2017; Minh *et al*. 2020) according to BIC. As this model was unavailable in BEAST v2.6.7; the next best model identified and available was GTR+F+I. All parameters were identical to our non-recombined ancient dataset, with the following two exceptions: i) the Proportion of Invariant sites was estimated with a starting value of 0.89 based upon the ModelFinder estimate, and ii) our origin prior followed a uniform distribution and was estimated with a starting value of 167 (1,850 CE) with lower and upper boundaries of 145 (1,872) and Infinity respectively based on historical reviews. Indeed, the earliest record of a mild form of a non-zoonotic equine influenza lacking abortions, thus resembling EHV-4, is the American equine epidemic of 1872-1873 (Williams 1924). Hence, this outbreak served as a guided hypothesis for our origins prior. Effective sample size (ESS) was >200 for all parameters with the first 10% iterations disregarded as burn-in using Tracer v.1.7.2 (Rambaut *et al*. 2018), thus confirming convergence. Maximum clade credibility trees were generated and visualised through BEAST v2.6.7 (Bouckaert *et al*. 2019) and FigTree v1.4.4 respectively, as described above.

### Positive Selection

Positive selection between our ancient and published modern sequences was tested using the CODEML branch-site model implemented in the PAML v.4.9 package (Yang 2007). The branch-site model compares the branch-site model A with the corresponding null model which fixes *ω*_2_=1, and has a likelihood ratio test of 1 degree of freedom (model=2; NSsites=2; fix_omega=0/1; omega =1.5/1). We used our previously generated phylogenetic tree from IQ-Tree based upon our ORF-concatenated alignment comprising the 28 EHV-4 sequences and the EHV-1 reference strain Ab4p as an outgroup to constrain the phylogenetic placement of our ancient sequence. The EHV-1 sequence was then removed, and the tree was saved in a Newick format. We then manually assigned the UR17x29 ancient sequence as a background branch, and the modern sequences as foreground branches (from Branch M onwards, Fig 1B). The codon frequencies were calculated from the average nucleotide frequencies. Testing for positive selection in subclades I and II followed the same protocol as above: positive selection in subclade I was tested by assigning the UR17x29 sequence and subclade II sequences as background branches and subclade I sequences as foreground branches, and vice-versa for positive selection in subclade II. We used the χ_1_ ^2^ with critical value 3.84 (df=1).

### Protein Structure prediction

Protein structure prediction for the eight versions of tegument protein UL14 observed in modern strains and UR17x29 was performed using I-TASSER (Yang *et al*. 2015). Amino acid sequences representing UL14 from ancient and modern populations were submitted to I-TASSER for automated protein structure prediction using default parameters. Unassigned nucleotides from the ancient genome were assumed to have the same nucleotides as the reference genome for I-TASSER submission. I-TASSER generates full-length atomic structural models from multiple threading alignments and predicts the functions from known protein function databases based on sequence and structure profile comparisons. I-TASSER generated five models, out of which the best model with highest C-score was chosen for structural analysis. Pdb files were visualised using RCSB PDB Mol* viewer (Sehnal *et al*. 2021). Lack of protein homolog structures of UL14 confirmed by X-ray crystallography or otherwise from other equine herpesviruses, makes it impossible to go beyond primary structure prediction.

## Acknowledgements

This project has received funding from the CNRS and University Paul Sabatier (AnimalFarm IRP), and the European Research Council (ERC) under the European Union’s Horizon 2020 research and innovation program (grant agreement 681605-PEGASUS). O.L. and K.M. were supported by the European Union’s Horizon 2020 research and innovation programme under the Marie Sklodowska-Curie grant agreements no. 895107 and no. 897931, respectively.

## Data Availability

The data is available on NCBI under BioSample: SAMEA9533417 with further details available at https://doi.org/10.1038/s41586-021-04018-9.

