## Supplementary Information for "Equine herpesvirus 4 infected domestic horses associated with Sintashta spoke-wheeled chariots around 4,000 years ago"

**Supplementary Figure S1:** Post-mortem DNA damage patterns for UR17x29 A) horse (host) and B) EHV-4 following USER treatment

**Supplementary Figure S2:** Phylogenetic relationships based on non-recombined data.

**Supplementary Figure S3:** Time-scaled EHV-4 evolutionary history based on modern dataset only.

**Supplementary Table S1:** Pathogen reference database for HAYSTAC and Kraken2 analyses

**Supplementary Table S2a-e**: HAYSTAC results for libraries ERR6466108- ERR6466112

**Supplementary Table S3a-e**: Kraken2 results for libraries ERR6466108- ERR6466112

**Supplementary Table S4:** EHV-1 and EHV-4 sequences used in this study with corresponding Genbank accession number, location and sampling date of isolates

**Supplementary Table S5:** Pairwise genetic distances computed from the maximum likelihood phylogenetic tree based on branch length

**Supplementary Table S6:** Whole genome recombination events and detection of recombination breakpoints

### A. UR17x29 – Horse (host)

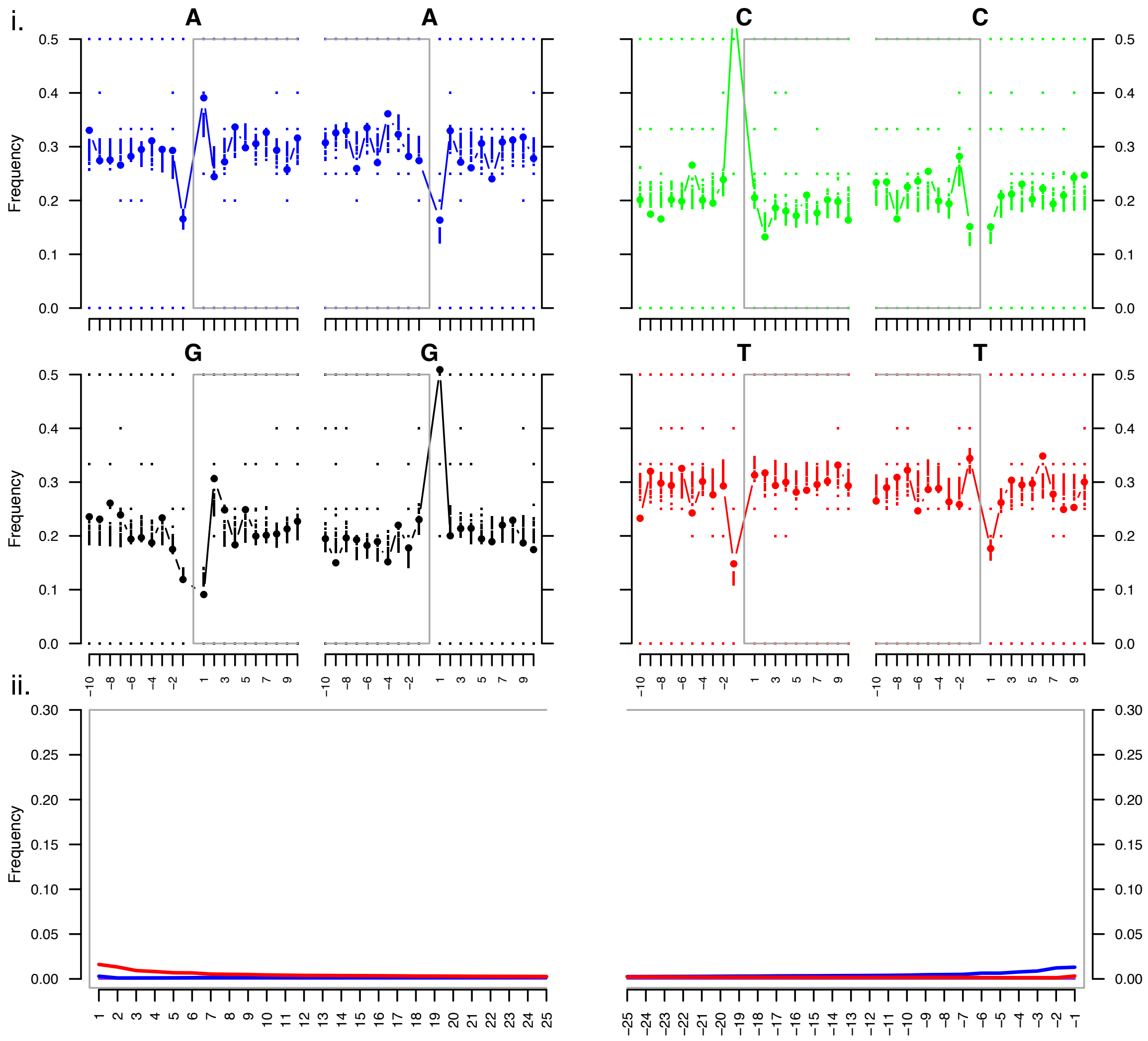

**B. UR17x29 – EHV-4**

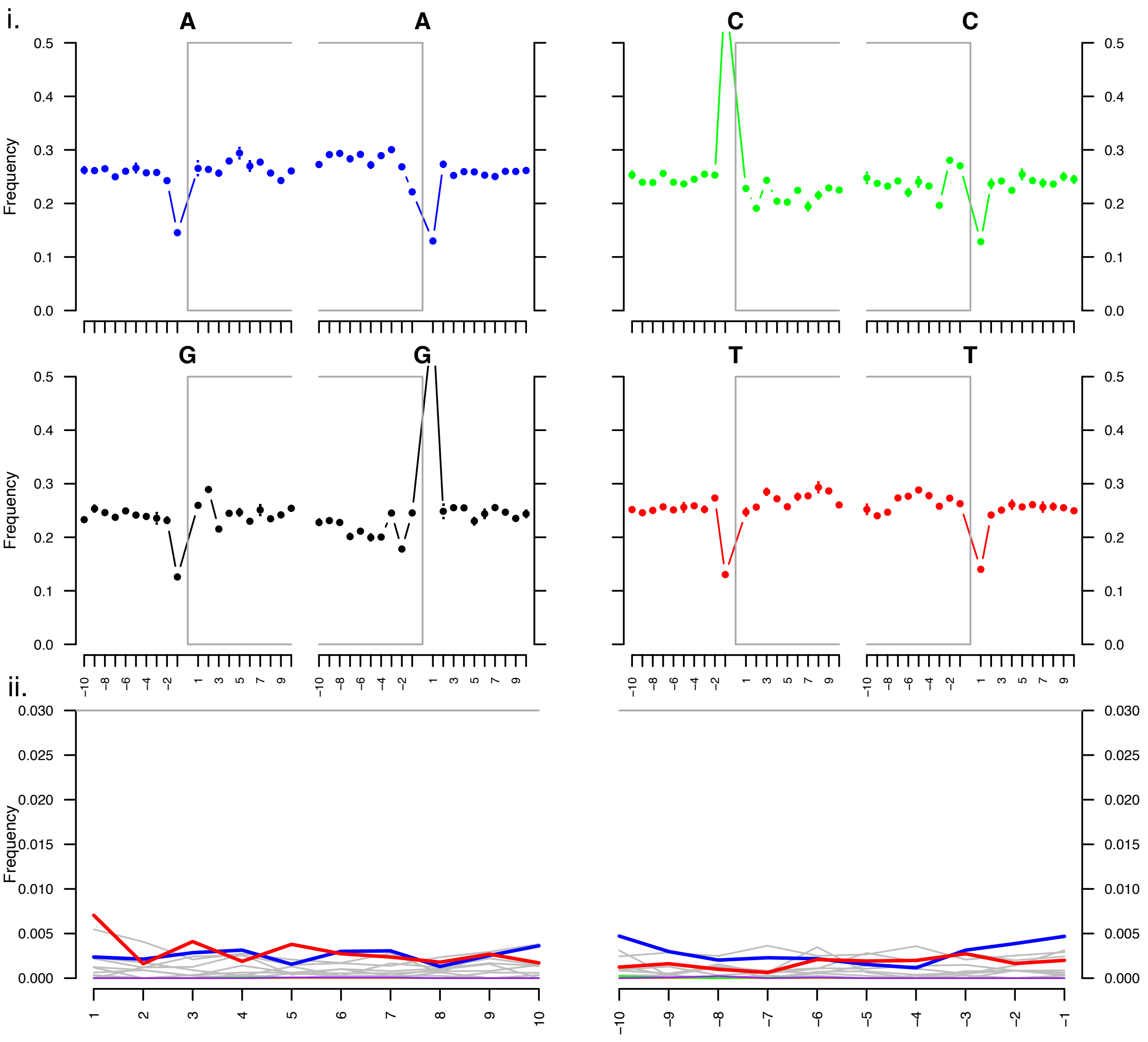


**Supplementary Figure S1: Post-mortem DNA damage patterns for UR17x29 A) horse (host) and B) EHV-4 following USER treatment.** i. Base composition of the first 10 nucleotides sequenced (1 to 10) and the 10 nucleotides preceding the aligned genomic region (-10 to -1). Nucleotide positions located within reads are indicated by the grey frame. ii. Nucleotide mis-incorporations. C→T and G→A nucleotide mis-incorporations are reported in red and blue, respectively.

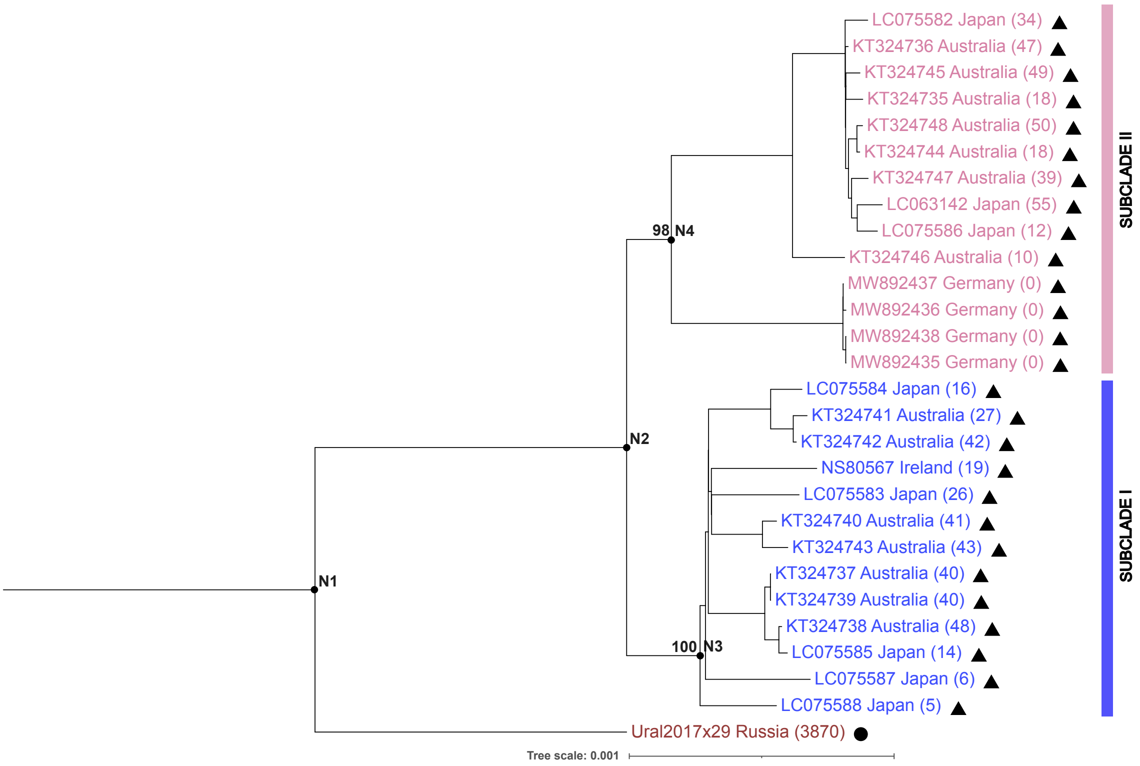


**Supplementary Figure S2: Phylogenetic relationships based on non-recombined data.** Maximum-Likelihood tree and node bootstrap support based on the non-recombined region spanning positions 4,254-90,128 of the EHV-4 reference genome (AF030027, [2]) and without tandem repeats. Sample names follow Figure 1.

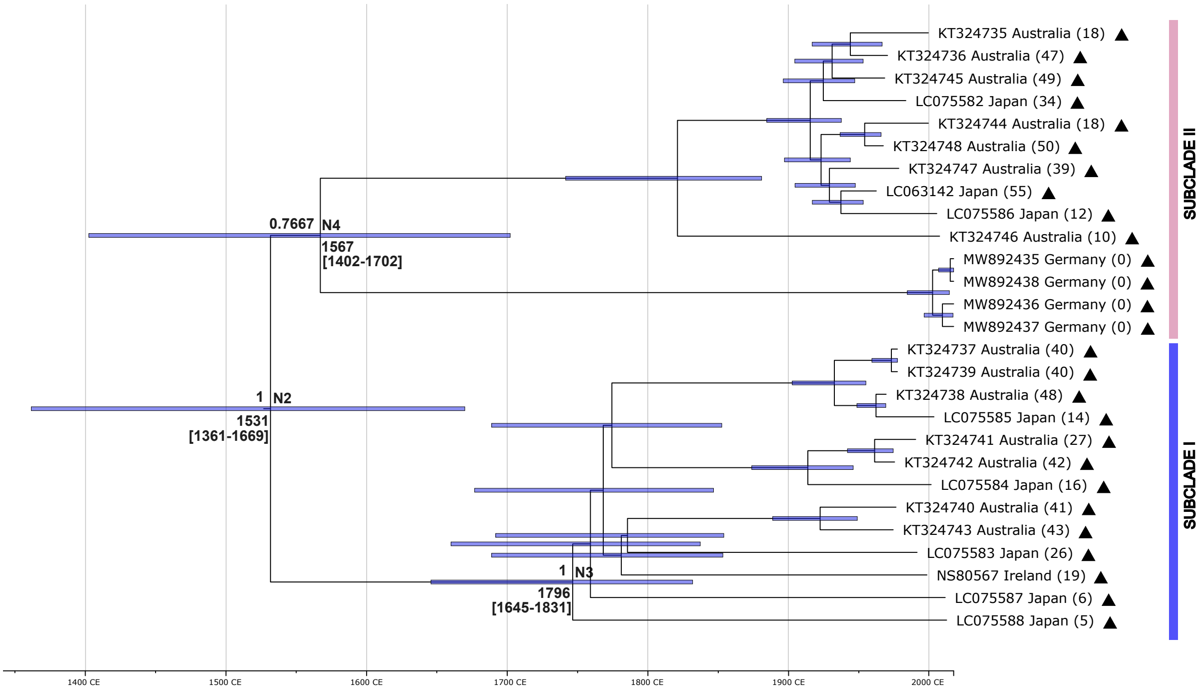


**Supplementary Figure S3: Time-scaled EHV-4 evolutionary history based on modern dataset only.** Bayesian phylogeny under the strict clock and birth-death skyline serial models based on the 27 previously published modern references. Node posterior supports are provided above nodes to the left, whilst median estimates of divergence times are provided below nodes, together with the 95% confidence interval in square brackets [Height_95%_HPD]. Sequence labels comprise Genbank accession numbers, sampling locations, and, in parentheses, sampling dates formatted as years Before Present (BP), with the 'Present' set to '2017' based on the most recently sampled isolates. The dates were used for tip-dating calibration. The symbols refer to Figure 1A.

**Table S1:** Reference database for HAYSTAC and Kraken2 analyses. The database is derived from the HOPS default pathogens list (n=354, https://github.com/rhuebler/HOPS/blob/external/Resources/default_list.txt) from which selected pathogens were removed (n=134, coloured in grey, see 'Reason for exclusion'). The resulting database was complemented with 27 horse-specific pathogens (marked by *), resulting in a final database comprising a total of 247 pathogens.

| Scientific_Name | Included in custom-built pathogen database? | Accession or Reason for exclusion |
|---|---|---|
| Actinobacillus_equuli_equuli* | Yes | GCF_000801145.1 / NZ_CP007715.1 |
| African_swine_fever_virus | Yes | GCF_003032865.1 |
| Aspergillus_fumigatus | Yes | GCF_000002655.1 |
| Bacillus_anthracis | Yes | GCF_000008445.1 / NC_007530.2 |
| Bacillus_cereus | Yes | GCF_002220285.1 / NZ_CP017060.1 |
| Bartonella_bacilliformis | Yes | GCF_000015445.1 / NC_008783.1 |
| Bartonella_henselae | Yes | GCF_000612965.1 / NZ_HG965802.1 |
| Bartonella_quintana | Yes | GCF_021869585.1 / NZ_CP091504.1 |
| BK_virus_strain_AS | No | Incomplete genome (protein-coding sequences only) |
| BK_virus_strain_GS | No | Incomplete genome (307bp origin-proximal region) |
| BK_virus_variant_RF | No | No sequences available |
| Blastomyces_dermatitidis | Yes | GCF_000003525.1 |
| Bordetella | No | Genus level - no specific reference genome |
| Bordetella_pertussis | Yes | GCF_004008975.1 /NZ_CP025371.1 |
| Bordetella_petrii | Yes | GCF_000067205.1 / NC_010170.1 |
| Borreliaceae | No | Family level - no specific reference genome |
| Borrelia | No | Genus level - no specific reference genome |
| Borreliella_afzelii | Yes | GCF_018141305.1 / NZ_CP042238.1 |
| Borreliella_burgdorferi | Yes | GCF_000181575.2 |
| Borreliella_garinii | Yes | GCF_000196215.1 / NC_006156.1 |
| Borrelia_recurrentis | Yes | GCF_000019705.1 |
| Brucellaceae | No | Family level - no specific reference genome |
| Brucella | No | Genus level - no specific reference genome |
| Brucella_abortus | Yes | GCF_000054005.1 |
| Brucella_melitensis | Yes | GCF_000007125.1 |
| Brucella_microti | Yes | GCF_000022745.1 |
| Brucella_ovis | Yes | GCF_000016845.1 |
| Brucella_suis | Yes | GCF_000007505.1 |
| Burkholderia_cepacia | Yes | GCF_009586235.1 |
| Burkholderia_mallei | Yes | GCF_002346025.1 |
| Burkholderia_pseudomallei | Yes | GCF_000756125.1 |
| Chlamydia_abortus | Yes | GCF_900416725.2 |
| Chlamydia_caviae | Yes | GCF_000007605.1 / NC_003361.3 |
| Chlamydia_felis | Yes | GCF_000009945.1 / NC_007899.1 |
| Chlamydia_muridarum | Yes | GCF_021065065.1 /NZ_CP042790.1 |
| Chlamydia_pecorum | Yes | GCF_020459125.1 |
| Chlamydia_pneumoniae | Yes | GCF_000024145.1 / NC_017285.1 |
| Chlamydia_psittaci | Yes | GCF_000204255.1 /NC_017287.1 |
| Chlamydia_trachomatis | Yes | GCF_000008725.1 / NC_000117.1 |
| Clostridium | No | Genus level - no specific reference genome |
| Clostridium_botulinum | Yes | GCF_000063585.1 |
| Clostridium_botulinum_BKT015925 | No | Excluded as substrain |
| Clostridium_botulinum_E3_str._Alaska_E43 | Yes | GCF_000020285.1 |
| Clostridium_perfringens* | Yes | GCF_020138775.1 / NZ_CP075979.1 |
| Clostridium_sporogenes | Yes | GCF_020450145.1 |
| Clostridium_tetani | Yes | GCF_000762305.1 / NZ_JRGJ01000001.1 |
| Clostridium_tetani_E88 | Yes | GCF_000007625.1 |
| Coccidioides_immitis | Yes | GCF_000149335.2 |
| Coccidioides_posadasii | Yes | GCF_000151335.2 |
| Corynebacterium_diphtheriae | Yes | GCF_002843135.1 |
| Corynebacterium_pseudotuberculosis* | Yes | GCF_000259155.4 / NC_017730.4 |
| Cowpox-Vaccinia_virus | No | Incomplete genome (protein-coding sequences only) |
| Cowpox_virus | Yes | GCF_000839185.1 |
| Coxiella_burnetii | Yes | NC_002971.4 |
| Cryptococcus_gattii | Yes | GCF_000185945.1 |
| Cryptococcus_neoformans | Yes | GCF_000091045.1 |
| Cynomolgus_Epstein-Barr_Virus_A4 | No | Incomplete genome (partial coding sequence) |
| Cynomolgus_Epstein-Barr_Virus_Si-IIA | No | Excluded as substrain |
| Cynomolgus_Epstein-Barr_Virus_TsB-B6 | No | Excluded as substrain |
| Eastern_equine_encephalitis* | Yes | KU840375.1 |
| Equid_alphaherpesvirus_1* | Yes | GCF_000844025.1 |
| Equid_alphaherpesvirus_3* | Yes | GCF_000921595.1 |
| Equid_alphaherpesvirus_8* | Yes | GCF_000894575.1 |
| Equid_alphaherpesvirus_9* | Yes | GCF_000883455.1 |
| Equid_alphaherpesvirus_4* | Yes | GCF_000846345.1 / NC_001844.1 |
| Equid_gammaherpesvirus_2* | Yes | GCF_000843985.2 |
| Equid_gammaherpesvirus_5* | Yes | GCF_000929435.1 |
| Equine_infectious_anemia* | Yes | NC_001450.1 |
| Equine_arteritis_virus* | Yes | GCF_000860865.1 / NC_002532.2 |
| Enterobius_vermicularis | No | Not assembled |
| Epstein-barr_virus_strain_ag876 | No | Excluded as substrain |
| Epstein-barr_virus_strain_p3hr-1 | No | Excluded as substrain |
| Epstein_Barr_type2* | Yes | NC_009334.1 |
| Escherichia_coli | Yes | GCF_000157115.2 / NZ_KQ235739.1 |
| Escherichia_coli_O111 | Yes | GCF_003293985.1 / NZ_QMIA01000099.1 |
| Escherichia_coli_O157-H7 | Yes | GCF_000008865.2 |
| Escherichia_coli_O26 | Yes | GCF_002734545.1 / NZ_BEKS01000001.1 |
| Fusobacterium | No | Genus level - no specific reference genome |
| Fusobacterium_nucleatum | Yes | GCF_003019295.1 |
| Francisella_tularensis | Yes | GCF_000833355.1 |
| Haemophilus_influenzae | Yes | GCF_000931575.1 |
| Haemophilus_influenzae_Rd_KW20 | No | RefSeq surppressed |
| Helicobacter_pylori | Yes | GCF_017821535.1 |
| Hepatitis_B_virus | Yes | GCF_000861825.2 |
| Recombinant_Hepatitis_C_virus_3a-2a_(SA1/JFH1) | No | Incomplete genome (protein-coding sequences only) |
| Hepatitis_delta_virus | Yes | NC_001653.2 |
| Hepatitis_E_virus | Yes | NC_038504.1 |
| Herpes_simplex_virus_1-strain_R-15 | No | Incomplete genome (partial ORFs) |
| Herpes_simplex_virus_type1-strain_17 | No | Excluded as substrain |
| Herpes_simplex_virus_type1-strain_A44 | No | Excluded as substrain |
| Herpes_simplex_virus_type1-strain_Angelotti | No | Excluded as substrain |
| Herpes_simplex_virus_type1-strain_CL101 | No | Excluded as substrain |
| Herpes_simplex_virus_type1-strain_CVG-2 | No | Excluded as substrain |
| Herpes_simplex_virus_type1-strain_F | No | Excluded as substrain |
| Herpes_simplex_virus_type1-strain_HFEM | No | Excluded as substrain |
| Herpes_simplex_virus_type1-strain_HZT | No | Excluded as substrain |
| Herpes_simplex_virus_type1-strain_MGH-10 | No | Excluded as substrain |
| Herpes_simplex_virus_type1-strain_MP | No | Excluded as substrain |
| Herpes_simplex_virus_type1-strain_Patton | No | Excluded as substrain |
| Herpes_simplex_virus_type1-strain_R19 | No | Excluded as substrain |
| Herpes_simplex_virus_type1-strain_RH2 | No | Excluded as substrain |
| Herpes_simplex_virus_type1-strain_SC16 | No | Excluded as substrain |
| Herpes_simplex_virus_unknown_type | No | Excluded as substrain |
| Horsepox* | Yes | KY349117.1 |
| Human_herpesvirus_1 | Yes | NC_001806.2 |
| Human_herpesvirus_6B | Yes | GCF_000846365.1 |
| Histoplasma_capsulatum | Yes | GCF_000150115.1 |
| Human_adenovirus_1 | Yes | GCF_000858645.1 |
| Human_adenovirus_11 | No | Excluded as substrain |
| Human_adenovirus_11+34 | No | Excluded as substrain |
| Human_adenovirus_11a | No | Excluded as substrain |
| Human_adenovirus_11p | Yes | AY163756.1 |
| Human_adenovirus_12 | Yes | X73487.1 |
| Human_adenovirus_14 | Yes | MG642756.1 |
| Human_adenovirus_14a | No | Excluded as substrain |
| Human_adenovirus_14p | No | Excluded as substrain |
| Human_adenovirus_14p1 | No | Excluded as substrain |
| Human_adenovirus_15 | Yes | AB562586.1 |
| Human_adenovirus_15-H9 | No | Excluded as substrain |
| Human_adenovirus_16 | Yes | AY601636.1 |
| Human_adenovirus_16p | No | Excluded as substrain |
| Human_adenovirus_17 | Yes | HQ910407.1 |

| | | |
|---|---|---|
| Human_adenovirus_18 | Yes | GU191019.1 |
| Human_adenovirus_19 | No | Excluded as substrain |
| Human_adenovirus_19a | Yes | AB448772.1 |
| Human_adenovirus_19C | No | Excluded as substrain |
| Human_adenovirus_19p | No | Excluded as substrain |
| Human_adenovirus_1_strain_Adenoid_71 | No | Excluded as substrain |
| Human_adenovirus_2 | Yes | GCF_000859465.1 |
| Human_adenovirus_20 | Yes | JN226749.1 |
| Human_adenovirus_21 | Yes | KF528688.1 |
| Human_adenovirus_21a | No | Excluded as substrain |
| Human_adenovirus_22 | Yes | FJ404771.1 |
| Human_adenovirus_22H8 | No | Excluded as substrain |
| Human_adenovirus_23 | Yes | JN226750.1 |
| Human_adenovirus_24 | Yes | JN226751.1 |
| Human_adenovirus_25 | Yes | JN226752.1 |
| Human_adenovirus_26 | Yes | GN062506.1 |
| Human_adenovirus_27 | Yes | JN226753.1 |
| Human_adenovirus_28 | Yes | FJ824826.1 |
| Human_adenovirus_29 | Yes | JN226754.1 |
| Human_adenovirus_30 | Yes | JN226755.1 |
| Human_adenovirus_31 | Yes | MZ983583.1 |
| Human_adenovirus_3+11p | No | Excluded as substrain |
| Human_adenovirus_32 | Yes | JN226756.1 |
| Human_adenovirus_33 | Yes | JN226758.1 |
| Human_adenovirus_34 | Yes | AY737797.1 |
| Human_adenovirus_34a | No | Excluded as substrain |
| Human_adenovirus_35 | Yes | GCF_000857885.1 |
| Human_adenovirus_35+11 | No | Excluded as substrain |
| Human_adenovirus_36 | Yes | GQ384080.1 |
| Human_adenovirus_3+7 | Yes | JN860679.1 |
| Human_adenovirus_38 | Yes | JN226759.1 |
| Human_adenovirus_38H44 | No | Excluded as substrain |
| Human_adenovirus_39 | Yes | JN226760.1 |
| Human_adenovirus_3p | No | Excluded as substrain |
| Human_adenovirus_40 | Yes | KU162869.1 |
| Human_adenovirus_41 | Yes | ON532818.1 |
| Human_adenovirus_41_Wisconsin | No | Excluded as substrain |
| Human_adenovirus_42 | Yes | JN226761.1 |
| Human_adenovirus_43 | Yes | KC529648.1 |
| Human_adenovirus_44 | Yes | JN226763.1 |
| Human_adenovirus_45 | Yes | JN226764.1 |
| Human_adenovirus_46 | Yes | AY875648.1 |
| Human_adenovirus_47 | Yes | JN226757.1 |
| Human_adenovirus_48 | Yes | EF153473.1 |
| Human_adenovirus_49 | Yes | DQ393829.1 |
| Human_adenovirus_4a | Yes | AP014849.1 |
| Human_adenovirus_5 | Yes | GCF_000857865.1 |
| Human_adenovirus_50 | No | Excluded as substrain |
| Human_adenovirus_50_prototype_strain_Wan | Yes | AY737798.1 |
| Human_adenovirus_51 | Yes | JN226765.1 |
| Human_adenovirus_52 | Yes | DQ923122.2 |
| Human_adenovirus_53 | Yes | AB605243.1 |
| Human_adenovirus_54 | Yes | GCF_000885675.1 |
| Human_adenovirus_55 | Yes | MT513753.1 |
| Human_adenovirus_56 | Yes | HM770721.2 |
| Human_adenovirus_57 | No | Incomplete genome (protein-coding sequences only) |
| Human_adenovirus_58 | Yes | HQ883276.1 |
| Human_adenovirus_59_MSJ-2011 | No | No sequences available |
| Human_adenovirus_6 | Yes | LC068712.1 |
| Human_adenovirus_60a | Yes | JN162672.1 |
| Human_adenovirus_61 | Yes | JF964962.1 |
| Human_adenovirus_62 | Yes | JN162671.1 |
| Human_adenovirus_63 | Yes | JN935766.1 |
| Human_adenovirus_64 | Yes | JQ326207.1 |
| Human_adenovirus_65 | Yes | AP012285.1 |
| Human_adenovirus_66 | Yes | JN860676.1 |
| Human_adenovirus_67 | Yes | AP012302.1 |
| Human_adenovirus_68 | Yes | JN860678.1 |
| Human_adenovirus_69 | Yes | JN226748.1 |
| Human_adenovirus_7 | Yes | GCF_000859485.1 |
| Human_adenovirus_71 | Yes | KF268207.1 |
| Human_adenovirus_7428 | No | Incomplete genome (partial coding sequence) |
| Human_adenovirus_7620 | No | Incomplete genome (partial coding sequence) |
| Human_adenovirus_7961 | No | Incomplete genome (partial coding sequence) |
| Human_adenovirus_7a | No | Incomplete genome (partial coding sequence) |
| Human_adenovirus_7d | No | Excluded as substrain |
| Human_adenovirus_7d2 | No | Excluded as substrain |
| Human_adenovirus_7h | No | Excluded as substrain |
| Human_adenovirus_7l | No | Excluded as substrain |
| Human_adenovirus_7m | No | Excluded as substrain |
| Human_adenovirus_7p | No | Excluded as substrain |
| Human_adenovirus_8E | No | Incomplete genome (partial coding sequence) |
| Human_adenovirus_9042 | No | Incomplete genome (partial coding sequence) |
| Human_adenovirus_9052 | No | Incomplete genome (partial coding sequence) |
| Human_adenovirus_9431 | No | No sequences available |
| Human_adenovirus_A02-2B-2 | No | No sequences available |
| Human_adenovirus_A03-2B-1 | No | Incomplete genome (partial coding sequence) |
| Human_adenovirus_AdV-Berlin | No | No sequences available |
| Human_adenovirus_AdV-Berlin003-02-DE | No | Incomplete genome (partial coding sequence) |
| Human_adenovirus_AdV-Berlin004-02-DE | No | Incomplete genome (partial coding sequence) |
| Human_adenovirus_AdV-Berlin131-01-DE | No | Incomplete genome (partial coding sequence) |
| Human_adenovirus_AdV-Berlin524-01-DE | No | Incomplete genome (partial coding sequence) |
| Human_adenovirus_AdV-Berlin604-01-DE | No | Incomplete genome (partial coding sequence) |
| Human_adenovirus_AdV-Berlin619-01-DE | No | Incomplete genome (partial coding sequence) |
| Human_adenovirus_AdV-Berlin643-01-DE | No | Incomplete genome (partial coding sequence) |
| Human_adenovirus_AdV-Berlin682-01-DE | No | Incomplete genome (partial coding sequence) |
| Human_adenovirus_AdV-Berlin718-01-DE | No | Incomplete genome (partial coding sequence) |
| Human_adenovirus_AdV-Berlin749-01-DE | No | Incomplete genome (partial coding sequence) |
| Human_adenovirus_AdV-Berlin769-01-DE | No | Incomplete genome (partial coding sequence) |
| Human_adenovirus_AdV-Berlin859-01-DE | No | Incomplete genome (partial coding sequence) |
| Human_adenovirus_ATCC_VR-1501 | No | Incomplete genome (partial coding sequence) |
| Human_adenovirus_ATCC_VR-1502 | No | Incomplete genome (partial coding sequence) |
| Human_adenovirus_B1 | Yes | NC_011203.1 |
| Human_adenovirus_B2 | Yes | JB741664.1 |
| Human_adenovirus_B3 | Yes | OQ518281.1 |
| Human_adenovirus_Chiba_E086-2012 | No | No sequences available |
| Human_adenovirus_D10 | Yes | JN226746.1 |
| Human_adenovirus_D13 | Yes | JN226747.1 |
| Human_adenovirus_D15H9 | No | Incomplete genome (partial gene) |
| Human_adenovirus_D17H29 | No | Incomplete genome (partial gene) |
| Human_adenovirus_D19H30 | No | Incomplete genome (partial gene) |
| Human_adenovirus_D30H44 | No | Incomplete genome (partial gene) |
| Human_adenovirus_D37 | Yes | LC215440.1 |
| Human_adenovirus_D37H13 | No | Excluded as substrain |
| Human_adenovirus_D37H17 | No | Excluded as substrain |
| Human_adenovirus_D46H13 | No | Excluded as substrain |
| Human_adenovirus_D8 | Yes | AB897885.1 |
| Human_adenovirus_D9 | Yes | AJ854486.1 |
| Human_adenovirus_E4 | Yes | OK323254.1 |
| Human_adenovirus_H02-2B-2 | No | Incomplete genome (partial coding sequence) |
| Human_adenovirus_H02-2B-3 | No | Incomplete genome (partial coding sequence) |
| Human_adenovirus_H03-2B-1 | No | Incomplete genome (partial coding sequence) |
| Human_adenovirus_H08-2B-2 | No | Incomplete genome (partial coding sequence) |
| Human_adenovirus_I02-2B-2 | No | Incomplete genome (partial coding sequence) |
| Human_adenovirus_I05-2B-2 | No | Incomplete genome (partial coding sequence) |
| Human_adenovirus_isolate_MY7-1 | No | Incomplete genome (partial gene) |
| Human_adenovirus_isolate_MY8-1 | No | Incomplete genome (partial gene) |
| Human_adenovirus_isolate_SGH2289 | No | Incomplete genome (partial gene) |
| Human_adenovirus_isolate_SGH5253 | No | Incomplete genome (partial gene) |
| Human_adenovirus_LZY153B | No | Incomplete genome (partial coding sequence) |
| Human_adenovirus_MKJ-Chennai-2010 | No | Incomplete genome (partial gene) |

| | | |
|---|---|---|
| Human_adenovirus_RKI-2797-04 | No | Incomplete genome (partial coding sequence) |
| Human_adenovirus_S02-2B-1 | No | Incomplete genome (partial coding sequence) |
| Human_adenovirus_S03-2B-1 | No | Incomplete genome (partial coding sequence) |
| Human_adenovirus_S03-2B-2 | No | Incomplete genome (partial coding sequence) |
| Human_adenovirus_S03-2B-3 | No | Incomplete genome (partial coding sequence) |
| Human_adenovirus_S05-2B-2 | No | Incomplete genome (partial coding sequence) |
| Human_adenovirus_S06-2B-3 | No | Incomplete genome (partial coding sequence) |
| Human_adenovirus_S07-2B-2 | No | Incomplete genome (partial coding sequence) |
| Human_adenovirus_S08-2B-3 | No | Incomplete genome (partial coding sequence) |
| Human_adenovirus_S10-1-6 | No | Incomplete genome (partial coding sequence) |
| Human_adenovirus_type_35p | No | Incomplete genome (protein-coding sequence only) |
| Human_alphaherpesvirus_1* | Yes | GCF_000859985.2 |
| Human_hepatitis_A_virus | Yes | LC373510.1 |
| Human_herpesvirus_3_strain_Dumas | Yes | X04370.1 |
| Human_herpesvirus_3_VZV-32 | No | Incomplete genome (partial ORFs) |
| Human_herpesvirus_3_strain_Oka_vaccine | No | No sequences available |
| Human_gammaherpesvirus_4 | Yes | GCF_002402265.1 |
| Human_mastadenovirus_C | Yes | NC_001405.1 |
| JC_polyomavirus | Yes | GCF_000863805.1 |
| JC_virus_x_BK_virus | No | Incomplete genome (regulatory region x3) |
| Klebsiella_oxytoca | Yes | GCF_003812925.1 |
| Klebsiella_pneumoniae_subsp._pneumoniae | Yes | GCF_000240185.1 |
| Klebsiella_pneumoniae_subsp._rhinoscleromatis | Yes | GCF_022809675.1 / NZ_CP094451.1 |
| Legionella_pneumophila | Yes | GCF_001753085.1 |
| Leptospira_alexanderi | Yes | GCF_002009845.1 |
| Leptospira_alstonii | Yes | GCF_001569395.1 |
| Leptospira_borgpetersenii | Yes | GCF_003516145.1 |
| Leptospira_broomii | Yes | GCF_000243715.2 |
| Leptospira_fainei | Yes | GCF_000306235.2 |
| Leptospira_inadai | Yes | GCF_000243675.2 |
| Leptospira_interrogans | Yes | GCF_002073495.2 |
| Leptospira_kirschneri | Yes | GCF_000243695.2 |
| Leptospira_kmetyi | Yes | GCF_003722295.1 |
| Leptospira_licerasiae | Yes | GCF_000526875.1 |
| Leptospira_noguchii | Yes | GCF_000306255.2 |
| Leptospira_santarosai | Yes | GCF_000313175.2 |
| Leptospira_weilii | Yes | GCF_006874765.1 |
| Leptospira_wolffii | Yes | GCF_004770635.1 |
| Marseilleviridae_tokyovirus* | Yes | AP017398.1 |
| Methanobrevibacter_oralis | Yes | GCF_912073625.1 |
| Moraxella_catarrhalis | Yes | GCF_002080125.1 |
| Mycobacteroides_abscessus | Yes | GCF_004028015.1 |
| Mycobacterium_africanum | Yes | GCF_001593225.1 / NZ_CP014617.1 |
| Mycobacterium_avium | Yes | GCF_022175585.1 |
| Mycobacterium_bovis | Yes | GCF_005156105.1 / NZ_CP039850.1 |
| Mycobacterium_canettii | Yes | GCF_000253375.1 |
| Mycobacterium_colombiense | Yes | GCF_000222105.3 |
| Mycobacterium_intracellulare | Yes | GCF_020736005.1 |
| Mycobacterium_kansasii | Yes | GCF_000157895.3 |
| Mycobacterium_leprae | Yes | GCF_003253775.1 |
| Mycobacterium_tuberculosis | Yes | GCF_000195955.2 / NC_000962.3 |
| Mycoplasma_pneumoniae | Yes | GCF_000733995.1 / NZ_CP008895.1 |
| Neisseria | No | Genus level - no specific reference genome |
| Neisseria_gonorrhoeae | Yes | GCF_013030075.1 |
| Neisseria_meningitidis | Yes | GCF_008330805.1 |
| Orthohepadnavirus | No | Genus level - no specific reference genome |
| Parvimonas_micra | Yes | GCF_003454775.1 |
| Peptostreptococcus_anaerobius | Yes | GCF_000178095.1 |
| Plasmodium_falciparum | Yes | GCF_000002765.5 |
| Plasmodium_vivax | Yes | GCF_000002415.2 |
| Pneumocystis_carinii | Yes | GCF_001477545.1 |
| Pneumocystis_jirovecii | Yes | GCF_001477535.1 |
| Porphyromonas_gingivalis | Yes | GCF_000010505.1 |
| Porphyromonas_gingivalis_W83 | Yes | GCF_000007585.1 |
| Prescottella_equi* | Yes | GCF_003013675.1 / NZ_CP027793.1 |
| Rickettsia_africae | Yes | GCF_000023005.1 |
| Rickettsia_akari | Yes | GCF_000018205.1 |
| Rickettsia_conorii | Yes | GCF_000007025.1 |
| Rickettsia_felis | Yes | GCF_000804505.1 |
| Rickettsia_japonica | Yes | GCF_002356735.1 |
| Rickettsia_prowazekii | Yes | GCF_000277165.1 |
| Rickettsia_rickettsii | Yes | GCF_000018225.1 |
| Rickettsia_sibirica | Yes | GCF_000166935.1 |
| Rickettsia_typhi | Yes | GCF_000277285.1 |
| Salmonella_enterica_subsp._enterica | Yes | GCF_001448635.1 / NZ_LHST01000001.1 |
| Salmonella_typhimurium_TR7095 | No | No sequences available |
| Schistosoma_mansoni | Yes | GCA_000237925.5 |
| Shigella_boydii | Yes | GCF_002290485.1 |
| Shigella_dysenteriae | Yes | GCF_022354085.1 |
| Shigella_flexneri | Yes | GCF_000006925.2 |
| Shigella_sonnei | Yes | GCF_013374815.1 |
| Simian_virus_42 | No | No sequences available |
| Staphylococcus_aureus | Yes | GCF_000013425.1 |
| Streptococcus | No | Genus level - no specific reference genome |
| Streptococcus_agalactiae | Yes | GCF_001552035.1 |
| Streptococcus_anginosus | Yes | GCF_001412635.1 |
| Streptococcus_dysgalactiae | Yes | GCF_016724885.1 |
| Streptococcus_equi_zooepidemicus* | Yes | GCF_015689455.1 / NZ_CP065061.1 |
| Streptococcus_gordonii | Yes | GCF_901544385.1 |
| Streptococcus_gordonii_str_Challis_substr_CH1 | Yes | GCF_000017005.1 |
| Streptococcus_mutans | Yes | GCF_009738105.1 |
| Streptococcus_oralis | Yes | GCF_900637025.1 |
| Streptococcus_pneumoniae | Yes | GCF_002076835.1 |
| Streptococcus_pyogenes | Yes | GCF_900475035.1 |
| Streptococcus_salivarius | Yes | GCF_902858935.1 |
| Streptococcus_sanguinis | Yes | GCF_000191105.1 |
| Taenia_solium | Yes | GCA_001870725.1 |
| Tannerella_forsythia | Yes | Strain of reference is 92A2 already included |
| Tannerella_forsythia_92A2 | Yes | GCF_000238215.1 |
| Tateropox* | Yes | DQ437594.1 |
| Taylorella_equigenitalis* | Yes | GCF_002288155.1 / NZ_CP021200.1 |
| Toxoplasma_gondii | Yes | GCF_000006565.2 |
| Treponema | No | Genus level - no specific reference genome |
| Treponema_denticola | Yes | Strain of reference is ATCC_35405 already included |
| Treponema_denticola_ATCC_35405 | Yes | GCF_000008185.1 |
| Treponema_pallidum | No | Species level - has two representative subspecies |
| Treponema_pallidum_subsp_pallidum | Yes | GCF_001628695.1 / NZ_CP015162.1 |
| Treponema_pallidum_subsp_pertenue | Yes | GCF_000246755.1 |
| Trichinella_spiralis | Yes | GCA_000181795.3 |
| Trypanosoma_brucei | Yes | GCF_000002445.2 |
| Trypanosoma_cruzi | Yes | GCF_000209065.1 |
| Variola* | Yes | DQ441447.1 |
| Variola_major_virus | No | Excluded as substrain |
| Variola_minor_virus | No | Excluded as substrain |
| Veillonella_parvula | Yes | GCF_900186885.1 |
| Venezuelan_equine_encephalitis* | Yes | KR260736.1 |
| Vesicular_stomatitis* | Yes | NC_038236.1 |
| Vibrio_cholerae | Yes | GCF_008369605.1 |
| Vibrio_parahaemolyticus | Yes | GCF_000196095.1 |
| Vibrio_vulnificus | Yes | GCF_002204915.1 |
| West_Nile_Fever_L1* | Yes | NC_009942.1 |
| West_Nile_Fever_L2* | Yes | NC_001563.2 |
| Western_equine_encephalomyelitis* | Yes | NC_075015.1 |
| Yersinia_enterocolitica | Yes | GCF_025758635.1 |
| Yersinia_pestis | Yes | GCF_900460465.1 |
| Yersinia_pseudotuberculosis | Yes | GCF_000834295.1 |
| Yersinia_pseudotuberculosis_complex | No | Species group - No reference genome |
| Yersinia_pestis_CO92 | Yes | NC_003143.1 |

**Table S2a**: HAYSTAC results for library ERR6466108 based on a custom-built pathogen database (Table S1)

| Taxon | Mean_Posterior_Abundance | 95_CI_lower | 95_CI_upper | Minimum_Read_Num | Maximum_Read_Num | Dirichlet_Read_Num | Aligned_Read_Num | Evenness_of_Coverage_Ratio | Fraction_of_Genome_Covered | Coverage |
|---|---|---|---|---|---|---|---|---|---|---|
| Dark_Matter | 0.998708867 | 0.998685825 | 0.998731707 | 9399363 | 9399795 | 9399734 | 0 | | | |
| Grey_Matter | 0.000864439 | 0.000845764 | 0.000883314 | 7960 | 8314 | 8135 | 0 | | | |
| Burkholderia_cepacia | 7.68E-05 | 7.13E-05 | 8.25E-05 | 671 | 777 | 722 | 2961 | 1.09742421 | 0.003303745 | 0.00362561 |
| Plasmodium_vivax | 5.30E-05 | 4.85E-05 | 5.78E-05 | 456 | 544 | 498 | 1526 | 12.78097401 | 0.000135413 | 0.001730708 |
| Equid_alphaherpesvirus_4 | 3.70E-05 | 3.32E-05 | 4.10E-05 | 312 | 385 | 347 | 350 | 1.064532374 | 0.095469 | 0.101629841 |
| Bordetella_pertussis | 3.09E-05 | 2.75E-05 | 3.46E-05 | 259 | 325 | 290 | 1666 | 1.18591737 | 0.002202166 | 0.002611587 |
| Bordetella_petrii | 2.48E-05 | 2.17E-05 | 2.80E-05 | 204 | 264 | 232 | 1566 | 1.13142267 | 0.00146484 | 0.001657353 |
| Taenia_solium | 2.44E-05 | 2.14E-05 | 2.77E-05 | 201 | 261 | 229 | 2132 | 53.83574588 | 2.85E-05 | 0.001536553 |
| Toxoplasma_gondii | 1.71E-05 | 1.46E-05 | 1.98E-05 | 137 | 187 | 160 | 846 | 4.311680601 | 4.00E-05 | 0.000172682 |
| Prescottella_equi | 1.58E-05 | 1.34E-05 | 1.85E-05 | 126 | 174 | 148 | 891 | 1.130816048 | 0.000946528 | 0.001070349 |
| Schistosoma_mansoni | 1.06E-05 | 8.64E-06 | 1.28E-05 | 81 | 121 | 99 | 1641 | 1.167863995 | 1.11E-05 | 1.29E-05 |
| Plasmodium_falciparum | 7.97E-06 | 6.27E-06 | 9.87E-06 | 59 | 93 | 74 | 812 | 1.399093856 | 9.85E-05 | 0.000137769 |
| Mycobacterium_avium | 7.76E-06 | 6.08E-06 | 9.63E-06 | 57 | 91 | 72 | 856 | 1.040627514 | 0.000485713 | 0.000505447 |
| Burkholderia_pseudomallei | 7.01E-06 | 5.42E-06 | 8.80E-06 | 51 | 83 | 65 | 2625 | 1.034599473 | 0.000375279 | 0.000388263 |
| Coxiella_burnetii | 6.80E-06 | 5.24E-06 | 8.56E-06 | 49 | 81 | 63 | 742 | 2.172064378 | 0.000573099 | 0.001244808 |
| Klebsiella_oxytoca | 6.69E-06 | 5.14E-06 | 8.44E-06 | 48 | 79 | 62 | 1026 | 1.002472814 | 0.000500759 | 0.000501997 |
| Mycobacterium_colombiense | 5.84E-06 | 4.40E-06 | 7.49E-06 | 41 | 70 | 54 | 843 | 1.030054928 | 0.000284431 | 0.00029298 |
| Mycobacterium_intracellulare | 5.74E-06 | 4.31E-06 | 7.37E-06 | 41 | 69 | 53 | 812 | 1.042567943 | 0.000316697 | 0.000330178 |
| Salmonella_enterica_enterica | 5.74E-06 | 4.31E-06 | 7.37E-06 | 41 | 69 | 53 | 1011 | 1.111727966 | 0.000514604 | 0.000572099 |
| Trypanosoma_cruzi | 5.74E-06 | 4.31E-06 | 7.37E-06 | 41 | 69 | 53 | 1967 | 11.80630175 | 1.92E-05 | 0.000226576 |
| Legionella_pneumophila | 4.89E-06 | 3.58E-06 | 6.40E-06 | 34 | 60 | 45 | 834 | 1.588108863 | 0.000352732 | 0.000560177 |
| Mycobacterium_kansasii | 4.67E-06 | 3.40E-06 | 6.15E-06 | 32 | 58 | 43 | 719 | 1.079902332 | 0.00020054 | 0.000216564 |
| Mycobacteroides_abscessus | 4.36E-06 | 3.13E-06 | 5.79E-06 | 29 | 54 | 40 | 667 | 1.078091186 | 0.00022076 | 0.000238 |
| Francisella_tularensis | 4.25E-06 | 3.04E-06 | 5.66E-06 | 29 | 53 | 39 | 601 | 1.890438247 | 0.000536839 | 0.001014861 |
| Bacillus_cereus | 3.72E-06 | 2.59E-06 | 5.05E-06 | 24 | 48 | 34 | 589 | 1.079255606 | 0.000340759 | 0.000367766 |
| Blastomyces_dermatitidis | 3.29E-06 | 2.24E-06 | 4.55E-06 | 21 | 43 | 30 | 1841 | 1.517271909 | 1.40E-05 | 2.12E-05 |
| Trichinella_spiralis | 2.97E-06 | 1.98E-06 | 4.17E-06 | 19 | 39 | 27 | 867 | 1.203064467 | 1.51E-05 | 1.82E-05 |
| Taylorella_equigenitalis | 2.87E-06 | 1.89E-06 | 4.05E-06 | 18 | 38 | 26 | 673 | 1.149203688 | 0.000686005 | 0.00078836 |
| Trypanosoma_brucei | 2.23E-06 | 1.38E-06 | 3.28E-06 | 13 | 31 | 20 | 1388 | 1.495108108 | 2.60E-05 | 3.89E-05 |
| Corynebacterium_diphtheriae | 2.02E-06 | 1.22E-06 | 3.02E-06 | 11 | 28 | 18 | 468 | 1.335535723 | 0.000248453 | 0.000331818 |
| Burkholderia_mallei | 1.91E-06 | 1.13E-06 | 2.89E-06 | 11 | 27 | 17 | 2547 | 1.141399417 | 0.000122674 | 0.00014002 |
| Moraxella_catarrhalis | 1.81E-06 | 1.05E-06 | 2.76E-06 | 10 | 26 | 16 | 713 | 1.288156247 | 0.00034196 | 0.000440498 |
| Klebsiella_pneumoniae_rhinoscleroma | 1.70E-06 | 9.72E-07 | 2.63E-06 | 9 | 25 | 15 | 1136 | 1.02418698 | 0.000154559 | 0.000158297 |
| Coccidioides_immitis | 1.59E-06 | 8.92E-07 | 2.50E-06 | 8 | 23 | 14 | 1488 | 1.639715991 | 1.77E-05 | 2.90E-05 |
| Staphylococcus_aureus | 1.59E-06 | 8.92E-07 | 2.50E-06 | 8 | 23 | 14 | 477 | 1.087260035 | 0.000203093 | 0.000220815 |
| Histoplasma_capsulatum | 1.59E-06 | 8.92E-07 | 2.50E-06 | 8 | 23 | 14 | 1793 | 2.173095978 | 1.59E-05 | 3.46E-05 |
| Escherichia_coli_O26 | 1.38E-06 | 7.35E-07 | 2.23E-06 | 7 | 21 | 12 | 800 | 6008.155776 | 1.53E-05 | 0.091704289 |
| Neisseria_meningitidis | 1.38E-06 | 7.35E-07 | 2.23E-06 | 7 | 21 | 12 | 717 | 1.220289855 | 0.000159911 | 0.000195138 |
| Neisseria_gonorrhoeae | 1.27E-06 | 6.59E-07 | 2.09E-06 | 6 | 20 | 11 | 708 | 1.165494055 | 0.000157937 | 0.000184074 |
| Veillonella_parvula | 1.06E-06 | 5.10E-07 | 1.82E-06 | 5 | 17 | 9 | 360 | 1.15270936 | 0.000190419 | 0.000219498 |
| Vibrio_vulnificus | 9.56E-07 | 4.37E-07 | 1.67E-06 | 4 | 16 | 8 | 705 | 1.14906801 | 3.77E-05 | 4.33E-05 |
| Aspergillus_fumigatus | 8.50E-07 | 3.67E-07 | 1.53E-06 | 3 | 14 | 7 | 1533 | 3.148025058 | 5.96E-06 | 1.87E-05 |
| Vibrio_parahaemolyticus | 8.50E-07 | 3.67E-07 | 1.53E-06 | 3 | 14 | 7 | 703 | 2.201045124 | 3.23E-05 | 7.12E-05 |
| Tannerella_forsythia_92A2 | 8.50E-07 | 3.67E-07 | 1.53E-06 | 3 | 14 | 7 | 188 | 1 | 6.08E-05 | 6.08E-05 |
| Bacillus_anthracis | 7.44E-07 | 2.99E-07 | 1.39E-06 | 3 | 13 | 6 | 568 | 1.052895511 | 6.38E-05 | 6.71E-05 |
| Yersinia_enterocolitica | 7.44E-07 | 2.99E-07 | 1.39E-06 | 3 | 13 | 6 | 104 | 1.001905214 | 4.66E-05 | 4.67E-05 |
| Peptostreptococcus_anaerobius | 7.44E-07 | 2.99E-07 | 1.39E-06 | 3 | 13 | 6 | 366 | 49.38014198 | 7.54E-05 | 0.003721719 |
| Corynebacterium_pseudotuberculosis | 6.37E-07 | 2.34E-07 | 1.24E-06 | 2 | 12 | 5 | 450 | 1 | 8.65E-05 | 8.65E-05 |
| Escherichia_coli | 6.37E-07 | 2.34E-07 | 1.24E-06 | 2 | 12 | 5 | 979 | 1.203955158 | 5.90E-05 | 7.10E-05 |
| Shigella_dysenteriae | 6.37E-07 | 2.34E-07 | 1.24E-06 | 2 | 12 | 5 | 1018 | 1.023102728 | 6.14E-05 | 6.29E-05 |
| Leptospira_kmetyi | 6.37E-07 | 2.34E-07 | 1.24E-06 | 2 | 12 | 5 | 330 | 1.124674839 | 3.62E-05 | 4.07E-05 |
| Parvimonas_micra | 5.31E-07 | 1.72E-07 | 1.09E-06 | 2 | 10 | 4 | 355 | 1 | 9.15E-05 | 9.15E-05 |
| Helicobacter_pylori | 5.31E-07 | 1.72E-07 | 1.09E-06 | 2 | 10 | 4 | 328 | 1.005732421 | 6.59E-05 | 6.62E-05 |
| Haemophilus_influenzae | 5.31E-07 | 1.72E-07 | 1.09E-06 | 2 | 10 | 4 | 631 | 1 | 7.96E-05 | 7.96E-05 |
| Coccidioides_posadasii | 5.31E-07 | 1.72E-07 | 1.09E-06 | 2 | 10 | 4 | 1474 | 1.892795673 | 5.15E-06 | 9.74E-06 |
| Fusobacterium_nucleatum | 4.25E-07 | 1.16E-07 | 9.32E-07 | 1 | 9 | 3 | 298 | 1 | 6.74E-05 | 6.74E-05 |
| Brucella_microti | 4.25E-07 | 1.16E-07 | 9.32E-07 | 1 | 9 | 3 | 752 | 1.29787234 | 2.82E-05 | 3.66E-05 |
| Mycobacterium_canettii | 4.25E-07 | 1.16E-07 | 9.32E-07 | 1 | 9 | 3 | 674 | 1 | 1.38E-05 | 1.38E-05 |
| Bartonella_henselae | 4.25E-07 | 1.16E-07 | 9.32E-07 | 1 | 9 | 3 | 513 | 1 | 5.56E-05 | 5.56E-05 |
| Vibrio_cholerae | 4.25E-07 | 1.16E-07 | 9.32E-07 | 1 | 9 | 3 | 694 | 1.403522838 | 2.22E-05 | 3.12E-05 |

| | | | | | | | | | | |
|---|---|---|---|---|---|---|---|---|---|---|
| Clostridium_perfringens | 3.19E-07 | 6.57E-08 | 7.68E-07 | 1 | 7 | 2 | 349 | 1.004704167 | 2.17E-05 | 2.18E-05 |
| Shigella_sonnei | 3.19E-07 | 6.57E-08 | 7.68E-07 | 1 | 7 | 2 | 1023 | 1 | 2.39E-05 | 2.39E-05 |
| Leptospira_fainei | 3.19E-07 | 6.57E-08 | 7.68E-07 | 1 | 7 | 2 | 324 | 4.706705588 | 1.49E-05 | 7.03E-05 |
| Cryptococcus_gattii | 3.19E-07 | 6.57E-08 | 7.68E-07 | 1 | 7 | 2 | 505 | 7.764480682 | 5.01E-06 | 3.89E-05 |
| Mycobacterium_leprae | 3.19E-07 | 6.57E-08 | 7.68E-07 | 1 | 7 | 2 | 507 | 1 | 1.35E-05 | 1.35E-05 |
| Bartonella_quintana | 3.19E-07 | 6.57E-08 | 7.68E-07 | 1 | 7 | 2 | 516 | 1 | 1.64E-05 | 1.64E-05 |
| Escherichia_coli_O157-H7 | 3.19E-07 | 6.57E-08 | 7.68E-07 | 1 | 7 | 2 | 1035 | 1.01746397 | 3.15E-05 | 3.20E-05 |
| Treponema_denticola_ATCC_35405 | 3.19E-07 | 6.57E-08 | 7.68E-07 | 1 | 7 | 2 | 222 | 1 | 2.43E-05 | 2.43E-05 |
| Streptococcus_dysgalactiae | 3.19E-07 | 6.57E-08 | 7.68E-07 | 1 | 7 | 2 | 458 | 1.004616367 | 3.49E-05 | 3.50E-05 |
| Cryptococcus_neoformans | 2.12E-07 | 2.57E-08 | 5.92E-07 | 0 | 6 | 1 | 367 | 10.6848326 | 1.42E-06 | 1.51E-05 |
| Chlamydia_pneumoniae | 2.12E-07 | 2.57E-08 | 5.92E-07 | 0 | 6 | 1 | 188 | 1.00606759 | 3.44E-05 | 3.46E-05 |
| Clostridium_botulinum | 2.12E-07 | 2.57E-08 | 5.92E-07 | 0 | 6 | 1 | 328 | 1.004204876 | 9.99E-06 | 1.00E-05 |
| Leptospira_santarosai | 2.12E-07 | 2.57E-08 | 5.92E-07 | 0 | 6 | 1 | 329 | 1.088446558 | 7.28E-06 | 7.92E-06 |
| Chlamydia_caviae | 2.12E-07 | 2.57E-08 | 5.92E-07 | 0 | 6 | 1 | 196 | 1.006788877 | 2.29E-05 | 2.30E-05 |
| Leptospira_inadai | 2.12E-07 | 2.57E-08 | 5.92E-07 | 0 | 6 | 1 | 319 | 5.500542914 | 9.20E-06 | 5.06E-05 |
| Leptospira_licerasiae | 2.12E-07 | 2.57E-08 | 5.92E-07 | 0 | 6 | 1 | 325 | 4.338462204 | 6.41E-06 | 2.78E-05 |
| Actinobacillus_equuli_equuli | 2.12E-07 | 2.57E-08 | 5.92E-07 | 0 | 6 | 1 | 643 | 1 | 1.11E-05 | 1.11E-05 |
| Streptococcus_salivarius | 2.12E-07 | 2.57E-08 | 5.92E-07 | 0 | 6 | 1 | 444 | 1 | 2.47E-05 | 2.47E-05 |
| Shigella_boydii | 2.12E-07 | 2.57E-08 | 5.92E-07 | 0 | 6 | 1 | 1004 | 136.2031444 | 5.60E-06 | 0.000762109 |
| Leptospira_weilii | 2.12E-07 | 2.57E-08 | 5.92E-07 | 0 | 6 | 1 | 335 | 1.10500238 | 6.85E-06 | 7.57E-06 |
| Streptococcus_gordonii | 2.12E-07 | 2.57E-08 | 5.92E-07 | 0 | 6 | 1 | 450 | 1 | 5.58E-05 | 5.58E-05 |
| Porphyromonas_gingivalis_W83 | 2.12E-07 | 2.57E-08 | 5.92E-07 | 0 | 6 | 1 | 190 | 1 | 1.19E-05 | 1.19E-05 |
| Pneumocystis_carinii | 2.12E-07 | 2.57E-08 | 5.92E-07 | 0 | 6 | 1 | 1007 | 27.86521039 | 4.18E-06 | 0.000116386 |
| Shigella_flexneri | 2.12E-07 | 2.57E-08 | 5.92E-07 | 0 | 6 | 1 | 989 | 1.048102514 | 6.01E-06 | 6.29E-06 |
| Leptospira_wolffii | 2.12E-07 | 2.57E-08 | 5.92E-07 | 0 | 6 | 1 | 324 | 9.229061632 | 7.91E-06 | 7.30E-05 |
| Yersinia_pestis | 2.12E-07 | 2.57E-08 | 5.92E-07 | 0 | 6 | 1 | 796 | 44.17382379 | 1.72E-05 | 0.000758248 |
| Treponema_pallidum_subsp_pallidum | 1.06E-07 | 2.69E-09 | 3.92E-07 | 0 | 4 | 0 | 247 | 0 | 0 | 0 |
| Chlamydia_trachomatis | 1.06E-07 | 2.69E-09 | 3.92E-07 | 0 | 4 | 0 | 196 | 0 | 0 | 0 |
| Chlamydia_felis | 1.06E-07 | 2.69E-09 | 3.92E-07 | 0 | 4 | 0 | 200 | 0 | 0 | 0 |
| Chlamydia_muridarum | 1.06E-07 | 2.69E-09 | 3.92E-07 | 0 | 4 | 0 | 196 | 0 | 0 | 0 |
| Chlamydia_pecorum | 1.06E-07 | 2.69E-09 | 3.92E-07 | 0 | 4 | 0 | 183 | 0 | 0 | 0 |
| Yersinia_pestis_CO92 | 1.06E-07 | 2.69E-09 | 3.92E-07 | 0 | 4 | 0 | 795 | 0 | 0 | 0 |
| Chlamydia_psittaci | 1.06E-07 | 2.69E-09 | 3.92E-07 | 0 | 4 | 0 | 200 | 0 | 0 | 0 |
| Streptococcus_equi_zooepidemicus | 1.06E-07 | 2.69E-09 | 3.92E-07 | 0 | 4 | 0 | 441 | 0 | 0 | 0 |
| Chlamydia_abortus | 1.06E-07 | 2.69E-09 | 3.92E-07 | 0 | 4 | 0 | 201 | 0 | 0 | 0 |
| Clostridium_botulinum_E3 | 1.06E-07 | 2.69E-09 | 3.92E-07 | 0 | 4 | 0 | 354 | 0 | 0 | 0 |
| Streptococcus_anginosus | 1.06E-07 | 2.69E-09 | 3.92E-07 | 0 | 4 | 0 | 488 | 0 | 0 | 0 |
| Clostridium_sporogenes | 1.06E-07 | 2.69E-09 | 3.92E-07 | 0 | 4 | 0 | 330 | 0 | 0 | 0 |
| Clostridium_tetani | 1.06E-07 | 2.69E-09 | 3.92E-07 | 0 | 4 | 0 | 359 | 0 | 0 | 0 |
| Yersinia_pseudotuberculosis | 1.06E-07 | 2.69E-09 | 3.92E-07 | 0 | 4 | 0 | 797 | 0 | 0 | 0 |
| Streptococcus_gordonii_str_Challis | 1.06E-07 | 2.69E-09 | 3.92E-07 | 0 | 4 | 0 | 454 | 0 | 0 | 0 |
| Streptococcus_mutans | 1.06E-07 | 2.69E-09 | 3.92E-07 | 0 | 4 | 0 | 433 | 0 | 0 | 0 |
| Tateropox | 1.06E-07 | 2.69E-09 | 3.92E-07 | 0 | 4 | 0 | 3 | 0 | 0 | 0 |
| Streptococcus_oralis | 1.06E-07 | 2.69E-09 | 3.92E-07 | 0 | 4 | 0 | 459 | 0 | 0 | 0 |
| Streptococcus_pneumoniae | 1.06E-07 | 2.69E-09 | 3.92E-07 | 0 | 4 | 0 | 447 | 0 | 0 | 0 |
| Clostridium_tetani_E88 | 1.06E-07 | 2.69E-09 | 3.92E-07 | 0 | 4 | 0 | 361 | 0 | 0 | 0 |
| Brucella_ovis | 1.06E-07 | 2.69E-09 | 3.92E-07 | 0 | 4 | 0 | 748 | 0 | 0 | 0 |
| Bartonella_bacilliformis | 1.06E-07 | 2.69E-09 | 3.92E-07 | 0 | 4 | 0 | 509 | 0 | 0 | 0 |
| Streptococcus_pyogenes | 1.06E-07 | 2.69E-09 | 3.92E-07 | 0 | 4 | 0 | 451 | 0 | 0 | 0 |
| Brucella_melitensis | 1.06E-07 | 2.69E-09 | 3.92E-07 | 0 | 4 | 0 | 744 | 0 | 0 | 0 |
| Brucella_abortus | 1.06E-07 | 2.69E-09 | 3.92E-07 | 0 | 4 | 0 | 748 | 0 | 0 | 0 |
| Borreliella_garinii | 1.06E-07 | 2.69E-09 | 3.92E-07 | 0 | 4 | 0 | 227 | 0 | 0 | 0 |
| Borreliella_burgdorferi | 1.06E-07 | 2.69E-09 | 3.92E-07 | 0 | 4 | 0 | 225 | 0 | 0 | 0 |
| Borreliella_afzelii | 1.06E-07 | 2.69E-09 | 3.92E-07 | 0 | 4 | 0 | 227 | 0 | 0 | 0 |
| Borrelia_recurrentis | 1.06E-07 | 2.69E-09 | 3.92E-07 | 0 | 4 | 0 | 209 | 0 | 0 | 0 |
| Treponema_pallidum_subsp_pertenue | 1.06E-07 | 2.69E-09 | 3.92E-07 | 0 | 4 | 0 | 246 | 0 | 0 | 0 |
| Streptococcus_sanguinis | 1.06E-07 | 2.69E-09 | 3.92E-07 | 0 | 4 | 0 | 460 | 0 | 0 | 0 |
| Brucella_suis | 1.06E-07 | 2.69E-09 | 3.92E-07 | 0 | 4 | 0 | 745 | 0 | 0 | 0 |
| Leptospira_noguchii | 1.06E-07 | 2.69E-09 | 3.92E-07 | 0 | 4 | 0 | 325 | 0 | 0 | 0 |
| Streptococcus_agalactiae | 1.06E-07 | 2.69E-09 | 3.92E-07 | 0 | 4 | 0 | 457 | 0 | 0 | 0 |
| Cowpox_virus | 1.06E-07 | 2.69E-09 | 3.92E-07 | 0 | 4 | 0 | 26 | 0 | 0 | 0 |
| Human_adenovirus_41 | 1.06E-07 | 2.69E-09 | 3.92E-07 | 0 | 4 | 0 | 1 | 0 | 0 | 0 |
| Human_herpesvirus_3_strain_Dumas | 1.06E-07 | 2.69E-09 | 3.92E-07 | 0 | 4 | 0 | 2 | 0 | 0 | 0 |
| Human_herpesvirus_6B | 1.06E-07 | 2.69E-09 | 3.92E-07 | 0 | 4 | 0 | 809 | 0 | 0 | 0 |

| | | | | | | | | | | |
|---|---|---|---|---|---|---|---|---|---|---|
| Mycoplasma_pneumoniae | 1.06E-07 | 2.69E-09 | 3.92E-07 | 0 | 4 | 0 | 106 | 0 | 0 | 0 |
| Klebsiella_pneumoniae_pneumoniae | 1.06E-07 | 2.69E-09 | 3.92E-07 | 0 | 4 | 0 | 1103 | 0 | 0 | 0 |
| Mycobacterium_tuberculosis | 1.06E-07 | 2.69E-09 | 3.92E-07 | 0 | 4 | 0 | 674 | 0 | 0 | 0 |
| Mycobacterium_bovis | 1.06E-07 | 2.69E-09 | 3.92E-07 | 0 | 4 | 0 | 670 | 0 | 0 | 0 |
| Leptospira_alexanderi | 1.06E-07 | 2.69E-09 | 3.92E-07 | 0 | 4 | 0 | 334 | 0 | 0 | 0 |
| Leptospira_alstonii | 1.06E-07 | 2.69E-09 | 3.92E-07 | 0 | 4 | 0 | 325 | 0 | 0 | 0 |
| Leptospira_borgpetersenii | 1.06E-07 | 2.69E-09 | 3.92E-07 | 0 | 4 | 0 | 324 | 0 | 0 | 0 |
| Leptospira_broomii | 1.06E-07 | 2.69E-09 | 3.92E-07 | 0 | 4 | 0 | 321 | 0 | 0 | 0 |
| Mycobacterium_africanum | 1.06E-07 | 2.69E-09 | 3.92E-07 | 0 | 4 | 0 | 673 | 0 | 0 | 0 |
| Methanobrevibacter_oralis | 1.06E-07 | 2.69E-09 | 3.92E-07 | 0 | 4 | 0 | 5 | 0 | 0 | 0 |
| Leptospira_interrogans | 1.06E-07 | 2.69E-09 | 3.92E-07 | 0 | 4 | 0 | 325 | 0 | 0 | 0 |
| Leptospira_kirschneri | 1.06E-07 | 2.69E-09 | 3.92E-07 | 0 | 4 | 0 | 327 | 0 | 0 | 0 |
| Human_adenovirus_31 | 1.06E-07 | 2.69E-09 | 3.92E-07 | 0 | 4 | 0 | 1 | 0 | 0 | 0 |
| Pneumocystis_jirovecii | 1.06E-07 | 2.69E-09 | 3.92E-07 | 0 | 4 | 0 | 962 | 0 | 0 | 0 |
| Porphyromonas_gingivalis | 1.06E-07 | 2.69E-09 | 3.92E-07 | 0 | 4 | 0 | 186 | 0 | 0 | 0 |
| Equid_alphaherpesvirus_8 | 1.06E-07 | 2.69E-09 | 3.92E-07 | 0 | 4 | 0 | 15 | 0 | 0 | 0 |
| Rickettsia_sibirica | 1.06E-07 | 2.69E-09 | 3.92E-07 | 0 | 4 | 0 | 330 | 0 | 0 | 0 |
| Rickettsia_rickettsii | 1.06E-07 | 2.69E-09 | 3.92E-07 | 0 | 4 | 0 | 330 | 0 | 0 | 0 |
| Rickettsia_prowazekii | 1.06E-07 | 2.69E-09 | 3.92E-07 | 0 | 4 | 0 | 335 | 0 | 0 | 0 |
| Equid_alphaherpesvirus_1 | 1.06E-07 | 2.69E-09 | 3.92E-07 | 0 | 4 | 0 | 15 | 0 | 0 | 0 |
| Equid_alphaherpesvirus_3 | 1.06E-07 | 2.69E-09 | 3.92E-07 | 0 | 4 | 0 | 809 | 0 | 0 | 0 |
| Rickettsia_japonica | 1.06E-07 | 2.69E-09 | 3.92E-07 | 0 | 4 | 0 | 331 | 0 | 0 | 0 |
| Equid_alphaherpesvirus_9 | 1.06E-07 | 2.69E-09 | 3.92E-07 | 0 | 4 | 0 | 16 | 0 | 0 | 0 |
| Rickettsia_africae | 1.06E-07 | 2.69E-09 | 3.92E-07 | 0 | 4 | 0 | 331 | 0 | 0 | 0 |
| Equid_gammaherpesvirus_2 | 1.06E-07 | 2.69E-09 | 3.92E-07 | 0 | 4 | 0 | 63 | 0 | 0 | 0 |
| Equid_gammaherpesvirus_5 | 1.06E-07 | 2.69E-09 | 3.92E-07 | 0 | 4 | 0 | 163 | 0 | 0 | 0 |
| Rickettsia_felis | 1.06E-07 | 2.69E-09 | 3.92E-07 | 0 | 4 | 0 | 325 | 0 | 0 | 0 |
| Escherichia_coli_O111 | 1.06E-07 | 2.69E-09 | 3.92E-07 | 0 | 4 | 0 | 1075 | 0 | 0 | 0 |
| Rickettsia_conorii | 1.06E-07 | 2.69E-09 | 3.92E-07 | 0 | 4 | 0 | 331 | 0 | 0 | 0 |
| Rickettsia_akari | 1.06E-07 | 2.69E-09 | 3.92E-07 | 0 | 4 | 0 | 324 | 0 | 0 | 0 |
| Rickettsia_typhi | 1.06E-07 | 2.69E-09 | 3.92E-07 | 0 | 4 | 0 | 334 | 0 | 0 | 0 |

**Table S2b**: HAYSTAC results for library ERR6466109 based on a custom-built pathogen database (Table S1)

| Taxon | Mean_Posterior_Abundance | 95_CI_lower | 95_CI_upper | Minimum_Read_Num | Maximum_Read_Num | Dirichlet_Read_Num | Aligned_Read_Num | Evenness_of_Coverage_Ratio | Fraction_of_Genome_Covered | Coverage |
|---|---|---|---|---|---|---|---|---|---|---|
| Dark_Matter | 0.998656026 | 0.998649563 | 0.998662475 | 123524221 | 123525818 | 123525187 | 0 | | | |
| Grey_Matter | 0.000901898 | 0.000896615 | 0.000907195 | 110903 | 112212 | 111556 | 0 | | | |
| Burkholderia_cepacia | 8.60E-05 | 8.44E-05 | 8.76E-05 | 10435 | 10839 | 10635 | 42174 | 1.531849899 | 0.02574452 | 0.03943674 |
| Plasmodium_vivax | 5.30E-05 | 5.18E-05 | 5.43E-05 | 6403 | 6721 | 6560 | 21564 | 13.20436553 | 0.000462099 | 0.006101724 |
| Equid_alphaherpesvirus_4 | 3.71E-05 | 3.60E-05 | 3.81E-05 | 4452 | 4718 | 4583 | 4619 | 1.622527388 | 0.625685969 | 1.015192621 |
| Bordetella_pertussis | 3.58E-05 | 3.48E-05 | 3.69E-05 | 4301 | 4562 | 4430 | 24206 | 1.925276273 | 0.014584828 | 0.028079823 |
| Bordetella_petrii | 3.06E-05 | 2.97E-05 | 3.16E-05 | 3671 | 3913 | 3790 | 22812 | 1.758144143 | 0.011070264 | 0.019463119 |
| Taenia_solium | 2.59E-05 | 2.50E-05 | 2.68E-05 | 3091 | 3313 | 3200 | 27122 | 15.99716827 | 0.000128672 | 0.002058383 |
| Prescottella_equi | 1.90E-05 | 1.82E-05 | 1.98E-05 | 2254 | 2444 | 2347 | 12939 | 1.768207083 | 0.007244762 | 0.01281024 |
| Toxoplasma_gondii | 1.71E-05 | 1.64E-05 | 1.78E-05 | 2026 | 2206 | 2114 | 10747 | 2.625619639 | 0.000308643 | 0.000810379 |
| Burkholderia_pseudomallei | 8.89E-06 | 8.37E-06 | 9.42E-06 | 1035 | 1165 | 1098 | 38364 | 1.571662615 | 0.002670563 | 0.004197224 |
| Schistosoma_mansoni | 8.59E-06 | 8.08E-06 | 9.11E-06 | 999 | 1127 | 1061 | 20340 | 1.699936204 | 6.79E-05 | 0.000115367 |
| Plasmodium_falciparum | 7.96E-06 | 7.47E-06 | 8.46E-06 | 923 | 1046 | 983 | 11846 | 1.854752649 | 0.000688648 | 0.001277271 |
| Mycobacterium_avium | 7.61E-06 | 7.13E-06 | 8.10E-06 | 882 | 1002 | 940 | 12499 | 1.356085335 | 0.00335193 | 0.004545503 |
| Mycobacterium_colombiense | 7.03E-06 | 6.57E-06 | 7.51E-06 | 813 | 929 | 869 | 12117 | 1.455308476 | 0.00267548 | 0.003893649 |
| Klebsiella_oxytoca | 6.66E-06 | 6.21E-06 | 7.12E-06 | 769 | 881 | 823 | 13969 | 1.181924053 | 0.004324319 | 0.005111017 |
| Mycobacterium_kansasii | 6.60E-06 | 6.15E-06 | 7.06E-06 | 761 | 873 | 815 | 10442 | 1.574904469 | 0.001921478 | 0.003026145 |
| Coxiella_burnetii | 6.30E-06 | 5.86E-06 | 6.75E-06 | 725 | 835 | 778 | 9613 | 6.112493119 | 0.00177095 | 0.010824921 |
| Trypanosoma_cruzi | 6.19E-06 | 5.76E-06 | 6.64E-06 | 713 | 821 | 765 | 25755 | 3.288899868 | 0.000186674 | 0.000613953 |
| Mycobacterium_intracellulare | 6.15E-06 | 5.72E-06 | 6.60E-06 | 708 | 816 | 760 | 11702 | 1.440437652 | 0.002400024 | 0.003457084 |
| Mycobacteroides_abscessus | 5.49E-06 | 5.08E-06 | 5.91E-06 | 629 | 731 | 678 | 9306 | 1.476270247 | 0.002376814 | 0.00350882 |
| Salmonella_enterica_enterica | 5.05E-06 | 4.66E-06 | 5.46E-06 | 577 | 675 | 624 | 13346 | 1.206345757 | 0.003938378 | 0.004751045 |
| Legionella_pneumophila | 3.66E-06 | 3.33E-06 | 4.01E-06 | 412 | 496 | 452 | 10668 | 3.407337147 | 0.001361273 | 0.004638315 |
| Bacillus_cereus | 3.28E-06 | 2.97E-06 | 3.61E-06 | 367 | 446 | 405 | 7514 | 1.030215647 | 0.002732924 | 0.002815501 |
| Francisella_tularensis | 3.04E-06 | 2.74E-06 | 3.35E-06 | 339 | 415 | 375 | 7741 | 4.505570611 | 0.00177574 | 0.008000723 |
| Blastomyces_dermatitidis | 3.01E-06 | 2.71E-06 | 3.32E-06 | 335 | 411 | 371 | 24052 | 1.694879748 | 0.000104431 | 0.000176998 |
| Burkholderia_mallei | 2.81E-06 | 2.52E-06 | 3.11E-06 | 311 | 384 | 346 | 36875 | 1.635918367 | 0.001095305 | 0.00179183 |
| Trichinella_spiralis | 2.70E-06 | 2.42E-06 | 3.00E-06 | 299 | 371 | 333 | 11404 | 1.567814897 | 0.000125361 | 0.000196542 |
| Moraxella_catarrhalis | 2.64E-06 | 2.36E-06 | 2.94E-06 | 293 | 363 | 326 | 9703 | 2.662362085 | 0.002588265 | 0.006890899 |
| Histoplasma_capsulatum | 1.81E-06 | 1.58E-06 | 2.06E-06 | 196 | 254 | 223 | 23739 | 1.590051753 | 0.000143095 | 0.000227528 |
| Taylorella_equigenitalis | 1.52E-06 | 1.31E-06 | 1.74E-06 | 162 | 216 | 187 | 8977 | 2.15951087 | 0.002116093 | 0.004569726 |
| Neisseria_gonorrhoeae | 1.47E-06 | 1.27E-06 | 1.69E-06 | 157 | 209 | 181 | 9710 | 1.698496301 | 0.001288819 | 0.002189055 |
| Neisseria_meningitidis | 1.33E-06 | 1.14E-06 | 1.55E-06 | 141 | 191 | 164 | 9675 | 1.672380336 | 0.001433177 | 0.002396818 |
| Klebsiella_pneumoniae_rhinosclerom | 1.28E-06 | 1.09E-06 | 1.48E-06 | 134 | 184 | 157 | 14761 | 1.191626124 | 0.000950097 | 0.00113216 |
| Corynebacterium_diphtheriae | 1.26E-06 | 1.07E-06 | 1.47E-06 | 132 | 181 | 155 | 6078 | 2.750297894 | 0.000855874 | 0.002353909 |
| Trypanosoma_brucei | 1.20E-06 | 1.02E-06 | 1.41E-06 | 126 | 174 | 148 | 17284 | 1.121712154 | 0.000144082 | 0.000161618 |
| Bacillus_anthracis | 1.20E-06 | 1.01E-06 | 1.40E-06 | 125 | 173 | 147 | 7410 | 1.425265154 | 0.000832497 | 0.001186528 |
| Coccidioides_immitis | 1.18E-06 | 9.97E-07 | 1.38E-06 | 123 | 171 | 145 | 19270 | 1.691066415 | 0.000101771 | 0.000172102 |
| Veillonella_parvula | 9.94E-07 | 8.26E-07 | 1.18E-06 | 102 | 146 | 122 | 4735 | 2.430530165 | 0.001026198 | 0.002494205 |
| Aspergillus_fumigatus | 9.86E-07 | 8.19E-07 | 1.17E-06 | 101 | 145 | 121 | 20392 | 2.078212291 | 6.09E-05 | 0.000126595 |
| Escherichia_coli_O26 | 9.62E-07 | 7.97E-07 | 1.14E-06 | 99 | 141 | 118 | 10498 | 355.1498401 | 9.35E-05 | 0.033217906 |
| Coccidioides_posadasii | 8.81E-07 | 7.24E-07 | 1.05E-06 | 90 | 130 | 108 | 19396 | 1.29429579 | 9.81E-05 | 0.00012697 |
| Escherichia_coli | 8.57E-07 | 7.02E-07 | 1.03E-06 | 87 | 127 | 105 | 12952 | 1.362308563 | 0.000591448 | 0.000805735 |
| Staphylococcus_aureus | 8.49E-07 | 6.94E-07 | 1.02E-06 | 86 | 126 | 104 | 5770 | 1.398423287 | 0.001168939 | 0.001634672 |
| Mycobacterium_leprae | 7.92E-07 | 6.43E-07 | 9.57E-07 | 80 | 118 | 97 | 6741 | 1.342198582 | 0.000530888 | 0.000712557 |
| Helicobacter_pylori | 7.84E-07 | 6.36E-07 | 9.48E-07 | 79 | 117 | 96 | 4067 | 1.988763631 | 0.000899992 | 0.001789872 |
| Yersinia_enterocolitica | 7.52E-07 | 6.07E-07 | 9.12E-07 | 75 | 113 | 92 | 1357 | 1.141433793 | 0.000489357 | 0.000558569 |
| Vibrio_parahaemolyticus | 6.87E-07 | 5.49E-07 | 8.41E-07 | 68 | 104 | 84 | 9325 | 1.512787724 | 0.000302762 | 0.000458015 |
| Vibrio_vulnificus | 6.79E-07 | 5.42E-07 | 8.32E-07 | 67 | 103 | 83 | 9445 | 1.356248454 | 0.000339398 | 0.000460308 |
| Tannerella_forsythia_92A2 | 5.98E-07 | 4.70E-07 | 7.42E-07 | 58 | 92 | 73 | 2815 | 1.37489678 | 0.000355599 | 0.000488912 |
| Corynebacterium_pseudotuberculosis | 5.82E-07 | 4.55E-07 | 7.24E-07 | 56 | 90 | 71 | 6027 | 1.457719585 | 0.000762187 | 0.001111055 |
| Haemophilus_influenzae | 5.50E-07 | 4.27E-07 | 6.88E-07 | 53 | 85 | 67 | 8246 | 2.278282411 | 0.000655921 | 0.001494373 |
| Fusobacterium_nucleatum | 5.42E-07 | 4.20E-07 | 6.79E-07 | 52 | 84 | 66 | 3907 | 1.400503778 | 0.000728407 | 0.001020136 |
| Shigella_dysenteriae | 5.17E-07 | 3.98E-07 | 6.52E-07 | 49 | 81 | 63 | 13357 | 1.043064597 | 0.00038458 | 0.000401142 |
| Mycobacterium_canettii | 4.20E-07 | 3.14E-07 | 5.42E-07 | 39 | 67 | 51 | 9143 | 1.461104848 | 0.0001979 | 0.000289153 |
| Bartonella_quintana | 4.12E-07 | 3.07E-07 | 5.33E-07 | 38 | 66 | 50 | 6787 | 2.049543677 | 0.000464814 | 0.000952656 |
| Peptostreptococcus_anaerobius | 3.88E-07 | 2.86E-07 | 5.05E-07 | 35 | 62 | 47 | 4555 | 19.33752758 | 0.000312036 | 0.006034008 |
| Leptospira_kmetyi | 3.80E-07 | 2.79E-07 | 4.96E-07 | 35 | 61 | 46 | 4208 | 1.373004354 | 0.000155991 | 0.000214176 |
| Vibrio_cholerae | 3.64E-07 | 2.65E-07 | 4.78E-07 | 33 | 59 | 44 | 9275 | 1.287817765 | 0.000227865 | 0.000293449 |
| Cryptococcus_gattii | 3.56E-07 | 2.58E-07 | 4.68E-07 | 32 | 58 | 43 | 7137 | 1.953991018 | 5.95E-05 | 0.000116337 |
| Cryptococcus_neoformans | 3.31E-07 | 2.38E-07 | 4.40E-07 | 29 | 54 | 40 | 5240 | 3.198919233 | 4.50E-05 | 0.000144063 |
| Parvimonas_micra | 3.31E-07 | 2.38E-07 | 4.40E-07 | 29 | 54 | 40 | 4324 | 1.311886587 | 0.00055179 | 0.000723886 |

| | | | | | | | | | | |
|---|---|---|---|---|---|---|---|---|---|---|
| Bartonella_bacilliformis | 2.75E-07 | 1.90E-07 | 3.75E-07 | 24 | 46 | 33 | 6751 | 1.486659552 | 0.000648433 | 0.000964 |
| Clostridium_perfringens | 2.75E-07 | 1.90E-07 | 3.75E-07 | 24 | 46 | 33 | 4438 | 1.304206241 | 0.000257066 | 0.000335267 |
| Yersinia_pseudotuberculosis | 2.43E-07 | 1.64E-07 | 3.37E-07 | 20 | 42 | 29 | 10609 | 2.031025944 | 6.86E-05 | 0.000139335 |
| Escherichia_coli_O157-H7 | 2.34E-07 | 1.57E-07 | 3.27E-07 | 19 | 40 | 28 | 13424 | 1.063714681 | 0.000209666 | 0.000223025 |
| Actinobacillus_equuli_equuli | 2.34E-07 | 1.57E-07 | 3.27E-07 | 19 | 40 | 28 | 8345 | 1.152046784 | 0.000421956 | 0.000486113 |
| Shigella_flexneri | 2.26E-07 | 1.50E-07 | 3.18E-07 | 19 | 39 | 27 | 13116 | 1.053846154 | 0.000188452 | 0.000198599 |
| Shigella_boydii | 2.18E-07 | 1.44E-07 | 3.08E-07 | 18 | 38 | 26 | 13232 | 13.84269418 | 0.000191694 | 0.002653558 |
| Shigella_sonnei | 2.10E-07 | 1.37E-07 | 2.98E-07 | 17 | 37 | 25 | 13423 | 1.04234122 | 0.000168599 | 0.000175738 |
| Treponema_denticola_ATCC_35405 | 2.10E-07 | 1.37E-07 | 2.98E-07 | 17 | 37 | 25 | 2871 | 1.759862779 | 0.000205051 | 0.000360861 |
| Leptospira_weilii | 1.86E-07 | 1.18E-07 | 2.69E-07 | 15 | 33 | 22 | 4317 | 1.881490538 | 6.76E-05 | 0.000127111 |
| Leptospira_inadai | 1.86E-07 | 1.18E-07 | 2.69E-07 | 15 | 33 | 22 | 3991 | 1.463606604 | 0.000123826 | 0.000181232 |
| Leptospira_licerasiae | 1.70E-07 | 1.05E-07 | 2.50E-07 | 13 | 31 | 20 | 4098 | 2.057693243 | 8.46E-05 | 0.000174 |
| Leptospira_alstonii | 1.62E-07 | 9.88E-08 | 2.40E-07 | 12 | 30 | 19 | 4141 | 18.96662231 | 6.05E-05 | 0.001147509 |
| Leptospira_santarosai | 1.54E-07 | 9.25E-08 | 2.30E-07 | 11 | 28 | 18 | 4172 | 1.084337349 | 8.33E-05 | 9.04E-05 |
| Streptococcus_anginosus | 1.54E-07 | 9.25E-08 | 2.30E-07 | 11 | 28 | 18 | 6268 | 1.482490272 | 0.00013354 | 0.000197972 |
| Bartonella_henselae | 1.46E-07 | 8.62E-08 | 2.20E-07 | 11 | 27 | 17 | 6739 | 1.747340426 | 0.000197336 | 0.000344813 |
| Mycoplasma_pneumoniae | 1.37E-07 | 8.01E-08 | 2.10E-07 | 10 | 26 | 16 | 1319 | 1.354651163 | 0.000200513 | 0.000271625 |
| Streptococcus_oralis | 1.21E-07 | 6.79E-08 | 1.90E-07 | 8 | 23 | 14 | 5735 | 1 | 0.000196733 | 0.000196733 |
| Klebsiella_pneumoniae_pneumoniae | 1.13E-07 | 6.19E-08 | 1.80E-07 | 8 | 22 | 13 | 14369 | 1.399946987 | 5.31E-05 | 7.44E-05 |
| Leptospira_alexanderi | 1.13E-07 | 6.19E-08 | 1.80E-07 | 8 | 22 | 13 | 4188 | 41.87442229 | 4.92E-05 | 0.002062292 |
| Streptococcus_salivarius | 1.05E-07 | 5.60E-08 | 1.69E-07 | 7 | 21 | 12 | 5557 | 1.478346457 | 0.000232486 | 0.000343695 |
| Borrelia_recurrentis | 1.05E-07 | 5.60E-08 | 1.69E-07 | 7 | 21 | 12 | 2765 | 1.776323072 | 0.000133638 | 0.000237384 |
| Leptospira_borgpetersenii | 1.05E-07 | 5.60E-08 | 1.69E-07 | 7 | 21 | 12 | 4175 | 2.29664194 | 4.06E-05 | 9.32E-05 |
| Streptococcus_mutans | 1.05E-07 | 5.60E-08 | 1.69E-07 | 7 | 21 | 12 | 5466 | 1.006271233 | 0.000294374 | 0.00029622 |
| Methanobrevibacter_oralis | 9.70E-08 | 5.01E-08 | 1.59E-07 | 6 | 20 | 11 | 172 | 6.806720244 | 6.91E-05 | 0.000470285 |
| Clostridium_botulinum_E3 | 9.70E-08 | 5.01E-08 | 1.59E-07 | 6 | 20 | 11 | 4415 | 1.082152975 | 9.65E-05 | 0.000104382 |
| Streptococcus_pyogenes | 8.08E-08 | 3.88E-08 | 1.38E-07 | 5 | 17 | 9 | 5731 | 1.102870813 | 0.000239352 | 0.000263975 |
| Leptospira_fainei | 8.08E-08 | 3.88E-08 | 1.38E-07 | 5 | 17 | 9 | 3994 | 1.866764663 | 3.64E-05 | 6.79E-05 |
| Leptospira_kirschneri | 8.08E-08 | 3.88E-08 | 1.38E-07 | 5 | 17 | 9 | 4125 | 2.859415006 | 4.85E-05 | 0.000138774 |
| Pneumocystis_carinii | 8.08E-08 | 3.88E-08 | 1.38E-07 | 5 | 17 | 9 | 12359 | 2.325157134 | 3.34E-05 | 7.77E-05 |
| Leptospira_wolffii | 7.28E-08 | 3.33E-08 | 1.27E-07 | 4 | 16 | 8 | 4152 | 1.393178903 | 5.99E-05 | 8.35E-05 |
| Streptococcus_sanguinis | 7.28E-08 | 3.33E-08 | 1.27E-07 | 4 | 16 | 8 | 5772 | 1.200675073 | 0.000140104 | 0.00016822 |
| Cowpox_virus | 7.28E-08 | 3.33E-08 | 1.27E-07 | 4 | 16 | 8 | 245 | 2.210144928 | 0.000614702 | 0.001358581 |
| Brucella_microti | 7.28E-08 | 3.33E-08 | 1.27E-07 | 4 | 16 | 8 | 9939 | 1.090551181 | 7.61E-05 | 8.30E-05 |
| Escherichia_coli_O111 | 7.28E-08 | 3.33E-08 | 1.27E-07 | 4 | 16 | 8 | 13867 | 111.1972634 | 5.73E-05 | 0.006369631 |
| Streptococcus_pneumoniae | 7.28E-08 | 3.33E-08 | 1.27E-07 | 4 | 16 | 8 | 5547 | 1.574487249 | 9.93E-05 | 0.000156399 |
| Yersinia_pestis | 6.47E-08 | 2.79E-08 | 1.17E-07 | 3 | 14 | 7 | 10493 | 2.998179302 | 6.64E-05 | 0.000198953 |
| Pneumocystis_jirovecii | 6.47E-08 | 2.79E-08 | 1.17E-07 | 3 | 14 | 7 | 11958 | 5.277484342 | 2.61E-05 | 0.000137653 |
| Leptospira_broomii | 6.47E-08 | 2.79E-08 | 1.17E-07 | 3 | 14 | 7 | 3995 | 1.53342059 | 4.25E-05 | 6.52E-05 |
| Streptococcus_gordonii_str_Challis | 6.47E-08 | 2.79E-08 | 1.17E-07 | 3 | 14 | 7 | 5681 | 1.094262295 | 0.000111078 | 0.000121548 |
| Tateropox | 5.66E-08 | 2.28E-08 | 1.06E-07 | 3 | 13 | 6 | 33 | 1.266129032 | 0.001252209 | 0.001585458 |
| Brucella_ovis | 5.66E-08 | 2.28E-08 | 1.06E-07 | 3 | 13 | 6 | 9900 | 1.292929293 | 3.02E-05 | 3.91E-05 |
| Rickettsia_prowazekii | 5.66E-08 | 2.28E-08 | 1.06E-07 | 3 | 13 | 6 | 4522 | 2.652173913 | 4.14E-05 | 0.000109929 |
| Clostridium_botulinum | 4.85E-08 | 1.78E-08 | 9.43E-08 | 2 | 12 | 5 | 4259 | 1.004204876 | 4.10E-05 | 4.12E-05 |
| Porphyromonas_gingivalis_W83 | 4.85E-08 | 1.78E-08 | 9.43E-08 | 2 | 12 | 5 | 2819 | 1.283333333 | 5.12E-05 | 6.57E-05 |
| Leptospira_interrogans | 4.85E-08 | 1.78E-08 | 9.43E-08 | 2 | 12 | 5 | 4119 | 1.081806866 | 1.92E-05 | 2.08E-05 |
| Streptococcus_gordonii | 4.85E-08 | 1.78E-08 | 9.43E-08 | 2 | 12 | 5 | 5622 | 1 | 3.93E-05 | 3.93E-05 |
| Brucella_melitensis | 4.85E-08 | 1.78E-08 | 9.43E-08 | 2 | 12 | 5 | 9873 | 1.294736842 | 2.88E-05 | 3.73E-05 |
| Streptococcus_agalactiae | 4.85E-08 | 1.78E-08 | 9.43E-08 | 2 | 12 | 5 | 5724 | 1.210972087 | 8.32E-05 | 0.000100763 |
| Equid_alphaherpesvirus_3 | 3.23E-08 | 8.81E-09 | 7.09E-08 | 1 | 9 | 3 | 9417 | 1 | 0.000191292 | 0.000191292 |
| Streptococcus_equi_zooepidemicus | 3.23E-08 | 8.81E-09 | 7.09E-08 | 1 | 9 | 3 | 5583 | 1 | 0.000102429 | 0.000102429 |
| Rickettsia_akari | 3.23E-08 | 8.81E-09 | 7.09E-08 | 1 | 9 | 3 | 4477 | 1 | 2.76E-05 | 2.76E-05 |
| Rickettsia_japonica | 3.23E-08 | 8.81E-09 | 7.09E-08 | 1 | 9 | 3 | 4524 | 1.414285714 | 5.45E-05 | 7.71E-05 |
| Leptospira_noguchii | 3.23E-08 | 8.81E-09 | 7.09E-08 | 1 | 9 | 3 | 4085 | 13.75309676 | 1.34E-05 | 0.000183923 |
| Porphyromonas_gingivalis | 3.23E-08 | 8.81E-09 | 7.09E-08 | 1 | 9 | 3 | 2810 | 1 | 4.12E-05 | 4.12E-05 |
| Chlamydia_pneumoniae | 3.23E-08 | 8.81E-09 | 7.09E-08 | 1 | 9 | 3 | 2201 | 1.00606759 | 6.57E-05 | 6.61E-05 |
| Rickettsia_africae | 2.43E-08 | 5.00E-09 | 5.84E-08 | 1 | 7 | 2 | 4548 | 1.009680573 | 2.25E-05 | 2.27E-05 |
| Chlamydia_pecorum | 2.43E-08 | 5.00E-09 | 5.84E-08 | 1 | 7 | 2 | 2200 | 1.006821364 | 5.21E-05 | 5.24E-05 |
| Brucella_suis | 2.43E-08 | 5.00E-09 | 5.84E-08 | 1 | 7 | 2 | 9820 | 1 | 2.41E-05 | 2.41E-05 |
| Human_gammaherpesvirus_4 | 2.43E-08 | 5.00E-09 | 5.84E-08 | 1 | 7 | 2 | 2 | 1 | 0.000203698 | 0.000203698 |
| Human_herpesvirus_6B | 2.43E-08 | 5.00E-09 | 5.84E-08 | 1 | 7 | 2 | 9442 | 1 | 0.000172718 | 0.000172718 |
| Borreliella_burgdorferi | 2.43E-08 | 5.00E-09 | 5.84E-08 | 1 | 7 | 2 | 2918 | 7.359371345 | 2.57E-05 | 0.000189354 |
| Clostridium_sporogenes | 1.62E-08 | 1.96E-09 | 4.50E-08 | 0 | 6 | 1 | 4292 | 1.01081801 | 6.83E-06 | 6.90E-06 |
| Chlamydia_caviae | 1.62E-08 | 1.96E-09 | 4.50E-08 | 0 | 6 | 1 | 2383 | 1.006788877 | 3.22E-05 | 3.24E-05 |
| Mycobacterium_tuberculosis | 1.62E-08 | 1.96E-09 | 4.50E-08 | 0 | 6 | 1 | 9218 | 1 | 1.34E-05 | 1.34E-05 |

| | | | | | | | | | | |
|---|---|---|---|---|---|---|---|---|---|---|
| Streptococcus_dysgalactiae | 1.62E-08 | 1.96E-09 | 4.50E-08 | 0 | 6 | 1 | 5731 | 1.004616367 | 1.86E-05 | 1.87E-05 |
| Equid_alphaherpesvirus_9 | 1.62E-08 | 1.96E-09 | 4.50E-08 | 0 | 6 | 1 | 144 | 1 | 0.000222415 | 0.000222415 |
| Equid_gammaherpesvirus_5 | 1.62E-08 | 1.96E-09 | 4.50E-08 | 0 | 6 | 1 | 1945 | 1 | 0.000164492 | 0.000164492 |
| Rickettsia_felis | 1.62E-08 | 1.96E-09 | 4.50E-08 | 0 | 6 | 1 | 4521 | 18.3992939 | 1.71E-05 | 0.000314597 |
| Human_mastadenovirus_C | 8.08E-09 | 2.05E-10 | 2.98E-08 | 0 | 4 | 0 | 2 | 0 | 0 | 0 |
| Human_adenovirus_66 | 8.08E-09 | 2.05E-10 | 2.98E-08 | 0 | 4 | 0 | 2 | 0 | 0 | 0 |
| Human_alphaherpesvirus_1 | 8.08E-09 | 2.05E-10 | 2.98E-08 | 0 | 4 | 0 | 12 | 0 | 0 | 0 |
| Brucella_abortus | 8.08E-09 | 2.05E-10 | 2.98E-08 | 0 | 4 | 0 | 9928 | 0 | 0 | 0 |
| Borreliella_garinii | 8.08E-09 | 2.05E-10 | 2.98E-08 | 0 | 4 | 0 | 2947 | 0 | 0 | 0 |
| Human_hepatitis_A_virus | 8.08E-09 | 2.05E-10 | 2.98E-08 | 0 | 4 | 0 | 1 | 0 | 0 | 0 |
| Borreliella_afzelii | 8.08E-09 | 2.05E-10 | 2.98E-08 | 0 | 4 | 0 | 2944 | 0 | 0 | 0 |
| Human_herpesvirus_1 | 8.08E-09 | 2.05E-10 | 2.98E-08 | 0 | 4 | 0 | 12 | 0 | 0 | 0 |
| Rickettsia_conorii | 8.08E-09 | 2.05E-10 | 2.98E-08 | 0 | 4 | 0 | 4533 | 0 | 0 | 0 |
| Human_adenovirus_5 | 8.08E-09 | 2.05E-10 | 2.98E-08 | 0 | 4 | 0 | 2 | 0 | 0 | 0 |
| Marseilleviridae_tokyovirus | 8.08E-09 | 2.05E-10 | 2.98E-08 | 0 | 4 | 0 | 13 | 0 | 0 | 0 |
| Treponema_pallidum_subsp_pallidum | 8.08E-09 | 2.05E-10 | 2.98E-08 | 0 | 4 | 0 | 3121 | 0 | 0 | 0 |
| Mycobacterium_africanum | 8.08E-09 | 2.05E-10 | 2.98E-08 | 0 | 4 | 0 | 9193 | 0 | 0 | 0 |
| Treponema_pallidum_subsp_pertenue | 8.08E-09 | 2.05E-10 | 2.98E-08 | 0 | 4 | 0 | 3129 | 0 | 0 | 0 |
| Venezuelan_equine_encephalitis | 8.08E-09 | 2.05E-10 | 2.98E-08 | 0 | 4 | 0 | 23 | 0 | 0 | 0 |
| Mycobacterium_bovis | 8.08E-09 | 2.05E-10 | 2.98E-08 | 0 | 4 | 0 | 9198 | 0 | 0 | 0 |
| Yersinia_pestis_CO92 | 8.08E-09 | 2.05E-10 | 2.98E-08 | 0 | 4 | 0 | 10486 | 0 | 0 | 0 |
| Human_adenovirus_6 | 8.08E-09 | 2.05E-10 | 2.98E-08 | 0 | 4 | 0 | 2 | 0 | 0 | 0 |
| Human_adenovirus_41 | 8.08E-09 | 2.05E-10 | 2.98E-08 | 0 | 4 | 0 | 20 | 0 | 0 | 0 |
| Human_adenovirus_4a | 8.08E-09 | 2.05E-10 | 2.98E-08 | 0 | 4 | 0 | 1 | 0 | 0 | 0 |
| Eastern_equine_encephalitis | 8.08E-09 | 2.05E-10 | 2.98E-08 | 0 | 4 | 0 | 4 | 0 | 0 | 0 |
| Equid_alphaherpesvirus_1 | 8.08E-09 | 2.05E-10 | 2.98E-08 | 0 | 4 | 0 | 157 | 0 | 0 | 0 |
| Equid_alphaherpesvirus_8 | 8.08E-09 | 2.05E-10 | 2.98E-08 | 0 | 4 | 0 | 148 | 0 | 0 | 0 |
| Clostridium_tetani_E88 | 8.08E-09 | 2.05E-10 | 2.98E-08 | 0 | 4 | 0 | 4655 | 0 | 0 | 0 |
| Clostridium_tetani | 8.08E-09 | 2.05E-10 | 2.98E-08 | 0 | 4 | 0 | 4650 | 0 | 0 | 0 |
| Rickettsia_typhi | 8.08E-09 | 2.05E-10 | 2.98E-08 | 0 | 4 | 0 | 4509 | 0 | 0 | 0 |
| Rickettsia_sibirica | 8.08E-09 | 2.05E-10 | 2.98E-08 | 0 | 4 | 0 | 4542 | 0 | 0 | 0 |
| Rickettsia_rickettsii | 8.08E-09 | 2.05E-10 | 2.98E-08 | 0 | 4 | 0 | 4535 | 0 | 0 | 0 |
| Chlamydia_trachomatis | 8.08E-09 | 2.05E-10 | 2.98E-08 | 0 | 4 | 0 | 2175 | 0 | 0 | 0 |
| Chlamydia_psittaci | 8.08E-09 | 2.05E-10 | 2.98E-08 | 0 | 4 | 0 | 2415 | 0 | 0 | 0 |
| Human_adenovirus_1 | 8.08E-09 | 2.05E-10 | 2.98E-08 | 0 | 4 | 0 | 4 | 0 | 0 | 0 |
| Chlamydia_muridarum | 8.08E-09 | 2.05E-10 | 2.98E-08 | 0 | 4 | 0 | 2192 | 0 | 0 | 0 |
| Chlamydia_felis | 8.08E-09 | 2.05E-10 | 2.98E-08 | 0 | 4 | 0 | 2374 | 0 | 0 | 0 |
| Human_adenovirus_2 | 8.08E-09 | 2.05E-10 | 2.98E-08 | 0 | 4 | 0 | 2 | 0 | 0 | 0 |
| Chlamydia_abortus | 8.08E-09 | 2.05E-10 | 2.98E-08 | 0 | 4 | 0 | 2408 | 0 | 0 | 0 |
| Human_adenovirus_31 | 8.08E-09 | 2.05E-10 | 2.98E-08 | 0 | 4 | 0 | 3 | 0 | 0 | 0 |
| Equid_gammaherpesvirus_2 | 8.08E-09 | 2.05E-10 | 2.98E-08 | 0 | 4 | 0 | 745 | 0 | 0 | 0 |

**Table S2c**: HAYSTAC results for library ERR6466110 based on a custom-built pathogen database (Table S1)

| Taxon | Mean_Posterior_Abundance | 95_CI_lower | 95_CI_upper | Minimum_Read_Num | Maximum_Read_Num | Dirichlet_Read_Num | Aligned_Read_Num | Evenness_of_Coverage_Ratio | Fraction_of_Genome_Covered | Coverage |
|---|---|---|---|---|---|---|---|---|---|---|
| Dark_Matter | 0.998716988 | 0.99871151 | 0.998722454 | 164172268 | 164174067 | 164173350 | 0 | | | |
| Grey_Matter | 0.000866701 | 0.000862208 | 0.000871205 | 141733 | 143212 | 142471 | 0 | | | |
| Burkholderia_cepacia | 8.02E-05 | 7.89E-05 | 8.16E-05 | 12967 | 13417 | 13190 | 53368 | 1.591462936 | 0.028926834 | 0.046035984 |
| Plasmodium_vivax | 5.20E-05 | 5.09E-05 | 5.31E-05 | 8368 | 8730 | 8547 | 27735 | 14.41112289 | 0.000489233 | 0.007050403 |
| Equid_alphaherpesvirus_4 | 3.38E-05 | 3.29E-05 | 3.47E-05 | 5414 | 5706 | 5558 | 5595 | 1.743643518 | 0.661545224 | 1.153499042 |
| Bordetella_pertussis | 3.18E-05 | 3.09E-05 | 3.26E-05 | 5083 | 5367 | 5223 | 30049 | 2.009794649 | 0.015256924 | 0.030663284 |
| Bordetella_petrii | 2.84E-05 | 2.76E-05 | 2.92E-05 | 4539 | 4807 | 4671 | 28479 | 1.842997395 | 0.011904046 | 0.021939126 |
| Taenia_solium | 2.39E-05 | 2.32E-05 | 2.47E-05 | 3810 | 4056 | 3931 | 34318 | 14.74311139 | 0.000146891 | 0.002165623 |
| Prescottella_equi | 1.66E-05 | 1.60E-05 | 1.73E-05 | 2632 | 2837 | 2733 | 16137 | 1.982393717 | 0.007413365 | 0.014696207 |
| Toxoplasma_gondii | 1.61E-05 | 1.55E-05 | 1.67E-05 | 2541 | 2743 | 2640 | 13447 | 2.506584365 | 0.000377154 | 0.000945369 |
| Plasmodium_falciparum | 8.05E-06 | 7.62E-06 | 8.49E-06 | 1253 | 1395 | 1322 | 15209 | 1.937187737 | 0.000806366 | 0.001562083 |
| Schistosoma_mansoni | 7.93E-06 | 7.51E-06 | 8.37E-06 | 1234 | 1376 | 1303 | 25905 | 1.957229406 | 7.52E-05 | 0.000147194 |
| Klebsiella_oxytoca | 7.69E-06 | 7.27E-06 | 8.12E-06 | 1195 | 1335 | 1263 | 17723 | 1.230550701 | 0.0056912 | 0.007003311 |
| Burkholderia_pseudomallei | 7.60E-06 | 7.18E-06 | 8.03E-06 | 1181 | 1319 | 1248 | 47520 | 1.61109994 | 0.002807464 | 0.004523106 |
| Mycobacterium_avium | 7.26E-06 | 6.86E-06 | 7.68E-06 | 1127 | 1263 | 1193 | 15868 | 1.405906736 | 0.003770824 | 0.005301427 |
| Coxiella_burnetii | 6.66E-06 | 6.27E-06 | 7.06E-06 | 1030 | 1160 | 1093 | 12168 | 6.850934893 | 0.001886554 | 0.012924658 |
| Mycobacterium_colombiense | 6.16E-06 | 5.79E-06 | 6.55E-06 | 952 | 1076 | 1012 | 15088 | 1.487200334 | 0.002924245 | 0.004348939 |
| Mycobacterium_intracellulare | 5.94E-06 | 5.57E-06 | 6.32E-06 | 916 | 1038 | 975 | 14884 | 1.489692361 | 0.002827245 | 0.004211726 |
| Trypanosoma_cruzi | 5.83E-06 | 5.46E-06 | 6.20E-06 | 898 | 1020 | 957 | 32835 | 3.213584329 | 0.000211947 | 0.00068111 |
| Mycobacterium_kansasii | 5.63E-06 | 5.28E-06 | 6.00E-06 | 867 | 987 | 925 | 13288 | 1.555288413 | 0.002192413 | 0.003409834 |
| Mycobacteroides_abscessus | 5.26E-06 | 4.92E-06 | 5.62E-06 | 808 | 924 | 864 | 11895 | 1.510160825 | 0.002684541 | 0.004054088 |
| Salmonella_enterica_enterica | 5.21E-06 | 4.87E-06 | 5.57E-06 | 801 | 915 | 856 | 17262 | 1.258467175 | 0.005022576 | 0.006320747 |
| Bacillus_cereus | 3.77E-06 | 3.48E-06 | 4.07E-06 | 572 | 670 | 619 | 9731 | 1.042563711 | 0.003811737 | 0.003973979 |
| Legionella_pneumophila | 3.58E-06 | 3.29E-06 | 3.87E-06 | 541 | 636 | 587 | 13857 | 3.724619008 | 0.001610126 | 0.005997105 |
| Blastomyces_dermatitidis | 3.24E-06 | 2.97E-06 | 3.52E-06 | 489 | 579 | 532 | 30823 | 1.758937106 | 0.000145131 | 0.000255277 |
| Burkholderia_mallei | 2.75E-06 | 2.50E-06 | 3.01E-06 | 411 | 495 | 451 | 45766 | 1.864713417 | 0.001116943 | 0.002082779 |
| Moraxella_catarrhalis | 2.72E-06 | 2.47E-06 | 2.98E-06 | 407 | 489 | 446 | 12552 | 2.863014767 | 0.003138811 | 0.008986461 |
| Francisella_tularensis | 2.69E-06 | 2.45E-06 | 2.95E-06 | 403 | 485 | 442 | 10064 | 5.066954644 | 0.001732964 | 0.008780851 |
| Trichinella_spiralis | 2.52E-06 | 2.28E-06 | 2.77E-06 | 375 | 455 | 413 | 14232 | 1.60951583 | 0.000147665 | 0.000237669 |
| Taylorella_equigenitalis | 1.82E-06 | 1.62E-06 | 2.03E-06 | 266 | 334 | 298 | 11517 | 2.451275343 | 0.002637641 | 0.006465584 |
| Histoplasma_capsulatum | 1.59E-06 | 1.40E-06 | 1.79E-06 | 230 | 294 | 260 | 30338 | 1.622758835 | 0.000156413 | 0.000253821 |
| Klebsiella_pneumoniae_rhinoscleroma | 1.39E-06 | 1.22E-06 | 1.58E-06 | 200 | 260 | 228 | 18871 | 1.27889586 | 0.001238296 | 0.001583652 |
| Coccidioides_immitis | 1.39E-06 | 1.22E-06 | 1.58E-06 | 200 | 260 | 228 | 24458 | 1.848193946 | 0.000121174 | 0.000223954 |
| Bacillus_anthracis | 1.39E-06 | 1.21E-06 | 1.57E-06 | 199 | 259 | 227 | 9531 | 1.387065888 | 0.001188788 | 0.001648927 |
| Corynebacterium_diphtheriae | 1.36E-06 | 1.19E-06 | 1.55E-06 | 196 | 254 | 223 | 8028 | 2.908717853 | 0.001046992 | 0.003045403 |
| Neisseria_gonorrhoeae | 1.36E-06 | 1.19E-06 | 1.55E-06 | 196 | 254 | 223 | 12104 | 1.602585691 | 0.001538387 | 0.002465397 |
| Trypanosoma_brucei | 1.27E-06 | 1.10E-06 | 1.44E-06 | 181 | 237 | 207 | 21969 | 1.366814807 | 0.000190984 | 0.00026104 |
| Neisseria_meningitidis | 1.22E-06 | 1.06E-06 | 1.40E-06 | 174 | 230 | 200 | 12131 | 1.566712899 | 0.001670959 | 0.002617913 |
| Aspergillus_fumigatus | 1.13E-06 | 9.75E-07 | 1.30E-06 | 160 | 214 | 185 | 26528 | 2.302521008 | 7.29E-05 | 0.000167841 |
| Veillonella_parvula | 1.11E-06 | 9.52E-07 | 1.27E-06 | 157 | 209 | 181 | 6170 | 2.99280271 | 0.001107806 | 0.003315445 |
| Escherichia_coli | 9.31E-07 | 7.89E-07 | 1.08E-06 | 130 | 178 | 152 | 16737 | 1.415685604 | 0.000702636 | 0.000994711 |
| Staphylococcus_aureus | 8.27E-07 | 6.94E-07 | 9.72E-07 | 114 | 160 | 135 | 7469 | 1.449906989 | 0.001333753 | 0.001933818 |
| Coccidioides_posadasii | 7.42E-07 | 6.16E-07 | 8.80E-07 | 101 | 145 | 121 | 24581 | 1.514135458 | 9.83E-05 | 0.000148816 |
| Vibrio_parahaemolyticus | 7.12E-07 | 5.89E-07 | 8.46E-07 | 97 | 139 | 116 | 11910 | 1.685372585 | 0.000420847 | 0.000709284 |
| Escherichia_coli_O26 | 6.81E-07 | 5.61E-07 | 8.13E-07 | 92 | 134 | 111 | 13407 | 573.8685688 | 4.90E-05 | 0.028110692 |
| Mycobacterium_leprae | 6.81E-07 | 5.61E-07 | 8.13E-07 | 92 | 134 | 111 | 8799 | 1.254405286 | 0.000569795 | 0.000714754 |
| Helicobacter_pylori | 6.51E-07 | 5.33E-07 | 7.80E-07 | 88 | 128 | 106 | 5256 | 2.854791354 | 0.000731936 | 0.002089526 |
| Haemophilus_influenzae | 5.96E-07 | 4.84E-07 | 7.20E-07 | 80 | 118 | 97 | 10394 | 2.682220434 | 0.000673253 | 0.001805814 |
| Yersinia_enterocolitica | 5.84E-07 | 4.73E-07 | 7.06E-07 | 78 | 116 | 95 | 1735 | 1.109569459 | 0.000532006 | 0.000590297 |
| Tannerella_forsythia_92A2 | 5.72E-07 | 4.62E-07 | 6.93E-07 | 76 | 114 | 93 | 3701 | 1.485100485 | 0.000423724 | 0.000629272 |
| Corynebacterium_pseudotuberculosis | 5.66E-07 | 4.57E-07 | 6.86E-07 | 75 | 113 | 92 | 7926 | 1.770444763 | 0.000869467 | 0.001539344 |
| Shigella_dysenteriae | 5.66E-07 | 4.57E-07 | 6.86E-07 | 75 | 113 | 92 | 17112 | 1.158334785 | 0.000600269 | 0.000695312 |
| Vibrio_cholerae | 5.60E-07 | 4.51E-07 | 6.80E-07 | 74 | 112 | 91 | 11967 | 1.292248988 | 0.000372607 | 0.000481501 |
| Fusobacterium_nucleatum | 5.29E-07 | 4.24E-07 | 6.46E-07 | 70 | 106 | 86 | 5071 | 2.068641618 | 0.000634833 | 0.001313242 |
| Vibrio_vulnificus | 5.11E-07 | 4.08E-07 | 6.26E-07 | 67 | 103 | 83 | 12110 | 1.293976063 | 0.000310675 | 0.000402006 |
| Peptostreptococcus_anaerobius | 4.56E-07 | 3.59E-07 | 5.65E-07 | 59 | 93 | 74 | 5790 | 33.12958484 | 0.000371083 | 0.012293825 |
| Mycobacterium_canettii | 3.77E-07 | 2.89E-07 | 4.77E-07 | 48 | 78 | 61 | 11793 | 1.34153264 | 0.000235829 | 0.000316372 |
| Cryptococcus_gattii | 3.53E-07 | 2.68E-07 | 4.49E-07 | 44 | 74 | 57 | 8898 | 1.726769249 | 7.96E-05 | 0.000137486 |
| Cryptococcus_neoformans | 3.35E-07 | 2.52E-07 | 4.29E-07 | 41 | 70 | 54 | 6661 | 2.356947182 | 5.73E-05 | 0.00013497 |
| Treponema_denticola_ATCC_35405 | 3.10E-07 | 2.31E-07 | 4.01E-07 | 38 | 66 | 50 | 3797 | 1.669565217 | 0.000323579 | 0.000540236 |

| | | | | | | | | | | |
|---|---|---|---|---|---|---|---|---|---|---|
| Escherichia_coli_O157-H7 | 2.74E-07 | 2.00E-07 | 3.59E-07 | 33 | 59 | 44 | 17303 | 1.06765659 | 0.000213777 | 0.000228241 |
| Bartonella_bacilliformis | 2.55E-07 | 1.84E-07 | 3.38E-07 | 30 | 56 | 41 | 8986 | 1.477 | 0.000692031 | 0.00102213 |
| Bartonella_henselae | 2.55E-07 | 1.84E-07 | 3.38E-07 | 30 | 56 | 41 | 8959 | 2.199255121 | 0.000281833 | 0.000619823 |
| Actinobacillus_equuli_equuli | 2.49E-07 | 1.79E-07 | 3.31E-07 | 29 | 54 | 40 | 10509 | 1.303064699 | 0.000362323 | 0.00047213 |
| Shigella_flexneri | 2.49E-07 | 1.79E-07 | 3.31E-07 | 29 | 54 | 40 | 16854 | 1.223083549 | 0.000240431 | 0.000294068 |
| Bartonella_quintana | 2.37E-07 | 1.69E-07 | 3.17E-07 | 28 | 52 | 38 | 8909 | 3.005586592 | 0.00032543 | 0.000978108 |
| Clostridium_perfringens | 2.25E-07 | 1.58E-07 | 3.03E-07 | 26 | 50 | 36 | 5615 | 1.351844478 | 0.000275689 | 0.000372689 |
| Parvimonas_micra | 2.19E-07 | 1.53E-07 | 2.96E-07 | 25 | 49 | 35 | 5546 | 1.769874477 | 0.000431444 | 0.000763601 |
| Leptospira_borgpetersenii | 2.13E-07 | 1.48E-07 | 2.89E-07 | 24 | 48 | 34 | 5405 | 3.848532419 | 4.66E-05 | 0.00017928 |
| Shigella_boydii | 1.89E-07 | 1.28E-07 | 2.61E-07 | 21 | 43 | 30 | 17015 | 8.307542971 | 0.000236042 | 0.001960932 |
| Leptospira_inadai | 1.89E-07 | 1.28E-07 | 2.61E-07 | 21 | 43 | 30 | 5244 | 2.898395968 | 0.000114629 | 0.000332239 |
| Brucella_microti | 1.82E-07 | 1.23E-07 | 2.53E-07 | 20 | 42 | 29 | 12801 | 2.064516129 | 0.000139331 | 0.000287652 |
| Leptospira_kmetyi | 1.64E-07 | 1.08E-07 | 2.32E-07 | 18 | 38 | 26 | 5424 | 1.807432432 | 6.70E-05 | 0.000121125 |
| Klebsiella_pneumoniae_pneumoniae | 1.46E-07 | 9.35E-08 | 2.10E-07 | 15 | 35 | 23 | 18353 | 1.044560615 | 8.32E-05 | 8.69E-05 |
| Leptospira_weilii | 1.28E-07 | 7.91E-08 | 1.88E-07 | 13 | 31 | 20 | 5610 | 1.629250668 | 8.03E-05 | 0.000130894 |
| Yersinia_pseudotuberculosis | 1.28E-07 | 7.91E-08 | 1.88E-07 | 13 | 31 | 20 | 13660 | 1.504372621 | 8.58E-05 | 0.000129006 |
| Leptospira_santarosai | 1.22E-07 | 7.43E-08 | 1.80E-07 | 12 | 30 | 19 | 5385 | 1.253276585 | 9.61E-05 | 0.000120495 |
| Shigella_sonnei | 1.22E-07 | 7.43E-08 | 1.80E-07 | 12 | 30 | 19 | 17231 | 1 | 0.000118418 | 0.000118418 |
| Leptospira_wolffii | 1.16E-07 | 6.96E-08 | 1.73E-07 | 11 | 28 | 18 | 5432 | 1.777126266 | 7.60E-05 | 0.000134977 |
| Leptospira_licerasiae | 1.09E-07 | 6.49E-08 | 1.66E-07 | 11 | 27 | 17 | 5357 | 2.938290987 | 8.41E-05 | 0.000247068 |
| Leptospira_broomii | 9.73E-08 | 5.56E-08 | 1.51E-07 | 9 | 25 | 15 | 5226 | 1.948019932 | 5.87E-05 | 0.000114331 |
| Leptospira_fainei | 9.73E-08 | 5.56E-08 | 1.51E-07 | 9 | 25 | 15 | 5224 | 1.844771974 | 6.16E-05 | 0.000113595 |
| Clostridium_botulinum_E3 | 9.12E-08 | 5.11E-08 | 1.43E-07 | 8 | 23 | 14 | 5652 | 1.02742616 | 0.000129521 | 0.000133073 |
| Tateropox | 8.52E-08 | 4.66E-08 | 1.35E-07 | 8 | 22 | 13 | 42 | 1.801886792 | 0.001070437 | 0.001928806 |
| Leptospira_alstonii | 8.52E-08 | 4.66E-08 | 1.35E-07 | 8 | 22 | 13 | 5348 | 9.57552867 | 4.94E-05 | 0.000473386 |
| Pneumocystis_carinii | 8.52E-08 | 4.66E-08 | 1.35E-07 | 8 | 22 | 13 | 15714 | 3.674162572 | 3.65E-05 | 0.000134278 |
| Brucella_melitensis | 7.91E-08 | 4.21E-08 | 1.28E-07 | 7 | 21 | 12 | 12702 | 2.498862898 | 4.31E-05 | 0.000107692 |
| Streptococcus_salivarius | 7.30E-08 | 3.77E-08 | 1.20E-07 | 6 | 20 | 11 | 7247 | 1.444196429 | 0.000205027 | 0.0002961 |
| Borrelia_recurrentis | 7.30E-08 | 3.77E-08 | 1.20E-07 | 6 | 20 | 11 | 3394 | 1.788465071 | 7.57E-05 | 0.000135341 |
| Leptospira_alexanderi | 7.30E-08 | 3.77E-08 | 1.20E-07 | 6 | 20 | 11 | 5483 | 47.54347214 | 3.84E-05 | 0.001826362 |
| Pneumocystis_jirovecii | 7.30E-08 | 3.77E-08 | 1.20E-07 | 6 | 20 | 11 | 15151 | 2.875867856 | 3.45E-05 | 9.93E-05 |
| Human_herpesvirus_6B | 6.69E-08 | 3.34E-08 | 1.12E-07 | 5 | 18 | 10 | 12101 | 1.393333333 | 0.000925275 | 0.001289216 |
| Streptococcus_pneumoniae | 6.69E-08 | 3.34E-08 | 1.12E-07 | 5 | 18 | 10 | 7220 | 1.663819112 | 0.000123935 | 0.000206205 |
| Streptococcus_sanguinis | 6.08E-08 | 2.92E-08 | 1.04E-07 | 5 | 17 | 9 | 7521 | 1.074170614 | 0.00014489 | 0.000155637 |
| Streptococcus_gordonii_str_Challis | 6.08E-08 | 2.92E-08 | 1.04E-07 | 5 | 17 | 9 | 7394 | 1 | 0.000122914 | 0.000122914 |
| Streptococcus_pyogenes | 5.47E-08 | 2.50E-08 | 9.59E-08 | 4 | 16 | 8 | 7413 | 1.075067024 | 0.000213585 | 0.000229618 |
| Yersinia_pestis | 5.47E-08 | 2.50E-08 | 9.59E-08 | 4 | 16 | 8 | 13552 | 2.998179302 | 6.64E-05 | 0.000198953 |
| Chlamydia_pneumoniae | 5.47E-08 | 2.50E-08 | 9.59E-08 | 4 | 16 | 8 | 2719 | 1.190612532 | 0.000135357 | 0.000161158 |
| Porphyromonas_gingivalis_W83 | 5.47E-08 | 2.50E-08 | 9.59E-08 | 4 | 16 | 8 | 3701 | 1.568047337 | 7.21E-05 | 0.00011308 |
| Streptococcus_oralis | 4.87E-08 | 2.10E-08 | 8.77E-08 | 3 | 14 | 7 | 7426 | 1 | 0.000132536 | 0.000132536 |
| Escherichia_coli_O111 | 4.87E-08 | 2.10E-08 | 8.77E-08 | 3 | 14 | 7 | 17934 | 38.75628882 | 4.31E-05 | 0.00167039 |
| Brucella_ovis | 4.87E-08 | 2.10E-08 | 8.77E-08 | 3 | 14 | 7 | 12806 | 1.609375 | 3.91E-05 | 6.29E-05 |
| Cowpox_virus | 4.26E-08 | 1.71E-08 | 7.94E-08 | 3 | 13 | 6 | 243 | 1.555555556 | 0.000801785 | 0.001247222 |
| Methanobrevibacter_oralis | 4.26E-08 | 1.71E-08 | 7.94E-08 | 3 | 13 | 6 | 186 | 2.557698676 | 4.55E-05 | 0.000116501 |
| Rickettsia_prowazekii | 4.26E-08 | 1.71E-08 | 7.94E-08 | 3 | 13 | 6 | 5915 | 1.259259259 | 4.87E-05 | 6.13E-05 |
| Rickettsia_felis | 3.65E-08 | 1.34E-08 | 7.10E-08 | 2 | 12 | 5 | 5924 | 5.053252563 | 6.14E-05 | 0.000310408 |
| Mycoplasma_pneumoniae | 3.65E-08 | 1.34E-08 | 7.10E-08 | 2 | 12 | 5 | 1696 | 2.388888889 | 8.39E-05 | 0.000200513 |
| Leptospira_kirschneri | 3.65E-08 | 1.34E-08 | 7.10E-08 | 2 | 12 | 5 | 5374 | 7.357596585 | 2.25E-05 | 0.000165192 |
| Streptococcus_agalactiae | 3.65E-08 | 1.34E-08 | 7.10E-08 | 2 | 12 | 5 | 7416 | 1.002383594 | 9.52E-05 | 9.55E-05 |
| Streptococcus_mutans | 3.65E-08 | 1.34E-08 | 7.10E-08 | 2 | 12 | 5 | 7037 | 1.006271233 | 6.26E-05 | 6.30E-05 |
| Chlamydia_muridarum | 3.04E-08 | 9.88E-09 | 6.23E-08 | 2 | 10 | 4 | 2739 | 1.176829268 | 0.000150344 | 0.000176929 |
| Marseilleviridae_tokyovirus | 3.04E-08 | 9.88E-09 | 6.23E-08 | 2 | 10 | 4 | 10 | 1.818181818 | 8.85E-05 | 0.000160984 |
| Brucella_abortus | 3.04E-08 | 9.88E-09 | 6.23E-08 | 2 | 10 | 4 | 12770 | 2.290474787 | 1.71E-05 | 3.91E-05 |
| Equid_gammaherpesvirus_2 | 2.43E-08 | 6.63E-09 | 5.33E-08 | 1 | 9 | 3 | 928 | 1 | 0.000433748 | 0.000433748 |
| Streptococcus_equi_zooepidemicus | 2.43E-08 | 6.63E-09 | 5.33E-08 | 1 | 9 | 3 | 7217 | 1 | 0.000121052 | 0.000121052 |
| Streptococcus_gordonii | 2.43E-08 | 6.63E-09 | 5.33E-08 | 1 | 9 | 3 | 7313 | 1 | 6.68E-05 | 6.68E-05 |
| Clostridium_botulinum | 2.43E-08 | 6.63E-09 | 5.33E-08 | 1 | 9 | 3 | 5466 | 1.004204876 | 3.59E-05 | 3.60E-05 |
| Leptospira_noguchii | 2.43E-08 | 6.63E-09 | 5.33E-08 | 1 | 9 | 3 | 5322 | 13.75309676 | 1.40E-05 | 0.000192681 |
| Streptococcus_dysgalactiae | 2.43E-08 | 6.63E-09 | 5.33E-08 | 1 | 9 | 3 | 7447 | 1.004616367 | 2.60E-05 | 2.62E-05 |
| Borreliella_burgdorferi | 1.82E-08 | 3.76E-09 | 4.39E-08 | 1 | 7 | 2 | 3666 | 7.359371345 | 2.12E-05 | 0.000155938 |
| Streptococcus_anginosus | 1.82E-08 | 3.76E-09 | 4.39E-08 | 1 | 7 | 2 | 8102 | 1 | 3.59E-05 | 3.59E-05 |
| Borreliella_afzelii | 1.82E-08 | 3.76E-09 | 4.39E-08 | 1 | 7 | 2 | 3701 | 1 | 3.16E-05 | 3.16E-05 |
| Chlamydia_felis | 1.82E-08 | 3.76E-09 | 4.39E-08 | 1 | 7 | 2 | 2934 | 1.006475517 | 2.56E-05 | 2.57E-05 |
| Porphyromonas_gingivalis | 1.82E-08 | 3.76E-09 | 4.39E-08 | 1 | 7 | 2 | 3700 | 1 | 2.55E-05 | 2.55E-05 |

| | | | | | | | | | | |
|---|---|---|---|---|---|---|---|---|---|---|
| Chlamydia_psittaci | 1.82E-08 | 3.76E-09 | 4.39E-08 | 1 | 7 | 2 | 2986 | 1.006446371 | 2.46E-05 | 2.48E-05 |
| Mycobacterium_bovis | 1.82E-08 | 3.76E-09 | 4.39E-08 | 1 | 7 | 2 | 11831 | 1 | 1.27E-05 | 1.27E-05 |
| Rickettsia_akari | 1.22E-08 | 1.47E-09 | 3.39E-08 | 0 | 6 | 1 | 5882 | 1 | 3.49E-05 | 3.49E-05 |
| Chlamydia_pecorum | 1.22E-08 | 1.47E-09 | 3.39E-08 | 0 | 6 | 1 | 2754 | 1.006821364 | 2.60E-05 | 2.62E-05 |
| Rickettsia_sibirica | 1.22E-08 | 1.47E-09 | 3.39E-08 | 0 | 6 | 1 | 5910 | 1 | 2.24E-05 | 2.24E-05 |
| Mycobacterium_tuberculosis | 1.22E-08 | 1.47E-09 | 3.39E-08 | 0 | 6 | 1 | 11841 | 1 | 7.48E-06 | 7.48E-06 |
| Clostridium_sporogenes | 1.22E-08 | 1.47E-09 | 3.39E-08 | 0 | 6 | 1 | 5519 | 1.01081801 | 6.83E-06 | 6.90E-06 |
| Leptospira_interrogans | 1.22E-08 | 1.47E-09 | 3.39E-08 | 0 | 6 | 1 | 5378 | 1.081806866 | 5.83E-06 | 6.31E-06 |
| Human_gammaherpesvirus_4 | 6.08E-09 | 1.54E-10 | 2.24E-08 | 0 | 4 | 0 | 3 | 0 | 0 | 0 |
| Human_alphaherpesvirus_1 | 6.08E-09 | 1.54E-10 | 2.24E-08 | 0 | 4 | 0 | 4 | 0 | 0 | 0 |
| Human_adenovirus_D9 | 6.08E-09 | 1.54E-10 | 2.24E-08 | 0 | 4 | 0 | 1 | 0 | 0 | 0 |
| Chlamydia_caviae | 6.08E-09 | 1.54E-10 | 2.24E-08 | 0 | 4 | 0 | 2940 | 0 | 0 | 0 |
| Chlamydia_abortus | 6.08E-09 | 1.54E-10 | 2.24E-08 | 0 | 4 | 0 | 2975 | 0 | 0 | 0 |
| Human_herpesvirus_1 | 6.08E-09 | 1.54E-10 | 2.24E-08 | 0 | 4 | 0 | 4 | 0 | 0 | 0 |
| Human_adenovirus_D37 | 6.08E-09 | 1.54E-10 | 2.24E-08 | 0 | 4 | 0 | 1 | 0 | 0 | 0 |
| Human_adenovirus_71 | 6.08E-09 | 1.54E-10 | 2.24E-08 | 0 | 4 | 0 | 1 | 0 | 0 | 0 |
| Brucella_suis | 6.08E-09 | 1.54E-10 | 2.24E-08 | 0 | 4 | 0 | 12694 | 0 | 0 | 0 |
| Treponema_pallidum_subsp_pertenue | 6.08E-09 | 1.54E-10 | 2.24E-08 | 0 | 4 | 0 | 4083 | 0 | 0 | 0 |
| Human_herpesvirus_3_strain_Dumas | 6.08E-09 | 1.54E-10 | 2.24E-08 | 0 | 4 | 0 | 3 | 0 | 0 | 0 |
| Treponema_pallidum_subsp_pallidum | 6.08E-09 | 1.54E-10 | 2.24E-08 | 0 | 4 | 0 | 4083 | 0 | 0 | 0 |
| Human_adenovirus_67 | 6.08E-09 | 1.54E-10 | 2.24E-08 | 0 | 4 | 0 | 1 | 0 | 0 | 0 |
| Borreliella_garinii | 6.08E-09 | 1.54E-10 | 2.24E-08 | 0 | 4 | 0 | 3689 | 0 | 0 | 0 |
| Mycobacterium_africanum | 6.08E-09 | 1.54E-10 | 2.24E-08 | 0 | 4 | 0 | 11825 | 0 | 0 | 0 |
| Rickettsia_africae | 6.08E-09 | 1.54E-10 | 2.24E-08 | 0 | 4 | 0 | 5924 | 0 | 0 | 0 |
| Yersinia_pestis_CO92 | 6.08E-09 | 1.54E-10 | 2.24E-08 | 0 | 4 | 0 | 13548 | 0 | 0 | 0 |
| Rickettsia_conorii | 6.08E-09 | 1.54E-10 | 2.24E-08 | 0 | 4 | 0 | 5925 | 0 | 0 | 0 |
| Rickettsia_japonica | 6.08E-09 | 1.54E-10 | 2.24E-08 | 0 | 4 | 0 | 5901 | 0 | 0 | 0 |
| Rickettsia_rickettsii | 6.08E-09 | 1.54E-10 | 2.24E-08 | 0 | 4 | 0 | 5918 | 0 | 0 | 0 |
| Rickettsia_typhi | 6.08E-09 | 1.54E-10 | 2.24E-08 | 0 | 4 | 0 | 5908 | 0 | 0 | 0 |
| Human_adenovirus_69 | 6.08E-09 | 1.54E-10 | 2.24E-08 | 0 | 4 | 0 | 1 | 0 | 0 | 0 |
| Chlamydia_trachomatis | 6.08E-09 | 1.54E-10 | 2.24E-08 | 0 | 4 | 0 | 2724 | 0 | 0 | 0 |
| Human_adenovirus_1 | 6.08E-09 | 1.54E-10 | 2.24E-08 | 0 | 4 | 0 | 4 | 0 | 0 | 0 |
| Human_adenovirus_66 | 6.08E-09 | 1.54E-10 | 2.24E-08 | 0 | 4 | 0 | 1 | 0 | 0 | 0 |
| Human_adenovirus_17 | 6.08E-09 | 1.54E-10 | 2.24E-08 | 0 | 4 | 0 | 1 | 0 | 0 | 0 |
| Human_adenovirus_20 | 6.08E-09 | 1.54E-10 | 2.24E-08 | 0 | 4 | 0 | 1 | 0 | 0 | 0 |
| Human_adenovirus_23 | 6.08E-09 | 1.54E-10 | 2.24E-08 | 0 | 4 | 0 | 1 | 0 | 0 | 0 |
| Human_adenovirus_25 | 6.08E-09 | 1.54E-10 | 2.24E-08 | 0 | 4 | 0 | 1 | 0 | 0 | 0 |
| Human_adenovirus_26 | 6.08E-09 | 1.54E-10 | 2.24E-08 | 0 | 4 | 0 | 1 | 0 | 0 | 0 |
| Equid_gammaherpesvirus_5 | 6.08E-09 | 1.54E-10 | 2.24E-08 | 0 | 4 | 0 | 2425 | 0 | 0 | 0 |
| Human_adenovirus_29 | 6.08E-09 | 1.54E-10 | 2.24E-08 | 0 | 4 | 0 | 1 | 0 | 0 | 0 |
| Equid_alphaherpesvirus_9 | 6.08E-09 | 1.54E-10 | 2.24E-08 | 0 | 4 | 0 | 185 | 0 | 0 | 0 |
| Equid_alphaherpesvirus_8 | 6.08E-09 | 1.54E-10 | 2.24E-08 | 0 | 4 | 0 | 195 | 0 | 0 | 0 |
| Variola | 6.08E-09 | 1.54E-10 | 2.24E-08 | 0 | 4 | 0 | 2 | 0 | 0 | 0 |
| Equid_alphaherpesvirus_3 | 6.08E-09 | 1.54E-10 | 2.24E-08 | 0 | 4 | 0 | 12074 | 0 | 0 | 0 |
| Equid_alphaherpesvirus_1 | 6.08E-09 | 1.54E-10 | 2.24E-08 | 0 | 4 | 0 | 193 | 0 | 0 | 0 |
| Eastern_equine_encephalitis | 6.08E-09 | 1.54E-10 | 2.24E-08 | 0 | 4 | 0 | 12 | 0 | 0 | 0 |
| Human_adenovirus_32 | 6.08E-09 | 1.54E-10 | 2.24E-08 | 0 | 4 | 0 | 1 | 0 | 0 | 0 |
| Human_adenovirus_38 | 6.08E-09 | 1.54E-10 | 2.24E-08 | 0 | 4 | 0 | 1 | 0 | 0 | 0 |
| Human_adenovirus_40 | 6.08E-09 | 1.54E-10 | 2.24E-08 | 0 | 4 | 0 | 1 | 0 | 0 | 0 |
| Human_adenovirus_41 | 6.08E-09 | 1.54E-10 | 2.24E-08 | 0 | 4 | 0 | 26 | 0 | 0 | 0 |
| Human_adenovirus_42 | 6.08E-09 | 1.54E-10 | 2.24E-08 | 0 | 4 | 0 | 1 | 0 | 0 | 0 |
| Human_adenovirus_45 | 6.08E-09 | 1.54E-10 | 2.24E-08 | 0 | 4 | 0 | 1 | 0 | 0 | 0 |
| Human_adenovirus_49 | 6.08E-09 | 1.54E-10 | 2.24E-08 | 0 | 4 | 0 | 1 | 0 | 0 | 0 |
| Venezuelan_equine_encephalitis | 6.08E-09 | 1.54E-10 | 2.24E-08 | 0 | 4 | 0 | 26 | 0 | 0 | 0 |
| Clostridium_tetani_E88 | 6.08E-09 | 1.54E-10 | 2.24E-08 | 0 | 4 | 0 | 5876 | 0 | 0 | 0 |
| Clostridium_tetani | 6.08E-09 | 1.54E-10 | 2.24E-08 | 0 | 4 | 0 | 5867 | 0 | 0 | 0 |
| Human_adenovirus_51 | 6.08E-09 | 1.54E-10 | 2.24E-08 | 0 | 4 | 0 | 1 | 0 | 0 | 0 |
| Human_adenovirus_53 | 6.08E-09 | 1.54E-10 | 2.24E-08 | 0 | 4 | 0 | 1 | 0 | 0 | 0 |
| Human_adenovirus_46 | 6.08E-09 | 1.54E-10 | 2.24E-08 | 0 | 4 | 0 | 1 | 0 | 0 | 0 |

**Table S2d**: HAYSTAC results for library ERR6466111 based on a custom-built pathogen database (Table S1)

| Taxon | Mean_Posterior_Abundance | 95_CI_lower | 95_CI_upper | Minimum_Read_Num | Maximum_Read_Num | Dirichlet_Read_Num | Aligned_Read_Num | Evenness_of_Coverage_Ratio | Fraction_of_Genome_Covered | Coverage |
|---|---|---|---|---|---|---|---|---|---|---|
| Dark_Matter | 0.998660508 | 0.998654473 | 0.99866653 | 141194536 | 141196241 | 141195554 | 0 | | | |
| Grey_Matter | 0.000897224 | 0.000892296 | 0.000902166 | 126157 | 127553 | 126853 | 0 | | | |
| Burkholderia_cepacia | 8.55E-05 | 8.40E-05 | 8.71E-05 | 11878 | 12309 | 12092 | 48048 | 1.55250182 | 0.02741444 | 0.042560968 |
| Plasmodium_vivax | 5.58E-05 | 5.45E-05 | 5.70E-05 | 7710 | 8058 | 7882 | 24487 | 13.32353314 | 0.00051137 | 0.006813259 |
| Equid_alphaherpesvirus_4 | 3.67E-05 | 3.57E-05 | 3.77E-05 | 5045 | 5327 | 5184 | 5254 | 1.71849943 | 0.645006422 | 1.108443168 |
| Bordetella_pertussis | 3.46E-05 | 3.37E-05 | 3.56E-05 | 4758 | 5032 | 4893 | 27284 | 2.024842696 | 0.015003787 | 0.030380309 |
| Bordetella_petrii | 2.92E-05 | 2.83E-05 | 3.01E-05 | 4008 | 4260 | 4132 | 25811 | 1.800812803 | 0.011260886 | 0.020278747 |
| Taenia_solium | 2.62E-05 | 2.54E-05 | 2.71E-05 | 3588 | 3826 | 3705 | 30732 | 14.67692678 | 0.000143563 | 0.002107058 |
| Prescottella_equi | 1.86E-05 | 1.79E-05 | 1.94E-05 | 2536 | 2738 | 2635 | 14716 | 1.806393431 | 0.007747444 | 0.013994932 |
| Toxoplasma_gondii | 1.70E-05 | 1.64E-05 | 1.77E-05 | 2313 | 2505 | 2407 | 12080 | 2.590012547 | 0.000344825 | 0.000893101 |
| Burkholderia_pseudomallei | 8.27E-06 | 7.80E-06 | 8.75E-06 | 1103 | 1237 | 1168 | 43814 | 1.592482952 | 0.002669857 | 0.004251702 |
| Plasmodium_falciparum | 8.04E-06 | 7.58E-06 | 8.52E-06 | 1072 | 1204 | 1136 | 12798 | 1.845801458 | 0.000732203 | 0.001351501 |
| Schistosoma_mansoni | 7.66E-06 | 7.21E-06 | 8.12E-06 | 1019 | 1148 | 1082 | 22928 | 1.771833344 | 6.73E-05 | 0.000119309 |
| Mycobacterium_avium | 7.29E-06 | 6.85E-06 | 7.74E-06 | 968 | 1094 | 1029 | 14371 | 1.371514819 | 0.003559814 | 0.004882338 |
| Mycobacterium_colombiense | 7.16E-06 | 6.72E-06 | 7.61E-06 | 951 | 1075 | 1011 | 14050 | 1.487655533 | 0.002942885 | 0.004377999 |
| Klebsiella_oxytoca | 7.03E-06 | 6.60E-06 | 7.47E-06 | 933 | 1057 | 993 | 15521 | 1.182292141 | 0.00461314 | 0.005454079 |
| Coxiella_burnetii | 6.77E-06 | 6.35E-06 | 7.20E-06 | 897 | 1019 | 956 | 10951 | 7.071946847 | 0.001691257 | 0.011960483 |
| Trypanosoma_cruzi | 6.25E-06 | 5.85E-06 | 6.67E-06 | 827 | 943 | 883 | 28593 | 2.850280884 | 0.000218674 | 0.000623283 |
| Mycobacterium_kansasii | 6.13E-06 | 5.73E-06 | 6.55E-06 | 810 | 926 | 866 | 12050 | 1.505539553 | 0.002115025 | 0.003184253 |
| Mycobacterium_intracellulare | 6.08E-06 | 5.68E-06 | 6.49E-06 | 803 | 917 | 858 | 13465 | 1.434018914 | 0.002554298 | 0.003662912 |
| Mycobacteroides_abscessus | 5.79E-06 | 5.40E-06 | 6.19E-06 | 763 | 875 | 817 | 10898 | 1.455666264 | 0.002699494 | 0.003929562 |
| Salmonella_enterica_enterica | 5.26E-06 | 4.89E-06 | 5.65E-06 | 691 | 798 | 743 | 15170 | 1.268600314 | 0.004359903 | 0.005530975 |
| Legionella_pneumophila | 4.39E-06 | 4.05E-06 | 4.74E-06 | 573 | 671 | 620 | 12380 | 3.923498954 | 0.001503393 | 0.005898561 |
| Bacillus_cereus | 3.44E-06 | 3.15E-06 | 3.76E-06 | 445 | 531 | 486 | 8577 | 1.042390661 | 0.0031847 | 0.003319701 |
| Francisella_tularensis | 3.41E-06 | 3.11E-06 | 3.72E-06 | 440 | 526 | 481 | 9250 | 4.385379521 | 0.002099234 | 0.009205938 |
| Blastomyces_dermatitidis | 3.15E-06 | 2.87E-06 | 3.45E-06 | 406 | 488 | 445 | 26774 | 1.74641668 | 0.000121441 | 0.000212086 |
| Burkholderia_mallei | 2.96E-06 | 2.69E-06 | 3.25E-06 | 380 | 460 | 418 | 42075 | 1.822248283 | 0.001067408 | 0.001945083 |
| Moraxella_catarrhalis | 2.69E-06 | 2.42E-06 | 2.96E-06 | 343 | 419 | 379 | 11006 | 2.684154751 | 0.002856017 | 0.007665991 |
| Trichinella_spiralis | 2.59E-06 | 2.33E-06 | 2.86E-06 | 329 | 404 | 365 | 12564 | 1.514155187 | 0.000136009 | 0.000205939 |
| Taylorella_equigenitalis | 1.63E-06 | 1.42E-06 | 1.84E-06 | 201 | 261 | 229 | 10161 | 2.975759346 | 0.001968887 | 0.005858933 |
| Histoplasma_capsulatum | 1.59E-06 | 1.39E-06 | 1.81E-06 | 197 | 255 | 224 | 26305 | 1.533363161 | 0.000128267 | 0.00019668 |
| Neisseria_gonorrhoeae | 1.56E-06 | 1.36E-06 | 1.78E-06 | 193 | 251 | 220 | 11045 | 1.795704418 | 0.001334405 | 0.002396196 |
| Corynebacterium_diphtheriae | 1.56E-06 | 1.36E-06 | 1.78E-06 | 193 | 251 | 220 | 7165 | 2.892148053 | 0.001123439 | 0.003249151 |
| Veillonella_parvula | 1.41E-06 | 1.23E-06 | 1.62E-06 | 173 | 229 | 199 | 5584 | 3.046275395 | 0.001246634 | 0.003797589 |
| Klebsiella_pneumoniae_rhinoscleroma | 1.37E-06 | 1.18E-06 | 1.56E-06 | 167 | 221 | 192 | 16812 | 1.220146509 | 0.001166579 | 0.001423397 |
| Trypanosoma_brucei | 1.26E-06 | 1.08E-06 | 1.45E-06 | 153 | 205 | 177 | 19341 | 1.294724502 | 0.000163564 | 0.00021177 |
| Neisseria_meningitidis | 1.23E-06 | 1.05E-06 | 1.42E-06 | 149 | 201 | 173 | 11057 | 1.432315522 | 0.0018216 | 0.002609106 |
| Bacillus_anthracis | 1.12E-06 | 9.57E-07 | 1.31E-06 | 135 | 185 | 158 | 8441 | 1.308373296 | 0.001046889 | 0.001369722 |
| Aspergillus_fumigatus | 1.10E-06 | 9.37E-07 | 1.28E-06 | 132 | 181 | 155 | 22769 | 2.061860465 | 7.32E-05 | 0.000150859 |
| Coccidioides_immitis | 1.06E-06 | 8.98E-07 | 1.24E-06 | 127 | 175 | 149 | 21510 | 1.652094407 | 0.000101496 | 0.00016768 |
| Escherichia_coli | 9.12E-07 | 7.62E-07 | 1.08E-06 | 108 | 152 | 128 | 14617 | 1.426288945 | 0.000653154 | 0.000931587 |
| Coccidioides_posadasii | 8.91E-07 | 7.42E-07 | 1.05E-06 | 105 | 149 | 125 | 21540 | 1.431901615 | 9.92E-05 | 0.000142059 |
| Staphylococcus_aureus | 8.49E-07 | 7.04E-07 | 1.01E-06 | 99 | 142 | 119 | 6528 | 1.56010607 | 0.001202966 | 0.001876754 |
| Mycobacterium_leprae | 8.28E-07 | 6.84E-07 | 9.84E-07 | 97 | 139 | 116 | 7930 | 1.244296578 | 0.000660159 | 0.000821433 |
| Helicobacter_pylori | 7.29E-07 | 5.95E-07 | 8.76E-07 | 84 | 124 | 102 | 4775 | 2.962564883 | 0.000747942 | 0.002215826 |
| Escherichia_coli_O26 | 7.21E-07 | 5.88E-07 | 8.68E-07 | 83 | 123 | 101 | 11840 | 70.83524837 | 8.89E-05 | 0.0062985 |
| Yersinia_enterocolitica | 6.79E-07 | 5.50E-07 | 8.21E-07 | 78 | 116 | 95 | 1485 | 1.128588199 | 0.000476387 | 0.000537645 |
| Vibrio_parahaemolyticus | 6.72E-07 | 5.44E-07 | 8.14E-07 | 77 | 115 | 94 | 10608 | 1.587285044 | 0.000371484 | 0.000589651 |
| Shigella_dysenteriae | 6.51E-07 | 5.25E-07 | 7.90E-07 | 74 | 112 | 91 | 15097 | 1.082036316 | 0.000593914 | 0.000642636 |
| Vibrio_cholerae | 5.80E-07 | 4.61E-07 | 7.12E-07 | 65 | 101 | 81 | 10580 | 1.236042903 | 0.000411511 | 0.000508645 |
| Fusobacterium_nucleatum | 5.52E-07 | 4.36E-07 | 6.81E-07 | 62 | 96 | 77 | 4525 | 2.012065295 | 0.0006463 | 0.001300398 |
| Peptostreptococcus_anaerobius | 5.23E-07 | 4.11E-07 | 6.49E-07 | 58 | 92 | 73 | 5250 | 56.59893959 | 0.000389325 | 0.022035388 |
| Mycobacterium_canettii | 4.74E-07 | 3.67E-07 | 5.94E-07 | 52 | 84 | 66 | 10679 | 1.575413223 | 0.000215972 | 0.000340245 |
| Vibrio_vulnificus | 4.60E-07 | 3.55E-07 | 5.78E-07 | 50 | 82 | 64 | 10772 | 1.227068173 | 0.000289572 | 0.000355325 |
| Parvimonas_micra | 4.53E-07 | 3.49E-07 | 5.70E-07 | 49 | 81 | 63 | 4958 | 1.426760563 | 0.000854463 | 0.001219114 |
| Tannerella_forsythia_92A2 | 4.31E-07 | 3.30E-07 | 5.46E-07 | 47 | 77 | 60 | 3240 | 1.380851064 | 0.000276022 | 0.000381146 |
| Haemophilus_influenzae | 4.24E-07 | 3.24E-07 | 5.38E-07 | 46 | 76 | 59 | 9260 | 2.119337979 | 0.000621798 | 0.0013178 |
| Cryptococcus_gattii | 4.10E-07 | 3.12E-07 | 5.22E-07 | 44 | 74 | 57 | 7697 | 1.410899868 | 8.11E-05 | 0.000114486 |
| Corynebacterium_pseudotuberculosis | 4.10E-07 | 3.12E-07 | 5.22E-07 | 44 | 74 | 57 | 7114 | 1.203134418 | 0.000689836 | 0.000829965 |
| Escherichia_coli_O157-H7 | 3.75E-07 | 2.81E-07 | 4.82E-07 | 40 | 68 | 52 | 15198 | 1.213339083 | 0.000237908 | 0.000288663 |

| | | | | | | | | | | |
|---|---|---|---|---|---|---|---|---|---|---|
| Bartonella_bacilliformis | 3.18E-07 | 2.32E-07 | 4.18E-07 | 33 | 59 | 44 | 7889 | 1.308238636 | 0.00097438 | 0.001274722 |
| Cryptococcus_neoformans | 3.11E-07 | 2.26E-07 | 4.10E-07 | 32 | 58 | 43 | 5746 | 1.666242568 | 5.46E-05 | 9.10E-05 |
| Treponema_denticola_ATCC_35405 | 2.90E-07 | 2.08E-07 | 3.85E-07 | 29 | 54 | 40 | 3247 | 1.568965517 | 0.000285594 | 0.000448087 |
| Leptospira_borgpetersenii | 2.83E-07 | 2.02E-07 | 3.77E-07 | 29 | 53 | 39 | 4633 | 3.14 | 7.51E-05 | 0.000235926 |
| Shigella_flexneri | 2.76E-07 | 1.96E-07 | 3.69E-07 | 28 | 52 | 38 | 14809 | 1 | 0.000289098 | 0.000289098 |
| Leptospira_kmetyi | 2.69E-07 | 1.90E-07 | 3.61E-07 | 27 | 51 | 37 | 4674 | 1.382 | 0.000113201 | 0.000156444 |
| Actinobacillus_equuli_equuli | 2.48E-07 | 1.72E-07 | 3.36E-07 | 24 | 48 | 34 | 9371 | 1.069714286 | 0.000359855 | 0.000384942 |
| Bartonella_henselae | 2.33E-07 | 1.61E-07 | 3.20E-07 | 23 | 45 | 32 | 7907 | 1.785714286 | 0.000396771 | 0.000708519 |
| Bartonella_quintana | 2.33E-07 | 1.61E-07 | 3.20E-07 | 23 | 45 | 32 | 7918 | 2.053304904 | 0.000284221 | 0.000583593 |
| Leptospira_licerasiae | 2.26E-07 | 1.55E-07 | 3.11E-07 | 22 | 44 | 31 | 4596 | 2.105194344 | 9.05E-05 | 0.000190518 |
| Shigella_boydii | 2.12E-07 | 1.43E-07 | 2.95E-07 | 20 | 42 | 29 | 14908 | 7.682196659 | 0.000187135 | 0.001437604 |
| Clostridium_perfringens | 2.12E-07 | 1.43E-07 | 2.95E-07 | 20 | 42 | 29 | 5124 | 1.165347923 | 0.000225314 | 0.00026257 |
| Escherichia_coli_O111 | 1.77E-07 | 1.14E-07 | 2.53E-07 | 16 | 36 | 24 | 15691 | 22.05796732 | 0.000136851 | 0.003018659 |
| Leptospira_broomii | 1.63E-07 | 1.03E-07 | 2.36E-07 | 15 | 33 | 22 | 4513 | 1.741348283 | 7.05E-05 | 0.0001228 |
| Leptospira_weilii | 1.49E-07 | 9.19E-08 | 2.18E-07 | 13 | 31 | 20 | 4811 | 2.214271178 | 5.91E-05 | 0.000130894 |
| Clostridium_botulinum_E3 | 1.41E-07 | 8.64E-08 | 2.10E-07 | 12 | 30 | 19 | 5236 | 1.222596965 | 0.000162038 | 0.000198107 |
| Leptospira_inadai | 1.27E-07 | 7.55E-08 | 1.93E-07 | 11 | 27 | 17 | 4506 | 1.741849496 | 9.71E-05 | 0.000169189 |
| Streptococcus_salivarius | 1.27E-07 | 7.55E-08 | 1.93E-07 | 11 | 27 | 17 | 6403 | 1.558028617 | 0.000287862 | 0.000448497 |
| Shigella_sonnei | 1.27E-07 | 7.55E-08 | 1.93E-07 | 11 | 27 | 17 | 15128 | 1 | 0.000114009 | 0.000114009 |
| Pneumocystis_carinii | 1.27E-07 | 7.55E-08 | 1.93E-07 | 11 | 27 | 17 | 13998 | 2.516615539 | 4.74E-05 | 0.000119237 |
| Yersinia_pseudotuberculosis | 1.20E-07 | 7.00E-08 | 1.84E-07 | 10 | 26 | 16 | 12018 | 2.141692506 | 3.74E-05 | 8.01E-05 |
| Leptospira_santarosai | 1.13E-07 | 6.47E-08 | 1.75E-07 | 9 | 25 | 15 | 4637 | 1.295373665 | 7.05E-05 | 9.14E-05 |
| Pneumocystis_jirovecii | 9.90E-08 | 5.41E-08 | 1.57E-07 | 8 | 22 | 13 | 13560 | 3.032529413 | 2.93E-05 | 8.88E-05 |
| Leptospira_alstonii | 9.90E-08 | 5.41E-08 | 1.57E-07 | 8 | 22 | 13 | 4596 | 22.97415304 | 5.23E-05 | 0.001200675 |
| Streptococcus_pneumoniae | 9.90E-08 | 5.41E-08 | 1.57E-07 | 8 | 22 | 13 | 6456 | 1.242784375 | 0.000108617 | 0.000134987 |
| Klebsiella_pneumoniae_pneumoniae | 9.90E-08 | 5.41E-08 | 1.57E-07 | 8 | 22 | 13 | 16279 | 1.159822475 | 7.66E-05 | 8.88E-05 |
| Leptospira_wolffii | 9.19E-08 | 4.90E-08 | 1.48E-07 | 7 | 21 | 12 | 4645 | 1.3802344 | 6.64E-05 | 9.16E-05 |
| Mycoplasma_pneumoniae | 8.49E-08 | 4.39E-08 | 1.39E-07 | 6 | 20 | 11 | 1602 | 1.603174603 | 0.000220331 | 0.00035323 |
| Tateropox | 8.49E-08 | 4.39E-08 | 1.39E-07 | 6 | 20 | 11 | 38 | 2.564655172 | 0.001171421 | 0.003004292 |
| Streptococcus_sanguinis | 8.49E-08 | 4.39E-08 | 1.39E-07 | 6 | 20 | 11 | 6730 | 1.169077254 | 0.000108341 | 0.00012666 |
| Rickettsia_typhi | 7.78E-08 | 3.88E-08 | 1.30E-07 | 5 | 18 | 10 | 5430 | 1.144736842 | 0.000136645 | 0.000156422 |
| Leptospira_alexanderi | 7.78E-08 | 3.88E-08 | 1.30E-07 | 5 | 18 | 10 | 4700 | 25.75142598 | 5.00E-05 | 0.001287267 |
| Leptospira_fainei | 7.78E-08 | 3.88E-08 | 1.30E-07 | 5 | 18 | 10 | 4515 | 1.335159917 | 7.14E-05 | 9.53E-05 |
| Leptospira_interrogans | 7.78E-08 | 3.88E-08 | 1.30E-07 | 5 | 18 | 10 | 4598 | 1.081806866 | 4.25E-05 | 4.60E-05 |
| Brucella_microti | 7.78E-08 | 3.88E-08 | 1.30E-07 | 5 | 18 | 10 | 11541 | 1.160784314 | 7.64E-05 | 8.87E-05 |
| Brucella_melitensis | 7.78E-08 | 3.88E-08 | 1.30E-07 | 5 | 18 | 10 | 11436 | 1.260416667 | 5.83E-05 | 7.34E-05 |
| Streptococcus_dysgalactiae | 7.07E-08 | 3.39E-08 | 1.21E-07 | 5 | 17 | 9 | 6594 | 1.337966344 | 0.000204539 | 0.000273666 |
| Human_herpesvirus_6B | 7.07E-08 | 3.39E-08 | 1.21E-07 | 5 | 17 | 9 | 10684 | 1.833333333 | 0.000851253 | 0.00156063 |
| Streptococcus_pyogenes | 6.37E-08 | 2.91E-08 | 1.11E-07 | 4 | 16 | 8 | 6576 | 1.46031746 | 0.000180373 | 0.000263402 |
| Yersinia_pestis | 6.37E-08 | 2.91E-08 | 1.11E-07 | 4 | 16 | 8 | 11928 | 44.17382379 | 6.11E-05 | 0.002700104 |
| Leptospira_noguchii | 6.37E-08 | 2.91E-08 | 1.11E-07 | 4 | 16 | 8 | 4560 | 13.18402181 | 2.14E-05 | 0.00028266 |
| Rickettsia_felis | 6.37E-08 | 2.91E-08 | 1.11E-07 | 4 | 16 | 8 | 5447 | 3.976993344 | 0.000161484 | 0.000642222 |
| Brucella_ovis | 5.66E-08 | 2.44E-08 | 1.02E-07 | 3 | 14 | 7 | 11511 | 1.464 | 3.82E-05 | 5.59E-05 |
| Rickettsia_prowazekii | 5.66E-08 | 2.44E-08 | 1.02E-07 | 3 | 14 | 7 | 5442 | 1.281481481 | 0.000121643 | 0.000155883 |
| Porphyromonas_gingivalis | 4.95E-08 | 1.99E-08 | 9.24E-08 | 3 | 13 | 6 | 3269 | 1.341666667 | 5.10E-05 | 6.84E-05 |
| Chlamydia_muridarum | 4.24E-08 | 1.56E-08 | 8.25E-08 | 2 | 12 | 5 | 2513 | 1.375722543 | 0.000158595 | 0.000218182 |
| Brucella_abortus | 4.24E-08 | 1.56E-08 | 8.25E-08 | 2 | 12 | 5 | 11496 | 1.545380579 | 2.78E-05 | 4.29E-05 |
| Borrelia_recurrentis | 4.24E-08 | 1.56E-08 | 8.25E-08 | 2 | 12 | 5 | 3042 | 1.674355099 | 8.21E-05 | 0.000137489 |
| Porphyromonas_gingivalis_W83 | 4.24E-08 | 1.56E-08 | 8.25E-08 | 2 | 12 | 5 | 3287 | 1 | 5.93E-05 | 5.93E-05 |
| Borreliella_afzelii | 4.24E-08 | 1.56E-08 | 8.25E-08 | 2 | 12 | 5 | 3370 | 1.375 | 7.84E-05 | 0.000107745 |
| Streptococcus_equi_zooepidemicus | 3.54E-08 | 1.15E-08 | 7.24E-08 | 2 | 10 | 4 | 6419 | 1 | 6.03E-05 | 6.03E-05 |
| Borreliella_garinii | 3.54E-08 | 1.15E-08 | 7.24E-08 | 2 | 10 | 4 | 3363 | 1.091422025 | 2.84E-05 | 3.10E-05 |
| Mycobacterium_tuberculosis | 3.54E-08 | 1.15E-08 | 7.24E-08 | 2 | 10 | 4 | 10716 | 1.769230769 | 8.84E-06 | 1.56E-05 |
| Equid_gammaherpesvirus_5 | 2.83E-08 | 7.71E-09 | 6.20E-08 | 1 | 9 | 3 | 2453 | 1 | 0.000515407 | 0.000515407 |
| Streptococcus_gordonii | 2.83E-08 | 7.71E-09 | 6.20E-08 | 1 | 9 | 3 | 6499 | 1.492063492 | 2.88E-05 | 4.30E-05 |
| Clostridium_sporogenes | 2.83E-08 | 7.71E-09 | 6.20E-08 | 1 | 9 | 3 | 4991 | 1.01081801 | 2.54E-05 | 2.57E-05 |
| Streptococcus_oralis | 2.83E-08 | 7.71E-09 | 6.20E-08 | 1 | 9 | 3 | 6632 | 1 | 0.000104579 | 0.000104579 |
| Chlamydia_caviae | 2.83E-08 | 7.71E-09 | 6.20E-08 | 1 | 9 | 3 | 2733 | 1.964466101 | 3.47E-05 | 6.82E-05 |
| Mycobacterium_africanum | 2.83E-08 | 7.71E-09 | 6.20E-08 | 1 | 9 | 3 | 10706 | 1.78125 | 7.12E-06 | 1.27E-05 |
| Clostridium_tetani | 2.12E-08 | 4.38E-09 | 5.11E-08 | 1 | 7 | 2 | 5299 | 99.6247902 | 1.68E-05 | 0.001678392 |
| Chlamydia_pneumoniae | 2.12E-08 | 4.38E-09 | 5.11E-08 | 1 | 7 | 2 | 2509 | 1.00606759 | 6.17E-05 | 6.20E-05 |
| Streptococcus_gordonii_str_Challis | 2.12E-08 | 4.38E-09 | 5.11E-08 | 1 | 7 | 2 | 6564 | 1 | 2.59E-05 | 2.59E-05 |
| Cowpox_virus | 2.12E-08 | 4.38E-09 | 5.11E-08 | 1 | 7 | 2 | 231 | 1 | 0.000409801 | 0.000409801 |
| Brucella_suis | 2.12E-08 | 4.38E-09 | 5.11E-08 | 1 | 7 | 2 | 11421 | 1 | 2.59E-05 | 2.59E-05 |

| | | | | | | | | | | |
|---|---|---|---|---|---|---|---|---|---|---|
| Clostridium_botulinum | 2.12E-08 | 4.38E-09 | 5.11E-08 | 1 | 7 | 2 | 4945 | 1.004204876 | 1.82E-05 | 1.83E-05 |
| Methanobrevibacter_oralis | 2.12E-08 | 4.38E-09 | 5.11E-08 | 1 | 7 | 2 | 149 | 353.0146341 | 3.28E-05 | 0.011562782 |
| Borreliella_burgdorferi | 2.12E-08 | 4.38E-09 | 5.11E-08 | 1 | 7 | 2 | 3341 | 2.804401528 | 4.62E-05 | 0.000129457 |
| Mycobacterium_bovis | 2.12E-08 | 4.38E-09 | 5.11E-08 | 1 | 7 | 2 | 10701 | 1 | 6.35E-06 | 6.35E-06 |
| Rickettsia_sibirica | 2.12E-08 | 4.38E-09 | 5.11E-08 | 1 | 7 | 2 | 5454 | 1 | 2.24E-05 | 2.24E-05 |
| Leptospira_kirschneri | 2.12E-08 | 4.38E-09 | 5.11E-08 | 1 | 7 | 2 | 4564 | 15.88523669 | 1.43E-05 | 0.000226962 |
| Chlamydia_felis | 1.41E-08 | 1.71E-09 | 3.94E-08 | 0 | 6 | 1 | 2736 | 1.006475517 | 2.47E-05 | 2.49E-05 |
| Chlamydia_psittaci | 1.41E-08 | 1.71E-09 | 3.94E-08 | 0 | 6 | 1 | 2778 | 1.006446371 | 2.29E-05 | 2.30E-05 |
| Rickettsia_akari | 1.41E-08 | 1.71E-09 | 3.94E-08 | 0 | 6 | 1 | 5379 | 1 | 3.01E-05 | 3.01E-05 |
| Rickettsia_conorii | 1.41E-08 | 1.71E-09 | 3.94E-08 | 0 | 6 | 1 | 5444 | 1 | 2.21E-05 | 2.21E-05 |
| Streptococcus_agalactiae | 1.41E-08 | 1.71E-09 | 3.94E-08 | 0 | 6 | 1 | 6580 | 1.002383594 | 2.26E-05 | 2.27E-05 |
| Rickettsia_japonica | 1.41E-08 | 1.71E-09 | 3.94E-08 | 0 | 6 | 1 | 5430 | 1 | 2.34E-05 | 2.34E-05 |
| Streptococcus_anginosus | 1.41E-08 | 1.71E-09 | 3.94E-08 | 0 | 6 | 1 | 7275 | 1 | 1.35E-05 | 1.35E-05 |
| Equid_gammaherpesvirus_2 | 1.41E-08 | 1.71E-09 | 3.94E-08 | 0 | 6 | 1 | 943 | 1 | 0.000168077 | 0.000168077 |
| Rickettsia_africae | 7.07E-09 | 1.79E-10 | 2.61E-08 | 0 | 4 | 0 | 5448 | 0 | 0 | 0 |
| Human_adenovirus_B1 | 7.07E-09 | 1.79E-10 | 2.61E-08 | 0 | 4 | 0 | 1 | 0 | 0 | 0 |
| Treponema_pallidum_subsp_pertenue | 7.07E-09 | 1.79E-10 | 2.61E-08 | 0 | 4 | 0 | 3618 | 0 | 0 | 0 |
| Marseilleviridae_tokyovirus | 7.07E-09 | 1.79E-10 | 2.61E-08 | 0 | 4 | 0 | 10 | 0 | 0 | 0 |
| Venezuelan_equine_encephalitis | 7.07E-09 | 1.79E-10 | 2.61E-08 | 0 | 4 | 0 | 23 | 0 | 0 | 0 |
| Yersinia_pestis_CO92 | 7.07E-09 | 1.79E-10 | 2.61E-08 | 0 | 4 | 0 | 11923 | 0 | 0 | 0 |
| Human_herpesvirus_3_strain_Dumas | 7.07E-09 | 1.79E-10 | 2.61E-08 | 0 | 4 | 0 | 1 | 0 | 0 | 0 |
| Human_herpesvirus_1 | 7.07E-09 | 1.79E-10 | 2.61E-08 | 0 | 4 | 0 | 15 | 0 | 0 | 0 |
| Human_alphaherpesvirus_1 | 7.07E-09 | 1.79E-10 | 2.61E-08 | 0 | 4 | 0 | 15 | 0 | 0 | 0 |
| Treponema_pallidum_subsp_pallidum | 7.07E-09 | 1.79E-10 | 2.61E-08 | 0 | 4 | 0 | 3617 | 0 | 0 | 0 |
| Chlamydia_pecorum | 7.07E-09 | 1.79E-10 | 2.61E-08 | 0 | 4 | 0 | 2561 | 0 | 0 | 0 |
| Human_adenovirus_66 | 7.07E-09 | 1.79E-10 | 2.61E-08 | 0 | 4 | 0 | 2 | 0 | 0 | 0 |
| Human_adenovirus_1 | 7.07E-09 | 1.79E-10 | 2.61E-08 | 0 | 4 | 0 | 4 | 0 | 0 | 0 |
| Equid_alphaherpesvirus_8 | 7.07E-09 | 1.79E-10 | 2.61E-08 | 0 | 4 | 0 | 201 | 0 | 0 | 0 |
| Equid_alphaherpesvirus_3 | 7.07E-09 | 1.79E-10 | 2.61E-08 | 0 | 4 | 0 | 10654 | 0 | 0 | 0 |
| Equid_alphaherpesvirus_1 | 7.07E-09 | 1.79E-10 | 2.61E-08 | 0 | 4 | 0 | 186 | 0 | 0 | 0 |
| Eastern_equine_encephalitis | 7.07E-09 | 1.79E-10 | 2.61E-08 | 0 | 4 | 0 | 6 | 0 | 0 | 0 |
| Rickettsia_rickettsii | 7.07E-09 | 1.79E-10 | 2.61E-08 | 0 | 4 | 0 | 5441 | 0 | 0 | 0 |
| Streptococcus_mutans | 7.07E-09 | 1.79E-10 | 2.61E-08 | 0 | 4 | 0 | 6243 | 0 | 0 | 0 |
| Clostridium_tetani_E88 | 7.07E-09 | 1.79E-10 | 2.61E-08 | 0 | 4 | 0 | 5308 | 0 | 0 | 0 |
| Human_adenovirus_50 | 7.07E-09 | 1.79E-10 | 2.61E-08 | 0 | 4 | 0 | 1 | 0 | 0 | 0 |
| Human_adenovirus_21 | 7.07E-09 | 1.79E-10 | 2.61E-08 | 0 | 4 | 0 | 2 | 0 | 0 | 0 |
| Human_adenovirus_40 | 7.07E-09 | 1.79E-10 | 2.61E-08 | 0 | 4 | 0 | 2 | 0 | 0 | 0 |
| Chlamydia_trachomatis | 7.07E-09 | 1.79E-10 | 2.61E-08 | 0 | 4 | 0 | 2497 | 0 | 0 | 0 |
| Human_adenovirus_41 | 7.07E-09 | 1.79E-10 | 2.61E-08 | 0 | 4 | 0 | 23 | 0 | 0 | 0 |
| Chlamydia_abortus | 7.07E-09 | 1.79E-10 | 2.61E-08 | 0 | 4 | 0 | 2770 | 0 | 0 | 0 |
| Human_adenovirus_4a | 7.07E-09 | 1.79E-10 | 2.61E-08 | 0 | 4 | 0 | 1 | 0 | 0 | 0 |
| Equid_alphaherpesvirus_9 | 7.07E-09 | 1.79E-10 | 2.61E-08 | 0 | 4 | 0 | 214 | 0 | 0 | 0 |

**Table S2e**: HAYSTAC results for library ERR6466112 based on a custom-built pathogen database (Table S1)

| Taxon | Mean_Posterior_Abundance | 95_CI_lower | 95_CI_upper | Minimum_Read_Num | Maximum_Read_Num | Dirichlet_Read_Num | Aligned_Read_Num | Evenness_of_Coverage_Ratio | Fraction_of_Genome_Covered | Coverage |
|---|---|---|---|---|---|---|---|---|---|---|
| Dark_Matter | 0.998647025 | 0.998641041 | 0.998652996 | 145087422 | 145089159 | 145088454 | 0 | | | |
| Grey_Matter | 0.000904016 | 0.000899136 | 0.000908909 | 130631 | 132051 | 131339 | 0 | | | |
| Burkholderia_cepacia | 8.54E-05 | 8.39E-05 | 8.69E-05 | 12196 | 12632 | 12412 | 49561 | 1.554817999 | 0.027440017 | 0.042664232 |
| Plasmodium_vivax | 5.53E-05 | 5.41E-05 | 5.65E-05 | 7856 | 8208 | 8030 | 25613 | 13.07168693 | 0.000529879 | 0.006926418 |
| Equid_alphaherpesvirus_4 | 3.60E-05 | 3.51E-05 | 3.70E-05 | 5093 | 5377 | 5233 | 5273 | 1.716726706 | 0.638715083 | 1.096499241 |
| Bordetella_pertussis | 3.57E-05 | 3.47E-05 | 3.66E-05 | 5040 | 5322 | 5179 | 28937 | 2.006166445 | 0.015111401 | 0.030315985 |
| Bordetella_petrii | 3.00E-05 | 2.91E-05 | 3.09E-05 | 4234 | 4493 | 4362 | 27135 | 1.79079572 | 0.011752191 | 0.021045774 |
| Taenia_solium | 2.60E-05 | 2.52E-05 | 2.69E-05 | 3662 | 3903 | 3781 | 31895 | 16.4142529 | 0.000146829 | 0.002410086 |
| Prescottella_equi | 1.89E-05 | 1.82E-05 | 1.96E-05 | 2649 | 2855 | 2750 | 15599 | 1.854182433 | 0.007773369 | 0.014413244 |
| Toxoplasma_gondii | 1.77E-05 | 1.70E-05 | 1.84E-05 | 2470 | 2668 | 2567 | 12837 | 2.500124665 | 0.000368139 | 0.000920394 |
| Mycobacterium_avium | 8.80E-06 | 8.32E-06 | 9.29E-06 | 1209 | 1349 | 1277 | 15191 | 1.399476813 | 0.004107853 | 0.005748846 |
| Plasmodium_falciparum | 8.55E-06 | 8.08E-06 | 9.03E-06 | 1174 | 1312 | 1241 | 13469 | 1.79953388 | 0.000802679 | 0.001444449 |
| Burkholderia_pseudomallei | 8.42E-06 | 7.95E-06 | 8.90E-06 | 1155 | 1292 | 1222 | 45695 | 1.584083862 | 0.002867729 | 0.004542724 |
| Klebsiella_oxytoca | 7.70E-06 | 7.25E-06 | 8.15E-06 | 1053 | 1184 | 1117 | 16186 | 1.196232123 | 0.005262392 | 0.006295042 |
| Schistosoma_mansoni | 7.25E-06 | 6.82E-06 | 7.69E-06 | 990 | 1118 | 1052 | 23892 | 1.723326122 | 6.50E-05 | 0.000111948 |
| Coxiella_burnetii | 6.99E-06 | 6.57E-06 | 7.43E-06 | 954 | 1079 | 1015 | 11354 | 6.719371118 | 0.00190082 | 0.012772314 |
| Mycobacterium_colombiense | 6.88E-06 | 6.46E-06 | 7.31E-06 | 938 | 1062 | 998 | 14564 | 1.452076667 | 0.002863667 | 0.004158264 |
| Mycobacterium_kansasii | 6.72E-06 | 6.30E-06 | 7.15E-06 | 916 | 1038 | 975 | 12596 | 1.546325228 | 0.002238329 | 0.003461185 |
| Trypanosoma_cruzi | 6.66E-06 | 6.24E-06 | 7.08E-06 | 907 | 1029 | 966 | 29786 | 3.09984046 | 0.000232161 | 0.000719663 |
| Mycobacterium_intracellulare | 6.60E-06 | 6.19E-06 | 7.03E-06 | 899 | 1021 | 958 | 14062 | 1.447054699 | 0.00281343 | 0.004071187 |
| Mycobacteroides_abscessus | 6.11E-06 | 5.71E-06 | 6.51E-06 | 830 | 946 | 886 | 11120 | 1.50242298 | 0.002802397 | 0.004210386 |
| Salmonella_enterica_enterica | 5.31E-06 | 4.95E-06 | 5.69E-06 | 718 | 827 | 771 | 15434 | 1.213115533 | 0.004506047 | 0.005466355 |
| Legionella_pneumophila | 3.87E-06 | 3.56E-06 | 4.19E-06 | 516 | 609 | 561 | 12764 | 3.215347465 | 0.001581017 | 0.005083518 |
| Bacillus_cereus | 3.81E-06 | 3.50E-06 | 4.14E-06 | 509 | 601 | 553 | 9187 | 1.047633434 | 0.00340245 | 0.00356452 |
| Blastomyces_dermatitidis | 3.21E-06 | 2.92E-06 | 3.51E-06 | 425 | 509 | 465 | 27741 | 1.71225098 | 0.000128106 | 0.00021935 |
| Burkholderia_mallei | 3.14E-06 | 2.86E-06 | 3.43E-06 | 415 | 499 | 455 | 44041 | 1.925473427 | 0.001170948 | 0.00225463 |
| Francisella_tularensis | 3.06E-06 | 2.78E-06 | 3.35E-06 | 404 | 486 | 443 | 9380 | 4.933896331 | 0.001836161 | 0.00905943 |
| Moraxella_catarrhalis | 2.93E-06 | 2.66E-06 | 3.22E-06 | 387 | 467 | 425 | 11539 | 3.241871136 | 0.002670997 | 0.00865903 |
| Trichinella_spiralis | 2.84E-06 | 2.57E-06 | 3.12E-06 | 373 | 453 | 411 | 13156 | 1.597039932 | 0.000146233 | 0.00023354 |
| Taylorella_equigenitalis | 1.95E-06 | 1.73E-06 | 2.19E-06 | 252 | 318 | 283 | 10830 | 2.543344387 | 0.002427757 | 0.006174621 |
| Histoplasma_capsulatum | 1.66E-06 | 1.46E-06 | 1.87E-06 | 212 | 272 | 240 | 27284 | 1.589999289 | 0.000134762 | 0.000214272 |
| Neisseria_gonorrhoeae | 1.60E-06 | 1.40E-06 | 1.81E-06 | 203 | 263 | 231 | 11554 | 1.860810588 | 0.001203174 | 0.00223888 |
| Corynebacterium_diphtheriae | 1.46E-06 | 1.27E-06 | 1.66E-06 | 184 | 241 | 211 | 7535 | 2.938882819 | 0.001079399 | 0.003172226 |
| Neisseria_meningitidis | 1.44E-06 | 1.25E-06 | 1.64E-06 | 182 | 238 | 208 | 11562 | 1.711851278 | 0.00179518 | 0.003073081 |
| Veillonella_parvula | 1.42E-06 | 1.24E-06 | 1.63E-06 | 180 | 236 | 206 | 5974 | 3.453310696 | 0.001104992 | 0.003815881 |
| Aspergillus_fumigatus | 1.33E-06 | 1.15E-06 | 1.52E-06 | 167 | 221 | 192 | 23750 | 1.802578269 | 9.24E-05 | 0.000166548 |
| Trypanosoma_brucei | 1.27E-06 | 1.09E-06 | 1.46E-06 | 158 | 212 | 183 | 20162 | 1.276033272 | 0.000170658 | 0.000217766 |
| Klebsiella_pneumoniae_rhinoscleroma | 1.24E-06 | 1.06E-06 | 1.43E-06 | 155 | 207 | 179 | 17269 | 1.176616642 | 0.000963212 | 0.001133331 |
| Coccidioides_immitis | 1.14E-06 | 9.75E-07 | 1.32E-06 | 142 | 192 | 165 | 22563 | 1.477707071 | 0.000114385 | 0.000169028 |
| Bacillus_anthracis | 1.07E-06 | 9.12E-07 | 1.25E-06 | 132 | 181 | 155 | 8944 | 1.395747972 | 0.00095659 | 0.001335158 |
| Staphylococcus_aureus | 9.43E-07 | 7.92E-07 | 1.11E-06 | 115 | 161 | 136 | 7110 | 1.567589143 | 0.001331981 | 0.002087999 |
| Coccidioides_posadasii | 9.36E-07 | 7.85E-07 | 1.10E-06 | 114 | 160 | 135 | 22597 | 1.444170129 | 0.00010554 | 0.000152418 |
| Yersinia_enterocolitica | 8.40E-07 | 6.97E-07 | 9.95E-07 | 101 | 145 | 121 | 1687 | 1.189751679 | 0.000596418 | 0.000709589 |
| Escherichia_coli | 8.33E-07 | 6.91E-07 | 9.88E-07 | 100 | 143 | 120 | 15102 | 1.464823355 | 0.000582716 | 0.000853576 |
| Helicobacter_pylori | 7.92E-07 | 6.54E-07 | 9.43E-07 | 95 | 137 | 114 | 4901 | 2.077996203 | 0.00111668 | 0.002320457 |
| Mycobacterium_leprae | 7.57E-07 | 6.22E-07 | 9.05E-07 | 90 | 131 | 109 | 8188 | 1.437141034 | 0.000491668 | 0.000706596 |
| Escherichia_coli_O26 | 6.88E-07 | 5.60E-07 | 8.30E-07 | 81 | 121 | 99 | 12246 | 185.7548023 | 6.27E-05 | 0.011637636 |
| Vibrio_parahaemolyticus | 6.81E-07 | 5.54E-07 | 8.22E-07 | 80 | 119 | 98 | 10921 | 1.656333511 | 0.000362192 | 0.000599911 |
| Fusobacterium_nucleatum | 6.75E-07 | 5.48E-07 | 8.14E-07 | 80 | 118 | 97 | 4917 | 2.26648161 | 0.000660979 | 0.001498096 |
| Tannerella_forsythia_92A2 | 5.92E-07 | 4.73E-07 | 7.23E-07 | 69 | 105 | 85 | 3455 | 1.356882739 | 0.000403169 | 0.000547053 |
| Corynebacterium_pseudotuberculosis | 5.71E-07 | 4.55E-07 | 7.01E-07 | 66 | 102 | 82 | 7461 | 1.39064143 | 0.000790878 | 0.001099828 |
| Vibrio_cholerae | 5.71E-07 | 4.55E-07 | 7.01E-07 | 66 | 102 | 82 | 11007 | 1.193784468 | 0.00042915 | 0.000512313 |
| Vibrio_vulnificus | 5.71E-07 | 4.55E-07 | 7.01E-07 | 66 | 102 | 82 | 11138 | 1.380200319 | 0.00025616 | 0.000353552 |
| Shigella_dysenteriae | 5.16E-07 | 4.06E-07 | 6.39E-07 | 59 | 93 | 74 | 15601 | 1.086573323 | 0.000546347 | 0.000593646 |
| Haemophilus_influenzae | 4.82E-07 | 3.76E-07 | 6.01E-07 | 55 | 87 | 69 | 9630 | 1.884724187 | 0.000765873 | 0.001443459 |
| Mycobacterium_canettii | 4.41E-07 | 3.39E-07 | 5.55E-07 | 49 | 81 | 63 | 11111 | 1.478079332 | 0.000213741 | 0.000315926 |
| Peptostreptococcus_anaerobius | 4.13E-07 | 3.15E-07 | 5.24E-07 | 46 | 76 | 59 | 5770 | 33.61526474 | 0.000351881 | 0.011828565 |
| Leptospira_kmetyi | 3.79E-07 | 2.85E-07 | 4.85E-07 | 41 | 70 | 54 | 5089 | 1.625544267 | 0.000155991 | 0.00025357 |
| Parvimonas_micra | 3.03E-07 | 2.20E-07 | 3.99E-07 | 32 | 58 | 43 | 5445 | 1.432806324 | 0.000608955 | 0.000872515 |
| Bartonella_henselae | 2.68E-07 | 1.91E-07 | 3.59E-07 | 28 | 52 | 38 | 8281 | 2.754464286 | 0.000235123 | 0.000647639 |

| | | | | | | | | | | |
|---|---|---|---|---|---|---|---|---|---|---|
| Leptospira_santarosai | 2.62E-07 | 1.85E-07 | 3.51E-07 | 27 | 51 | 37 | 5072 | 1.355679702 | 0.000134802 | 0.000182749 |
| Escherichia_coli_O157-H7 | 2.62E-07 | 1.85E-07 | 3.51E-07 | 27 | 51 | 37 | 15733 | 1.443592136 | 0.000151396 | 0.000218554 |
| Actinobacillus_equuli_equuli | 2.62E-07 | 1.85E-07 | 3.51E-07 | 27 | 51 | 37 | 9672 | 1.332867133 | 0.000294053 | 0.000391934 |
| Bartonella_bacilliformis | 2.48E-07 | 1.74E-07 | 3.35E-07 | 25 | 49 | 35 | 8272 | 1.198595787 | 0.000689955 | 0.000826978 |
| Shigella_flexneri | 2.41E-07 | 1.68E-07 | 3.27E-07 | 24 | 48 | 34 | 15252 | 1.023335621 | 0.00030173 | 0.000308771 |
| Cryptococcus_neoformans | 2.41E-07 | 1.68E-07 | 3.27E-07 | 24 | 48 | 34 | 6106 | 1.78345776 | 4.28E-05 | 7.64E-05 |
| Cryptococcus_gattii | 2.34E-07 | 1.62E-07 | 3.19E-07 | 24 | 46 | 33 | 8123 | 1.885287206 | 3.60E-05 | 6.79E-05 |
| Leptospira_licerasiae | 2.34E-07 | 1.62E-07 | 3.19E-07 | 24 | 46 | 33 | 4971 | 2.934343213 | 7.79E-05 | 0.000228614 |
| Treponema_denticola_ATCC_35405 | 2.27E-07 | 1.56E-07 | 3.11E-07 | 23 | 45 | 32 | 3586 | 1.668161435 | 0.000235298 | 0.000392515 |
| Clostridium_perfringens | 2.27E-07 | 1.56E-07 | 3.11E-07 | 23 | 45 | 32 | 5529 | 1.312113652 | 0.000286375 | 0.000375757 |
| Leptospira_borgpetersenii | 2.20E-07 | 1.51E-07 | 3.03E-07 | 22 | 44 | 31 | 5033 | 2.875 | 6.01E-05 | 0.000172812 |
| Bartonella_quintana | 2.06E-07 | 1.39E-07 | 2.87E-07 | 20 | 42 | 29 | 8296 | 1.512733447 | 0.000356943 | 0.000539959 |
| Leptospira_weilii | 1.93E-07 | 1.28E-07 | 2.70E-07 | 19 | 39 | 27 | 5278 | 1.392710372 | 9.91E-05 | 0.000137955 |
| Shigella_boydii | 1.79E-07 | 1.17E-07 | 2.54E-07 | 17 | 37 | 25 | 15288 | 10.94307809 | 0.000162473 | 0.001777959 |
| Leptospira_inadai | 1.72E-07 | 1.11E-07 | 2.46E-07 | 16 | 36 | 24 | 4913 | 1.377265487 | 0.000130331 | 0.000179501 |
| Escherichia_coli_O111 | 1.72E-07 | 1.11E-07 | 2.46E-07 | 16 | 36 | 24 | 16122 | 99.58216899 | 9.84E-05 | 0.009794507 |
| Yersinia_pseudotuberculosis | 1.72E-07 | 1.11E-07 | 2.46E-07 | 16 | 36 | 24 | 12412 | 3.924635862 | 2.98E-05 | 0.00011678 |
| Shigella_sonnei | 1.65E-07 | 1.06E-07 | 2.38E-07 | 15 | 35 | 23 | 15565 | 1 | 0.000161041 | 0.000161041 |
| Leptospira_alstonii | 1.38E-07 | 8.41E-08 | 2.04E-07 | 12 | 30 | 19 | 4984 | 6.301376212 | 8.26E-05 | 0.000520686 |
| Borrelia_recurrentis | 1.38E-07 | 8.41E-08 | 2.04E-07 | 12 | 30 | 19 | 3283 | 1.593069779 | 0.000240709 | 0.000383466 |
| Leptospira_wolffii | 1.38E-07 | 8.41E-08 | 2.04E-07 | 12 | 30 | 19 | 5050 | 1.340803844 | 6.73E-05 | 9.03E-05 |
| Mycoplasma_pneumoniae | 1.38E-07 | 8.41E-08 | 2.04E-07 | 12 | 30 | 19 | 1808 | 1.517110266 | 0.000306599 | 0.000465144 |
| Streptococcus_salivarius | 1.10E-07 | 6.29E-08 | 1.70E-07 | 9 | 25 | 15 | 6962 | 1.598290598 | 0.00021418 | 0.000342322 |
| Leptospira_broomii | 1.03E-07 | 5.78E-08 | 1.62E-07 | 8 | 23 | 14 | 4915 | 1.378300386 | 5.76E-05 | 7.93E-05 |
| Pneumocystis_carinii | 1.03E-07 | 5.78E-08 | 1.62E-07 | 8 | 23 | 14 | 14500 | 2.952376732 | 4.23E-05 | 0.000124855 |
| Brucella_microti | 1.03E-07 | 5.78E-08 | 1.62E-07 | 8 | 23 | 14 | 11992 | 1.132034632 | 0.000138432 | 0.00015671 |
| Leptospira_noguchii | 9.64E-08 | 5.27E-08 | 1.53E-07 | 8 | 22 | 13 | 4961 | 3.623826909 | 4.25E-05 | 0.000153848 |
| Clostridium_botulinum_E3 | 9.64E-08 | 5.27E-08 | 1.53E-07 | 8 | 22 | 13 | 5598 | 1 | 9.37E-05 | 9.37E-05 |
| Human_herpesvirus_6B | 9.64E-08 | 5.27E-08 | 1.53E-07 | 8 | 22 | 13 | 11119 | 2.262711864 | 0.000727883 | 0.001646989 |
| Leptospira_kirschneri | 8.95E-08 | 4.76E-08 | 1.44E-07 | 7 | 21 | 12 | 4983 | 6.02910703 | 3.36E-05 | 0.000202364 |
| Streptococcus_pyogenes | 8.95E-08 | 4.76E-08 | 1.44E-07 | 7 | 21 | 12 | 7216 | 1.245231608 | 0.000210149 | 0.000261684 |
| Klebsiella_pneumoniae_pneumoniae | 8.26E-08 | 4.27E-08 | 1.35E-07 | 6 | 20 | 11 | 16727 | 1.159734757 | 5.39E-05 | 6.25E-05 |
| Rickettsia_prowazekii | 8.26E-08 | 4.27E-08 | 1.35E-07 | 6 | 20 | 11 | 5693 | 2.149253731 | 0.000120742 | 0.000259505 |
| Leptospira_alexanderi | 8.26E-08 | 4.27E-08 | 1.35E-07 | 6 | 20 | 11 | 5138 | 213.4181705 | 2.88E-05 | 0.006148774 |
| Leptospira_fainei | 7.57E-08 | 3.78E-08 | 1.27E-07 | 5 | 18 | 10 | 4892 | 1.538832392 | 5.90E-05 | 9.08E-05 |
| Streptococcus_gordonii | 7.57E-08 | 3.78E-08 | 1.27E-07 | 5 | 18 | 10 | 7028 | 1 | 9.20E-05 | 9.20E-05 |
| Streptococcus_dysgalactiae | 7.57E-08 | 3.78E-08 | 1.27E-07 | 5 | 18 | 10 | 7199 | 1.115172236 | 9.72E-05 | 0.000108346 |
| Streptococcus_oralis | 6.88E-08 | 3.30E-08 | 1.18E-07 | 5 | 17 | 9 | 7191 | 1 | 0.000251094 | 0.000251094 |
| Brucella_suis | 6.88E-08 | 3.30E-08 | 1.18E-07 | 5 | 17 | 9 | 11899 | 1 | 7.27E-05 | 7.27E-05 |
| Pneumocystis_jirovecii | 6.88E-08 | 3.30E-08 | 1.18E-07 | 5 | 17 | 9 | 14025 | 4.685822703 | 2.86E-05 | 0.000133941 |
| Streptococcus_agalactiae | 6.19E-08 | 2.83E-08 | 1.08E-07 | 4 | 16 | 8 | 7090 | 1.002383594 | 0.000155835 | 0.000156206 |
| Streptococcus_sanguinis | 6.19E-08 | 2.83E-08 | 1.08E-07 | 4 | 16 | 8 | 7265 | 1.074170614 | 0.000121395 | 0.000130399 |
| Yersinia_pestis | 5.51E-08 | 2.38E-08 | 9.93E-08 | 3 | 14 | 7 | 12296 | 44.17382379 | 7.72E-05 | 0.003412117 |
| Streptococcus_pneumoniae | 5.51E-08 | 2.38E-08 | 9.93E-08 | 3 | 14 | 7 | 6943 | 1.867957854 | 7.10E-05 | 0.00013266 |
| Tateropox | 5.51E-08 | 2.38E-08 | 9.93E-08 | 3 | 14 | 7 | 44 | 1.480916031 | 0.001322898 | 0.001959101 |
| Porphyromonas_gingivalis_W83 | 5.51E-08 | 2.38E-08 | 9.93E-08 | 3 | 14 | 7 | 3429 | 1.18079096 | 7.55E-05 | 8.92E-05 |
| Rickettsia_felis | 4.82E-08 | 1.94E-08 | 8.99E-08 | 3 | 13 | 6 | 5702 | 3.410091885 | 9.12E-05 | 0.00031097 |
| Methanobrevibacter_oralis | 4.13E-08 | 1.52E-08 | 8.03E-08 | 2 | 12 | 5 | 160 | 684.634442 | 1.69E-05 | 0.011562782 |
| Streptococcus_gordonii_str_Challis | 4.13E-08 | 1.52E-08 | 8.03E-08 | 2 | 12 | 5 | 7092 | 1 | 6.33E-05 | 6.33E-05 |
| Leptospira_interrogans | 3.44E-08 | 1.12E-08 | 7.05E-08 | 2 | 10 | 4 | 4959 | 1.081806866 | 1.40E-05 | 1.52E-05 |
| Clostridium_botulinum | 3.44E-08 | 1.12E-08 | 7.05E-08 | 2 | 10 | 4 | 5350 | 1.004204876 | 4.92E-05 | 4.94E-05 |
| Streptococcus_equi_zooepidemicus | 2.75E-08 | 7.50E-09 | 6.03E-08 | 1 | 9 | 3 | 6967 | 1 | 4.46E-05 | 4.46E-05 |
| Porphyromonas_gingivalis | 2.75E-08 | 7.50E-09 | 6.03E-08 | 1 | 9 | 3 | 3444 | 1 | 2.80E-05 | 2.80E-05 |
| Borreliella_burgdorferi | 2.75E-08 | 7.50E-09 | 6.03E-08 | 1 | 9 | 3 | 3499 | 2.030607384 | 4.16E-05 | 8.45E-05 |
| Chlamydia_pneumoniae | 2.75E-08 | 7.50E-09 | 6.03E-08 | 1 | 9 | 3 | 2741 | 1.00606759 | 5.37E-05 | 5.40E-05 |
| Streptococcus_mutans | 2.75E-08 | 7.50E-09 | 6.03E-08 | 1 | 9 | 3 | 6826 | 1.006271233 | 5.08E-05 | 5.11E-05 |
| Brucella_ovis | 2.75E-08 | 7.50E-09 | 6.03E-08 | 1 | 9 | 3 | 11991 | 1.551405012 | 3.18E-05 | 4.93E-05 |
| Clostridium_tetani | 2.06E-08 | 4.26E-09 | 4.97E-08 | 1 | 7 | 2 | 5857 | 99.6247902 | 1.68E-05 | 0.001678392 |
| Chlamydia_trachomatis | 2.06E-08 | 4.26E-09 | 4.97E-08 | 1 | 7 | 2 | 2734 | 1 | 5.56E-05 | 5.56E-05 |
| Brucella_melitensis | 2.06E-08 | 4.26E-09 | 4.97E-08 | 1 | 7 | 2 | 11880 | 1.556309349 | 1.94E-05 | 3.02E-05 |
| Mycobacterium_tuberculosis | 2.06E-08 | 4.26E-09 | 4.97E-08 | 1 | 7 | 2 | 11135 | 1.666666667 | 9.52E-06 | 1.59E-05 |
| Mycobacterium_africanum | 2.06E-08 | 4.26E-09 | 4.97E-08 | 1 | 7 | 2 | 11125 | 1 | 8.68E-06 | 8.68E-06 |
| Streptococcus_anginosus | 2.06E-08 | 4.26E-09 | 4.97E-08 | 1 | 7 | 2 | 7892 | 1 | 1.40E-05 | 1.40E-05 |
| Rickettsia_conorii | 2.06E-08 | 4.26E-09 | 4.97E-08 | 1 | 7 | 2 | 5703 | 1 | 4.89E-05 | 4.89E-05 |

| | | | | | | | | | | |
|---|---|---|---|---|---|---|---|---|---|---|
| Borreliella_afzelii | 1.38E-08 | 1.67E-09 | 3.83E-08 | 0 | 6 | 1 | 3536 | 1 | 4.68E-05 | 4.68E-05 |
| Cowpox_virus | 1.38E-08 | 1.67E-09 | 3.83E-08 | 0 | 6 | 1 | 254 | 1 | 0.000253899 | 0.000253899 |
| Rickettsia_rickettsii | 1.38E-08 | 1.67E-09 | 3.83E-08 | 0 | 6 | 1 | 5708 | 1 | 2.70E-05 | 2.70E-05 |
| Yersinia_pestis_CO92 | 1.38E-08 | 1.67E-09 | 3.83E-08 | 0 | 6 | 1 | 12293 | 1 | 8.81E-06 | 8.81E-06 |
| Brucella_abortus | 1.38E-08 | 1.67E-09 | 3.83E-08 | 0 | 6 | 1 | 11940 | 1.545380579 | 8.24E-06 | 1.27E-05 |
| Clostridium_sporogenes | 1.38E-08 | 1.67E-09 | 3.83E-08 | 0 | 6 | 1 | 5410 | 1.01081801 | 8.71E-06 | 8.81E-06 |
| Rickettsia_typhi | 1.38E-08 | 1.67E-09 | 3.83E-08 | 0 | 6 | 1 | 5681 | 1 | 3.96E-05 | 3.96E-05 |
| Rickettsia_akari | 1.38E-08 | 1.67E-09 | 3.83E-08 | 0 | 6 | 1 | 5669 | 1 | 3.98E-05 | 3.98E-05 |
| Chlamydia_caviae | 1.38E-08 | 1.67E-09 | 3.83E-08 | 0 | 6 | 1 | 2946 | 1.006788877 | 4.82E-05 | 4.86E-05 |
| Chlamydia_muridarum | 1.38E-08 | 1.67E-09 | 3.83E-08 | 0 | 6 | 1 | 2747 | 1 | 4.77E-05 | 4.77E-05 |
| Eastern_equine_encephalitis | 6.88E-09 | 1.74E-10 | 2.54E-08 | 0 | 4 | 0 | 4 | 0 | 0 | 0 |
| Borreliella_garinii | 6.88E-09 | 1.74E-10 | 2.54E-08 | 0 | 4 | 0 | 3523 | 0 | 0 | 0 |
| Rickettsia_japonica | 6.88E-09 | 1.74E-10 | 2.54E-08 | 0 | 4 | 0 | 5693 | 0 | 0 | 0 |
| Treponema_pallidum_subsp_pertenue | 6.88E-09 | 1.74E-10 | 2.54E-08 | 0 | 4 | 0 | 3874 | 0 | 0 | 0 |
| Marseilleviridae_tokyovirus | 6.88E-09 | 1.74E-10 | 2.54E-08 | 0 | 4 | 0 | 11 | 0 | 0 | 0 |
| Treponema_pallidum_subsp_pallidum | 6.88E-09 | 1.74E-10 | 2.54E-08 | 0 | 4 | 0 | 3871 | 0 | 0 | 0 |
| Human_herpesvirus_1 | 6.88E-09 | 1.74E-10 | 2.54E-08 | 0 | 4 | 0 | 13 | 0 | 0 | 0 |
| Venezuelan_equine_encephalitis | 6.88E-09 | 1.74E-10 | 2.54E-08 | 0 | 4 | 0 | 28 | 0 | 0 | 0 |
| Rickettsia_africae | 6.88E-09 | 1.74E-10 | 2.54E-08 | 0 | 4 | 0 | 5709 | 0 | 0 | 0 |
| Mycobacterium_bovis | 6.88E-09 | 1.74E-10 | 2.54E-08 | 0 | 4 | 0 | 11110 | 0 | 0 | 0 |
| Rickettsia_sibirica | 6.88E-09 | 1.74E-10 | 2.54E-08 | 0 | 4 | 0 | 5693 | 0 | 0 | 0 |
| Human_adenovirus_4a | 6.88E-09 | 1.74E-10 | 2.54E-08 | 0 | 4 | 0 | 2 | 0 | 0 | 0 |
| Human_hepatitis_A_virus | 6.88E-09 | 1.74E-10 | 2.54E-08 | 0 | 4 | 0 | 1 | 0 | 0 | 0 |
| Human_alphaherpesvirus_1 | 6.88E-09 | 1.74E-10 | 2.54E-08 | 0 | 4 | 0 | 13 | 0 | 0 | 0 |
| Equid_alphaherpesvirus_3 | 6.88E-09 | 1.74E-10 | 2.54E-08 | 0 | 4 | 0 | 11080 | 0 | 0 | 0 |
| Equid_alphaherpesvirus_8 | 6.88E-09 | 1.74E-10 | 2.54E-08 | 0 | 4 | 0 | 176 | 0 | 0 | 0 |
| Equid_alphaherpesvirus_9 | 6.88E-09 | 1.74E-10 | 2.54E-08 | 0 | 4 | 0 | 184 | 0 | 0 | 0 |
| Equid_gammaherpesvirus_5 | 6.88E-09 | 1.74E-10 | 2.54E-08 | 0 | 4 | 0 | 2513 | 0 | 0 | 0 |
| Clostridium_tetani_E88 | 6.88E-09 | 1.74E-10 | 2.54E-08 | 0 | 4 | 0 | 5857 | 0 | 0 | 0 |
| Human_adenovirus_1 | 6.88E-09 | 1.74E-10 | 2.54E-08 | 0 | 4 | 0 | 7 | 0 | 0 | 0 |
| Human_adenovirus_31 | 6.88E-09 | 1.74E-10 | 2.54E-08 | 0 | 4 | 0 | 2 | 0 | 0 | 0 |
| Human_adenovirus_41 | 6.88E-09 | 1.74E-10 | 2.54E-08 | 0 | 4 | 0 | 23 | 0 | 0 | 0 |
| Chlamydia_psittaci | 6.88E-09 | 1.74E-10 | 2.54E-08 | 0 | 4 | 0 | 3013 | 0 | 0 | 0 |
| Human_adenovirus_49 | 6.88E-09 | 1.74E-10 | 2.54E-08 | 0 | 4 | 0 | 2 | 0 | 0 | 0 |
| Chlamydia_pecorum | 6.88E-09 | 1.74E-10 | 2.54E-08 | 0 | 4 | 0 | 2756 | 0 | 0 | 0 |
| Chlamydia_felis | 6.88E-09 | 1.74E-10 | 2.54E-08 | 0 | 4 | 0 | 2954 | 0 | 0 | 0 |
| Chlamydia_abortus | 6.88E-09 | 1.74E-10 | 2.54E-08 | 0 | 4 | 0 | 3002 | 0 | 0 | 0 |
| Equid_alphaherpesvirus_1 | 6.88E-09 | 1.74E-10 | 2.54E-08 | 0 | 4 | 0 | 186 | 0 | 0 | 0 |
| Human_adenovirus_61 | 6.88E-09 | 1.74E-10 | 2.54E-08 | 0 | 4 | 0 | 2 | 0 | 0 | 0 |
| Equid_gammaherpesvirus_2 | 6.88E-09 | 1.74E-10 | 2.54E-08 | 0 | 4 | 0 | 975 | 0 | 0 | 0 |

**Table S3a:** Results of the Kraken2 analysis for library ERR6466108 based on a custom-built pathogen list (Table S1)

| Sr | Percentage of fragments covered | Number of fragments covered | Number of fragments assigned | rank code | NCBI taxonomic ID | Name |
|---|---|---|---|---|---|---|
| 1 | 99.73 | 9388640 | 9388640 | U | 0 | unclassified |
| 2 | 0.27 | 24988 | 0 | R | 1 | root |
| 3 | 0.26 | 24737 | 150 | R1 | 131567 | cellular organisms |
| 4 | 0.24 | 22611 | 671 | D | 2 | Bacteria |
| 5 | 0.18 | 17089 | 495 | P | 1224 | Pseudomonadota |
| 6 | 0.13 | 12285 | 118 | C | 28216 | Betaproteobacteria |
| 7 | 0.13 | 11859 | 433 | O | 80840 | Burkholderiales |
| 8 | 0.08 | 7379 | 0 | F | 119060 | Burkholderiaceae |
| 9 | 0.08 | 7379 | 1112 | G | 32008 | Burkholderia |
| 10 | 0.04 | 3514 | 0 | G1 | 87882 | Burkholderia cepacia complex |
| 11 | 0.04 | 3514 | 3514 | S | 292 | Burkholderia cepacia |
| 12 | 0.03 | 2753 | 2360 | G1 | 111527 | pseudomallei group |
| 13 | 0.00 | 300 | 300 | S | 28450 | Burkholderia pseudomallei |
| 14 | 0.00 | 93 | 93 | S | 13373 | Burkholderia mallei |
| 15 | 0.04 | 4047 | 33 | F | 506 | Alcaligenaceae |
| 16 | 0.04 | 3950 | 378 | G | 517 | Bordetella |
| 17 | 0.02 | 1846 | 1846 | S | 520 | Bordetella pertussis |
| 18 | 0.02 | 1726 | 1726 | S | 94624 | Bordetella petrii |
| 19 | 0.00 | 64 | 0 | G | 29574 | Taylorella |
| 20 | 0.00 | 64 | 64 | S | 29575 | Taylorella equigenitalis |
| 21 | 0.00 | 308 | 0 | O | 206351 | Neisseriales |
| 22 | 0.00 | 308 | 0 | F | 481 | Neisseriaceae |
| 23 | 0.00 | 308 | 142 | G | 482 | Neisseria |
| 24 | 0.00 | 69 | 69 | S | 485 | Neisseria gonorrhoeae |
| 25 | 0.00 | 57 | 57 | S | 487 | Neisseria meningitidis |
| 26 | 0.00 | 40 | 40 | S | 483 | Neisseria cinerea |
| 27 | 0.04 | 3427 | 268 | C | 1236 | Gammaproteobacteria |
| 28 | 0.02 | 2223 | 77 | O | 91347 | Enterobacterales |
| 29 | 0.02 | 2022 | 443 | F | 543 | Enterobacteriaceae |
| 30 | 0.01 | 1188 | 0 | F1 | 2890311 | Klebsiella/Raoultella group |
| 31 | 0.01 | 1188 | 95 | G | 570 | Klebsiella |
| 32 | 0.01 | 634 | 603 | S | 573 | Klebsiella pneumoniae |
| 33 | 0.00 | 31 | 31 | S1 | 39831 | Klebsiella pneumoniae subsp. rhinoscleromatis |
| 34 | 0.00 | 459 | 459 | S | 571 | Klebsiella oxytoca |
| 35 | 0.00 | 272 | 0 | G | 590 | Salmonella |
| 36 | 0.00 | 272 | 0 | S | 28901 | Salmonella enterica |
| 37 | 0.00 | 272 | 0 | S1 | 59201 | Salmonella enterica subsp. enterica |
| 38 | 0.00 | 272 | 272 | S2 | 594 | Salmonella enterica subsp. enterica serovar Gallinarum |
| 39 | 0.00 | 99 | 0 | G | 561 | Escherichia |
| 40 | 0.00 | 99 | 68 | S | 562 | Escherichia coli |
| 41 | 0.00 | 23 | 23 | S1 | 83334 | Escherichia coli O157:H7 |
| 42 | 0.00 | 4 | 4 | S1 | 168927 | Escherichia coli O111:H- |
| 43 | 0.00 | 4 | 4 | S1 | 244319 | Escherichia coli O26:H11 |
| 44 | 0.00 | 20 | 11 | G | 620 | Shigella |
| 45 | 0.00 | 9 | 9 | S | 623 | Shigella flexneri |
| 46 | 0.00 | 124 | 0 | F | 1903411 | Yersiniaceae |
| 47 | 0.00 | 124 | 43 | G | 629 | Yersinia |
| 48 | 0.00 | 50 | 34 | G1 | 1649845 | Yersinia pseudotuberculosis complex |
| 49 | 0.00 | 16 | 16 | S | 632 | Yersinia pestis |
| 50 | 0.00 | 31 | 31 | S | 630 | Yersinia enterocolitica |
| 51 | 0.00 | 466 | 36 | O | 118969 | Legionellales |
| 52 | 0.00 | 251 | 0 | F | 118968 | Coxiellaceae |
| 53 | 0.00 | 251 | 0 | G | 776 | Coxiella |
| 54 | 0.00 | 251 | 251 | S | 777 | Coxiella burnetii |
| 55 | 0.00 | 179 | 0 | F | 444 | Legionellaceae |
| 56 | 0.00 | 179 | 0 | G | 445 | Legionella |
| 57 | 0.00 | 179 | 179 | S | 446 | Legionella pneumophila |
| 58 | 0.00 | 166 | 0 | O | 2887326 | Moraxellales |
| 59 | 0.00 | 166 | 0 | F | 468 | Moraxellaceae |
| 60 | 0.00 | 166 | 0 | G | 475 | Moraxella |
| 61 | 0.00 | 166 | 166 | S | 480 | Moraxella catarrhalis |
| 62 | 0.00 | 133 | 0 | O | 135623 | Vibrionales |
| 63 | 0.00 | 133 | 0 | F | 641 | Vibrionaceae |
| 64 | 0.00 | 133 | 39 | G | 662 | Vibrio |
| 65 | 0.00 | 36 | 0 | G1 | 717610 | Vibrio harveyi group |
| 66 | 0.00 | 36 | 36 | S | 670 | Vibrio parahaemolyticus |
| 67 | 0.00 | 32 | 32 | S | 666 | Vibrio cholerae |
| 68 | 0.00 | 26 | 26 | S | 672 | Vibrio vulnificus |
| 69 | 0.00 | 112 | 0 | O | 72273 | Thiotrichales |
| 70 | 0.00 | 112 | 0 | F | 34064 | Francisellaceae |
| 71 | 0.00 | 112 | 0 | G | 262 | Francisella |
| 72 | 0.00 | 112 | 112 | S | 263 | Francisella tularensis |
| 73 | 0.00 | 59 | 0 | O | 135625 | Pasteurellales |
| 74 | 0.00 | 59 | 13 | F | 712 | Pasteurellaceae |
| 75 | 0.00 | 26 | 0 | G | 713 | Actinobacillus |
| 76 | 0.00 | 26 | 0 | S | 718 | Actinobacillus equuli |
| 77 | 0.00 | 26 | 26 | S1 | 202947 | Actinobacillus equuli subsp. equuli |
| 78 | 0.00 | 20 | 0 | G | 724 | Haemophilus |
| 79 | 0.00 | 20 | 0 | S | 727 | Haemophilus influenzae |
| 80 | 0.00 | 20 | 20 | S1 | 725 | Haemophilus influenzae biotype aegyptius |
| 81 | 0.01 | 882 | 8 | C | 28211 | Alphaproteobacteria |
| 82 | 0.01 | 811 | 127 | O | 356 | Hyphomicrobiales |
| 83 | 0.01 | 580 | 0 | F | 118882 | Brucellaceae |
| 84 | 0.01 | 580 | 0 | F1 | 2826938 | Brucella/Ochrobactrum group |
| 85 | 0.01 | 580 | 562 | G | 234 | Brucella |
| 86 | 0.00 | 8 | 0 | S | 236 | Brucella ovis |
| 87 | 0.00 | 8 | 8 | S1 | 444178 | Brucella ovis ATCC 25840 |
| 88 | 0.00 | 5 | 0 | S | 444163 | Brucella microti |
| 89 | 0.00 | 5 | 5 | S1 | 568815 | Brucella microti CCM 4915 |
| 90 | 0.00 | 4 | 0 | S | 29459 | Brucella melitensis |
| 91 | 0.00 | 4 | 0 | S1 | 644337 | Brucella melitensis bv. 1 |
| 92 | 0.00 | 4 | 4 | S2 | 224914 | Brucella melitensis bv. 1 str. 16M |
| 93 | 0.00 | 1 | 0 | S | 235 | Brucella abortus |
| 94 | 0.00 | 1 | 1 | S1 | 359391 | Brucella abortus 2308 |
| 95 | 0.00 | 104 | 0 | F | 772 | Bartonellaceae |
| 96 | 0.00 | 104 | 69 | G | 773 | Bartonella |
| 97 | 0.00 | 16 | 16 | S | 38323 | Bartonella henselae |
| 98 | 0.00 | 14 | 0 | S | 774 | Bartonella bacilliformis |
| 99 | 0.00 | 14 | 14 | S1 | 360095 | Bartonella bacilliformis KC583 |
| 100 | 0.00 | 5 | 5 | S | 803 | Bartonella quintana |
| 101 | 0.00 | 63 | 0 | O | 766 | Rickettsiales |
| 102 | 0.00 | 63 | 0 | F | 775 | Rickettsiaceae |
| 103 | 0.00 | 63 | 0 | F1 | 33988 | Rickettsieae |
| 104 | 0.00 | 63 | 42 | G | 780 | Rickettsia |
| 105 | 0.00 | 19 | 11 | G1 | 114277 | spotted fever group |
| 106 | 0.00 | 3 | 0 | S | 786 | Rickettsia akari |
| 107 | 0.00 | 3 | 3 | S1 | 293614 | Rickettsia akari str. Hartford |
| 108 | 0.00 | 2 | 2 | S | 35790 | Rickettsia japonica |
| 109 | 0.00 | 2 | 2 | S | 42862 | Rickettsia felis |
| 110 | 0.00 | 1 | 0 | G2 | 266068 | Rickettsia sibirica subgroup |

| | | | | | | |
|---|---|---|---|---|---|---|
| 111 | 0.00 | 1 | 0 | S | 35793 | Rickettsia sibirica |
| 112 | 0.00 | 1 | 1 | S1 | 272951 | Rickettsia sibirica 246 |
| 113 | 0.00 | 2 | 0 | G1 | 114292 | typhus group |
| 114 | 0.00 | 2 | 2 | S | 785 | Rickettsia typhi |
| 115 | 0.05 | 4388 | 30 | D1 | 1783272 | Terrabacteria group |
| 116 | 0.04 | 3694 | 0 | P | 201174 | Actinomycetota |
| 117 | 0.04 | 3694 | 0 | C | 1760 | Actinomycetes |
| 118 | 0.04 | 3694 | 301 | O | 85007 | Mycobacteriales |
| 119 | 0.03 | 2408 | 100 | F | 1762 | Mycobacteriaceae |
| 120 | 0.02 | 2056 | 217 | G | 1763 | Mycobacterium |
| 121 | 0.01 | 1239 | 246 | G1 | 120793 | Mycobacterium avium complex (MAC) |
| 122 | 0.00 | 419 | 419 | S | 1764 | Mycobacterium avium |
| 123 | 0.00 | 304 | 304 | S | 339268 | Mycobacterium colombiense |
| 124 | 0.00 | 270 | 270 | S | 1767 | Mycobacterium intracellulare |
| 125 | 0.00 | 338 | 338 | S | 1768 | Mycobacterium kansasii |
| 126 | 0.00 | 210 | 181 | G1 | 77643 | Mycobacterium tuberculosis complex |
| 127 | 0.00 | 15 | 15 | S | 1773 | Mycobacterium tuberculosis |
| 128 | 0.00 | 14 | 14 | S | 78331 | Mycobacterium canettii |
| 129 | 0.00 | 52 | 52 | S | 1769 | Mycobacterium leprae |
| 130 | 0.00 | 252 | 0 | G | 670516 | Mycobacteroides |
| 131 | 0.00 | 252 | 0 | S | 36809 | Mycobacteroides abscessus |
| 132 | 0.00 | 252 | 252 | S1 | 319705 | Mycobacteroides abscessus subsp. bolletii |
| 133 | 0.01 | 776 | 0 | F | 85025 | Nocardiaceae |
| 134 | 0.01 | 776 | 0 | G | 2979332 | Prescottella |
| 135 | 0.01 | 776 | 776 | S | 43767 | Prescottella equi |
| 136 | 0.00 | 209 | 0 | F | 1653 | Corynebacteriaceae |
| 137 | 0.00 | 209 | 32 | G | 1716 | Corynebacterium |
| 138 | 0.00 | 145 | 145 | S | 1717 | Corynebacterium diphtheriae |
| 139 | 0.00 | 32 | 0 | S | 1719 | Corynebacterium pseudotuberculosis |
| 140 | 0.00 | 32 | 32 | S1 | 1087451 | Corynebacterium pseudotuberculosis 31 |
| 141 | 0.01 | 656 | 60 | P | 1239 | Bacillota |
| 142 | 0.00 | 356 | 23 | C | 91061 | Bacilli |
| 143 | 0.00 | 271 | 28 | O | 1385 | Bacillales |
| 144 | 0.00 | 183 | 0 | F | 186817 | Bacillaceae |
| 145 | 0.00 | 183 | 0 | G | 1386 | Bacillus |
| 146 | 0.00 | 183 | 110 | G1 | 86661 | Bacillus cereus group |
| 147 | 0.00 | 49 | 49 | S | 1396 | Bacillus cereus |
| 148 | 0.00 | 24 | 0 | S | 1392 | Bacillus anthracis |
| 149 | 0.00 | 24 | 24 | S1 | 261594 | Bacillus anthracis str. 'Ames Ancestor' |
| 150 | 0.00 | 60 | 0 | F | 90964 | Staphylococcaceae |
| 151 | 0.00 | 60 | 0 | G | 1279 | Staphylococcus |
| 152 | 0.00 | 60 | 60 | S | 1280 | Staphylococcus aureus |
| 153 | 0.00 | 62 | 0 | O | 186826 | Lactobacillales |
| 154 | 0.00 | 62 | 0 | F | 1300 | Streptococcaceae |
| 155 | 0.00 | 62 | 36 | G | 1301 | Streptococcus |
| 156 | 0.00 | 6 | 6 | S | 1305 | Streptococcus sanguinis |
| 157 | 0.00 | 4 | 3 | S | 1302 | Streptococcus gordonii |
| 158 | 0.00 | 1 | 0 | S1 | 29390 | Streptococcus gordonii str. Challis |
| 159 | 0.00 | 1 | 1 | S2 | 467705 | Streptococcus gordonii str. Challis substr. CH1 |
| 160 | 0.00 | 4 | 4 | S | 1313 | Streptococcus pneumoniae |
| 161 | 0.00 | 3 | 3 | S | 1309 | Streptococcus mutans |
| 162 | 0.00 | 3 | 3 | S | 1314 | Streptococcus pyogenes |
| 163 | 0.00 | 2 | 2 | S | 1304 | Streptococcus salivarius |
| 164 | 0.00 | 2 | 0 | G1 | 119603 | Streptococcus dysgalactiae group |
| 165 | 0.00 | 1 | 1 | S | 1334 | Streptococcus dysgalactiae |
| 166 | 0.00 | 1 | 0 | S | 1336 | Streptococcus equi |
| 167 | 0.00 | 1 | 1 | S1 | 40041 | Streptococcus equi subsp. zooepidemicus |
| 168 | 0.00 | 1 | 1 | S | 1311 | Streptococcus agalactiae |
| 169 | 0.00 | 1 | 0 | G1 | 671232 | Streptococcus anginosus group |
| 170 | 0.00 | 1 | 1 | S | 1328 | Streptococcus anginosus |
| 171 | 0.00 | 175 | 0 | C | 186801 | Clostridia |
| 172 | 0.00 | 175 | 4 | O | 186802 | Eubacteriales |
| 173 | 0.00 | 109 | 0 | F | 31979 | Clostridiaceae |
| 174 | 0.00 | 109 | 48 | G | 1485 | Clostridium |
| 175 | 0.00 | 19 | 0 | S | 1491 | Clostridium botulinum |
| 176 | 0.00 | 19 | 0 | S1 | 36830 | Clostridium botulinum E |
| 177 | 0.00 | 19 | 19 | S2 | 508767 | Clostridium botulinum E3 str. Alaska E43 |
| 178 | 0.00 | 16 | 16 | S | 1488 | Clostridium acetobutylicum |
| 179 | 0.00 | 16 | 16 | S | 1502 | Clostridium perfringens |
| 180 | 0.00 | 9 | 8 | S | 1513 | Clostridium tetani |
| 181 | 0.00 | 1 | 1 | S1 | 212717 | Clostridium tetani E88 |
| 182 | 0.00 | 1 | 1 | S | 1509 | Clostridium sporogenes |
| 183 | 0.00 | 62 | 0 | F | 186804 | Peptostreptococcaceae |
| 184 | 0.00 | 62 | 0 | G | 1257 | Peptostreptococcus |
| 185 | 0.00 | 62 | 62 | S | 1261 | Peptostreptococcus anaerobius |
| 186 | 0.00 | 38 | 0 | C | 909932 | Negativicutes |
| 187 | 0.00 | 38 | 0 | O | 1843489 | Veillonellales |
| 188 | 0.00 | 38 | 0 | F | 31977 | Veillonellaceae |
| 189 | 0.00 | 38 | 0 | G | 29465 | Veillonella |
| 190 | 0.00 | 38 | 38 | S | 29466 | Veillonella parvula |
| 191 | 0.00 | 27 | 0 | C | 1737404 | Tissierellia |
| 192 | 0.00 | 27 | 0 | O | 1737405 | Tissierellales |
| 193 | 0.00 | 27 | 0 | F | 1570339 | Peptoniphilaceae |
| 194 | 0.00 | 27 | 0 | G | 543311 | Parvimonas |
| 195 | 0.00 | 27 | 27 | S | 33033 | Parvimonas micra |
| 196 | 0.00 | 8 | 0 | P | 544448 | Mycoplasmatota |
| 197 | 0.00 | 8 | 0 | O | 2790996 | Mycoplasmoidales |
| 198 | 0.00 | 8 | 0 | F | 2895623 | Metamycoplasmataceae |
| 199 | 0.00 | 8 | 0 | G | 2995234 | Mycoplasmoides |
| 200 | 0.00 | 8 | 0 | S | 2104 | Mycoplasmoides pneumoniae |
| 201 | 0.00 | 8 | 8 | S1 | 1441379 | Mycoplasmoides pneumoniae M29 |
| 202 | 0.00 | 254 | 0 | P | 203691 | Spirochaetota |
| 203 | 0.00 | 254 | 1 | C | 203692 | Spirochaetia |
| 204 | 0.00 | 142 | 0 | O | 1643688 | Leptospirales |
| 205 | 0.00 | 142 | 0 | F | 170 | Leptospiraceae |
| 206 | 0.00 | 142 | 62 | G | 171 | Leptospira |
| 207 | 0.00 | 17 | 17 | S | 408139 | Leptospira kmetyi |
| 208 | 0.00 | 12 | 0 | S | 48782 | Leptospira fainei |
| 209 | 0.00 | 12 | 0 | S1 | 293072 | Leptospira fainei serovar Hurstbridge |
| 210 | 0.00 | 12 | 12 | S2 | 1193011 | Leptospira fainei serovar Hurstbridge str. BUT 6 |
| 211 | 0.00 | 11 | 11 | S | 447106 | Leptospira licerasiae |
| 212 | 0.00 | 10 | 0 | S | 29506 | Leptospira inadai |
| 213 | 0.00 | 10 | 10 | S1 | 293084 | Leptospira inadai serovar Lyme |
| 214 | 0.00 | 9 | 9 | S | 28182 | Leptospira noguchii |
| 215 | 0.00 | 6 | 6 | S | 409998 | Leptospira wolffii |
| 216 | 0.00 | 4 | 4 | S | 28183 | Leptospira santarosai |
| 217 | 0.00 | 3 | 3 | S | 173 | Leptospira interrogans |
| 218 | 0.00 | 3 | 3 | S | 29507 | Leptospira kirschneri |
| 219 | 0.00 | 2 | 2 | S | 28452 | Leptospira alstonii |
| 220 | 0.00 | 1 | 1 | S | 174 | Leptospira borgpetersenii |
| 221 | 0.00 | 1 | 1 | S | 100053 | Leptospira alexanderi |
| 222 | 0.00 | 1 | 0 | S | 301541 | Leptospira broomii |
| 223 | 0.00 | 1 | 0 | S1 | 1324404 | Leptospira broomii serovar Hurstbridge |

| | | | | | | |
|---|---|---|---|---|---|---|
| 224 | 0.00 | 1 | 1 | S2 | 1049789 | Leptospira broomii serovar Hurstbridge str. 5399 |
| 225 | 0.00 | 111 | 6 | O | 136 | Spirochaetales |
| 226 | 0.00 | 82 | 0 | F | 2845253 | Treponemataceae |
| 227 | 0.00 | 82 | 7 | G | 157 | Treponema |
| 228 | 0.00 | 49 | 49 | S | 156 | Treponema zuelzerae |
| 229 | 0.00 | 15 | 15 | S | 160 | Treponema pallidum |
| 230 | 0.00 | 11 | 8 | S | 158 | Treponema denticola |
| 231 | 0.00 | 3 | 3 | S1 | 243275 | Treponema denticola ATCC 35405 |
| 232 | 0.00 | 23 | 7 | F | 1643685 | Borreliaceae |
| 233 | 0.00 | 9 | 7 | G | 64895 | Borreliella |
| 234 | 0.00 | 2 | 0 | S | 139 | Borreliella burgdorferi |
| 235 | 0.00 | 2 | 2 | S1 | 445984 | Borreliella burgdorferi Bol26 |
| 236 | 0.00 | 7 | 0 | G | 138 | Borrelia |
| 237 | 0.00 | 7 | 0 | S | 44449 | Borrelia recurrentis |
| 238 | 0.00 | 7 | 7 | S1 | 412418 | Borrelia recurrentis A1 |
| 239 | 0.00 | 77 | 0 | D1 | 1783270 | FCB group |
| 240 | 0.00 | 77 | 0 | D2 | 68336 | Bacteroidota/Chlorobiota group |
| 241 | 0.00 | 77 | 0 | P | 976 | Bacteroidota |
| 242 | 0.00 | 77 | 0 | C | 200643 | Bacteroidia |
| 243 | 0.00 | 77 | 25 | O | 171549 | Bacteroidales |
| 244 | 0.00 | 27 | 0 | F | 2005525 | Tannerellaceae |
| 245 | 0.00 | 27 | 0 | G | 195950 | Tannerella |
| 246 | 0.00 | 27 | 27 | S | 28112 | Tannerella forsythia |
| 247 | 0.00 | 25 | 0 | F | 171551 | Porphyromonadaceae |
| 248 | 0.00 | 25 | 0 | G | 836 | Porphyromonas |
| 249 | 0.00 | 25 | 20 | S | 837 | Porphyromonas gingivalis |
| 250 | 0.00 | 5 | 5 | S1 | 242619 | Porphyromonas gingivalis W83 |
| 251 | 0.00 | 47 | 0 | P | 32066 | Fusobacteriota |
| 252 | 0.00 | 47 | 0 | C | 203490 | Fusobacteriia |
| 253 | 0.00 | 47 | 0 | O | 203491 | Fusobacteriales |
| 254 | 0.00 | 47 | 0 | F | 203492 | Fusobacteriaceae |
| 255 | 0.00 | 47 | 15 | G | 848 | Fusobacterium |
| 256 | 0.00 | 24 | 24 | S | 851 | Fusobacterium nucleatum |
| 257 | 0.00 | 8 | 8 | S | 849 | Fusobacterium gonidiaformans |
| 258 | 0.00 | 44 | 0 | D1 | 1783257 | PVC group |
| 259 | 0.00 | 44 | 0 | P | 204428 | Chlamydiota |
| 260 | 0.00 | 44 | 0 | C | 204429 | Chlamydiia |
| 261 | 0.00 | 44 | 0 | O | 51291 | Chlamydiales |
| 262 | 0.00 | 44 | 0 | F | 809 | Chlamydiaceae |
| 263 | 0.00 | 44 | 0 | F1 | 1113537 | Chlamydia/Chlamydophila group |
| 264 | 0.00 | 44 | 39 | G | 810 | Chlamydia |
| 265 | 0.00 | 2 | 0 | S | 83558 | Chlamydia pneumoniae |
| 266 | 0.00 | 2 | 2 | S1 | 406984 | Chlamydia pneumoniae LPCoLN |
| 267 | 0.00 | 2 | 2 | S | 85991 | Chlamydia pecorum |
| 268 | 0.00 | 1 | 0 | S | 83554 | Chlamydia psittaci |
| 269 | 0.00 | 1 | 1 | S1 | 331636 | Chlamydia psittaci 6BC |
| 270 | 0.00 | 41 | 0 | P | 29547 | Campylobacterota |
| 271 | 0.00 | 41 | 0 | C | 3031852 | Epsilonproteobacteria |
| 272 | 0.00 | 41 | 0 | O | 213849 | Campylobacterales |
| 273 | 0.00 | 41 | 0 | F | 72293 | Helicobacteraceae |
| 274 | 0.00 | 41 | 0 | G | 209 | Helicobacter |
| 275 | 0.00 | 41 | 41 | S | 210 | Helicobacter pylori |
| 276 | 0.02 | 1974 | 123 | D | 2759 | Eukaryota |
| 277 | 0.01 | 982 | 0 | D1 | 2698737 | Sar |
| 278 | 0.01 | 982 | 0 | D2 | 33630 | Alveolata |
| 279 | 0.01 | 982 | 10 | P | 5794 | Apicomplexa |
| 280 | 0.01 | 582 | 0 | C | 422676 | Aconoidasida |
| 281 | 0.01 | 582 | 0 | O | 5819 | Haemosporida |
| 282 | 0.01 | 582 | 0 | F | 1639119 | Plasmodiidae |
| 283 | 0.01 | 582 | 1 | G | 5820 | Plasmodium |
| 284 | 0.01 | 577 | 0 | G1 | 418103 | Plasmodium (Plasmodium) |
| 285 | 0.01 | 577 | 577 | S | 5855 | Plasmodium vivax |
| 286 | 0.00 | 4 | 0 | G1 | 418107 | Plasmodium (Laverania) |
| 287 | 0.00 | 4 | 0 | S | 5833 | Plasmodium falciparum |
| 288 | 0.00 | 4 | 4 | S1 | 36329 | Plasmodium falciparum 3D7 |
| 289 | 0.00 | 390 | 0 | C | 1280412 | Conoidasida |
| 290 | 0.00 | 390 | 0 | C1 | 5796 | Coccidia |
| 291 | 0.00 | 390 | 0 | O | 75739 | Eucoccidiorida |
| 292 | 0.00 | 390 | 0 | O1 | 423054 | Eimeriorina |
| 293 | 0.00 | 390 | 0 | F | 5809 | Sarcocystidae |
| 294 | 0.00 | 390 | 0 | G | 5810 | Toxoplasma |
| 295 | 0.00 | 390 | 0 | S | 5811 | Toxoplasma gondii |
| 296 | 0.00 | 390 | 390 | S1 | 508771 | Toxoplasma gondii ME49 |
| 297 | 0.01 | 847 | 25 | D1 | 33154 | Opisthokonta |
| 298 | 0.01 | 743 | 0 | K | 33208 | Metazoa |
| 299 | 0.01 | 743 | 0 | K1 | 6072 | Eumetazoa |
| 300 | 0.01 | 743 | 0 | K2 | 33213 | Bilateria |
| 301 | 0.01 | 743 | 7 | K3 | 33317 | Protostomia |
| 302 | 0.01 | 697 | 0 | K4 | 2697495 | Spiralia |
| 303 | 0.01 | 697 | 0 | K5 | 1206795 | Lophotrochozoa |
| 304 | 0.01 | 697 | 10 | P | 6157 | Platyhelminthes |
| 305 | 0.01 | 595 | 0 | C | 6199 | Cestoda |
| 306 | 0.01 | 595 | 0 | C1 | 6200 | Eucestoda |
| 307 | 0.01 | 595 | 0 | O | 6201 | Cyclophyllidea |
| 308 | 0.01 | 595 | 0 | F | 6208 | Taeniidae |
| 309 | 0.01 | 595 | 0 | G | 6202 | Taenia |
| 310 | 0.01 | 595 | 595 | S | 6204 | Taenia solium |
| 311 | 0.00 | 92 | 0 | C | 6178 | Trematoda |
| 312 | 0.00 | 92 | 0 | C1 | 6179 | Digenea |
| 313 | 0.00 | 92 | 0 | O | 6180 | Strigeidida |
| 314 | 0.00 | 92 | 0 | O1 | 31244 | Schistosomatoidea |
| 315 | 0.00 | 92 | 0 | F | 31245 | Schistosomatidae |
| 316 | 0.00 | 92 | 0 | G | 6181 | Schistosoma |
| 317 | 0.00 | 92 | 92 | S | 6183 | Schistosoma mansoni |
| 318 | 0.00 | 39 | 0 | K4 | 1206794 | Ecdysozoa |
| 319 | 0.00 | 39 | 0 | P | 6231 | Nematoda |
| 320 | 0.00 | 39 | 0 | C | 119088 | Enoplea |
| 321 | 0.00 | 39 | 0 | C1 | 1457286 | Dorylaimia |
| 322 | 0.00 | 39 | 0 | O | 6329 | Trichinellida |
| 323 | 0.00 | 39 | 0 | F | 6332 | Trichinellidae |
| 324 | 0.00 | 39 | 0 | G | 6333 | Trichinella |
| 325 | 0.00 | 39 | 39 | S | 6334 | Trichinella spiralis |
| 326 | 0.00 | 79 | 0 | K | 4751 | Fungi |
| 327 | 0.00 | 79 | 5 | K1 | 451864 | Dikarya |
| 328 | 0.00 | 51 | 0 | P | 4890 | Ascomycota |
| 329 | 0.00 | 43 | 0 | P1 | 716545 | saccharomyceta |
| 330 | 0.00 | 43 | 0 | P2 | 147538 | Pezizomycotina |
| 331 | 0.00 | 43 | 0 | P3 | 716546 | leotiomyceta |
| 332 | 0.00 | 43 | 0 | C | 147545 | Eurotiomycetes |
| 333 | 0.00 | 43 | 1 | C1 | 451871 | Eurotiomycetidae |
| 334 | 0.00 | 28 | 1 | O | 33183 | Onygenales |
| 335 | 0.00 | 20 | 1 | F | 299071 | Ajellomycetaceae |
| 336 | 0.00 | 17 | 0 | G | 229219 | Blastomyces |

| | | | | | | |
|---|---|---|---|---|---|---|
| 337 | 0.00 | 17 | 0 | S | 5039 | Blastomyces dermatitidis |
| 338 | 0.00 | 17 | 17 | S1 | 559297 | Blastomyces dermatitidis ER-3 |
| 339 | 0.00 | 2 | 0 | G | 5036 | Histoplasma |
| 340 | 0.00 | 2 | 0 | S | 5037 | Histoplasma capsulatum |
| 341 | 0.00 | 2 | 2 | S1 | 447093 | Histoplasma capsulatum G186AR |
| 342 | 0.00 | 7 | 0 | F | 33184 | Onygenaceae |
| 343 | 0.00 | 7 | 0 | G | 5500 | Coccidioides |
| 344 | 0.00 | 5 | 0 | S | 5501 | Coccidioides immitis |
| 345 | 0.00 | 5 | 5 | S1 | 246410 | Coccidioides immitis RS |
| 346 | 0.00 | 2 | 0 | S | 199306 | Coccidioides posadasii |
| 347 | 0.00 | 2 | 2 | S1 | 222929 | Coccidioides posadasii C735 delta SOWgp |
| 348 | 0.00 | 14 | 0 | O | 5042 | Eurotiales |
| 349 | 0.00 | 14 | 0 | F | 1131492 | Aspergillaceae |
| 350 | 0.00 | 14 | 0 | G | 5052 | Aspergillus |
| 351 | 0.00 | 14 | 0 | G1 | 2720872 | Aspergillus subgen. Fumigati |
| 352 | 0.00 | 14 | 0 | S | 746128 | Aspergillus fumigatus |
| 353 | 0.00 | 14 | 14 | S1 | 330879 | Aspergillus fumigatus Af293 |
| 354 | 0.00 | 8 | 0 | P1 | 451866 | Taphrinomycotina |
| 355 | 0.00 | 8 | 0 | C | 147553 | Pneumocystidomycetes |
| 356 | 0.00 | 8 | 0 | O | 37987 | Pneumocystidales |
| 357 | 0.00 | 8 | 0 | F | 44281 | Pneumocystidaceae |
| 358 | 0.00 | 8 | 1 | G | 4753 | Pneumocystis |
| 359 | 0.00 | 5 | 0 | S | 4754 | Pneumocystis carinii |
| 360 | 0.00 | 5 | 5 | S1 | 1408658 | Pneumocystis carinii B80 |
| 361 | 0.00 | 2 | 0 | S | 42068 | Pneumocystis jirovecii |
| 362 | 0.00 | 2 | 2 | S1 | 1408657 | Pneumocystis jirovecii RU7 |
| 363 | 0.00 | 23 | 0 | P | 5204 | Basidiomycota |
| 364 | 0.00 | 23 | 0 | P1 | 5302 | Agaricomycotina |
| 365 | 0.00 | 23 | 0 | C | 155616 | Tremellomycetes |
| 366 | 0.00 | 23 | 0 | O | 5234 | Tremellales |
| 367 | 0.00 | 23 | 0 | F | 1884633 | Cryptococcaceae |
| 368 | 0.00 | 23 | 11 | G | 5206 | Cryptococcus |
| 369 | 0.00 | 8 | 0 | G1 | 1884637 | Cryptococcus gattii species complex |
| 370 | 0.00 | 8 | 0 | S | 37769 | Cryptococcus gattii VGI |
| 371 | 0.00 | 8 | 8 | S1 | 367775 | Cryptococcus gattii WM276 |
| 372 | 0.00 | 4 | 0 | G1 | 1897064 | Cryptococcus neoformans species complex |
| 373 | 0.00 | 4 | 0 | S | 5207 | Cryptococcus neoformans |
| 374 | 0.00 | 4 | 0 | S1 | 40410 | Cryptococcus neoformans var. neoformans |
| 375 | 0.00 | 4 | 4 | S2 | 214684 | Cryptococcus neoformans var. neoformans JEC21 |
| 376 | 0.00 | 22 | 0 | D1 | 2611352 | Discoba |
| 377 | 0.00 | 22 | 0 | P | 33682 | Euglenozoa |
| 378 | 0.00 | 22 | 0 | C | 5653 | Kinetoplastea |
| 379 | 0.00 | 22 | 0 | C1 | 2704647 | Metakinetoplastina |
| 380 | 0.00 | 22 | 0 | O | 2704949 | Trypanosomatida |
| 381 | 0.00 | 22 | 0 | F | 5654 | Trypanosomatidae |
| 382 | 0.00 | 22 | 0 | G | 5690 | Trypanosoma |
| 383 | 0.00 | 14 | 0 | G1 | 47570 | Schizotrypanum |
| 384 | 0.00 | 14 | 14 | S | 5693 | Trypanosoma cruzi |
| 385 | 0.00 | 8 | 0 | G1 | 39700 | Trypanozoon |
| 386 | 0.00 | 8 | 0 | S | 5691 | Trypanosoma brucei |
| 387 | 0.00 | 8 | 0 | S1 | 5702 | Trypanosoma brucei brucei |
| 388 | 0.00 | 8 | 8 | S2 | 185431 | Trypanosoma brucei brucei TREU927 |
| 389 | 0.00 | 2 | 0 | D | 2157 | Archaea |
| 390 | 0.00 | 2 | 0 | P | 28890 | Euryarchaeota |
| 391 | 0.00 | 2 | 0 | P1 | 2283794 | Methanomada group |
| 392 | 0.00 | 2 | 0 | C | 183925 | Methanobacteria |
| 393 | 0.00 | 2 | 0 | O | 2158 | Methanobacteriales |
| 394 | 0.00 | 2 | 0 | F | 2159 | Methanobacteriaceae |
| 395 | 0.00 | 2 | 0 | G | 2172 | Methanobrevibacter |
| 396 | 0.00 | 2 | 2 | S | 66851 | Methanobrevibacter oralis |
| 397 | 0.00 | 251 | 0 | D | 10239 | Viruses |
| 398 | 0.00 | 251 | 0 | D1 | 2731341 | Duplodnaviria |
| 399 | 0.00 | 251 | 0 | K | 2731360 | Heunggongvirae |
| 400 | 0.00 | 251 | 0 | P | 2731361 | Peploviricota |
| 401 | 0.00 | 251 | 0 | C | 2731363 | Herviviricetes |
| 402 | 0.00 | 251 | 0 | O | 548681 | Herpesvirales |
| 403 | 0.00 | 251 | 0 | F | 10292 | Herpesviridae |
| 404 | 0.00 | 251 | 0 | F1 | 10293 | Alphaherpesvirinae |
| 405 | 0.00 | 251 | 0 | G | 10319 | Varicellovirus |
| 406 | 0.00 | 250 | 0 | S | 3050278 | Varicellovirus equidalpha4 |
| 407 | 0.00 | 250 | 250 | S1 | 10331 | Equid alphaherpesvirus 4 |
| 408 | 0.00 | 1 | 0 | S | 3050277 | Varicellovirus equidalpha3 |
| 409 | 0.00 | 1 | 1 | S1 | 80341 | Equid alphaherpesvirus 3 |

**Table S3b:** Results of the Kraken2 analysis for library ERR6466109 based on a custom-built pathogen list (Table S1)

| Sr | Percentage of fragments covered | Number of fragments covered | Number of fragments assigned | rank code | NCBI taxonomic ID | Name |
|---|---|---|---|---|---|---|
| 1 | 99.75 | 123408800 | 123408800 | U | 0 | unclassified |
| 2 | 0.25 | 308909 | 1 | R | 1 | root |
| 3 | 0.25 | 305681 | 1978 | R1 | 131567 | cellular organisms |
| 4 | 0.23 | 280226 | 8128 | D | 2 | Bacteria |
| 5 | 0.17 | 209926 | 5858 | P | 1224 | Pseudomonadota |
| 6 | 0.12 | 151362 | 1388 | C | 28216 | Betaproteobacteria |
| 7 | 0.12 | 146158 | 5285 | O | 80840 | Burkholderiales |
| 8 | 0.07 | 91170 | 0 | F | 119060 | Burkholderiaceae |
| 9 | 0.07 | 91170 | 13104 | G | 32008 | Burkholderia |
| 10 | 0.04 | 43363 | 0 | G1 | 87882 | Burkholderia cepacia complex |
| 11 | 0.04 | 43363 | 43363 | S | 292 | Burkholderia cepacia |
| 12 | 0.03 | 34703 | 29216 | G1 | 111527 | pseudomallei group |
| 13 | 0.00 | 4111 | 4111 | S | 28450 | Burkholderia pseudomallei |
| 14 | 0.00 | 1376 | 1376 | S | 13373 | Burkholderia mallei |
| 15 | 0.04 | 49703 | 443 | F | 506 | Alcaligenaceae |
| 16 | 0.04 | 48647 | 4708 | G | 517 | Bordetella |
| 17 | 0.02 | 22837 | 22837 | S | 520 | Bordetella pertussis |
| 18 | 0.02 | 21102 | 21102 | S | 94624 | Bordetella petrii |
| 19 | 0.00 | 613 | 0 | G | 29574 | Taylorella |
| 20 | 0.00 | 613 | 613 | S | 29575 | Taylorella equigenitalis |
| 21 | 0.00 | 3816 | 0 | O | 206351 | Neisseriales |
| 22 | 0.00 | 3816 | 0 | F | 481 | Neisseriaceae |
| 23 | 0.00 | 3816 | 1808 | G | 482 | Neisseria |
| 24 | 0.00 | 911 | 911 | S | 485 | Neisseria gonorrhoeae |
| 25 | 0.00 | 600 | 600 | S | 487 | Neisseria meningitidis |
| 26 | 0.00 | 497 | 497 | S | 483 | Neisseria cinerea |
| 27 | 0.03 | 41762 | 3516 | C | 1236 | Gammaproteobacteria |
| 28 | 0.02 | 27230 | 925 | O | 91347 | Enterobacterales |
| 29 | 0.02 | 24542 | 5390 | F | 543 | Enterobacteriaceae |
| 30 | 0.01 | 13807 | 0 | F1 | 2890311 | Klebsiella/Raoultella group |
| 31 | 0.01 | 13807 | 1128 | G | 570 | Klebsiella |
| 32 | 0.01 | 7481 | 6976 | S | 573 | Klebsiella pneumoniae |
| 33 | 0.00 | 505 | 505 | S1 | 39831 | Klebsiella pneumoniae subsp. rhinoscleromatis |
| 34 | 0.00 | 5198 | 5198 | S | 571 | Klebsiella oxytoca |
| 35 | 0.00 | 3754 | 0 | G | 590 | Salmonella |
| 36 | 0.00 | 3754 | 0 | S | 28901 | Salmonella enterica |
| 37 | 0.00 | 3754 | 0 | S1 | 59201 | Salmonella enterica subsp. enterica |
| 38 | 0.00 | 3754 | 3754 | S2 | 594 | Salmonella enterica subsp. enterica serovar Gallinarum |
| 39 | 0.00 | 1340 | 0 | G | 561 | Escherichia |
| 40 | 0.00 | 1340 | 983 | S | 562 | Escherichia coli |
| 41 | 0.00 | 285 | 285 | S1 | 83334 | Escherichia coli O157:H7 |
| 42 | 0.00 | 54 | 54 | S1 | 168927 | Escherichia coli O111:H- |
| 43 | 0.00 | 18 | 18 | S1 | 244319 | Escherichia coli O26:H11 |
| 44 | 0.00 | 251 | 107 | G | 620 | Shigella |
| 45 | 0.00 | 111 | 111 | S | 623 | Shigella flexneri |
| 46 | 0.00 | 14 | 14 | S | 621 | Shigella boydii |
| 47 | 0.00 | 13 | 13 | S | 622 | Shigella dysenteriae |
| 48 | 0.00 | 6 | 6 | S | 624 | Shigella sonnei |
| 49 | 0.00 | 1763 | 0 | F | 1903411 | Yersiniaceae |
| 50 | 0.00 | 1763 | 754 | G | 629 | Yersinia |
| 51 | 0.00 | 574 | 380 | G1 | 1649845 | Yersinia pseudotuberculosis complex |
| 52 | 0.00 | 172 | 172 | S | 632 | Yersinia pestis |
| 53 | 0.00 | 22 | 22 | S | 633 | Yersinia pseudotuberculosis |
| 54 | 0.00 | 435 | 435 | S | 630 | Yersinia enterocolitica |
| 55 | 0.00 | 5163 | 318 | O | 118969 | Legionellales |
| 56 | 0.00 | 2846 | 0 | F | 118968 | Coxiellaceae |
| 57 | 0.00 | 2846 | 0 | G | 776 | Coxiella |
| 58 | 0.00 | 2846 | 2846 | S | 777 | Coxiella burnetii |
| 59 | 0.00 | 1999 | 0 | F | 444 | Legionellaceae |
| 60 | 0.00 | 1999 | 0 | G | 445 | Legionella |
| 61 | 0.00 | 1999 | 1999 | S | 446 | Legionella pneumophila |
| 62 | 0.00 | 2111 | 0 | O | 2887326 | Moraxellales |
| 63 | 0.00 | 2111 | 0 | F | 468 | Moraxellaceae |
| 64 | 0.00 | 2111 | 0 | G | 475 | Moraxella |
| 65 | 0.00 | 2111 | 2111 | S | 480 | Moraxella catarrhalis |
| 66 | 0.00 | 1790 | 0 | O | 135623 | Vibrionales |
| 67 | 0.00 | 1790 | 0 | F | 641 | Vibrionaceae |
| 68 | 0.00 | 1790 | 559 | G | 662 | Vibrio |
| 69 | 0.00 | 483 | 483 | S | 666 | Vibrio cholerae |
| 70 | 0.00 | 470 | 0 | G1 | 717610 | Vibrio harveyi group |
| 71 | 0.00 | 470 | 470 | S | 670 | Vibrio parahaemolyticus |
| 72 | 0.00 | 278 | 278 | S | 672 | Vibrio vulnificus |
| 73 | 0.00 | 1055 | 0 | O | 72273 | Thiotrichales |
| 74 | 0.00 | 1055 | 0 | F | 34064 | Francisellaceae |
| 75 | 0.00 | 1055 | 0 | G | 262 | Francisella |
| 76 | 0.00 | 1055 | 1055 | S | 263 | Francisella tularensis |
| 77 | 0.00 | 897 | 0 | O | 135625 | Pasteurellales |
| 78 | 0.00 | 897 | 266 | F | 712 | Pasteurellaceae |
| 79 | 0.00 | 318 | 0 | G | 713 | Actinobacillus |
| 80 | 0.00 | 318 | 0 | S | 718 | Actinobacillus equuli |
| 81 | 0.00 | 318 | 318 | S1 | 202947 | Actinobacillus equuli subsp. equuli |
| 82 | 0.00 | 313 | 0 | G | 724 | Haemophilus |
| 83 | 0.00 | 313 | 0 | S | 727 | Haemophilus influenzae |
| 84 | 0.00 | 313 | 313 | S1 | 725 | Haemophilus influenzae biotype aegyptius |
| 85 | 0.01 | 10944 | 116 | C | 28211 | Alphaproteobacteria |
| 86 | 0.01 | 10136 | 1509 | O | 356 | Hyphomicrobiales |
| 87 | 0.01 | 7048 | 0 | F | 118882 | Brucellaceae |
| 88 | 0.01 | 7048 | 0 | F1 | 2826938 | Brucella/Ochrobactrum group |
| 89 | 0.01 | 7048 | 6797 | G | 234 | Brucella |
| 90 | 0.00 | 153 | 0 | S | 236 | Brucella ovis |
| 91 | 0.00 | 153 | 153 | S1 | 444178 | Brucella ovis ATCC 25840 |
| 92 | 0.00 | 55 | 0 | S | 444163 | Brucella microti |
| 93 | 0.00 | 55 | 55 | S1 | 568815 | Brucella microti CCM 4915 |
| 94 | 0.00 | 34 | 0 | S | 29459 | Brucella melitensis |
| 95 | 0.00 | 34 | 0 | S1 | 644337 | Brucella melitensis bv. 1 |
| 96 | 0.00 | 34 | 34 | S2 | 224914 | Brucella melitensis bv. 1 str. 16M |
| 97 | 0.00 | 8 | 0 | S | 235 | Brucella abortus |
| 98 | 0.00 | 8 | 8 | S1 | 359391 | Brucella abortus 2308 |
| 99 | 0.00 | 1 | 0 | S | 29461 | Brucella suis |
| 100 | 0.00 | 1 | 1 | S1 | 204722 | Brucella suis 1330 |
| 101 | 0.00 | 1579 | 0 | F | 772 | Bartonellaceae |
| 102 | 0.00 | 1579 | 1017 | G | 773 | Bartonella |
| 103 | 0.00 | 220 | 220 | S | 38323 | Bartonella henselae |
| 104 | 0.00 | 171 | 0 | S | 774 | Bartonella bacilliformis |
| 105 | 0.00 | 171 | 171 | S1 | 360095 | Bartonella bacilliformis KC583 |
| 106 | 0.00 | 171 | 171 | S | 803 | Bartonella quintana |
| 107 | 0.00 | 692 | 0 | O | 766 | Rickettsiales |
| 108 | 0.00 | 692 | 0 | F | 775 | Rickettsiaceae |
| 109 | 0.00 | 692 | 0 | F1 | 33988 | Rickettsieae |
| 110 | 0.00 | 692 | 485 | G | 780 | Rickettsia |

| | | | | | | |
|---|---|---|---|---|---|---|
| 111 | 0.00 | 172 | 110 | G1 | 114277 | spotted fever group |
| 112 | 0.00 | 24 | 0 | S | 786 | Rickettsia akari |
| 113 | 0.00 | 24 | 24 | S1 | 293614 | Rickettsia akari str. Hartford |
| 114 | 0.00 | 19 | 19 | S | 42862 | Rickettsia felis |
| 115 | 0.00 | 18 | 18 | S | 35790 | Rickettsia japonica |
| 116 | 0.00 | 1 | 0 | G2 | 266068 | Rickettsia sibirica subgroup |
| 117 | 0.00 | 1 | 0 | S | 35793 | Rickettsia sibirica |
| 118 | 0.00 | 1 | 1 | S1 | 272951 | Rickettsia sibirica 246 |
| 119 | 0.00 | 35 | 1 | G1 | 114292 | typhus group |
| 120 | 0.00 | 22 | 22 | S | 782 | Rickettsia prowazekii |
| 121 | 0.00 | 12 | 12 | S | 785 | Rickettsia typhi |
| 122 | 0.05 | 56339 | 297 | D1 | 1783272 | Terrabacteria group |
| 123 | 0.04 | 47910 | 0 | P | 201174 | Actinomycetota |
| 124 | 0.04 | 47910 | 0 | C | 1760 | Actinomycetes |
| 125 | 0.04 | 47910 | 3885 | O | 85007 | Mycobacteriales |
| 126 | 0.03 | 32202 | 1244 | F | 1762 | Mycobacteriaceae |
| 127 | 0.02 | 27613 | 2895 | G | 1763 | Mycobacterium |
| 128 | 0.01 | 16651 | 3464 | G1 | 120793 | Mycobacterium avium complex (MAC) |
| 129 | 0.00 | 5434 | 5434 | S | 1764 | Mycobacterium avium |
| 130 | 0.00 | 3941 | 3941 | S | 1767 | Mycobacterium intracellulare |
| 131 | 0.00 | 3812 | 3812 | S | 339268 | Mycobacterium colombiense |
| 132 | 0.00 | 4502 | 4502 | S | 1768 | Mycobacterium kansasii |
| 133 | 0.00 | 3080 | 2612 | G1 | 77643 | Mycobacterium tuberculosis complex |
| 134 | 0.00 | 271 | 271 | S | 78331 | Mycobacterium canettii |
| 135 | 0.00 | 197 | 197 | S | 1773 | Mycobacterium tuberculosis |
| 136 | 0.00 | 485 | 485 | S | 1769 | Mycobacterium leprae |
| 137 | 0.00 | 3345 | 0 | G | 670516 | Mycobacteroides |
| 138 | 0.00 | 3345 | 0 | S | 36809 | Mycobacteroides abscessus |
| 139 | 0.00 | 3345 | 3345 | S1 | 319705 | Mycobacteroides abscessus subsp. bolletii |
| 140 | 0.01 | 9798 | 0 | F | 85025 | Nocardiaceae |
| 141 | 0.01 | 9798 | 0 | G | 2979332 | Prescottella |
| 142 | 0.01 | 9798 | 9798 | S | 43767 | Prescottella equi |
| 143 | 0.00 | 2025 | 0 | F | 1653 | Corynebacteriaceae |
| 144 | 0.00 | 2025 | 452 | G | 1716 | Corynebacterium |
| 145 | 0.00 | 1194 | 1194 | S | 1717 | Corynebacterium diphtheriae |
| 146 | 0.00 | 379 | 0 | S | 1719 | Corynebacterium pseudotuberculosis |
| 147 | 0.00 | 379 | 379 | S1 | 1087451 | Corynebacterium pseudotuberculosis 31 |
| 148 | 0.01 | 8043 | 696 | P | 1239 | Bacillota |
| 149 | 0.00 | 4474 | 344 | C | 91061 | Bacilli |
| 150 | 0.00 | 3309 | 251 | O | 1385 | Bacillales |
| 151 | 0.00 | 2285 | 0 | F | 186817 | Bacillaceae |
| 152 | 0.00 | 2285 | 0 | G | 1386 | Bacillus |
| 153 | 0.00 | 2285 | 1374 | G1 | 86661 | Bacillus cereus group |
| 154 | 0.00 | 564 | 564 | S | 1396 | Bacillus cereus |
| 155 | 0.00 | 347 | 0 | S | 1392 | Bacillus anthracis |
| 156 | 0.00 | 347 | 347 | S1 | 261594 | Bacillus anthracis str. 'Ames Ancestor' |
| 157 | 0.00 | 773 | 0 | F | 90964 | Staphylococcaceae |
| 158 | 0.00 | 773 | 0 | G | 1279 | Staphylococcus |
| 159 | 0.00 | 773 | 773 | S | 1280 | Staphylococcus aureus |
| 160 | 0.00 | 821 | 0 | O | 186826 | Lactobacillales |
| 161 | 0.00 | 821 | 0 | F | 1300 | Streptococcaceae |
| 162 | 0.00 | 821 | 423 | G | 1301 | Streptococcus |
| 163 | 0.00 | 78 | 0 | G1 | 119603 | Streptococcus dysgalactiae group |
| 164 | 0.00 | 68 | 0 | S | 1336 | Streptococcus equi |
| 165 | 0.00 | 68 | 68 | S1 | 40041 | Streptococcus equi subsp. zooepidemicus |
| 166 | 0.00 | 10 | 10 | S | 1334 | Streptococcus dysgalactiae |
| 167 | 0.00 | 51 | 42 | S | 1302 | Streptococcus gordonii |
| 168 | 0.00 | 9 | 0 | S1 | 29390 | Streptococcus gordonii str. Challis |
| 169 | 0.00 | 9 | 9 | S2 | 467705 | Streptococcus gordonii str. Challis substr. CH1 |
| 170 | 0.00 | 51 | 51 | S | 1305 | Streptococcus sanguinis |
| 171 | 0.00 | 47 | 47 | S | 1309 | Streptococcus mutans |
| 172 | 0.00 | 42 | 0 | G1 | 671232 | Streptococcus anginosus group |
| 173 | 0.00 | 42 | 42 | S | 1328 | Streptococcus anginosus |
| 174 | 0.00 | 40 | 40 | S | 1313 | Streptococcus pneumoniae |
| 175 | 0.00 | 38 | 38 | S | 1304 | Streptococcus salivarius |
| 176 | 0.00 | 20 | 20 | S | 1303 | Streptococcus oralis |
| 177 | 0.00 | 16 | 16 | S | 1314 | Streptococcus pyogenes |
| 178 | 0.00 | 15 | 15 | S | 1311 | Streptococcus agalactiae |
| 179 | 0.00 | 2051 | 0 | C | 186801 | Clostridia |
| 180 | 0.00 | 2051 | 52 | O | 186802 | Eubacteriales |
| 181 | 0.00 | 1325 | 0 | F | 31979 | Clostridiaceae |
| 182 | 0.00 | 1325 | 599 | G | 1485 | Clostridium |
| 183 | 0.00 | 242 | 242 | S | 1502 | Clostridium perfringens |
| 184 | 0.00 | 188 | 11 | S | 1491 | Clostridium botulinum |
| 185 | 0.00 | 177 | 0 | S1 | 36830 | Clostridium botulinum E |
| 186 | 0.00 | 177 | 177 | S2 | 508767 | Clostridium botulinum E3 str. Alaska E43 |
| 187 | 0.00 | 142 | 142 | S | 1513 | Clostridium tetani |
| 188 | 0.00 | 129 | 129 | S | 1488 | Clostridium acetobutylicum |
| 189 | 0.00 | 25 | 25 | S | 1509 | Clostridium sporogenes |
| 190 | 0.00 | 674 | 0 | F | 186804 | Peptostreptococcaceae |
| 191 | 0.00 | 674 | 0 | G | 1257 | Peptostreptococcus |
| 192 | 0.00 | 674 | 674 | S | 1261 | Peptostreptococcus anaerobius |
| 193 | 0.00 | 488 | 0 | C | 909932 | Negativicutes |
| 194 | 0.00 | 488 | 0 | O | 1843489 | Veillonellales |
| 195 | 0.00 | 488 | 0 | F | 31977 | Veillonellaceae |
| 196 | 0.00 | 488 | 0 | G | 29465 | Veillonella |
| 197 | 0.00 | 488 | 488 | S | 29466 | Veillonella parvula |
| 198 | 0.00 | 334 | 0 | C | 1737404 | Tissierellia |
| 199 | 0.00 | 334 | 0 | O | 1737405 | Tissierellales |
| 200 | 0.00 | 334 | 0 | F | 1570339 | Peptoniphilaceae |
| 201 | 0.00 | 334 | 0 | G | 543311 | Parvimonas |
| 202 | 0.00 | 334 | 334 | S | 33033 | Parvimonas micra |
| 203 | 0.00 | 89 | 0 | P | 544448 | Mycoplasmatota |
| 204 | 0.00 | 89 | 0 | O | 2790996 | Mycoplasmoidales |
| 205 | 0.00 | 89 | 0 | F | 2895623 | Metamycoplasmataceae |
| 206 | 0.00 | 89 | 0 | G | 2995234 | Mycoplasmoides |
| 207 | 0.00 | 89 | 0 | S | 2104 | Mycoplasmoides pneumoniae |
| 208 | 0.00 | 89 | 89 | S1 | 1441379 | Mycoplasmoides pneumoniae M29 |
| 209 | 0.00 | 3249 | 0 | P | 203691 | Spirochaetota |
| 210 | 0.00 | 3249 | 12 | C | 203692 | Spirochaetia |
| 211 | 0.00 | 1695 | 0 | O | 1643688 | Leptospirales |
| 212 | 0.00 | 1695 | 0 | F | 170 | Leptospiraceae |
| 213 | 0.00 | 1695 | 793 | G | 171 | Leptospira |
| 214 | 0.00 | 224 | 224 | S | 408139 | Leptospira kmetyi |
| 215 | 0.00 | 164 | 164 | S | 447106 | Leptospira licerasiae |
| 216 | 0.00 | 70 | 70 | S | 409998 | Leptospira wolffii |
| 217 | 0.00 | 63 | 63 | S | 28182 | Leptospira noguchii |
| 218 | 0.00 | 61 | 0 | S | 48782 | Leptospira fainei |
| 219 | 0.00 | 61 | 0 | S1 | 293072 | Leptospira fainei serovar Hurstbridge |
| 220 | 0.00 | 61 | 61 | S2 | 1193011 | Leptospira fainei serovar Hurstbridge str. BUT 6 |
| 221 | 0.00 | 58 | 58 | S | 28183 | Leptospira santarosai |
| 222 | 0.00 | 54 | 0 | S | 29506 | Leptospira inadai |
| 223 | 0.00 | 54 | 54 | S1 | 293084 | Leptospira inadai serovar Lyme |

| | | | | | | |
|---|---|---|---|---|---|---|
| 224 | 0.00 | 44 | 44 | S | 174 | Leptospira borgpetersenii |
| 225 | 0.00 | 34 | 34 | S | 28452 | Leptospira alstonii |
| 226 | 0.00 | 30 | 30 | S | 100053 | Leptospira alexanderi |
| 227 | 0.00 | 28 | 28 | S | 173 | Leptospira interrogans |
| 228 | 0.00 | 28 | 28 | S | 29507 | Leptospira kirschneri |
| 229 | 0.00 | 23 | 23 | S | 28184 | Leptospira weilii |
| 230 | 0.00 | 21 | 0 | S | 301541 | Leptospira broomii |
| 231 | 0.00 | 21 | 0 | S1 | 1324404 | Leptospira broomii serovar Hurstbridge |
| 232 | 0.00 | 21 | 21 | S2 | 1049789 | Leptospira broomii serovar Hurstbridge str. 5399 |
| 233 | 0.00 | 1542 | 69 | O | 136 | Spirochaetales |
| 234 | 0.00 | 1171 | 0 | F | 2845253 | Treponemataceae |
| 235 | 0.00 | 1171 | 131 | G | 157 | Treponema |
| 236 | 0.00 | 785 | 785 | S | 156 | Treponema zuelzerae |
| 237 | 0.00 | 132 | 130 | S | 160 | Treponema pallidum |
| 238 | 0.00 | 2 | 2 | S1 | 168 | Treponema pallidum subsp. pertenue |
| 239 | 0.00 | 123 | 123 | S | 158 | Treponema denticola |
| 240 | 0.00 | 302 | 69 | F | 1643685 | Borreliaceae |
| 241 | 0.00 | 142 | 121 | G | 64895 | Borreliella |
| 242 | 0.00 | 19 | 0 | S | 139 | Borreliella burgdorferi |
| 243 | 0.00 | 19 | 19 | S1 | 445984 | Borreliella burgdorferi Bol26 |
| 244 | 0.00 | 2 | 0 | S | 664662 | Borreliella bavariensis |
| 245 | 0.00 | 2 | 2 | S1 | 290434 | Borrelia garinii subsp. bavariensis PBi |
| 246 | 0.00 | 91 | 0 | G | 138 | Borrelia |
| 247 | 0.00 | 91 | 0 | S | 44449 | Borrelia recurrentis |
| 248 | 0.00 | 91 | 91 | S1 | 412418 | Borrelia recurrentis A1 |
| 249 | 0.00 | 1041 | 0 | D1 | 1783270 | FCB group |
| 250 | 0.00 | 1041 | 0 | D2 | 68336 | Bacteroidota/Chlorobiota group |
| 251 | 0.00 | 1041 | 0 | P | 976 | Bacteroidota |
| 252 | 0.00 | 1041 | 0 | C | 200643 | Bacteroidia |
| 253 | 0.00 | 1041 | 339 | O | 171549 | Bacteroidales |
| 254 | 0.00 | 428 | 0 | F | 2005525 | Tannerellaceae |
| 255 | 0.00 | 428 | 0 | G | 195950 | Tannerella |
| 256 | 0.00 | 428 | 406 | S | 28112 | Tannerella forsythia |
| 257 | 0.00 | 22 | 22 | S1 | 203275 | Tannerella forsythia 92A2 |
| 258 | 0.00 | 274 | 0 | F | 171551 | Porphyromonadaceae |
| 259 | 0.00 | 274 | 0 | G | 836 | Porphyromonas |
| 260 | 0.00 | 274 | 230 | S | 837 | Porphyromonas gingivalis |
| 261 | 0.00 | 44 | 44 | S1 | 242619 | Porphyromonas gingivalis W83 |
| 262 | 0.00 | 612 | 0 | D1 | 1783257 | PVC group |
| 263 | 0.00 | 612 | 0 | P | 204428 | Chlamydiota |
| 264 | 0.00 | 612 | 0 | C | 204429 | Chlamydiia |
| 265 | 0.00 | 612 | 0 | O | 51291 | Chlamydiales |
| 266 | 0.00 | 612 | 0 | F | 809 | Chlamydiaceae |
| 267 | 0.00 | 612 | 0 | F1 | 1113537 | Chlamydia/Chlamydophila group |
| 268 | 0.00 | 612 | 531 | G | 810 | Chlamydia |
| 269 | 0.00 | 32 | 0 | S | 83557 | Chlamydia caviae |
| 270 | 0.00 | 32 | 32 | S1 | 227941 | Chlamydia caviae GPIC |
| 271 | 0.00 | 21 | 0 | S | 83558 | Chlamydia pneumoniae |
| 272 | 0.00 | 21 | 21 | S1 | 406984 | Chlamydia pneumoniae LPCoLN |
| 273 | 0.00 | 9 | 9 | S | 83560 | Chlamydia muridarum |
| 274 | 0.00 | 8 | 8 | S | 85991 | Chlamydia pecorum |
| 275 | 0.00 | 6 | 0 | S | 83556 | Chlamydia felis |
| 276 | 0.00 | 6 | 6 | S1 | 264202 | Chlamydia felis Fe/C-56 |
| 277 | 0.00 | 3 | 0 | S | 83554 | Chlamydia psittaci |
| 278 | 0.00 | 3 | 3 | S1 | 331636 | Chlamydia psittaci 6BC |
| 279 | 0.00 | 2 | 2 | S | 83555 | Chlamydia abortus |
| 280 | 0.00 | 555 | 0 | P | 32066 | Fusobacteriota |
| 281 | 0.00 | 555 | 0 | C | 203490 | Fusobacteriia |
| 282 | 0.00 | 555 | 0 | O | 203491 | Fusobacteriales |
| 283 | 0.00 | 555 | 0 | F | 203492 | Fusobacteriaceae |
| 284 | 0.00 | 555 | 197 | G | 848 | Fusobacterium |
| 285 | 0.00 | 214 | 214 | S | 851 | Fusobacterium nucleatum |
| 286 | 0.00 | 144 | 144 | S | 849 | Fusobacterium gonidiaformans |
| 287 | 0.00 | 376 | 0 | P | 29547 | Campylobacterota |
| 288 | 0.00 | 376 | 0 | C | 3031852 | Epsilonproteobacteria |
| 289 | 0.00 | 376 | 0 | O | 213849 | Campylobacterales |
| 290 | 0.00 | 376 | 0 | F | 72293 | Helicobacteraceae |
| 291 | 0.00 | 376 | 0 | G | 209 | Helicobacter |
| 292 | 0.00 | 376 | 376 | S | 210 | Helicobacter pylori |
| 293 | 0.02 | 23431 | 1855 | D | 2759 | Eukaryota |
| 294 | 0.01 | 11107 | 0 | D1 | 2698737 | Sar |
| 295 | 0.01 | 11107 | 0 | D2 | 33630 | Alveolata |
| 296 | 0.01 | 11107 | 61 | P | 5794 | Apicomplexa |
| 297 | 0.01 | 6921 | 0 | C | 422676 | Aconoidasida |
| 298 | 0.01 | 6921 | 0 | O | 5819 | Haemosporida |
| 299 | 0.01 | 6921 | 0 | F | 1639119 | Plasmodiidae |
| 300 | 0.01 | 6921 | 25 | G | 5820 | Plasmodium |
| 301 | 0.01 | 6888 | 0 | G1 | 418103 | Plasmodium (Plasmodium) |
| 302 | 0.01 | 6888 | 6888 | S | 5855 | Plasmodium vivax |
| 303 | 0.00 | 8 | 0 | G1 | 418107 | Plasmodium (Laverania) |
| 304 | 0.00 | 8 | 0 | S | 5833 | Plasmodium falciparum |
| 305 | 0.00 | 8 | 8 | S1 | 36329 | Plasmodium falciparum 3D7 |
| 306 | 0.00 | 4125 | 0 | C | 1280412 | Conoidasida |
| 307 | 0.00 | 4125 | 0 | C1 | 5796 | Coccidia |
| 308 | 0.00 | 4125 | 0 | O | 75739 | Eucoccidiorida |
| 309 | 0.00 | 4125 | 0 | O1 | 423054 | Eimeriorina |
| 310 | 0.00 | 4125 | 0 | F | 5809 | Sarcocystidae |
| 311 | 0.00 | 4125 | 0 | G | 5810 | Toxoplasma |
| 312 | 0.00 | 4125 | 0 | S | 5811 | Toxoplasma gondii |
| 313 | 0.00 | 4113 | 4113 | S1 | 508771 | Toxoplasma gondii ME49 |
| 314 | 0.00 | 12 | 12 | S1 | 383379 | Toxoplasma gondii RH |
| 315 | 0.01 | 10121 | 241 | D1 | 33154 | Opisthokonta |
| 316 | 0.01 | 8927 | 0 | K | 33208 | Metazoa |
| 317 | 0.01 | 8927 | 0 | K1 | 6072 | Eumetazoa |
| 318 | 0.01 | 8927 | 0 | K2 | 33213 | Bilateria |
| 319 | 0.01 | 8927 | 59 | K3 | 33317 | Protostomia |
| 320 | 0.01 | 8498 | 0 | K4 | 2697495 | Spiralia |
| 321 | 0.01 | 8498 | 0 | K5 | 1206795 | Lophotrochozoa |
| 322 | 0.01 | 8498 | 112 | P | 6157 | Platyhelminthes |
| 323 | 0.01 | 7153 | 0 | C | 6199 | Cestoda |
| 324 | 0.01 | 7153 | 0 | C1 | 6200 | Eucestoda |
| 325 | 0.01 | 7153 | 0 | O | 6201 | Cyclophyllidea |
| 326 | 0.01 | 7153 | 0 | F | 6208 | Taeniidae |
| 327 | 0.01 | 7153 | 0 | G | 6202 | Taenia |
| 328 | 0.01 | 7153 | 7153 | S | 6204 | Taenia solium |
| 329 | 0.00 | 1233 | 0 | C | 6178 | Trematoda |
| 330 | 0.00 | 1233 | 0 | C1 | 6179 | Digenea |
| 331 | 0.00 | 1233 | 0 | O | 6180 | Strigeidida |
| 332 | 0.00 | 1233 | 0 | O1 | 31244 | Schistosomatoidea |
| 333 | 0.00 | 1233 | 0 | F | 31245 | Schistosomatidae |
| 334 | 0.00 | 1233 | 0 | G | 6181 | Schistosoma |
| 335 | 0.00 | 1233 | 1233 | S | 6183 | Schistosoma mansoni |
| 336 | 0.00 | 370 | 0 | K4 | 1206794 | Ecdysozoa |

| | | | | | | |
|---|---|---|---|---|---|---|
| 337 | 0.00 | 370 | 0 | P | 6231 | Nematoda |
| 338 | 0.00 | 370 | 0 | C | 119088 | Enoplea |
| 339 | 0.00 | 370 | 0 | C1 | 1457286 | Dorylaimia |
| 340 | 0.00 | 370 | 0 | O | 6329 | Trichinellida |
| 341 | 0.00 | 370 | 0 | F | 6332 | Trichinellidae |
| 342 | 0.00 | 370 | 0 | G | 6333 | Trichinella |
| 343 | 0.00 | 370 | 370 | S | 6334 | Trichinella spiralis |
| 344 | 0.00 | 953 | 0 | K | 4751 | Fungi |
| 345 | 0.00 | 953 | 89 | K1 | 451864 | Dikarya |
| 346 | 0.00 | 642 | 0 | P | 4890 | Ascomycota |
| 347 | 0.00 | 573 | 0 | P1 | 716545 | saccharomyceta |
| 348 | 0.00 | 573 | 0 | P2 | 147538 | Pezizomycotina |
| 349 | 0.00 | 573 | 0 | P3 | 716546 | leotiomyceta |
| 350 | 0.00 | 573 | 0 | C | 147545 | Eurotiomycetes |
| 351 | 0.00 | 573 | 34 | C1 | 451871 | Eurotiomycetidae |
| 352 | 0.00 | 394 | 7 | O | 33183 | Onygenales |
| 353 | 0.00 | 303 | 11 | F | 299071 | Ajellomycetaceae |
| 354 | 0.00 | 203 | 0 | G | 229219 | Blastomyces |
| 355 | 0.00 | 203 | 0 | S | 5039 | Blastomyces dermatitidis |
| 356 | 0.00 | 203 | 203 | S1 | 559297 | Blastomyces dermatitidis ER-3 |
| 357 | 0.00 | 89 | 0 | G | 5036 | Histoplasma |
| 358 | 0.00 | 89 | 0 | S | 5037 | Histoplasma capsulatum |
| 359 | 0.00 | 89 | 89 | S1 | 447093 | Histoplasma capsulatum G186AR |
| 360 | 0.00 | 84 | 0 | F | 33184 | Onygenaceae |
| 361 | 0.00 | 84 | 21 | G | 5500 | Coccidioides |
| 362 | 0.00 | 37 | 0 | S | 199306 | Coccidioides posadasii |
| 363 | 0.00 | 37 | 37 | S1 | 222929 | Coccidioides posadasii C735 delta SOWgp |
| 364 | 0.00 | 26 | 0 | S | 5501 | Coccidioides immitis |
| 365 | 0.00 | 26 | 26 | S1 | 246410 | Coccidioides immitis RS |
| 366 | 0.00 | 145 | 0 | O | 5042 | Eurotiales |
| 367 | 0.00 | 145 | 0 | F | 1131492 | Aspergillaceae |
| 368 | 0.00 | 145 | 0 | G | 5052 | Aspergillus |
| 369 | 0.00 | 145 | 0 | G1 | 2720872 | Aspergillus subgen. Fumigati |
| 370 | 0.00 | 145 | 0 | S | 746128 | Aspergillus fumigatus |
| 371 | 0.00 | 145 | 145 | S1 | 330879 | Aspergillus fumigatus Af293 |
| 372 | 0.00 | 69 | 0 | P1 | 451866 | Taphrinomycotina |
| 373 | 0.00 | 69 | 0 | C | 147553 | Pneumocystidomycetes |
| 374 | 0.00 | 69 | 0 | O | 37987 | Pneumocystidales |
| 375 | 0.00 | 69 | 0 | F | 44281 | Pneumocystidaceae |
| 376 | 0.00 | 69 | 12 | G | 4753 | Pneumocystis |
| 377 | 0.00 | 41 | 0 | S | 4754 | Pneumocystis carinii |
| 378 | 0.00 | 41 | 41 | S1 | 1408658 | Pneumocystis carinii B80 |
| 379 | 0.00 | 16 | 0 | S | 42068 | Pneumocystis jirovecii |
| 380 | 0.00 | 16 | 16 | S1 | 1408657 | Pneumocystis jirovecii RU7 |
| 381 | 0.00 | 222 | 0 | P | 5204 | Basidiomycota |
| 382 | 0.00 | 222 | 0 | P1 | 5302 | Agaricomycotina |
| 383 | 0.00 | 222 | 0 | C | 155616 | Tremellomycetes |
| 384 | 0.00 | 222 | 0 | O | 5234 | Tremellales |
| 385 | 0.00 | 222 | 0 | F | 1884633 | Cryptococcaceae |
| 386 | 0.00 | 222 | 139 | G | 5206 | Cryptococcus |
| 387 | 0.00 | 53 | 0 | G1 | 1884637 | Cryptococcus gattii species complex |
| 388 | 0.00 | 53 | 0 | S | 37769 | Cryptococcus gattii VGI |
| 389 | 0.00 | 53 | 53 | S1 | 367775 | Cryptococcus gattii WM276 |
| 390 | 0.00 | 30 | 0 | G1 | 1897064 | Cryptococcus neoformans species complex |
| 391 | 0.00 | 30 | 0 | S | 5207 | Cryptococcus neoformans |
| 392 | 0.00 | 30 | 0 | S1 | 40410 | Cryptococcus neoformans var. neoformans |
| 393 | 0.00 | 30 | 30 | S2 | 214684 | Cryptococcus neoformans var. neoformans JEC21 |
| 394 | 0.00 | 348 | 0 | D1 | 2611352 | Discoba |
| 395 | 0.00 | 348 | 0 | P | 33682 | Euglenozoa |
| 396 | 0.00 | 348 | 0 | C | 5653 | Kinetoplastea |
| 397 | 0.00 | 348 | 0 | C1 | 2704647 | Metakinetoplastina |
| 398 | 0.00 | 348 | 0 | O | 2704949 | Trypanosomatida |
| 399 | 0.00 | 348 | 0 | F | 5654 | Trypanosomatidae |
| 400 | 0.00 | 348 | 20 | G | 5690 | Trypanosoma |
| 401 | 0.00 | 263 | 0 | G1 | 47570 | Schizotrypanum |
| 402 | 0.00 | 263 | 263 | S | 5693 | Trypanosoma cruzi |
| 403 | 0.00 | 65 | 0 | G1 | 39700 | Trypanozoon |
| 404 | 0.00 | 65 | 0 | S | 5691 | Trypanosoma brucei |
| 405 | 0.00 | 65 | 0 | S1 | 5702 | Trypanosoma brucei brucei |
| 406 | 0.00 | 65 | 65 | S2 | 185431 | Trypanosoma brucei brucei TREU927 |
| 407 | 0.00 | 46 | 0 | D | 2157 | Archaea |
| 408 | 0.00 | 46 | 0 | P | 28890 | Euryarchaeota |
| 409 | 0.00 | 46 | 0 | P1 | 2283794 | Methanomada group |
| 410 | 0.00 | 46 | 0 | C | 183925 | Methanobacteria |
| 411 | 0.00 | 46 | 0 | O | 2158 | Methanobacteriales |
| 412 | 0.00 | 46 | 0 | F | 2159 | Methanobacteriaceae |
| 413 | 0.00 | 46 | 0 | G | 2172 | Methanobrevibacter |
| 414 | 0.00 | 46 | 46 | S | 66851 | Methanobrevibacter oralis |
| 415 | 0.00 | 3227 | 0 | D | 10239 | Viruses |
| 416 | 0.00 | 3226 | 0 | D1 | 2731341 | Duplodnaviria |
| 417 | 0.00 | 3226 | 0 | K | 2731360 | Heunggongvirae |
| 418 | 0.00 | 3226 | 0 | P | 2731361 | Peploviricota |
| 419 | 0.00 | 3226 | 0 | C | 2731363 | Herviviricetes |
| 420 | 0.00 | 3226 | 0 | O | 548681 | Herpesvirales |
| 421 | 0.00 | 3226 | 0 | F | 10292 | Herpesviridae |
| 422 | 0.00 | 3226 | 0 | F1 | 10293 | Alphaherpesvirinae |
| 423 | 0.00 | 3226 | 8 | G | 10319 | Varicellovirus |
| 424 | 0.00 | 3216 | 0 | S | 3050278 | Varicellovirus equidalpha4 |
| 425 | 0.00 | 3216 | 3216 | S1 | 10331 | Equid alphaherpesvirus 4 |
| 426 | 0.00 | 1 | 0 | S | 3050277 | Varicellovirus equidalpha3 |
| 427 | 0.00 | 1 | 1 | S1 | 80341 | Equid alphaherpesvirus 3 |
| 428 | 0.00 | 1 | 0 | S | 3050279 | Varicellovirus equidalpha8 |
| 429 | 0.00 | 1 | 1 | S1 | 39637 | Equid alphaherpesvirus 8 |
| 430 | 0.00 | 1 | 0 | D1 | 2732004 | Varidnaviria |
| 431 | 0.00 | 1 | 0 | K | 2732005 | Bamfordvirae |
| 432 | 0.00 | 1 | 0 | P | 2732007 | Nucleocytoviricota |
| 433 | 0.00 | 1 | 0 | C | 2732525 | Pokkesviricetes |
| 434 | 0.00 | 1 | 0 | O | 2732527 | Chitovirales |
| 435 | 0.00 | 1 | 0 | F | 10240 | Poxviridae |
| 436 | 0.00 | 1 | 0 | F1 | 10241 | Chordopoxvirinae |
| 437 | 0.00 | 1 | 0 | G | 10242 | Orthopoxvirus |
| 438 | 0.00 | 1 | 1 | S | 10243 | Cowpox virus |

**Table S3c:** Results of the Kraken2 analysis for library ERR6466110 based on a custom-built pathogen list (Table S1)

| Sr | Percentage of fragments covered | Number of fragments covered | Number of fragments assigned | rank code | NCBI taxonomic ID | Name |
|---|---|---|---|---|---|---|
| 1 | 99.74 | 163983957 | 163983957 | U | 0 | unclassified |
| 2 | 0.26 | 431490 | 0 | R | 1 | root |
| 3 | 0.26 | 427420 | 2825 | R1 | 131567 | cellular organisms |
| 4 | 0.24 | 392611 | 11194 | D | 2 | Bacteria |
| 5 | 0.18 | 293810 | 7924 | P | 1224 | Pseudomonadota |
| 6 | 0.13 | 212839 | 2074 | C | 28216 | Betaproteobacteria |
| 7 | 0.12 | 205289 | 7285 | O | 80840 | Burkholderiales |
| 8 | 0.08 | 128179 | 0 | F | 119060 | Burkholderiaceae |
| 9 | 0.08 | 128179 | 18275 | G | 32008 | Burkholderia |
| 10 | 0.04 | 61148 | 0 | G1 | 87882 | Burkholderia cepacia complex |
| 11 | 0.04 | 61148 | 61148 | S | 292 | Burkholderia cepacia |
| 12 | 0.03 | 48756 | 41151 | G1 | 111527 | pseudomallei group |
| 13 | 0.00 | 5753 | 5753 | S | 28450 | Burkholderia pseudomallei |
| 14 | 0.00 | 1852 | 1852 | S | 13373 | Burkholderia mallei |
| 15 | 0.04 | 69825 | 575 | F | 506 | Alcaligenaceae |
| 16 | 0.04 | 68255 | 6767 | G | 517 | Bordetella |
| 17 | 0.02 | 32098 | 32098 | S | 520 | Bordetella pertussis |
| 18 | 0.02 | 29390 | 29390 | S | 94624 | Bordetella petrii |
| 19 | 0.00 | 995 | 0 | G | 29574 | Taylorella |
| 20 | 0.00 | 995 | 995 | S | 29575 | Taylorella equigenitalis |
| 21 | 0.00 | 5476 | 0 | O | 206351 | Neisseriales |
| 22 | 0.00 | 5476 | 0 | F | 481 | Neisseriaceae |
| 23 | 0.00 | 5476 | 2406 | G | 482 | Neisseria |
| 24 | 0.00 | 1422 | 1422 | S | 485 | Neisseria gonorrhoeae |
| 25 | 0.00 | 915 | 915 | S | 487 | Neisseria meningitidis |
| 26 | 0.00 | 733 | 733 | S | 483 | Neisseria cinerea |
| 27 | 0.04 | 57637 | 4671 | C | 1236 | Gammaproteobacteria |
| 28 | 0.02 | 37572 | 1248 | O | 91347 | Enterobacterales |
| 29 | 0.02 | 34038 | 7547 | F | 543 | Enterobacteriaceae |
| 30 | 0.01 | 19006 | 0 | F1 | 2890311 | Klebsiella/Raoultella group |
| 31 | 0.01 | 19006 | 1485 | G | 570 | Klebsiella |
| 32 | 0.01 | 10142 | 9505 | S | 573 | Klebsiella pneumoniae |
| 33 | 0.00 | 637 | 637 | S1 | 39831 | Klebsiella pneumoniae subsp. rhinoscleromatis |
| 34 | 0.00 | 7379 | 7379 | S | 571 | Klebsiella oxytoca |
| 35 | 0.00 | 5343 | 0 | G | 590 | Salmonella |
| 36 | 0.00 | 5343 | 0 | S | 28901 | Salmonella enterica |
| 37 | 0.00 | 5343 | 0 | S1 | 59201 | Salmonella enterica subsp. enterica |
| 38 | 0.00 | 5343 | 5343 | S2 | 594 | Salmonella enterica subsp. enterica serovar Gallinarum |
| 39 | 0.00 | 1825 | 0 | G | 561 | Escherichia |
| 40 | 0.00 | 1825 | 1355 | S | 562 | Escherichia coli |
| 41 | 0.00 | 352 | 352 | S1 | 83334 | Escherichia coli O157:H7 |
| 42 | 0.00 | 63 | 63 | S1 | 168927 | Escherichia coli O111:H- |
| 43 | 0.00 | 55 | 55 | S1 | 244319 | Escherichia coli O26:H11 |
| 44 | 0.00 | 317 | 122 | G | 620 | Shigella |
| 45 | 0.00 | 144 | 144 | S | 623 | Shigella flexneri |
| 46 | 0.00 | 29 | 29 | S | 621 | Shigella boydii |
| 47 | 0.00 | 19 | 19 | S | 622 | Shigella dysenteriae |
| 48 | 0.00 | 3 | 3 | S | 624 | Shigella sonnei |
| 49 | 0.00 | 2286 | 0 | F | 1903411 | Yersiniaceae |
| 50 | 0.00 | 2286 | 1052 | G | 629 | Yersinia |
| 51 | 0.00 | 675 | 479 | G1 | 1649845 | Yersinia pseudotuberculosis complex |
| 52 | 0.00 | 174 | 171 | S | 632 | Yersinia pestis |
| 53 | 0.00 | 3 | 3 | S1 | 214092 | Yersinia pestis CO92 |
| 54 | 0.00 | 22 | 22 | S | 633 | Yersinia pseudotuberculosis |
| 55 | 0.00 | 559 | 559 | S | 630 | Yersinia enterocolitica |
| 56 | 0.00 | 7374 | 489 | O | 118969 | Legionellales |
| 57 | 0.00 | 4066 | 0 | F | 118968 | Coxiellaceae |
| 58 | 0.00 | 4066 | 0 | G | 776 | Coxiella |
| 59 | 0.00 | 4066 | 4066 | S | 777 | Coxiella burnetii |
| 60 | 0.00 | 2819 | 0 | F | 444 | Legionellaceae |
| 61 | 0.00 | 2819 | 0 | G | 445 | Legionella |
| 62 | 0.00 | 2819 | 2819 | S | 446 | Legionella pneumophila |
| 63 | 0.00 | 2901 | 0 | O | 2887326 | Moraxellales |
| 64 | 0.00 | 2901 | 0 | F | 468 | Moraxellaceae |
| 65 | 0.00 | 2901 | 0 | G | 475 | Moraxella |
| 66 | 0.00 | 2901 | 2901 | S | 480 | Moraxella catarrhalis |
| 67 | 0.00 | 2398 | 0 | O | 135623 | Vibrionales |
| 68 | 0.00 | 2398 | 0 | F | 641 | Vibrionaceae |
| 69 | 0.00 | 2398 | 775 | G | 662 | Vibrio |
| 70 | 0.00 | 677 | 677 | S | 666 | Vibrio cholerae |
| 71 | 0.00 | 552 | 0 | G1 | 717610 | Vibrio harveyi group |
| 72 | 0.00 | 552 | 552 | S | 670 | Vibrio parahaemolyticus |
| 73 | 0.00 | 394 | 394 | S | 672 | Vibrio vulnificus |
| 74 | 0.00 | 1404 | 0 | O | 72273 | Thiotrichales |
| 75 | 0.00 | 1404 | 0 | F | 34064 | Francisellaceae |
| 76 | 0.00 | 1404 | 0 | G | 262 | Francisella |
| 77 | 0.00 | 1404 | 1404 | S | 263 | Francisella tularensis |
| 78 | 0.00 | 1317 | 0 | O | 135625 | Pasteurellales |
| 79 | 0.00 | 1317 | 384 | F | 712 | Pasteurellaceae |
| 80 | 0.00 | 496 | 0 | G | 724 | Haemophilus |
| 81 | 0.00 | 496 | 0 | S | 727 | Haemophilus influenzae |
| 82 | 0.00 | 496 | 496 | S1 | 725 | Haemophilus influenzae biotype aegyptius |
| 83 | 0.00 | 437 | 0 | G | 713 | Actinobacillus |
| 84 | 0.00 | 437 | 0 | S | 718 | Actinobacillus equuli |
| 85 | 0.00 | 437 | 437 | S1 | 202947 | Actinobacillus equuli subsp. equuli |
| 86 | 0.01 | 15410 | 138 | C | 28211 | Alphaproteobacteria |
| 87 | 0.01 | 14177 | 2107 | O | 356 | Hyphomicrobiales |
| 88 | 0.01 | 9942 | 0 | F | 118882 | Brucellaceae |
| 89 | 0.01 | 9942 | 0 | F1 | 2826938 | Brucella/Ochrobactrum group |
| 90 | 0.01 | 9942 | 9656 | G | 234 | Brucella |
| 91 | 0.00 | 122 | 0 | S | 236 | Brucella ovis |
| 92 | 0.00 | 122 | 122 | S1 | 444178 | Brucella ovis ATCC 25840 |
| 93 | 0.00 | 97 | 0 | S | 444163 | Brucella microti |
| 94 | 0.00 | 97 | 97 | S1 | 568815 | Brucella microti CCM 4915 |
| 95 | 0.00 | 36 | 0 | S | 29459 | Brucella melitensis |
| 96 | 0.00 | 36 | 0 | S1 | 644337 | Brucella melitensis bv. 1 |
| 97 | 0.00 | 36 | 36 | S2 | 224914 | Brucella melitensis bv. 1 str. 16M |
| 98 | 0.00 | 20 | 0 | S | 235 | Brucella abortus |
| 99 | 0.00 | 20 | 20 | S1 | 359391 | Brucella abortus 2308 |
| 100 | 0.00 | 11 | 0 | S | 29461 | Brucella suis |
| 101 | 0.00 | 11 | 11 | S1 | 204722 | Brucella suis 1330 |
| 102 | 0.00 | 2128 | 0 | F | 772 | Bartonellaceae |
| 103 | 0.00 | 2128 | 1311 | G | 773 | Bartonella |
| 104 | 0.00 | 351 | 351 | S | 38323 | Bartonella henselae |
| 105 | 0.00 | 260 | 0 | S | 774 | Bartonella bacilliformis |
| 106 | 0.00 | 260 | 260 | S1 | 360095 | Bartonella bacilliformis KC583 |
| 107 | 0.00 | 206 | 206 | S | 803 | Bartonella quintana |
| 108 | 0.00 | 1095 | 0 | O | 766 | Rickettsiales |
| 109 | 0.00 | 1095 | 0 | F | 775 | Rickettsiaceae |
| 110 | 0.00 | 1095 | 0 | F1 | 33988 | Rickettsieae |
| 111 | 0.00 | 1095 | 736 | G | 780 | Rickettsia |
| 112 | 0.00 | 306 | 173 | G1 | 114277 | spotted fever group |
| 113 | 0.00 | 66 | 66 | S | 42862 | Rickettsia felis |

| | | | | | | |
|---|---|---|---|---|---|---|
| 114 | 0.00 | 43 | 0 | S | 786 | Rickettsia akari |
| 115 | 0.00 | 43 | 43 | S1 | 293614 | Rickettsia akari str. Hartford |
| 116 | 0.00 | 13 | 13 | S | 35790 | Rickettsia japonica |
| 117 | 0.00 | 11 | 0 | G2 | 266068 | Rickettsia sibirica subgroup |
| 118 | 0.00 | 11 | 0 | S | 35793 | Rickettsia sibirica |
| 119 | 0.00 | 11 | 11 | S1 | 272951 | Rickettsia sibirica 246 |
| 120 | 0.00 | 53 | 3 | G1 | 114292 | typhus group |
| 121 | 0.00 | 33 | 33 | S | 785 | Rickettsia typhi |
| 122 | 0.00 | 17 | 17 | S | 782 | Rickettsia prowazekii |
| 123 | 0.05 | 79460 | 432 | D1 | 1783272 | Terrabacteria group |
| 124 | 0.04 | 67626 | 0 | P | 201174 | Actinomycetota |
| 125 | 0.04 | 67626 | 0 | C | 1760 | Actinomycetes |
| 126 | 0.04 | 67626 | 5948 | O | 85007 | Mycobacteriales |
| 127 | 0.03 | 45055 | 1766 | F | 1762 | Mycobacteriaceae |
| 128 | 0.02 | 38565 | 4263 | G | 1763 | Mycobacterium |
| 129 | 0.01 | 23262 | 4729 | G1 | 120793 | Mycobacterium avium complex (MAC) |
| 130 | 0.00 | 7732 | 7732 | S | 1764 | Mycobacterium avium |
| 131 | 0.00 | 5664 | 5664 | S | 1767 | Mycobacterium intracellulare |
| 132 | 0.00 | 5137 | 5137 | S | 339268 | Mycobacterium colombiense |
| 133 | 0.00 | 6046 | 6046 | S | 1768 | Mycobacterium kansasii |
| 134 | 0.00 | 4211 | 3508 | G1 | 77643 | Mycobacterium tuberculosis complex |
| 135 | 0.00 | 366 | 361 | S | 1773 | Mycobacterium tuberculosis |
| 136 | 0.00 | 5 | 5 | S1 | 1765 | Mycobacterium tuberculosis variant bovis |
| 137 | 0.00 | 337 | 337 | S | 78331 | Mycobacterium canettii |
| 138 | 0.00 | 783 | 783 | S | 1769 | Mycobacterium leprae |
| 139 | 0.00 | 4724 | 0 | G | 670516 | Mycobacteroides |
| 140 | 0.00 | 4724 | 0 | S | 36809 | Mycobacteroides abscessus |
| 141 | 0.00 | 4724 | 4724 | S1 | 319705 | Mycobacteroides abscessus subsp. bolletii |
| 142 | 0.01 | 13824 | 0 | F | 85025 | Nocardiaceae |
| 143 | 0.01 | 13824 | 0 | G | 2979332 | Prescottella |
| 144 | 0.01 | 13824 | 13824 | S | 43767 | Prescottella equi |
| 145 | 0.00 | 2799 | 0 | F | 1653 | Corynebacteriaceae |
| 146 | 0.00 | 2799 | 600 | G | 1716 | Corynebacterium |
| 147 | 0.00 | 1747 | 1747 | S | 1717 | Corynebacterium diphtheriae |
| 148 | 0.00 | 452 | 0 | S | 1719 | Corynebacterium pseudotuberculosis |
| 149 | 0.00 | 452 | 452 | S1 | 1087451 | Corynebacterium pseudotuberculosis 31 |
| 150 | 0.01 | 11263 | 943 | P | 1239 | Bacillota |
| 151 | 0.00 | 6406 | 436 | C | 91061 | Bacilli |
| 152 | 0.00 | 4717 | 373 | O | 1385 | Bacillales |
| 153 | 0.00 | 3317 | 0 | F | 186817 | Bacillaceae |
| 154 | 0.00 | 3317 | 0 | G | 1386 | Bacillus |
| 155 | 0.00 | 3317 | 1931 | G1 | 86661 | Bacillus cereus group |
| 156 | 0.00 | 880 | 880 | S | 1396 | Bacillus cereus |
| 157 | 0.00 | 506 | 0 | S | 1392 | Bacillus anthracis |
| 158 | 0.00 | 506 | 506 | S1 | 261594 | Bacillus anthracis str. 'Ames Ancestor' |
| 159 | 0.00 | 1027 | 0 | F | 90964 | Staphylococcaceae |
| 160 | 0.00 | 1027 | 0 | G | 1279 | Staphylococcus |
| 161 | 0.00 | 1027 | 1027 | S | 1280 | Staphylococcus aureus |
| 162 | 0.00 | 1253 | 0 | O | 186826 | Lactobacillales |
| 163 | 0.00 | 1253 | 0 | F | 1300 | Streptococcaceae |
| 164 | 0.00 | 1253 | 634 | G | 1301 | Streptococcus |
| 165 | 0.00 | 134 | 0 | G1 | 119603 | Streptococcus dysgalactiae group |
| 166 | 0.00 | 110 | 0 | S | 1336 | Streptococcus equi |
| 167 | 0.00 | 110 | 110 | S1 | 40041 | Streptococcus equi subsp. zooepidemicus |
| 168 | 0.00 | 24 | 24 | S | 1334 | Streptococcus dysgalactiae |
| 169 | 0.00 | 103 | 103 | S | 1309 | Streptococcus mutans |
| 170 | 0.00 | 98 | 89 | S | 1302 | Streptococcus gordonii |
| 171 | 0.00 | 9 | 0 | S1 | 29390 | Streptococcus gordonii str. Challis |
| 172 | 0.00 | 9 | 9 | S2 | 467705 | Streptococcus gordonii str. Challis substr. CH1 |
| 173 | 0.00 | 60 | 60 | S | 1304 | Streptococcus salivarius |
| 174 | 0.00 | 56 | 0 | G1 | 671232 | Streptococcus anginosus group |
| 175 | 0.00 | 56 | 56 | S | 1328 | Streptococcus anginosus |
| 176 | 0.00 | 49 | 49 | S | 1313 | Streptococcus pneumoniae |
| 177 | 0.00 | 46 | 46 | S | 1305 | Streptococcus sanguinis |
| 178 | 0.00 | 37 | 37 | S | 1311 | Streptococcus agalactiae |
| 179 | 0.00 | 25 | 25 | S | 1314 | Streptococcus pyogenes |
| 180 | 0.00 | 11 | 11 | S | 1303 | Streptococcus oralis |
| 181 | 0.00 | 2868 | 0 | C | 186801 | Clostridia |
| 182 | 0.00 | 2868 | 83 | O | 186802 | Eubacteriales |
| 183 | 0.00 | 1821 | 0 | F | 31979 | Clostridiaceae |
| 184 | 0.00 | 1821 | 780 | G | 1485 | Clostridium |
| 185 | 0.00 | 352 | 352 | S | 1502 | Clostridium perfringens |
| 186 | 0.00 | 259 | 1 | S | 1491 | Clostridium botulinum |
| 187 | 0.00 | 258 | 0 | S1 | 36830 | Clostridium botulinum E |
| 188 | 0.00 | 258 | 258 | S2 | 508767 | Clostridium botulinum E3 str. Alaska E43 |
| 189 | 0.00 | 206 | 206 | S | 1513 | Clostridium tetani |
| 190 | 0.00 | 173 | 173 | S | 1488 | Clostridium acetobutylicum |
| 191 | 0.00 | 51 | 51 | S | 1509 | Clostridium sporogenes |
| 192 | 0.00 | 964 | 0 | F | 186804 | Peptostreptococcaceae |
| 193 | 0.00 | 964 | 0 | G | 1257 | Peptostreptococcus |
| 194 | 0.00 | 964 | 964 | S | 1261 | Peptostreptococcus anaerobius |
| 195 | 0.00 | 633 | 0 | C | 909932 | Negativicutes |
| 196 | 0.00 | 633 | 0 | O | 1843489 | Veillonellales |
| 197 | 0.00 | 633 | 0 | F | 31977 | Veillonellaceae |
| 198 | 0.00 | 633 | 0 | G | 29465 | Veillonella |
| 199 | 0.00 | 633 | 633 | S | 29466 | Veillonella parvula |
| 200 | 0.00 | 413 | 0 | C | 1737404 | Tissierellia |
| 201 | 0.00 | 413 | 0 | O | 1737405 | Tissierellales |
| 202 | 0.00 | 413 | 0 | F | 1570339 | Peptoniphilaceae |
| 203 | 0.00 | 413 | 0 | G | 543311 | Parvimonas |
| 204 | 0.00 | 413 | 413 | S | 33033 | Parvimonas micra |
| 205 | 0.00 | 139 | 0 | P | 544448 | Mycoplasmatota |
| 206 | 0.00 | 139 | 0 | O | 2790996 | Mycoplasmoidales |
| 207 | 0.00 | 139 | 0 | F | 2895623 | Metamycoplasmataceae |
| 208 | 0.00 | 139 | 0 | G | 2995234 | Mycoplasmoides |
| 209 | 0.00 | 139 | 0 | S | 2104 | Mycoplasmoides pneumoniae |
| 210 | 0.00 | 139 | 139 | S1 | 1441379 | Mycoplasmoides pneumoniae M29 |
| 211 | 0.00 | 4508 | 0 | P | 203691 | Spirochaetota |
| 212 | 0.00 | 4508 | 17 | C | 203692 | Spirochaetia |
| 213 | 0.00 | 2386 | 0 | O | 1643688 | Leptospirales |
| 214 | 0.00 | 2386 | 0 | F | 170 | Leptospiraceae |
| 215 | 0.00 | 2386 | 1115 | G | 171 | Leptospira |
| 216 | 0.00 | 336 | 336 | S | 408139 | Leptospira kmetyi |
| 217 | 0.00 | 239 | 239 | S | 447106 | Leptospira licerasiae |
| 218 | 0.00 | 112 | 112 | S | 409998 | Leptospira wolffii |
| 219 | 0.00 | 88 | 0 | S | 48782 | Leptospira fainei |
| 220 | 0.00 | 88 | 0 | S1 | 293072 | Leptospira fainei serovar Hurstbridge |
| 221 | 0.00 | 88 | 88 | S2 | 1193011 | Leptospira fainei serovar Hurstbridge str. BUT 6 |
| 222 | 0.00 | 76 | 0 | S | 29506 | Leptospira inadai |
| 223 | 0.00 | 76 | 76 | S1 | 293084 | Leptospira inadai serovar Lyme |
| 224 | 0.00 | 63 | 63 | S | 28183 | Leptospira santarosai |
| 225 | 0.00 | 62 | 62 | S | 28182 | Leptospira noguchii |
| 226 | 0.00 | 62 | 62 | S | 28452 | Leptospira alstonii |
| 227 | 0.00 | 51 | 0 | S | 301541 | Leptospira broomii |
| 228 | 0.00 | 51 | 0 | S1 | 1324404 | Leptospira broomii serovar Hurstbridge |
| 229 | 0.00 | 51 | 51 | S2 | 1049789 | Leptospira broomii serovar Hurstbridge str. 5399 |

| | | | | | | |
|---|---|---|---|---|---|---|
| 230 | 0.00 | 47 | 47 | S | 29507 | Leptospira kirschneri |
| 231 | 0.00 | 38 | 38 | S | 174 | Leptospira borgpetersenii |
| 232 | 0.00 | 38 | 38 | S | 28184 | Leptospira weilii |
| 233 | 0.00 | 32 | 32 | S | 173 | Leptospira interrogans |
| 234 | 0.00 | 27 | 27 | S | 100053 | Leptospira alexanderi |
| 235 | 0.00 | 2105 | 79 | O | 136 | Spirochaetales |
| 236 | 0.00 | 1628 | 0 | F | 2845253 | Treponemataceae |
| 237 | 0.00 | 1628 | 194 | G | 157 | Treponema |
| 238 | 0.00 | 1040 | 1040 | S | 156 | Treponema zuelzerae |
| 239 | 0.00 | 238 | 232 | S | 160 | Treponema pallidum |
| 240 | 0.00 | 6 | 6 | S1 | 168 | Treponema pallidum subsp. pertenue |
| 241 | 0.00 | 156 | 152 | S | 158 | Treponema denticola |
| 242 | 0.00 | 4 | 4 | S1 | 243275 | Treponema denticola ATCC 35405 |
| 243 | 0.00 | 398 | 113 | F | 1643685 | Borreliaceae |
| 244 | 0.00 | 175 | 167 | G | 64895 | Borreliella |
| 245 | 0.00 | 7 | 0 | S | 139 | Borreliella burgdorferi |
| 246 | 0.00 | 7 | 7 | S1 | 445984 | Borreliella burgdorferi Bol26 |
| 247 | 0.00 | 1 | 0 | S | 664662 | Borreliella bavariensis |
| 248 | 0.00 | 1 | 1 | S1 | 290434 | Borrelia garinii subsp. bavariensis PBi |
| 249 | 0.00 | 110 | 0 | G | 138 | Borrelia |
| 250 | 0.00 | 110 | 0 | S | 44449 | Borrelia recurrentis |
| 251 | 0.00 | 110 | 110 | S1 | 412418 | Borrelia recurrentis A1 |
| 252 | 0.00 | 1517 | 0 | D1 | 1783270 | FCB group |
| 253 | 0.00 | 1517 | 0 | D2 | 68336 | Bacteroidota/Chlorobiota group |
| 254 | 0.00 | 1517 | 0 | P | 976 | Bacteroidota |
| 255 | 0.00 | 1517 | 0 | C | 200643 | Bacteroidia |
| 256 | 0.00 | 1517 | 560 | O | 171549 | Bacteroidales |
| 257 | 0.00 | 561 | 0 | F | 2005525 | Tannerellaceae |
| 258 | 0.00 | 561 | 0 | G | 195950 | Tannerella |
| 259 | 0.00 | 561 | 546 | S | 28112 | Tannerella forsythia |
| 260 | 0.00 | 15 | 15 | S1 | 203275 | Tannerella forsythia 92A2 |
| 261 | 0.00 | 396 | 0 | F | 171551 | Porphyromonadaceae |
| 262 | 0.00 | 396 | 0 | G | 836 | Porphyromonas |
| 263 | 0.00 | 396 | 347 | S | 837 | Porphyromonas gingivalis |
| 264 | 0.00 | 49 | 49 | S1 | 242619 | Porphyromonas gingivalis W83 |
| 265 | 0.00 | 821 | 0 | P | 32066 | Fusobacteriota |
| 266 | 0.00 | 821 | 0 | C | 203490 | Fusobacteriia |
| 267 | 0.00 | 821 | 0 | O | 203491 | Fusobacteriales |
| 268 | 0.00 | 821 | 0 | F | 203492 | Fusobacteriaceae |
| 269 | 0.00 | 821 | 249 | G | 848 | Fusobacterium |
| 270 | 0.00 | 359 | 359 | S | 851 | Fusobacterium nucleatum |
| 271 | 0.00 | 213 | 213 | S | 849 | Fusobacterium gonidiaformans |
| 272 | 0.00 | 764 | 0 | D1 | 1783257 | PVC group |
| 273 | 0.00 | 764 | 0 | P | 204428 | Chlamydiota |
| 274 | 0.00 | 764 | 0 | C | 204429 | Chlamydiia |
| 275 | 0.00 | 764 | 0 | O | 51291 | Chlamydiales |
| 276 | 0.00 | 764 | 0 | F | 809 | Chlamydiaceae |
| 277 | 0.00 | 764 | 0 | F1 | 1113537 | Chlamydia/Chlamydophila group |
| 278 | 0.00 | 764 | 685 | G | 810 | Chlamydia |
| 279 | 0.00 | 24 | 0 | S | 83558 | Chlamydia pneumoniae |
| 280 | 0.00 | 24 | 24 | S1 | 406984 | Chlamydia pneumoniae LPCoLN |
| 281 | 0.00 | 20 | 0 | S | 83557 | Chlamydia caviae |
| 282 | 0.00 | 20 | 20 | S1 | 227941 | Chlamydia caviae GPIC |
| 283 | 0.00 | 16 | 0 | S | 83556 | Chlamydia felis |
| 284 | 0.00 | 16 | 16 | S1 | 264202 | Chlamydia felis Fe/C-56 |
| 285 | 0.00 | 9 | 9 | S | 85991 | Chlamydia pecorum |
| 286 | 0.00 | 6 | 6 | S | 83560 | Chlamydia muridarum |
| 287 | 0.00 | 2 | 0 | S | 813 | Chlamydia trachomatis |
| 288 | 0.00 | 2 | 2 | S1 | 272561 | Chlamydia trachomatis D/UW-3/CX |
| 289 | 0.00 | 2 | 0 | S | 83554 | Chlamydia psittaci |
| 290 | 0.00 | 2 | 2 | S1 | 331636 | Chlamydia psittaci 6BC |
| 291 | 0.00 | 537 | 0 | P | 29547 | Campylobacterota |
| 292 | 0.00 | 537 | 0 | C | 3031852 | Epsilonproteobacteria |
| 293 | 0.00 | 537 | 0 | O | 213849 | Campylobacterales |
| 294 | 0.00 | 537 | 0 | F | 72293 | Helicobacteraceae |
| 295 | 0.00 | 537 | 0 | G | 209 | Helicobacter |
| 296 | 0.00 | 537 | 537 | S | 210 | Helicobacter pylori |
| 297 | 0.02 | 31935 | 2509 | D | 2759 | Eukaryota |
| 298 | 0.01 | 15238 | 0 | D1 | 2698737 | Sar |
| 299 | 0.01 | 15238 | 0 | D2 | 33630 | Alveolata |
| 300 | 0.01 | 15238 | 74 | P | 5794 | Apicomplexa |
| 301 | 0.01 | 9423 | 0 | C | 422676 | Aconoidasida |
| 302 | 0.01 | 9423 | 0 | O | 5819 | Haemosporida |
| 303 | 0.01 | 9423 | 0 | F | 1639119 | Plasmodiidae |
| 304 | 0.01 | 9423 | 43 | G | 5820 | Plasmodium |
| 305 | 0.01 | 9354 | 0 | G1 | 418103 | Plasmodium (Plasmodium) |
| 306 | 0.01 | 9354 | 9354 | S | 5855 | Plasmodium vivax |
| 307 | 0.00 | 26 | 0 | G1 | 418107 | Plasmodium (Laverania) |
| 308 | 0.00 | 26 | 0 | S | 5833 | Plasmodium falciparum |
| 309 | 0.00 | 26 | 26 | S1 | 36329 | Plasmodium falciparum 3D7 |
| 310 | 0.00 | 5741 | 0 | C | 1280412 | Conoidasida |
| 311 | 0.00 | 5741 | 0 | C1 | 5796 | Coccidia |
| 312 | 0.00 | 5741 | 0 | O | 75739 | Eucoccidiorida |
| 313 | 0.00 | 5741 | 0 | O1 | 423054 | Eimeriorina |
| 314 | 0.00 | 5741 | 0 | F | 5809 | Sarcocystidae |
| 315 | 0.00 | 5741 | 0 | G | 5810 | Toxoplasma |
| 316 | 0.00 | 5741 | 0 | S | 5811 | Toxoplasma gondii |
| 317 | 0.00 | 5734 | 5734 | S1 | 508771 | Toxoplasma gondii ME49 |
| 318 | 0.00 | 7 | 7 | S1 | 383379 | Toxoplasma gondii RH |
| 319 | 0.01 | 13693 | 278 | D1 | 33154 | Opisthokonta |
| 320 | 0.01 | 12111 | 0 | K | 33208 | Metazoa |
| 321 | 0.01 | 12111 | 0 | K1 | 6072 | Eumetazoa |
| 322 | 0.01 | 12111 | 0 | K2 | 33213 | Bilateria |
| 323 | 0.01 | 12111 | 103 | K3 | 33317 | Protostomia |
| 324 | 0.01 | 11495 | 0 | K4 | 2697495 | Spiralia |
| 325 | 0.01 | 11495 | 0 | K5 | 1206795 | Lophotrochozoa |
| 326 | 0.01 | 11495 | 186 | P | 6157 | Platyhelminthes |
| 327 | 0.01 | 9412 | 0 | C | 6199 | Cestoda |
| 328 | 0.01 | 9412 | 0 | C1 | 6200 | Eucestoda |
| 329 | 0.01 | 9412 | 0 | O | 6201 | Cyclophyllidea |
| 330 | 0.01 | 9412 | 0 | F | 6208 | Taeniidae |
| 331 | 0.01 | 9412 | 0 | G | 6202 | Taenia |
| 332 | 0.01 | 9412 | 9412 | S | 6204 | Taenia solium |
| 333 | 0.00 | 1897 | 0 | C | 6178 | Trematoda |
| 334 | 0.00 | 1897 | 0 | C1 | 6179 | Digenea |
| 335 | 0.00 | 1897 | 0 | O | 6180 | Strigeidida |
| 336 | 0.00 | 1897 | 0 | O1 | 31244 | Schistosomatoidea |
| 337 | 0.00 | 1897 | 0 | F | 31245 | Schistosomatidae |
| 338 | 0.00 | 1897 | 0 | G | 6181 | Schistosoma |
| 339 | 0.00 | 1897 | 1897 | S | 6183 | Schistosoma mansoni |
| 340 | 0.00 | 513 | 0 | K4 | 1206794 | Ecdysozoa |
| 341 | 0.00 | 513 | 0 | P | 6231 | Nematoda |
| 342 | 0.00 | 513 | 0 | C | 119088 | Enoplea |
| 343 | 0.00 | 513 | 0 | C1 | 1457286 | Dorylaimia |
| 344 | 0.00 | 513 | 0 | O | 6329 | Trichinellida |
| 345 | 0.00 | 513 | 0 | F | 6332 | Trichinellidae |

| | | | | | | |
|---|---|---|---|---|---|---|
| 346 | 0.00 | 513 | 0 | G | 6333 | Trichinella |
| 347 | 0.00 | 513 | 513 | S | 6334 | Trichinella spiralis |
| 348 | 0.00 | 1304 | 0 | K | 4751 | Fungi |
| 349 | 0.00 | 1304 | 74 | K1 | 451864 | Dikarya |
| 350 | 0.00 | 902 | 2 | P | 4890 | Ascomycota |
| 351 | 0.00 | 818 | 0 | P1 | 716545 | saccharomyceta |
| 352 | 0.00 | 818 | 0 | P2 | 147538 | Pezizomycotina |
| 353 | 0.00 | 818 | 0 | P3 | 716546 | leotiomyceta |
| 354 | 0.00 | 818 | 0 | C | 147545 | Eurotiomycetes |
| 355 | 0.00 | 818 | 35 | C1 | 451871 | Eurotiomycetidae |
| 356 | 0.00 | 597 | 11 | O | 33183 | Onygenales |
| 357 | 0.00 | 471 | 30 | F | 299071 | Ajellomycetaceae |
| 358 | 0.00 | 296 | 0 | G | 229219 | Blastomyces |
| 359 | 0.00 | 296 | 0 | S | 5039 | Blastomyces dermatitidis |
| 360 | 0.00 | 296 | 296 | S1 | 559297 | Blastomyces dermatitidis ER-3 |
| 361 | 0.00 | 145 | 0 | G | 5036 | Histoplasma |
| 362 | 0.00 | 145 | 0 | S | 5037 | Histoplasma capsulatum |
| 363 | 0.00 | 145 | 145 | S1 | 447093 | Histoplasma capsulatum G186AR |
| 364 | 0.00 | 115 | 0 | F | 33184 | Onygenaceae |
| 365 | 0.00 | 115 | 34 | G | 5500 | Coccidioides |
| 366 | 0.00 | 44 | 0 | S | 199306 | Coccidioides posadasii |
| 367 | 0.00 | 44 | 44 | S1 | 222929 | Coccidioides posadasii C735 delta SOWgp |
| 368 | 0.00 | 37 | 0 | S | 5501 | Coccidioides immitis |
| 369 | 0.00 | 37 | 37 | S1 | 246410 | Coccidioides immitis RS |
| 370 | 0.00 | 186 | 0 | O | 5042 | Eurotiales |
| 371 | 0.00 | 186 | 0 | F | 1131492 | Aspergillaceae |
| 372 | 0.00 | 186 | 0 | G | 5052 | Aspergillus |
| 373 | 0.00 | 186 | 0 | G1 | 2720872 | Aspergillus subgen. Fumigati |
| 374 | 0.00 | 186 | 0 | S | 746128 | Aspergillus fumigatus |
| 375 | 0.00 | 186 | 186 | S1 | 330879 | Aspergillus fumigatus Af293 |
| 376 | 0.00 | 82 | 0 | P1 | 451866 | Taphrinomycotina |
| 377 | 0.00 | 82 | 0 | C | 147553 | Pneumocystidomycetes |
| 378 | 0.00 | 82 | 0 | O | 37987 | Pneumocystidales |
| 379 | 0.00 | 82 | 0 | F | 44281 | Pneumocystidaceae |
| 380 | 0.00 | 82 | 13 | G | 4753 | Pneumocystis |
| 381 | 0.00 | 49 | 0 | S | 4754 | Pneumocystis carinii |
| 382 | 0.00 | 49 | 49 | S1 | 1408658 | Pneumocystis carinii B80 |
| 383 | 0.00 | 20 | 0 | S | 42068 | Pneumocystis jirovecii |
| 384 | 0.00 | 20 | 20 | S1 | 1408657 | Pneumocystis jirovecii RU7 |
| 385 | 0.00 | 328 | 0 | P | 5204 | Basidiomycota |
| 386 | 0.00 | 328 | 0 | P1 | 5302 | Agaricomycotina |
| 387 | 0.00 | 328 | 0 | C | 155616 | Tremellomycetes |
| 388 | 0.00 | 328 | 0 | O | 5234 | Tremellales |
| 389 | 0.00 | 328 | 0 | F | 1884633 | Cryptococcaceae |
| 390 | 0.00 | 328 | 192 | G | 5206 | Cryptococcus |
| 391 | 0.00 | 95 | 0 | G1 | 1884637 | Cryptococcus gattii species complex |
| 392 | 0.00 | 95 | 0 | S | 37769 | Cryptococcus gattii VGI |
| 393 | 0.00 | 95 | 95 | S1 | 367775 | Cryptococcus gattii WM276 |
| 394 | 0.00 | 41 | 0 | G1 | 1897064 | Cryptococcus neoformans species complex |
| 395 | 0.00 | 41 | 0 | S | 5207 | Cryptococcus neoformans |
| 396 | 0.00 | 41 | 0 | S1 | 40410 | Cryptococcus neoformans var. neoformans |
| 397 | 0.00 | 41 | 41 | S2 | 214684 | Cryptococcus neoformans var. neoformans JEC21 |
| 398 | 0.00 | 495 | 0 | D1 | 2611352 | Discoba |
| 399 | 0.00 | 495 | 0 | P | 33682 | Euglenozoa |
| 400 | 0.00 | 495 | 0 | C | 5653 | Kinetoplastea |
| 401 | 0.00 | 495 | 0 | C1 | 2704647 | Metakinetoplastina |
| 402 | 0.00 | 495 | 0 | O | 2704949 | Trypanosomatida |
| 403 | 0.00 | 495 | 0 | F | 5654 | Trypanosomatidae |
| 404 | 0.00 | 495 | 30 | G | 5690 | Trypanosoma |
| 405 | 0.00 | 351 | 0 | G1 | 47570 | Schizotrypanum |
| 406 | 0.00 | 351 | 351 | S | 5693 | Trypanosoma cruzi |
| 407 | 0.00 | 114 | 0 | G1 | 39700 | Trypanozoon |
| 408 | 0.00 | 114 | 0 | S | 5691 | Trypanosoma brucei |
| 409 | 0.00 | 114 | 0 | S1 | 5702 | Trypanosoma brucei brucei |
| 410 | 0.00 | 114 | 114 | S2 | 185431 | Trypanosoma brucei brucei TREU927 |
| 411 | 0.00 | 49 | 0 | D | 2157 | Archaea |
| 412 | 0.00 | 49 | 0 | P | 28890 | Euryarchaeota |
| 413 | 0.00 | 49 | 0 | P1 | 2283794 | Methanomada group |
| 414 | 0.00 | 49 | 0 | C | 183925 | Methanobacteria |
| 415 | 0.00 | 49 | 0 | O | 2158 | Methanobacteriales |
| 416 | 0.00 | 49 | 0 | F | 2159 | Methanobacteriaceae |
| 417 | 0.00 | 49 | 0 | G | 2172 | Methanobrevibacter |
| 418 | 0.00 | 49 | 49 | S | 66851 | Methanobrevibacter oralis |
| 419 | 0.00 | 4070 | 0 | D | 10239 | Viruses |
| 420 | 0.00 | 4062 | 0 | D1 | 2731341 | Duplodnaviria |
| 421 | 0.00 | 4062 | 0 | K | 2731360 | Heunggongvirae |
| 422 | 0.00 | 4062 | 0 | P | 2731361 | Peploviricota |
| 423 | 0.00 | 4062 | 0 | C | 2731363 | Herviviricetes |
| 424 | 0.00 | 4062 | 0 | O | 548681 | Herpesvirales |
| 425 | 0.00 | 4062 | 0 | F | 10292 | Herpesviridae |
| 426 | 0.00 | 4062 | 0 | F1 | 10293 | Alphaherpesvirinae |
| 427 | 0.00 | 4062 | 13 | G | 10319 | Varicellovirus |
| 428 | 0.00 | 4043 | 0 | S | 3050278 | Varicellovirus equidalpha4 |
| 429 | 0.00 | 4043 | 4043 | S1 | 10331 | Equid alphaherpesvirus 4 |
| 430 | 0.00 | 6 | 0 | S | 3050277 | Varicellovirus equidalpha3 |
| 431 | 0.00 | 6 | 6 | S1 | 80341 | Equid alphaherpesvirus 3 |
| 432 | 0.00 | 8 | 0 | D1 | 2732004 | Varidnaviria |
| 433 | 0.00 | 8 | 0 | K | 2732005 | Bamfordvirae |
| 434 | 0.00 | 6 | 0 | P | 2732007 | Nucleocytoviricota |
| 435 | 0.00 | 6 | 0 | C | 2732523 | Megaviricetes |
| 436 | 0.00 | 6 | 0 | O | 2732555 | Pimascovirales |
| 437 | 0.00 | 6 | 0 | F | 944644 | Marseilleviridae |
| 438 | 0.00 | 6 | 0 | G | 1513458 | Marseillevirus |
| 439 | 0.00 | 6 | 0 | G1 | 1813598 | unclassified Marseillevirus |
| 440 | 0.00 | 6 | 6 | S | 1826170 | Tokyovirus A1 |
| 441 | 0.00 | 2 | 0 | P | 2732008 | Preplasmiviricota |
| 442 | 0.00 | 2 | 0 | C | 2732529 | Tectiliviricetes |
| 443 | 0.00 | 2 | 0 | O | 2732559 | Rowavirales |
| 444 | 0.00 | 2 | 0 | F | 10508 | Adenoviridae |
| 445 | 0.00 | 2 | 0 | G | 10509 | Mastadenovirus |
| 446 | 0.00 | 2 | 2 | S | 129951 | Human mastadenovirus C |

**Table S3d:** Results of the Kraken2 analysis for library ERR6466111 based on a custom-built pathogen list (Table S1)

| Sr | Percentage of fragments covered | Number of fragments covered | Number of fragments assigned | rank code | NCBI taxonomic ID | Name |
|---|---|---|---|---|---|---|
| 1 | 99.75 | 141063866 | 141063866 | U | 0 | unclassified |
| 2 | 0.25 | 351989 | 1 | R | 1 | root |
| 3 | 0.25 | 348311 | 2227 | R1 | 131567 | cellular organisms |
| 4 | 0.23 | 319693 | 9206 | D | 2 | Bacteria |
| 5 | 0.17 | 238827 | 6303 | P | 1224 | Pseudomonadota |
| 6 | 0.12 | 173718 | 1739 | C | 28216 | Betaproteobacteria |
| 7 | 0.12 | 167608 | 6290 | O | 80840 | Burkholderiales |
| 8 | 0.07 | 104025 | 0 | F | 119060 | Burkholderiaceae |
| 9 | 0.07 | 104025 | 14461 | G | 32008 | Burkholderia |
| 10 | 0.03 | 49410 | 0 | G1 | 87882 | Burkholderia cepacia complex |
| 11 | 0.03 | 49410 | 49410 | S | 292 | Burkholderia cepacia |
| 12 | 0.03 | 40154 | 33565 | G1 | 111527 | pseudomallei group |
| 13 | 0.00 | 4927 | 4927 | S | 28450 | Burkholderia pseudomallei |
| 14 | 0.00 | 1662 | 1662 | S | 13373 | Burkholderia mallei |
| 15 | 0.04 | 57293 | 493 | F | 506 | Alcaligenaceae |
| 16 | 0.04 | 56034 | 5732 | G | 517 | Bordetella |
| 17 | 0.02 | 25961 | 25961 | S | 520 | Bordetella pertussis |
| 18 | 0.02 | 24341 | 24341 | S | 94624 | Bordetella petrii |
| 19 | 0.00 | 766 | 0 | G | 29574 | Taylorella |
| 20 | 0.00 | 766 | 766 | S | 29575 | Taylorella equigenitalis |
| 21 | 0.00 | 4371 | 0 | O | 206351 | Neisseriales |
| 22 | 0.00 | 4371 | 0 | F | 481 | Neisseriaceae |
| 23 | 0.00 | 4371 | 1947 | G | 482 | Neisseria |
| 24 | 0.00 | 1123 | 1123 | S | 485 | Neisseria gonorrhoeae |
| 25 | 0.00 | 675 | 675 | S | 487 | Neisseria meningitidis |
| 26 | 0.00 | 626 | 626 | S | 483 | Neisseria cinerea |
| 27 | 0.03 | 46734 | 3955 | C | 1236 | Gammaproteobacteria |
| 28 | 0.02 | 30129 | 1003 | O | 91347 | Enterobacterales |
| 29 | 0.02 | 27256 | 5955 | F | 543 | Enterobacteriaceae |
| 30 | 0.01 | 15324 | 0 | F1 | 2890311 | Klebsiella/Raoultella group |
| 31 | 0.01 | 15324 | 1187 | G | 570 | Klebsiella |
| 32 | 0.01 | 8313 | 7825 | S | 573 | Klebsiella pneumoniae |
| 33 | 0.00 | 488 | 488 | S1 | 39831 | Klebsiella pneumoniae subsp. rhinoscleromatis |
| 34 | 0.00 | 5824 | 5824 | S | 571 | Klebsiella oxytoca |
| 35 | 0.00 | 4225 | 0 | G | 590 | Salmonella |
| 36 | 0.00 | 4225 | 0 | S | 28901 | Salmonella enterica |
| 37 | 0.00 | 4225 | 0 | S1 | 59201 | Salmonella enterica subsp. enterica |
| 38 | 0.00 | 4225 | 4225 | S2 | 594 | Salmonella enterica subsp. enterica serovar Gallinarum |
| 39 | 0.00 | 1476 | 0 | G | 561 | Escherichia |
| 40 | 0.00 | 1476 | 1044 | S | 562 | Escherichia coli |
| 41 | 0.00 | 298 | 298 | S1 | 83334 | Escherichia coli O157:H7 |
| 42 | 0.00 | 98 | 98 | S1 | 168927 | Escherichia coli O111:H- |
| 43 | 0.00 | 36 | 36 | S1 | 244319 | Escherichia coli O26:H11 |
| 44 | 0.00 | 276 | 119 | G | 620 | Shigella |
| 45 | 0.00 | 116 | 116 | S | 623 | Shigella flexneri |
| 46 | 0.00 | 25 | 25 | S | 622 | Shigella dysenteriae |
| 47 | 0.00 | 15 | 15 | S | 621 | Shigella boydii |
| 48 | 0.00 | 1 | 1 | S | 624 | Shigella sonnei |
| 49 | 0.00 | 1870 | 0 | F | 1903411 | Yersiniaceae |
| 50 | 0.00 | 1870 | 840 | G | 629 | Yersinia |
| 51 | 0.00 | 561 | 394 | G1 | 1649845 | Yersinia pseudotuberculosis complex |
| 52 | 0.00 | 145 | 143 | S | 632 | Yersinia pestis |
| 53 | 0.00 | 2 | 2 | S1 | 214092 | Yersinia pestis CO92 |
| 54 | 0.00 | 22 | 22 | S | 633 | Yersinia pseudotuberculosis |
| 55 | 0.00 | 469 | 469 | S | 630 | Yersinia enterocolitica |
| 56 | 0.00 | 5983 | 363 | O | 118969 | Legionellales |
| 57 | 0.00 | 3255 | 0 | F | 118968 | Coxiellaceae |
| 58 | 0.00 | 3255 | 0 | G | 776 | Coxiella |
| 59 | 0.00 | 3255 | 3255 | S | 777 | Coxiella burnetii |
| 60 | 0.00 | 2365 | 0 | F | 444 | Legionellaceae |
| 61 | 0.00 | 2365 | 0 | G | 445 | Legionella |
| 62 | 0.00 | 2365 | 2365 | S | 446 | Legionella pneumophila |
| 63 | 0.00 | 2407 | 0 | O | 2887326 | Moraxellales |
| 64 | 0.00 | 2407 | 0 | F | 468 | Moraxellaceae |
| 65 | 0.00 | 2407 | 0 | G | 475 | Moraxella |
| 66 | 0.00 | 2407 | 2407 | S | 480 | Moraxella catarrhalis |
| 67 | 0.00 | 1919 | 0 | O | 135623 | Vibrionales |
| 68 | 0.00 | 1919 | 0 | F | 641 | Vibrionaceae |
| 69 | 0.00 | 1919 | 595 | G | 662 | Vibrio |
| 70 | 0.00 | 588 | 588 | S | 666 | Vibrio cholerae |
| 71 | 0.00 | 425 | 0 | G1 | 717610 | Vibrio harveyi group |
| 72 | 0.00 | 425 | 425 | S | 670 | Vibrio parahaemolyticus |
| 73 | 0.00 | 311 | 311 | S | 672 | Vibrio vulnificus |
| 74 | 0.00 | 1292 | 0 | O | 72273 | Thiotrichales |
| 75 | 0.00 | 1292 | 0 | F | 34064 | Francisellaceae |
| 76 | 0.00 | 1292 | 0 | G | 262 | Francisella |
| 77 | 0.00 | 1292 | 1292 | S | 263 | Francisella tularensis |
| 78 | 0.00 | 1049 | 0 | O | 135625 | Pasteurellales |
| 79 | 0.00 | 1049 | 305 | F | 712 | Pasteurellaceae |
| 80 | 0.00 | 373 | 0 | G | 713 | Actinobacillus |
| 81 | 0.00 | 373 | 0 | S | 718 | Actinobacillus equuli |
| 82 | 0.00 | 373 | 373 | S1 | 202947 | Actinobacillus equuli subsp. equuli |
| 83 | 0.00 | 371 | 0 | G | 724 | Haemophilus |
| 84 | 0.00 | 371 | 0 | S | 727 | Haemophilus influenzae |
| 85 | 0.00 | 371 | 371 | S1 | 725 | Haemophilus influenzae biotype aegyptius |
| 86 | 0.01 | 12072 | 100 | C | 28211 | Alphaproteobacteria |
| 87 | 0.01 | 11130 | 1700 | O | 356 | Hyphomicrobiales |
| 88 | 0.01 | 7726 | 0 | F | 118882 | Brucellaceae |
| 89 | 0.01 | 7726 | 0 | F1 | 2826938 | Brucella/Ochrobactrum group |
| 90 | 0.01 | 7726 | 7520 | G | 234 | Brucella |
| 91 | 0.00 | 116 | 0 | S | 236 | Brucella ovis |
| 92 | 0.00 | 116 | 116 | S1 | 444178 | Brucella ovis ATCC 25840 |
| 93 | 0.00 | 55 | 0 | S | 444163 | Brucella microti |
| 94 | 0.00 | 55 | 55 | S1 | 568815 | Brucella microti CCM 4915 |
| 95 | 0.00 | 17 | 0 | S | 29459 | Brucella melitensis |
| 96 | 0.00 | 17 | 0 | S1 | 644337 | Brucella melitensis bv. 1 |
| 97 | 0.00 | 17 | 17 | S2 | 224914 | Brucella melitensis bv. 1 str. 16M |
| 98 | 0.00 | 14 | 0 | S | 235 | Brucella abortus |
| 99 | 0.00 | 14 | 14 | S1 | 359391 | Brucella abortus 2308 |
| 100 | 0.00 | 4 | 0 | S | 29461 | Brucella suis |
| 101 | 0.00 | 4 | 4 | S1 | 204722 | Brucella suis 1330 |
| 102 | 0.00 | 1704 | 0 | F | 772 | Bartonellaceae |
| 103 | 0.00 | 1704 | 1095 | G | 773 | Bartonella |
| 104 | 0.00 | 261 | 261 | S | 38323 | Bartonella henselae |
| 105 | 0.00 | 195 | 0 | S | 774 | Bartonella bacilliformis |
| 106 | 0.00 | 195 | 195 | S1 | 360095 | Bartonella bacilliformis KC583 |
| 107 | 0.00 | 153 | 153 | S | 803 | Bartonella quintana |
| 108 | 0.00 | 842 | 0 | O | 766 | Rickettsiales |
| 109 | 0.00 | 842 | 0 | F | 775 | Rickettsiaceae |
| 110 | 0.00 | 842 | 0 | F1 | 33988 | Rickettsieae |
| 111 | 0.00 | 842 | 543 | G | 780 | Rickettsia |
| 112 | 0.00 | 257 | 161 | G1 | 114277 | spotted fever group |
| 113 | 0.00 | 36 | 0 | S | 786 | Rickettsia akari |
| 114 | 0.00 | 36 | 36 | S1 | 293614 | Rickettsia akari str. Hartford |
| 115 | 0.00 | 35 | 35 | S | 42862 | Rickettsia felis |
| 116 | 0.00 | 25 | 25 | S | 35790 | Rickettsia japonica |

| | | | | | | |
|---|---|---|---|---|---|---|
| 117 | 0.00 | 42 | 10 | G1 | 114292 | typhus group |
| 118 | 0.00 | 20 | 20 | S | 785 | Rickettsia typhi |
| 119 | 0.00 | 12 | 12 | S | 782 | Rickettsia prowazekii |
| 120 | 0.05 | 65000 | 334 | D1 | 1783272 | Terrabacteria group |
| 121 | 0.04 | 55331 | 0 | P | 201174 | Actinomycetota |
| 122 | 0.04 | 55331 | 0 | C | 1760 | Actinomycetes |
| 123 | 0.04 | 55331 | 4549 | O | 85007 | Mycobacteriales |
| 124 | 0.03 | 37231 | 1360 | F | 1762 | Mycobacteriaceae |
| 125 | 0.02 | 32000 | 3322 | G | 1763 | Mycobacterium |
| 126 | 0.01 | 19397 | 3825 | G1 | 120793 | Mycobacterium avium complex (MAC) |
| 127 | 0.00 | 6370 | 6370 | S | 1764 | Mycobacterium avium |
| 128 | 0.00 | 4650 | 4650 | S | 339268 | Mycobacterium colombiense |
| 129 | 0.00 | 4552 | 4552 | S | 1767 | Mycobacterium intracellulare |
| 130 | 0.00 | 5017 | 5017 | S | 1768 | Mycobacterium kansasii |
| 131 | 0.00 | 3547 | 3007 | G1 | 77643 | Mycobacterium tuberculosis complex |
| 132 | 0.00 | 309 | 309 | S | 78331 | Mycobacterium canettii |
| 133 | 0.00 | 231 | 231 | S | 1773 | Mycobacterium tuberculosis |
| 134 | 0.00 | 717 | 717 | S | 1769 | Mycobacterium leprae |
| 135 | 0.00 | 3871 | 0 | G | 670516 | Mycobacteroides |
| 136 | 0.00 | 3871 | 0 | S | 36809 | Mycobacteroides abscessus |
| 137 | 0.00 | 3871 | 3871 | S1 | 319705 | Mycobacteroides abscessus subsp. bolletii |
| 138 | 0.01 | 11243 | 0 | F | 85025 | Nocardiaceae |
| 139 | 0.01 | 11243 | 0 | G | 2979332 | Prescottella |
| 140 | 0.01 | 11243 | 11243 | S | 43767 | Prescottella equi |
| 141 | 0.00 | 2308 | 0 | F | 1653 | Corynebacteriaceae |
| 142 | 0.00 | 2308 | 539 | G | 1716 | Corynebacterium |
| 143 | 0.00 | 1405 | 1405 | S | 1717 | Corynebacterium diphtheriae |
| 144 | 0.00 | 364 | 0 | S | 1719 | Corynebacterium pseudotuberculosis |
| 145 | 0.00 | 364 | 364 | S1 | 1087451 | Corynebacterium pseudotuberculosis 31 |
| 146 | 0.01 | 9224 | 724 | P | 1239 | Bacillota |
| 147 | 0.00 | 5119 | 345 | C | 91061 | Bacilli |
| 148 | 0.00 | 3719 | 283 | O | 1385 | Bacillales |
| 149 | 0.00 | 2575 | 0 | F | 186817 | Bacillaceae |
| 150 | 0.00 | 2575 | 0 | G | 1386 | Bacillus |
| 151 | 0.00 | 2575 | 1556 | G1 | 86661 | Bacillus cereus group |
| 152 | 0.00 | 651 | 651 | S | 1396 | Bacillus cereus |
| 153 | 0.00 | 368 | 0 | S | 1392 | Bacillus anthracis |
| 154 | 0.00 | 368 | 368 | S1 | 261594 | Bacillus anthracis str. 'Ames Ancestor' |
| 155 | 0.00 | 861 | 0 | F | 90964 | Staphylococcaceae |
| 156 | 0.00 | 861 | 0 | G | 1279 | Staphylococcus |
| 157 | 0.00 | 861 | 861 | S | 1280 | Staphylococcus aureus |
| 158 | 0.00 | 1055 | 0 | O | 186826 | Lactobacillales |
| 159 | 0.00 | 1055 | 0 | F | 1300 | Streptococcaceae |
| 160 | 0.00 | 1055 | 622 | G | 1301 | Streptococcus |
| 161 | 0.00 | 81 | 0 | G1 | 119603 | Streptococcus dysgalactiae group |
| 162 | 0.00 | 62 | 0 | S | 1336 | Streptococcus equi |
| 163 | 0.00 | 62 | 62 | S1 | 40041 | Streptococcus equi subsp. zooepidemicus |
| 164 | 0.00 | 19 | 19 | S | 1334 | Streptococcus dysgalactiae |
| 165 | 0.00 | 71 | 71 | S | 1309 | Streptococcus mutans |
| 166 | 0.00 | 61 | 51 | S | 1302 | Streptococcus gordonii |
| 167 | 0.00 | 10 | 0 | S1 | 29390 | Streptococcus gordonii str. Challis |
| 168 | 0.00 | 10 | 10 | S2 | 467705 | Streptococcus gordonii str. Challis substr. CH1 |
| 169 | 0.00 | 50 | 50 | S | 1305 | Streptococcus sanguinis |
| 170 | 0.00 | 46 | 46 | S | 1304 | Streptococcus salivarius |
| 171 | 0.00 | 36 | 36 | S | 1313 | Streptococcus pneumoniae |
| 172 | 0.00 | 31 | 0 | G1 | 671232 | Streptococcus anginosus group |
| 173 | 0.00 | 31 | 31 | S | 1328 | Streptococcus anginosus |
| 174 | 0.00 | 23 | 23 | S | 1314 | Streptococcus pyogenes |
| 175 | 0.00 | 18 | 18 | S | 1311 | Streptococcus agalactiae |
| 176 | 0.00 | 16 | 16 | S | 1303 | Streptococcus oralis |
| 177 | 0.00 | 2519 | 0 | C | 186801 | Clostridia |
| 178 | 0.00 | 2519 | 71 | O | 186802 | Eubacteriales |
| 179 | 0.00 | 1633 | 0 | F | 31979 | Clostridiaceae |
| 180 | 0.00 | 1633 | 723 | G | 1485 | Clostridium |
| 181 | 0.00 | 292 | 292 | S | 1502 | Clostridium perfringens |
| 182 | 0.00 | 238 | 10 | S | 1491 | Clostridium botulinum |
| 183 | 0.00 | 228 | 0 | S1 | 36830 | Clostridium botulinum E |
| 184 | 0.00 | 228 | 228 | S2 | 508767 | Clostridium botulinum E3 str. Alaska E43 |
| 185 | 0.00 | 170 | 170 | S | 1488 | Clostridium acetobutylicum |
| 186 | 0.00 | 167 | 164 | S | 1513 | Clostridium tetani |
| 187 | 0.00 | 3 | 3 | S1 | 212717 | Clostridium tetani E88 |
| 188 | 0.00 | 43 | 43 | S | 1509 | Clostridium sporogenes |
| 189 | 0.00 | 815 | 0 | F | 186804 | Peptostreptococcaceae |
| 190 | 0.00 | 815 | 0 | G | 1257 | Peptostreptococcus |
| 191 | 0.00 | 815 | 815 | S | 1261 | Peptostreptococcus anaerobius |
| 192 | 0.00 | 522 | 0 | C | 909932 | Negativicutes |
| 193 | 0.00 | 522 | 0 | O | 1843489 | Veillonellales |
| 194 | 0.00 | 522 | 0 | F | 31977 | Veillonellaceae |
| 195 | 0.00 | 522 | 0 | G | 29465 | Veillonella |
| 196 | 0.00 | 522 | 522 | S | 29466 | Veillonella parvula |
| 197 | 0.00 | 340 | 0 | C | 1737404 | Tissierellia |
| 198 | 0.00 | 340 | 0 | O | 1737405 | Tissierellales |
| 199 | 0.00 | 340 | 0 | F | 1570339 | Peptoniphilaceae |
| 200 | 0.00 | 340 | 0 | G | 543311 | Parvimonas |
| 201 | 0.00 | 340 | 340 | S | 33033 | Parvimonas micra |
| 202 | 0.00 | 111 | 0 | P | 544448 | Mycoplasmatota |
| 203 | 0.00 | 111 | 0 | O | 2790996 | Mycoplasmoidales |
| 204 | 0.00 | 111 | 0 | F | 2895623 | Metamycoplasmataceae |
| 205 | 0.00 | 111 | 0 | G | 2995234 | Mycoplasmoides |
| 206 | 0.00 | 111 | 0 | S | 2104 | Mycoplasmoides pneumoniae |
| 207 | 0.00 | 111 | 111 | S1 | 1441379 | Mycoplasmoides pneumoniae M29 |
| 208 | 0.00 | 3601 | 0 | P | 203691 | Spirochaetota |
| 209 | 0.00 | 3601 | 11 | C | 203692 | Spirochaetia |
| 210 | 0.00 | 1898 | 0 | O | 1643688 | Leptospirales |
| 211 | 0.00 | 1898 | 0 | F | 170 | Leptospiraceae |
| 212 | 0.00 | 1898 | 820 | G | 171 | Leptospira |
| 213 | 0.00 | 290 | 290 | S | 408139 | Leptospira kmetyi |
| 214 | 0.00 | 191 | 191 | S | 447106 | Leptospira licerasiae |
| 215 | 0.00 | 88 | 88 | S | 409998 | Leptospira wolffii |
| 216 | 0.00 | 77 | 0 | S | 29506 | Leptospira inadai |
| 217 | 0.00 | 77 | 77 | S1 | 293084 | Leptospira inadai serovar Lyme |
| 218 | 0.00 | 71 | 0 | S | 48782 | Leptospira fainei |
| 219 | 0.00 | 71 | 0 | S1 | 293072 | Leptospira fainei serovar Hurstbridge |
| 220 | 0.00 | 71 | 71 | S2 | 1193011 | Leptospira fainei serovar Hurstbridge str. BUT 6 |
| 221 | 0.00 | 65 | 65 | S | 28183 | Leptospira santarosai |
| 222 | 0.00 | 54 | 54 | S | 28182 | Leptospira noguchii |
| 223 | 0.00 | 48 | 48 | S | 28452 | Leptospira alstonii |
| 224 | 0.00 | 39 | 0 | S | 301541 | Leptospira broomii |
| 225 | 0.00 | 39 | 0 | S1 | 1324404 | Leptospira broomii serovar Hurstbridge |
| 226 | 0.00 | 39 | 39 | S2 | 1049789 | Leptospira broomii serovar Hurstbridge str. 5399 |
| 227 | 0.00 | 38 | 38 | S | 173 | Leptospira interrogans |
| 228 | 0.00 | 36 | 36 | S | 100053 | Leptospira alexanderi |
| 229 | 0.00 | 30 | 30 | S | 174 | Leptospira borgpetersenii |
| 230 | 0.00 | 29 | 29 | S | 29507 | Leptospira kirschneri |
| 231 | 0.00 | 22 | 22 | S | 28184 | Leptospira weilii |
| 232 | 0.00 | 1692 | 76 | O | 136 | Spirochaetales |
| 233 | 0.00 | 1276 | 0 | F | 2845253 | Treponemataceae |
| 234 | 0.00 | 1276 | 143 | G | 157 | Treponema |
| 235 | 0.00 | 782 | 782 | S | 156 | Treponema zuelzerae |

| | | | | | | |
|---|---|---|---|---|---|---|
| 236 | 0.00 | 219 | 214 | S | 160 | Treponema pallidum |
| 237 | 0.00 | 5 | 5 | S1 | 168 | Treponema pallidum subsp. pertenue |
| 238 | 0.00 | 132 | 130 | S | 158 | Treponema denticola |
| 239 | 0.00 | 2 | 2 | S1 | 243275 | Treponema denticola ATCC 35405 |
| 240 | 0.00 | 340 | 73 | F | 1643685 | Borreliaceae |
| 241 | 0.00 | 155 | 143 | G | 64895 | Borreliella |
| 242 | 0.00 | 8 | 0 | S | 139 | Borreliella burgdorferi |
| 243 | 0.00 | 8 | 8 | S1 | 445984 | Borreliella burgdorferi Bol26 |
| 244 | 0.00 | 4 | 0 | S | 664662 | Borreliella bavariensis |
| 245 | 0.00 | 4 | 4 | S1 | 290434 | Borrelia garinii subsp. bavariensis PBi |
| 246 | 0.00 | 112 | 0 | G | 138 | Borrelia |
| 247 | 0.00 | 112 | 0 | S | 44449 | Borrelia recurrentis |
| 248 | 0.00 | 112 | 112 | S1 | 412418 | Borrelia recurrentis A1 |
| 249 | 0.00 | 1197 | 0 | D1 | 1783270 | FCB group |
| 250 | 0.00 | 1197 | 0 | D2 | 68336 | Bacteroidota/Chlorobiota group |
| 251 | 0.00 | 1197 | 0 | P | 976 | Bacteroidota |
| 252 | 0.00 | 1197 | 0 | C | 200643 | Bacteroidia |
| 253 | 0.00 | 1197 | 419 | O | 171549 | Bacteroidales |
| 254 | 0.00 | 450 | 0 | F | 2005525 | Tannerellaceae |
| 255 | 0.00 | 450 | 0 | G | 195950 | Tannerella |
| 256 | 0.00 | 450 | 443 | S | 28112 | Tannerella forsythia |
| 257 | 0.00 | 7 | 7 | S1 | 203275 | Tannerella forsythia 92A2 |
| 258 | 0.00 | 328 | 0 | F | 171551 | Porphyromonadaceae |
| 259 | 0.00 | 328 | 0 | G | 836 | Porphyromonas |
| 260 | 0.00 | 328 | 299 | S | 837 | Porphyromonas gingivalis |
| 261 | 0.00 | 29 | 29 | S1 | 242619 | Porphyromonas gingivalis W83 |
| 262 | 0.00 | 761 | 0 | D1 | 1783257 | PVC group |
| 263 | 0.00 | 761 | 0 | P | 204428 | Chlamydiota |
| 264 | 0.00 | 761 | 0 | C | 204429 | Chlamydiia |
| 265 | 0.00 | 761 | 0 | O | 51291 | Chlamydiales |
| 266 | 0.00 | 761 | 0 | F | 809 | Chlamydiaceae |
| 267 | 0.00 | 761 | 0 | F1 | 1113537 | Chlamydia/Chlamydophila group |
| 268 | 0.00 | 761 | 645 | G | 810 | Chlamydia |
| 269 | 0.00 | 35 | 0 | S | 83557 | Chlamydia caviae |
| 270 | 0.00 | 35 | 35 | S1 | 227941 | Chlamydia caviae GPIC |
| 271 | 0.00 | 33 | 0 | S | 83558 | Chlamydia pneumoniae |
| 272 | 0.00 | 33 | 33 | S1 | 406984 | Chlamydia pneumoniae LPCoLN |
| 273 | 0.00 | 18 | 0 | S | 83554 | Chlamydia psittaci |
| 274 | 0.00 | 18 | 18 | S1 | 331636 | Chlamydia psittaci 6BC |
| 275 | 0.00 | 9 | 9 | S | 85991 | Chlamydia pecorum |
| 276 | 0.00 | 8 | 8 | S | 83555 | Chlamydia abortus |
| 277 | 0.00 | 6 | 0 | S | 83556 | Chlamydia felis |
| 278 | 0.00 | 6 | 6 | S1 | 264202 | Chlamydia felis Fe/C-56 |
| 279 | 0.00 | 6 | 6 | S | 83560 | Chlamydia muridarum |
| 280 | 0.00 | 1 | 0 | S | 813 | Chlamydia trachomatis |
| 281 | 0.00 | 1 | 1 | S1 | 272561 | Chlamydia trachomatis D/UW-3/CX |
| 282 | 0.00 | 637 | 0 | P | 32066 | Fusobacteriota |
| 283 | 0.00 | 637 | 0 | C | 203490 | Fusobacteriia |
| 284 | 0.00 | 637 | 0 | O | 203491 | Fusobacteriales |
| 285 | 0.00 | 637 | 0 | F | 203492 | Fusobacteriaceae |
| 286 | 0.00 | 637 | 223 | G | 848 | Fusobacterium |
| 287 | 0.00 | 258 | 258 | S | 851 | Fusobacterium nucleatum |
| 288 | 0.00 | 156 | 156 | S | 849 | Fusobacterium gonidiaformans |
| 289 | 0.00 | 464 | 0 | P | 29547 | Campylobacterota |
| 290 | 0.00 | 464 | 0 | C | 3031852 | Epsilonproteobacteria |
| 291 | 0.00 | 464 | 0 | O | 213849 | Campylobacterales |
| 292 | 0.00 | 464 | 0 | F | 72293 | Helicobacteraceae |
| 293 | 0.00 | 464 | 0 | G | 209 | Helicobacter |
| 294 | 0.00 | 464 | 464 | S | 210 | Helicobacter pylori |
| 295 | 0.02 | 26350 | 2109 | D | 2759 | Eukaryota |
| 296 | 0.01 | 12619 | 0 | D1 | 2698737 | Sar |
| 297 | 0.01 | 12619 | 0 | D2 | 33630 | Alveolata |
| 298 | 0.01 | 12619 | 77 | P | 5794 | Apicomplexa |
| 299 | 0.01 | 7878 | 0 | C | 422676 | Aconoidasida |
| 300 | 0.01 | 7878 | 0 | O | 5819 | Haemosporida |
| 301 | 0.01 | 7878 | 0 | F | 1639119 | Plasmodiidae |
| 302 | 0.01 | 7878 | 24 | G | 5820 | Plasmodium |
| 303 | 0.01 | 7844 | 0 | G1 | 418103 | Plasmodium (Plasmodium) |
| 304 | 0.01 | 7844 | 7844 | S | 5855 | Plasmodium vivax |
| 305 | 0.00 | 10 | 0 | G1 | 418107 | Plasmodium (Laverania) |
| 306 | 0.00 | 10 | 0 | S | 5833 | Plasmodium falciparum |
| 307 | 0.00 | 10 | 10 | S1 | 36329 | Plasmodium falciparum 3D7 |
| 308 | 0.00 | 4664 | 0 | C | 1280412 | Conoidasida |
| 309 | 0.00 | 4664 | 0 | C1 | 5796 | Coccidia |
| 310 | 0.00 | 4664 | 0 | O | 75739 | Eucoccidiorida |
| 311 | 0.00 | 4664 | 0 | O1 | 423054 | Eimeriorina |
| 312 | 0.00 | 4664 | 0 | F | 5809 | Sarcocystidae |
| 313 | 0.00 | 4664 | 0 | G | 5810 | Toxoplasma |
| 314 | 0.00 | 4664 | 0 | S | 5811 | Toxoplasma gondii |
| 315 | 0.00 | 4656 | 4656 | S1 | 508771 | Toxoplasma gondii ME49 |
| 316 | 0.00 | 8 | 8 | S1 | 383379 | Toxoplasma gondii RH |
| 317 | 0.01 | 11240 | 311 | D1 | 33154 | Opisthokonta |
| 318 | 0.01 | 9794 | 0 | K | 33208 | Metazoa |
| 319 | 0.01 | 9794 | 0 | K1 | 6072 | Eumetazoa |
| 320 | 0.01 | 9794 | 0 | K2 | 33213 | Bilateria |
| 321 | 0.01 | 9794 | 58 | K3 | 33317 | Protostomia |
| 322 | 0.01 | 9390 | 0 | K4 | 2697495 | Spiralia |
| 323 | 0.01 | 9390 | 0 | K5 | 1206795 | Lophotrochozoa |
| 324 | 0.01 | 9390 | 154 | P | 6157 | Platyhelminthes |
| 325 | 0.01 | 7846 | 0 | C | 6199 | Cestoda |
| 326 | 0.01 | 7846 | 0 | C1 | 6200 | Eucestoda |
| 327 | 0.01 | 7846 | 0 | O | 6201 | Cyclophyllidea |
| 328 | 0.01 | 7846 | 0 | F | 6208 | Taeniidae |
| 329 | 0.01 | 7846 | 0 | G | 6202 | Taenia |
| 330 | 0.01 | 7846 | 7846 | S | 6204 | Taenia solium |
| 331 | 0.00 | 1390 | 0 | C | 6178 | Trematoda |
| 332 | 0.00 | 1390 | 0 | C1 | 6179 | Digenea |
| 333 | 0.00 | 1390 | 0 | O | 6180 | Strigeidida |
| 334 | 0.00 | 1390 | 0 | O1 | 31244 | Schistosomatoidea |
| 335 | 0.00 | 1390 | 0 | F | 31245 | Schistosomatidae |
| 336 | 0.00 | 1390 | 0 | G | 6181 | Schistosoma |
| 337 | 0.00 | 1390 | 1390 | S | 6183 | Schistosoma mansoni |
| 338 | 0.00 | 346 | 0 | K4 | 1206794 | Ecdysozoa |
| 339 | 0.00 | 346 | 0 | P | 6231 | Nematoda |
| 340 | 0.00 | 346 | 0 | C | 119088 | Enoplea |
| 341 | 0.00 | 346 | 0 | C1 | 1457286 | Dorylaimia |
| 342 | 0.00 | 346 | 0 | O | 6329 | Trichinellida |
| 343 | 0.00 | 346 | 0 | F | 6332 | Trichinellidae |
| 344 | 0.00 | 346 | 0 | G | 6333 | Trichinella |
| 345 | 0.00 | 346 | 346 | S | 6334 | Trichinella spiralis |
| 346 | 0.00 | 1135 | 0 | K | 4751 | Fungi |
| 347 | 0.00 | 1135 | 73 | K1 | 451864 | Dikarya |
| 348 | 0.00 | 779 | 0 | P | 4890 | Ascomycota |
| 349 | 0.00 | 709 | 0 | P1 | 716545 | saccharomyceta |
| 350 | 0.00 | 709 | 0 | P2 | 147538 | Pezizomycotina |
| 351 | 0.00 | 709 | 0 | P3 | 716546 | leotiomyceta |
| 352 | 0.00 | 709 | 0 | C | 147545 | Eurotiomycetes |
| 353 | 0.00 | 709 | 25 | C1 | 451871 | Eurotiomycetidae |
| 354 | 0.00 | 494 | 6 | O | 33183 | Onygenales |

| | | | | | | |
|---|---|---|---|---|---|---|
| 355 | 0.00 | 404 | 21 | F | 299071 | Ajellomycetaceae |
| 356 | 0.00 | 245 | 0 | G | 229219 | Blastomyces |
| 357 | 0.00 | 245 | 0 | S | 5039 | Blastomyces dermatitidis |
| 358 | 0.00 | 245 | 245 | S1 | 559297 | Blastomyces dermatitidis ER-3 |
| 359 | 0.00 | 138 | 0 | G | 5036 | Histoplasma |
| 360 | 0.00 | 138 | 0 | S | 5037 | Histoplasma capsulatum |
| 361 | 0.00 | 138 | 138 | S1 | 447093 | Histoplasma capsulatum G186AR |
| 362 | 0.00 | 84 | 0 | F | 33184 | Onygenaceae |
| 363 | 0.00 | 84 | 35 | G | 5500 | Coccidioides |
| 364 | 0.00 | 29 | 0 | S | 199306 | Coccidioides posadasii |
| 365 | 0.00 | 29 | 29 | S1 | 222929 | Coccidioides posadasii C735 delta SOWgp |
| 366 | 0.00 | 20 | 0 | S | 5501 | Coccidioides immitis |
| 367 | 0.00 | 20 | 20 | S1 | 246410 | Coccidioides immitis RS |
| 368 | 0.00 | 190 | 0 | O | 5042 | Eurotiales |
| 369 | 0.00 | 190 | 0 | F | 1131492 | Aspergillaceae |
| 370 | 0.00 | 190 | 0 | G | 5052 | Aspergillus |
| 371 | 0.00 | 190 | 0 | G1 | 2720872 | Aspergillus subgen. Fumigati |
| 372 | 0.00 | 190 | 0 | S | 746128 | Aspergillus fumigatus |
| 373 | 0.00 | 190 | 190 | S1 | 330879 | Aspergillus fumigatus Af293 |
| 374 | 0.00 | 70 | 0 | P1 | 451866 | Taphrinomycotina |
| 375 | 0.00 | 70 | 0 | C | 147553 | Pneumocystidomycetes |
| 376 | 0.00 | 70 | 0 | O | 37987 | Pneumocystidales |
| 377 | 0.00 | 70 | 0 | F | 44281 | Pneumocystidaceae |
| 378 | 0.00 | 70 | 19 | G | 4753 | Pneumocystis |
| 379 | 0.00 | 32 | 0 | S | 4754 | Pneumocystis carinii |
| 380 | 0.00 | 32 | 32 | S1 | 1408658 | Pneumocystis carinii B80 |
| 381 | 0.00 | 19 | 0 | S | 42068 | Pneumocystis jirovecii |
| 382 | 0.00 | 19 | 19 | S1 | 1408657 | Pneumocystis jirovecii RU7 |
| 383 | 0.00 | 283 | 0 | P | 5204 | Basidiomycota |
| 384 | 0.00 | 283 | 0 | P1 | 5302 | Agaricomycotina |
| 385 | 0.00 | 283 | 0 | C | 155616 | Tremellomycetes |
| 386 | 0.00 | 283 | 0 | O | 5234 | Tremellales |
| 387 | 0.00 | 283 | 0 | F | 1884633 | Cryptococcaceae |
| 388 | 0.00 | 283 | 179 | G | 5206 | Cryptococcus |
| 389 | 0.00 | 85 | 0 | G1 | 1884637 | Cryptococcus gattii species complex |
| 390 | 0.00 | 85 | 0 | S | 37769 | Cryptococcus gattii VGI |
| 391 | 0.00 | 85 | 85 | S1 | 367775 | Cryptococcus gattii WM276 |
| 392 | 0.00 | 19 | 0 | G1 | 1897064 | Cryptococcus neoformans species complex |
| 393 | 0.00 | 19 | 0 | S | 5207 | Cryptococcus neoformans |
| 394 | 0.00 | 19 | 0 | S1 | 40410 | Cryptococcus neoformans var. neoformans |
| 395 | 0.00 | 19 | 19 | S2 | 214684 | Cryptococcus neoformans var. neoformans JEC21 |
| 396 | 0.00 | 382 | 0 | D1 | 2611352 | Discoba |
| 397 | 0.00 | 382 | 0 | P | 33682 | Euglenozoa |
| 398 | 0.00 | 382 | 0 | C | 5653 | Kinetoplastea |
| 399 | 0.00 | 382 | 0 | C1 | 2704647 | Metakinetoplastina |
| 400 | 0.00 | 382 | 0 | O | 2704949 | Trypanosomatida |
| 401 | 0.00 | 382 | 0 | F | 5654 | Trypanosomatidae |
| 402 | 0.00 | 382 | 22 | G | 5690 | Trypanosoma |
| 403 | 0.00 | 276 | 0 | G1 | 47570 | Schizotrypanum |
| 404 | 0.00 | 276 | 276 | S | 5693 | Trypanosoma cruzi |
| 405 | 0.00 | 84 | 0 | G1 | 39700 | Trypanozoon |
| 406 | 0.00 | 84 | 0 | S | 5691 | Trypanosoma brucei |
| 407 | 0.00 | 84 | 0 | S1 | 5702 | Trypanosoma brucei brucei |
| 408 | 0.00 | 84 | 84 | S2 | 185431 | Trypanosoma brucei brucei TREU927 |
| 409 | 0.00 | 41 | 0 | D | 2157 | Archaea |
| 410 | 0.00 | 41 | 0 | P | 28890 | Euryarchaeota |
| 411 | 0.00 | 41 | 0 | P1 | 2283794 | Methanomada group |
| 412 | 0.00 | 41 | 0 | C | 183925 | Methanobacteria |
| 413 | 0.00 | 41 | 0 | O | 2158 | Methanobacteriales |
| 414 | 0.00 | 41 | 0 | F | 2159 | Methanobacteriaceae |
| 415 | 0.00 | 41 | 0 | G | 2172 | Methanobrevibacter |
| 416 | 0.00 | 41 | 41 | S | 66851 | Methanobrevibacter oralis |
| 417 | 0.00 | 3677 | 0 | D | 10239 | Viruses |
| 418 | 0.00 | 3675 | 0 | D1 | 2731341 | Duplodnaviria |
| 419 | 0.00 | 3675 | 0 | K | 2731360 | Heunggongvirae |
| 420 | 0.00 | 3675 | 0 | P | 2731361 | Peploviricota |
| 421 | 0.00 | 3675 | 0 | C | 2731363 | Herviviricetes |
| 422 | 0.00 | 3675 | 0 | O | 548681 | Herpesvirales |
| 423 | 0.00 | 3675 | 0 | F | 10292 | Herpesviridae |
| 424 | 0.00 | 3675 | 0 | F1 | 10293 | Alphaherpesvirinae |
| 425 | 0.00 | 3675 | 17 | G | 10319 | Varicellovirus |
| 426 | 0.00 | 3658 | 0 | S | 3050278 | Varicellovirus equidalpha4 |
| 427 | 0.00 | 3658 | 3658 | S1 | 10331 | Equid alphaherpesvirus 4 |
| 428 | 0.00 | 2 | 0 | D1 | 2732004 | Varidnaviria |
| 429 | 0.00 | 2 | 0 | K | 2732005 | Bamfordvirae |
| 430 | 0.00 | 2 | 0 | P | 2732007 | Nucleocytoviricota |
| 431 | 0.00 | 2 | 0 | C | 2732523 | Megaviricetes |
| 432 | 0.00 | 2 | 0 | O | 2732555 | Pimascovirales |
| 433 | 0.00 | 2 | 0 | F | 944644 | Marseilleviridae |
| 434 | 0.00 | 2 | 0 | G | 1513458 | Marseillevirus |
| 435 | 0.00 | 2 | 0 | G1 | 1813598 | unclassified Marseillevirus |
| 436 | 0.00 | 2 | 2 | S | 1826170 | Tokyovirus A1 |

**Table S3e:** Results of the Kraken2 analysis for library ERR6466112 based on a custom-built pathogen list (Table S1)

| Sr | Percentage of fragments covered | Number of fragments covered | Number of fragments assigned | rank code | NCBI taxonomic ID | Name |
|---|---|---|---|---|---|---|
| 1 | 99.76 | 144963738 | 144963738 | U | 0 | unclassified |
| 2 | 0.24 | 353960 | 1 | R | 1 | root |
| 3 | 0.24 | 350297 | 2356 | R1 | 131567 | cellular organisms |
| 4 | 0.22 | 321385 | 9676 | D | 2 | Bacteria |
| 5 | 0.16 | 239503 | 6447 | P | 1224 | Pseudomonadota |
| 6 | 0.12 | 173754 | 1723 | C | 28216 | Betaproteobacteria |
| 7 | 0.12 | 167678 | 6125 | O | 80840 | Burkholderiales |
| 8 | 0.07 | 104214 | 0 | F | 119060 | Burkholderiaceae |
| 9 | 0.07 | 104214 | 14962 | G | 32008 | Burkholderia |
| 10 | 0.03 | 49342 | 0 | G1 | 87882 | Burkholderia cepacia complex |
| 11 | 0.03 | 49342 | 49342 | S | 292 | Burkholderia cepacia |
| 12 | 0.03 | 39910 | 33642 | G1 | 111527 | pseudomallei group |
| 13 | 0.00 | 4786 | 4786 | S | 28450 | Burkholderia pseudomallei |
| 14 | 0.00 | 1482 | 1482 | S | 13373 | Burkholderia mallei |
| 15 | 0.04 | 57339 | 501 | F | 506 | Alcaligenaceae |
| 16 | 0.04 | 56019 | 5715 | G | 517 | Bordetella |
| 17 | 0.02 | 26142 | 26142 | S | 520 | Bordetella pertussis |
| 18 | 0.02 | 24162 | 24162 | S | 94624 | Bordetella petrii |
| 19 | 0.00 | 819 | 0 | G | 29574 | Taylorella |
| 20 | 0.00 | 819 | 819 | S | 29575 | Taylorella equigenitalis |
| 21 | 0.00 | 4353 | 0 | O | 206351 | Neisseriales |
| 22 | 0.00 | 4353 | 0 | F | 481 | Neisseriaceae |
| 23 | 0.00 | 4353 | 2072 | G | 482 | Neisseria |
| 24 | 0.00 | 983 | 983 | S | 485 | Neisseria gonorrhoeae |
| 25 | 0.00 | 664 | 664 | S | 487 | Neisseria meningitidis |
| 26 | 0.00 | 634 | 634 | S | 483 | Neisseria cinerea |
| 27 | 0.03 | 46826 | 3803 | C | 1236 | Gammaproteobacteria |
| 28 | 0.02 | 30399 | 872 | O | 91347 | Enterobacterales |
| 29 | 0.02 | 27620 | 6029 | F | 543 | Enterobacteriaceae |
| 30 | 0.01 | 15677 | 0 | F1 | 2890311 | Klebsiella/Raoultella group |
| 31 | 0.01 | 15677 | 1258 | G | 570 | Klebsiella |
| 32 | 0.01 | 8634 | 8070 | S | 573 | Klebsiella pneumoniae |
| 33 | 0.00 | 564 | 564 | S1 | 39831 | Klebsiella pneumoniae subsp. rhinoscleromatis |
| 34 | 0.00 | 5785 | 5785 | S | 571 | Klebsiella oxytoca |
| 35 | 0.00 | 4230 | 0 | G | 590 | Salmonella |
| 36 | 0.00 | 4230 | 0 | S | 28901 | Salmonella enterica |
| 37 | 0.00 | 4230 | 0 | S1 | 59201 | Salmonella enterica subsp. enterica |
| 38 | 0.00 | 4230 | 4230 | S2 | 594 | Salmonella enterica subsp. enterica serovar Gallinarum |
| 39 | 0.00 | 1440 | 0 | G | 561 | Escherichia |
| 40 | 0.00 | 1440 | 1018 | S | 562 | Escherichia coli |
| 41 | 0.00 | 318 | 318 | S1 | 83334 | Escherichia coli O157:H7 |
| 42 | 0.00 | 74 | 74 | S1 | 168927 | Escherichia coli O111:H- |
| 43 | 0.00 | 30 | 30 | S1 | 244319 | Escherichia coli O26:H11 |
| 44 | 0.00 | 244 | 107 | G | 620 | Shigella |
| 45 | 0.00 | 95 | 95 | S | 623 | Shigella flexneri |
| 46 | 0.00 | 23 | 23 | S | 621 | Shigella boydii |
| 47 | 0.00 | 18 | 18 | S | 622 | Shigella dysenteriae |
| 48 | 0.00 | 1 | 1 | S | 624 | Shigella sonnei |
| 49 | 0.00 | 1907 | 0 | F | 1903411 | Yersiniaceae |
| 50 | 0.00 | 1907 | 949 | G | 629 | Yersinia |
| 51 | 0.00 | 517 | 347 | G1 | 1649845 | Yersinia pseudotuberculosis complex |
| 52 | 0.00 | 145 | 145 | S | 632 | Yersinia pestis |
| 53 | 0.00 | 25 | 25 | S | 633 | Yersinia pseudotuberculosis |
| 54 | 0.00 | 441 | 441 | S | 630 | Yersinia enterocolitica |
| 55 | 0.00 | 5919 | 339 | O | 118969 | Legionellales |
| 56 | 0.00 | 3256 | 0 | F | 118968 | Coxiellaceae |
| 57 | 0.00 | 3256 | 0 | G | 776 | Coxiella |
| 58 | 0.00 | 3256 | 3256 | S | 777 | Coxiella burnetii |
| 59 | 0.00 | 2324 | 0 | F | 444 | Legionellaceae |
| 60 | 0.00 | 2324 | 0 | G | 445 | Legionella |
| 61 | 0.00 | 2324 | 2324 | S | 446 | Legionella pneumophila |
| 62 | 0.00 | 2384 | 0 | O | 2887326 | Moraxellales |
| 63 | 0.00 | 2384 | 0 | F | 468 | Moraxellaceae |
| 64 | 0.00 | 2384 | 0 | G | 475 | Moraxella |
| 65 | 0.00 | 2384 | 2384 | S | 480 | Moraxella catarrhalis |
| 66 | 0.00 | 2116 | 0 | O | 135623 | Vibrionales |
| 67 | 0.00 | 2116 | 0 | F | 641 | Vibrionaceae |
| 68 | 0.00 | 2116 | 695 | G | 662 | Vibrio |
| 69 | 0.00 | 671 | 671 | S | 666 | Vibrio cholerae |
| 70 | 0.00 | 420 | 0 | G1 | 717610 | Vibrio harveyi group |
| 71 | 0.00 | 420 | 420 | S | 670 | Vibrio parahaemolyticus |
| 72 | 0.00 | 330 | 330 | S | 672 | Vibrio vulnificus |
| 73 | 0.00 | 1147 | 0 | O | 72273 | Thiotrichales |
| 74 | 0.00 | 1147 | 0 | F | 34064 | Francisellaceae |
| 75 | 0.00 | 1147 | 0 | G | 262 | Francisella |
| 76 | 0.00 | 1147 | 1147 | S | 263 | Francisella tularensis |
| 77 | 0.00 | 1058 | 0 | O | 135625 | Pasteurellales |
| 78 | 0.00 | 1058 | 336 | F | 712 | Pasteurellaceae |
| 79 | 0.00 | 365 | 0 | G | 724 | Haemophilus |
| 80 | 0.00 | 365 | 0 | S | 727 | Haemophilus influenzae |
| 81 | 0.00 | 365 | 365 | S1 | 725 | Haemophilus influenzae biotype aegyptius |
| 82 | 0.00 | 357 | 0 | G | 713 | Actinobacillus |
| 83 | 0.00 | 357 | 0 | S | 718 | Actinobacillus equuli |
| 84 | 0.00 | 357 | 357 | S1 | 202947 | Actinobacillus equuli subsp. equuli |
| 85 | 0.01 | 12476 | 78 | C | 28211 | Alphaproteobacteria |
| 86 | 0.01 | 11483 | 1689 | O | 356 | Hyphomicrobiales |
| 87 | 0.01 | 8046 | 0 | F | 118882 | Brucellaceae |
| 88 | 0.01 | 8046 | 0 | F1 | 2826938 | Brucella/Ochrobactrum group |
| 89 | 0.01 | 8046 | 7841 | G | 234 | Brucella |
| 90 | 0.00 | 104 | 0 | S | 236 | Brucella ovis |
| 91 | 0.00 | 104 | 104 | S1 | 444178 | Brucella ovis ATCC 25840 |
| 92 | 0.00 | 65 | 0 | S | 444163 | Brucella microti |
| 93 | 0.00 | 65 | 65 | S1 | 568815 | Brucella microti CCM 4915 |
| 94 | 0.00 | 26 | 0 | S | 29459 | Brucella melitensis |
| 95 | 0.00 | 26 | 0 | S1 | 644337 | Brucella melitensis bv. 1 |
| 96 | 0.00 | 26 | 26 | S2 | 224914 | Brucella melitensis bv. 1 str. 16M |
| 97 | 0.00 | 7 | 0 | S | 235 | Brucella abortus |
| 98 | 0.00 | 7 | 7 | S1 | 359391 | Brucella abortus 2308 |
| 99 | 0.00 | 3 | 0 | S | 29461 | Brucella suis |
| 100 | 0.00 | 3 | 3 | S1 | 204722 | Brucella suis 1330 |
| 101 | 0.00 | 1748 | 0 | F | 772 | Bartonellaceae |
| 102 | 0.00 | 1748 | 1061 | G | 773 | Bartonella |
| 103 | 0.00 | 289 | 289 | S | 38323 | Bartonella henselae |
| 104 | 0.00 | 249 | 0 | S | 774 | Bartonella bacilliformis |
| 105 | 0.00 | 249 | 249 | S1 | 360095 | Bartonella bacilliformis KC583 |
| 106 | 0.00 | 149 | 149 | S | 803 | Bartonella quintana |
| 107 | 0.00 | 915 | 0 | O | 766 | Rickettsiales |
| 108 | 0.00 | 915 | 0 | F | 775 | Rickettsiaceae |
| 109 | 0.00 | 915 | 0 | F1 | 33988 | Rickettsieae |
| 110 | 0.00 | 915 | 623 | G | 780 | Rickettsia |
| 111 | 0.00 | 258 | 150 | G1 | 114277 | spotted fever group |
| 112 | 0.00 | 47 | 47 | S | 42862 | Rickettsia felis |
| 113 | 0.00 | 29 | 29 | S | 35790 | Rickettsia japonica |
| 114 | 0.00 | 28 | 0 | S | 786 | Rickettsia akari |
| 115 | 0.00 | 28 | 28 | S1 | 293614 | Rickettsia akari str. Hartford |
| 116 | 0.00 | 3 | 3 | S | 783 | Rickettsia rickettsii |

| | | | | | | |
|---|---|---|---|---|---|---|
| 117 | 0.00 | 1 | 0 | G2 | 266068 | Rickettsia sibirica subgroup |
| 118 | 0.00 | 1 | 0 | S | 35793 | Rickettsia sibirica |
| 119 | 0.00 | 1 | 1 | S1 | 272951 | Rickettsia sibirica 246 |
| 120 | 0.00 | 34 | 0 | G1 | 114292 | typhus group |
| 121 | 0.00 | 17 | 17 | S | 782 | Rickettsia prowazekii |
| 122 | 0.00 | 17 | 17 | S | 785 | Rickettsia typhi |
| 123 | 0.05 | 65421 | 419 | D1 | 1783272 | Terrabacteria group |
| 124 | 0.04 | 55734 | 0 | P | 201174 | Actinomycetota |
| 125 | 0.04 | 55734 | 0 | C | 1760 | Actinomycetes |
| 126 | 0.04 | 55734 | 4641 | O | 85007 | Mycobacteriales |
| 127 | 0.03 | 37451 | 1439 | F | 1762 | Mycobacteriaceae |
| 128 | 0.02 | 32078 | 3444 | G | 1763 | Mycobacterium |
| 129 | 0.01 | 19233 | 4023 | G1 | 120793 | Mycobacterium avium complex (MAC) |
| 130 | 0.00 | 6076 | 6076 | S | 1764 | Mycobacterium avium |
| 131 | 0.00 | 4568 | 4568 | S | 1767 | Mycobacterium intracellulare |
| 132 | 0.00 | 4566 | 4566 | S | 339268 | Mycobacterium colombiense |
| 133 | 0.00 | 5101 | 5101 | S | 1768 | Mycobacterium kansasii |
| 134 | 0.00 | 3696 | 3087 | G1 | 77643 | Mycobacterium tuberculosis complex |
| 135 | 0.00 | 360 | 360 | S | 78331 | Mycobacterium canettii |
| 136 | 0.00 | 249 | 249 | S | 1773 | Mycobacterium tuberculosis |
| 137 | 0.00 | 604 | 604 | S | 1769 | Mycobacterium leprae |
| 138 | 0.00 | 3934 | 0 | G | 670516 | Mycobacteroides |
| 139 | 0.00 | 3934 | 0 | S | 36809 | Mycobacteroides abscessus |
| 140 | 0.00 | 3934 | 3934 | S1 | 319705 | Mycobacteroides abscessus subsp. bolletii |
| 141 | 0.01 | 11239 | 0 | F | 85025 | Nocardiaceae |
| 142 | 0.01 | 11239 | 0 | G | 2979332 | Prescottella |
| 143 | 0.01 | 11239 | 11239 | S | 43767 | Prescottella equi |
| 144 | 0.00 | 2403 | 0 | F | 1653 | Corynebacteriaceae |
| 145 | 0.00 | 2403 | 537 | G | 1716 | Corynebacterium |
| 146 | 0.00 | 1467 | 1467 | S | 1717 | Corynebacterium diphtheriae |
| 147 | 0.00 | 399 | 0 | S | 1719 | Corynebacterium pseudotuberculosis |
| 148 | 0.00 | 399 | 399 | S1 | 1087451 | Corynebacterium pseudotuberculosis 31 |
| 149 | 0.01 | 9187 | 732 | P | 1239 | Bacillota |
| 150 | 0.00 | 5252 | 370 | C | 91061 | Bacilli |
| 151 | 0.00 | 3859 | 280 | O | 1385 | Bacillales |
| 152 | 0.00 | 2749 | 0 | F | 186817 | Bacillaceae |
| 153 | 0.00 | 2749 | 0 | G | 1386 | Bacillus |
| 154 | 0.00 | 2749 | 1614 | G1 | 86661 | Bacillus cereus group |
| 155 | 0.00 | 722 | 722 | S | 1396 | Bacillus cereus |
| 156 | 0.00 | 413 | 0 | S | 1392 | Bacillus anthracis |
| 157 | 0.00 | 413 | 413 | S1 | 261594 | Bacillus anthracis str. 'Ames Ancestor' |
| 158 | 0.00 | 830 | 0 | F | 90964 | Staphylococcaceae |
| 159 | 0.00 | 830 | 0 | G | 1279 | Staphylococcus |
| 160 | 0.00 | 830 | 830 | S | 1280 | Staphylococcus aureus |
| 161 | 0.00 | 1023 | 0 | O | 186826 | Lactobacillales |
| 162 | 0.00 | 1023 | 0 | F | 1300 | Streptococcaceae |
| 163 | 0.00 | 1023 | 602 | G | 1301 | Streptococcus |
| 164 | 0.00 | 91 | 91 | S | 1309 | Streptococcus mutans |
| 165 | 0.00 | 69 | 62 | S | 1302 | Streptococcus gordonii |
| 166 | 0.00 | 7 | 0 | S1 | 29390 | Streptococcus gordonii str. Challis |
| 167 | 0.00 | 7 | 7 | S2 | 467705 | Streptococcus gordonii str. Challis substr. CH1 |
| 168 | 0.00 | 66 | 0 | G1 | 119603 | Streptococcus dysgalactiae group |
| 169 | 0.00 | 56 | 0 | S | 1336 | Streptococcus equi |
| 170 | 0.00 | 56 | 56 | S1 | 40041 | Streptococcus equi subsp. zooepidemicus |
| 171 | 0.00 | 10 | 10 | S | 1334 | Streptococcus dysgalactiae |
| 172 | 0.00 | 45 | 45 | S | 1304 | Streptococcus salivarius |
| 173 | 0.00 | 33 | 33 | S | 1305 | Streptococcus sanguinis |
| 174 | 0.00 | 33 | 0 | G1 | 671232 | Streptococcus anginosus group |
| 175 | 0.00 | 33 | 33 | S | 1328 | Streptococcus anginosus |
| 176 | 0.00 | 29 | 29 | S | 1313 | Streptococcus pneumoniae |
| 177 | 0.00 | 23 | 23 | S | 1311 | Streptococcus agalactiae |
| 178 | 0.00 | 17 | 17 | S | 1314 | Streptococcus pyogenes |
| 179 | 0.00 | 15 | 15 | S | 1303 | Streptococcus oralis |
| 180 | 0.00 | 2323 | 0 | C | 186801 | Clostridia |
| 181 | 0.00 | 2323 | 78 | O | 186802 | Eubacteriales |
| 182 | 0.00 | 1492 | 0 | F | 31979 | Clostridiaceae |
| 183 | 0.00 | 1492 | 658 | G | 1485 | Clostridium |
| 184 | 0.00 | 281 | 281 | S | 1502 | Clostridium perfringens |
| 185 | 0.00 | 233 | 5 | S | 1491 | Clostridium botulinum |
| 186 | 0.00 | 228 | 0 | S1 | 36830 | Clostridium botulinum E |
| 187 | 0.00 | 228 | 228 | S2 | 508767 | Clostridium botulinum E3 str. Alaska E43 |
| 188 | 0.00 | 153 | 153 | S | 1513 | Clostridium tetani |
| 189 | 0.00 | 141 | 141 | S | 1488 | Clostridium acetobutylicum |
| 190 | 0.00 | 26 | 26 | S | 1509 | Clostridium sporogenes |
| 191 | 0.00 | 753 | 0 | F | 186804 | Peptostreptococcaceae |
| 192 | 0.00 | 753 | 0 | G | 1257 | Peptostreptococcus |
| 193 | 0.00 | 753 | 753 | S | 1261 | Peptostreptococcus anaerobius |
| 194 | 0.00 | 490 | 0 | C | 909932 | Negativicutes |
| 195 | 0.00 | 490 | 0 | O | 1843489 | Veillonellales |
| 196 | 0.00 | 490 | 0 | F | 31977 | Veillonellaceae |
| 197 | 0.00 | 490 | 0 | G | 29465 | Veillonella |
| 198 | 0.00 | 490 | 490 | S | 29466 | Veillonella parvula |
| 199 | 0.00 | 390 | 0 | C | 1737404 | Tissierellia |
| 200 | 0.00 | 390 | 0 | O | 1737405 | Tissierellales |
| 201 | 0.00 | 390 | 0 | F | 1570339 | Peptoniphilaceae |
| 202 | 0.00 | 390 | 0 | G | 543311 | Parvimonas |
| 203 | 0.00 | 390 | 390 | S | 33033 | Parvimonas micra |
| 204 | 0.00 | 81 | 0 | P | 544448 | Mycoplasmatota |
| 205 | 0.00 | 81 | 0 | O | 2790996 | Mycoplasmoidales |
| 206 | 0.00 | 81 | 0 | F | 2895623 | Metamycoplasmataceae |
| 207 | 0.00 | 81 | 0 | G | 2995234 | Mycoplasmoides |
| 208 | 0.00 | 81 | 0 | S | 2104 | Mycoplasmoides pneumoniae |
| 209 | 0.00 | 81 | 81 | S1 | 1441379 | Mycoplasmoides pneumoniae M29 |
| 210 | 0.00 | 3807 | 0 | P | 203691 | Spirochaetota |
| 211 | 0.00 | 3807 | 9 | C | 203692 | Spirochaetia |
| 212 | 0.00 | 1999 | 0 | O | 1643688 | Leptospirales |
| 213 | 0.00 | 1999 | 0 | F | 170 | Leptospiraceae |
| 214 | 0.00 | 1999 | 915 | G | 171 | Leptospira |
| 215 | 0.00 | 277 | 277 | S | 408139 | Leptospira kmetyi |
| 216 | 0.00 | 169 | 169 | S | 447106 | Leptospira licerasiae |
| 217 | 0.00 | 107 | 107 | S | 409998 | Leptospira wolffii |
| 218 | 0.00 | 89 | 0 | S | 48782 | Leptospira fainei |
| 219 | 0.00 | 89 | 0 | S1 | 293072 | Leptospira fainei serovar Hurstbridge |
| 220 | 0.00 | 89 | 89 | S2 | 1193011 | Leptospira fainei serovar Hurstbridge str. BUT 6 |
| 221 | 0.00 | 88 | 88 | S | 28183 | Leptospira santarosai |
| 222 | 0.00 | 63 | 63 | S | 28182 | Leptospira noguchii |
| 223 | 0.00 | 61 | 0 | S | 29506 | Leptospira inadai |
| 224 | 0.00 | 61 | 61 | S1 | 293084 | Leptospira inadai serovar Lyme |
| 225 | 0.00 | 58 | 58 | S | 28452 | Leptospira alstonii |
| 226 | 0.00 | 43 | 43 | S | 174 | Leptospira borgpetersenii |
| 227 | 0.00 | 38 | 38 | S | 100053 | Leptospira alexanderi |
| 228 | 0.00 | 28 | 28 | S | 28184 | Leptospira weilii |
| 229 | 0.00 | 26 | 0 | S | 301541 | Leptospira broomii |
| 230 | 0.00 | 26 | 0 | S1 | 1324404 | Leptospira broomii serovar Hurstbridge |
| 231 | 0.00 | 26 | 26 | S2 | 1049789 | Leptospira broomii serovar Hurstbridge str. 5399 |
| 232 | 0.00 | 23 | 23 | S | 29507 | Leptospira kirschneri |
| 233 | 0.00 | 14 | 14 | S | 173 | Leptospira interrogans |
| 234 | 0.00 | 1799 | 49 | O | 136 | Spirochaetales |
| 235 | 0.00 | 1432 | 0 | F | 2845253 | Treponemataceae |

| | | | | | | |
|---|---|---|---|---|---|---|
| 236 | 0.00 | 1432 | 138 | G | 157 | Treponema |
| 237 | 0.00 | 938 | 938 | S | 156 | Treponema zuelzerae |
| 238 | 0.00 | 206 | 202 | S | 160 | Treponema pallidum |
| 239 | 0.00 | 4 | 4 | S1 | 168 | Treponema pallidum subsp. pertenue |
| 240 | 0.00 | 150 | 149 | S | 158 | Treponema denticola |
| 241 | 0.00 | 1 | 1 | S1 | 243275 | Treponema denticola ATCC 35405 |
| 242 | 0.00 | 318 | 113 | F | 1643685 | Borreliaceae |
| 243 | 0.00 | 135 | 122 | G | 64895 | Borreliella |
| 244 | 0.00 | 9 | 0 | S | 139 | Borreliella burgdorferi |
| 245 | 0.00 | 9 | 9 | S1 | 445984 | Borreliella burgdorferi Bol26 |
| 246 | 0.00 | 2 | 2 | S | 29518 | Borreliella afzelii |
| 247 | 0.00 | 2 | 0 | S | 664662 | Borreliella bavariensis |
| 248 | 0.00 | 2 | 2 | S1 | 290434 | Borrelia garinii subsp. bavariensis PBi |
| 249 | 0.00 | 70 | 0 | G | 138 | Borrelia |
| 250 | 0.00 | 70 | 0 | S | 44449 | Borrelia recurrentis |
| 251 | 0.00 | 70 | 70 | S1 | 412418 | Borrelia recurrentis A1 |
| 252 | 0.00 | 1150 | 0 | D1 | 1783270 | FCB group |
| 253 | 0.00 | 1150 | 0 | D2 | 68336 | Bacteroidota/Chlorobiota group |
| 254 | 0.00 | 1150 | 0 | P | 976 | Bacteroidota |
| 255 | 0.00 | 1150 | 0 | C | 200643 | Bacteroidia |
| 256 | 0.00 | 1150 | 417 | O | 171549 | Bacteroidales |
| 257 | 0.00 | 429 | 0 | F | 2005525 | Tannerellaceae |
| 258 | 0.00 | 429 | 0 | G | 195950 | Tannerella |
| 259 | 0.00 | 429 | 424 | S | 28112 | Tannerella forsythia |
| 260 | 0.00 | 5 | 5 | S1 | 203275 | Tannerella forsythia 92A2 |
| 261 | 0.00 | 304 | 0 | F | 171551 | Porphyromonadaceae |
| 262 | 0.00 | 304 | 0 | G | 836 | Porphyromonas |
| 263 | 0.00 | 304 | 268 | S | 837 | Porphyromonas gingivalis |
| 264 | 0.00 | 36 | 36 | S1 | 242619 | Porphyromonas gingivalis W83 |
| 265 | 0.00 | 682 | 0 | D1 | 1783257 | PVC group |
| 266 | 0.00 | 682 | 0 | P | 204428 | Chlamydiota |
| 267 | 0.00 | 682 | 0 | C | 204429 | Chlamydiia |
| 268 | 0.00 | 682 | 0 | O | 51291 | Chlamydiales |
| 269 | 0.00 | 682 | 0 | F | 809 | Chlamydiaceae |
| 270 | 0.00 | 682 | 0 | F1 | 1113537 | Chlamydia/Chlamydophila group |
| 271 | 0.00 | 682 | 566 | G | 810 | Chlamydia |
| 272 | 0.00 | 45 | 0 | S | 83558 | Chlamydia pneumoniae |
| 273 | 0.00 | 45 | 45 | S1 | 406984 | Chlamydia pneumoniae LPCoLN |
| 274 | 0.00 | 43 | 0 | S | 83557 | Chlamydia caviae |
| 275 | 0.00 | 43 | 43 | S1 | 227941 | Chlamydia caviae GPIC |
| 276 | 0.00 | 9 | 9 | S | 85991 | Chlamydia pecorum |
| 277 | 0.00 | 7 | 7 | S | 83560 | Chlamydia muridarum |
| 278 | 0.00 | 6 | 0 | S | 83556 | Chlamydia felis |
| 279 | 0.00 | 6 | 6 | S1 | 264202 | Chlamydia felis Fe/C-56 |
| 280 | 0.00 | 4 | 0 | S | 813 | Chlamydia trachomatis |
| 281 | 0.00 | 4 | 4 | S1 | 272561 | Chlamydia trachomatis D/UW-3/CX |
| 282 | 0.00 | 1 | 0 | S | 83554 | Chlamydia psittaci |
| 283 | 0.00 | 1 | 1 | S1 | 331636 | Chlamydia psittaci 6BC |
| 284 | 0.00 | 1 | 1 | S | 83555 | Chlamydia abortus |
| 285 | 0.00 | 664 | 0 | P | 32066 | Fusobacteriota |
| 286 | 0.00 | 664 | 0 | C | 203490 | Fusobacteriia |
| 287 | 0.00 | 664 | 0 | O | 203491 | Fusobacteriales |
| 288 | 0.00 | 664 | 0 | F | 203492 | Fusobacteriaceae |
| 289 | 0.00 | 664 | 237 | G | 848 | Fusobacterium |
| 290 | 0.00 | 272 | 272 | S | 851 | Fusobacterium nucleatum |
| 291 | 0.00 | 155 | 155 | S | 849 | Fusobacterium gonidiaformans |
| 292 | 0.00 | 482 | 0 | P | 29547 | Campylobacterota |
| 293 | 0.00 | 482 | 0 | C | 3031852 | Epsilonproteobacteria |
| 294 | 0.00 | 482 | 0 | O | 213849 | Campylobacterales |
| 295 | 0.00 | 482 | 0 | F | 72293 | Helicobacteraceae |
| 296 | 0.00 | 482 | 0 | G | 209 | Helicobacter |
| 297 | 0.00 | 482 | 482 | S | 210 | Helicobacter pylori |
| 298 | 0.02 | 26515 | 2066 | D | 2759 | Eukaryota |
| 299 | 0.01 | 12703 | 0 | D1 | 2698737 | Sar |
| 300 | 0.01 | 12703 | 0 | D2 | 33630 | Alveolata |
| 301 | 0.01 | 12703 | 59 | P | 5794 | Apicomplexa |
| 302 | 0.01 | 7878 | 0 | C | 422676 | Aconoidasida |
| 303 | 0.01 | 7878 | 0 | O | 5819 | Haemosporida |
| 304 | 0.01 | 7878 | 0 | F | 1639119 | Plasmodiidae |
| 305 | 0.01 | 7878 | 33 | G | 5820 | Plasmodium |
| 306 | 0.01 | 7822 | 0 | G1 | 418103 | Plasmodium (Plasmodium) |
| 307 | 0.01 | 7822 | 7822 | S | 5855 | Plasmodium vivax |
| 308 | 0.00 | 23 | 0 | G1 | 418107 | Plasmodium (Laverania) |
| 309 | 0.00 | 23 | 0 | S | 5833 | Plasmodium falciparum |
| 310 | 0.00 | 23 | 23 | S1 | 36329 | Plasmodium falciparum 3D7 |
| 311 | 0.00 | 4766 | 0 | C | 1280412 | Conoidasida |
| 312 | 0.00 | 4766 | 0 | C1 | 5796 | Coccidia |
| 313 | 0.00 | 4766 | 0 | O | 75739 | Eucoccidiorida |
| 314 | 0.00 | 4766 | 0 | O1 | 423054 | Eimeriorina |
| 315 | 0.00 | 4766 | 0 | F | 5809 | Sarcocystidae |
| 316 | 0.00 | 4766 | 0 | G | 5810 | Toxoplasma |
| 317 | 0.00 | 4766 | 0 | S | 5811 | Toxoplasma gondii |
| 318 | 0.00 | 4747 | 4747 | S1 | 508771 | Toxoplasma gondii ME49 |
| 319 | 0.00 | 19 | 19 | S1 | 383379 | Toxoplasma gondii RH |
| 320 | 0.01 | 11363 | 262 | D1 | 33154 | Opisthokonta |
| 321 | 0.01 | 9999 | 0 | K | 33208 | Metazoa |
| 322 | 0.01 | 9999 | 0 | K1 | 6072 | Eumetazoa |
| 323 | 0.01 | 9999 | 0 | K2 | 33213 | Bilateria |
| 324 | 0.01 | 9999 | 84 | K3 | 33317 | Protostomia |
| 325 | 0.01 | 9513 | 0 | K4 | 2697495 | Spiralia |
| 326 | 0.01 | 9513 | 0 | K5 | 1206795 | Lophotrochozoa |
| 327 | 0.01 | 9513 | 177 | P | 6157 | Platyhelminthes |
| 328 | 0.01 | 7901 | 0 | C | 6199 | Cestoda |
| 329 | 0.01 | 7901 | 0 | C1 | 6200 | Eucestoda |
| 330 | 0.01 | 7901 | 0 | O | 6201 | Cyclophyllidea |
| 331 | 0.01 | 7901 | 0 | F | 6208 | Taeniidae |
| 332 | 0.01 | 7901 | 0 | G | 6202 | Taenia |
| 333 | 0.01 | 7901 | 7901 | S | 6204 | Taenia solium |
| 334 | 0.00 | 1435 | 0 | C | 6178 | Trematoda |
| 335 | 0.00 | 1435 | 0 | C1 | 6179 | Digenea |
| 336 | 0.00 | 1435 | 0 | O | 6180 | Strigeidida |
| 337 | 0.00 | 1435 | 0 | O1 | 31244 | Schistosomatoidea |
| 338 | 0.00 | 1435 | 0 | F | 31245 | Schistosomatidae |
| 339 | 0.00 | 1435 | 0 | G | 6181 | Schistosoma |
| 340 | 0.00 | 1435 | 1435 | S | 6183 | Schistosoma mansoni |
| 341 | 0.00 | 402 | 0 | K4 | 1206794 | Ecdysozoa |
| 342 | 0.00 | 402 | 0 | P | 6231 | Nematoda |
| 343 | 0.00 | 402 | 0 | C | 119088 | Enoplea |
| 344 | 0.00 | 402 | 0 | C1 | 1457286 | Dorylaimia |
| 345 | 0.00 | 402 | 0 | O | 6329 | Trichinellida |
| 346 | 0.00 | 402 | 0 | F | 6332 | Trichinellidae |
| 347 | 0.00 | 402 | 0 | G | 6333 | Trichinella |
| 348 | 0.00 | 402 | 402 | S | 6334 | Trichinella spiralis |
| 349 | 0.00 | 1102 | 0 | K | 4751 | Fungi |
| 350 | 0.00 | 1102 | 84 | K1 | 451864 | Dikarya |
| 351 | 0.00 | 748 | 4 | P | 4890 | Ascomycota |
| 352 | 0.00 | 676 | 0 | P1 | 716545 | saccharomyceta |
| 353 | 0.00 | 676 | 0 | P2 | 147538 | Pezizomycotina |
| 354 | 0.00 | 676 | 0 | P3 | 716546 | leotiomyceta |

| | | | | | | |
|---|---|---|---|---|---|---|
| 355 | 0.00 | 676 | 0 | C | 147545 | Eurotiomycetes |
| 356 | 0.00 | 676 | 27 | C1 | 451871 | Eurotiomycetidae |
| 357 | 0.00 | 491 | 7 | O | 33183 | Onygenales |
| 358 | 0.00 | 391 | 23 | F | 299071 | Ajellomycetaceae |
| 359 | 0.00 | 238 | 0 | G | 229219 | Blastomyces |
| 360 | 0.00 | 238 | 0 | S | 5039 | Blastomyces dermatitidis |
| 361 | 0.00 | 238 | 238 | S1 | 559297 | Blastomyces dermatitidis ER-3 |
| 362 | 0.00 | 130 | 0 | G | 5036 | Histoplasma |
| 363 | 0.00 | 130 | 0 | S | 5037 | Histoplasma capsulatum |
| 364 | 0.00 | 130 | 130 | S1 | 447093 | Histoplasma capsulatum G186AR |
| 365 | 0.00 | 93 | 0 | F | 33184 | Onygenaceae |
| 366 | 0.00 | 93 | 39 | G | 5500 | Coccidioides |
| 367 | 0.00 | 40 | 0 | S | 199306 | Coccidioides posadasii |
| 368 | 0.00 | 40 | 40 | S1 | 222929 | Coccidioides posadasii C735 delta SOWgp |
| 369 | 0.00 | 14 | 0 | S | 5501 | Coccidioides immitis |
| 370 | 0.00 | 14 | 14 | S1 | 246410 | Coccidioides immitis RS |
| 371 | 0.00 | 158 | 0 | O | 5042 | Eurotiales |
| 372 | 0.00 | 158 | 0 | F | 1131492 | Aspergillaceae |
| 373 | 0.00 | 158 | 0 | G | 5052 | Aspergillus |
| 374 | 0.00 | 158 | 0 | G1 | 2720872 | Aspergillus subgen. Fumigati |
| 375 | 0.00 | 158 | 0 | S | 746128 | Aspergillus fumigatus |
| 376 | 0.00 | 158 | 158 | S1 | 330879 | Aspergillus fumigatus Af293 |
| 377 | 0.00 | 68 | 0 | P1 | 451866 | Taphrinomycotina |
| 378 | 0.00 | 68 | 0 | C | 147553 | Pneumocystidomycetes |
| 379 | 0.00 | 68 | 0 | O | 37987 | Pneumocystidales |
| 380 | 0.00 | 68 | 0 | F | 44281 | Pneumocystidaceae |
| 381 | 0.00 | 68 | 13 | G | 4753 | Pneumocystis |
| 382 | 0.00 | 41 | 0 | S | 4754 | Pneumocystis carinii |
| 383 | 0.00 | 41 | 41 | S1 | 1408658 | Pneumocystis carinii B80 |
| 384 | 0.00 | 14 | 0 | S | 42068 | Pneumocystis jirovecii |
| 385 | 0.00 | 14 | 14 | S1 | 1408657 | Pneumocystis jirovecii RU7 |
| 386 | 0.00 | 270 | 0 | P | 5204 | Basidiomycota |
| 387 | 0.00 | 270 | 0 | P1 | 5302 | Agaricomycotina |
| 388 | 0.00 | 270 | 0 | C | 155616 | Tremellomycetes |
| 389 | 0.00 | 270 | 0 | O | 5234 | Tremellales |
| 390 | 0.00 | 270 | 0 | F | 1884633 | Cryptococcaceae |
| 391 | 0.00 | 270 | 166 | G | 5206 | Cryptococcus |
| 392 | 0.00 | 80 | 0 | G1 | 1884637 | Cryptococcus gattii species complex |
| 393 | 0.00 | 80 | 0 | S | 37769 | Cryptococcus gattii VGI |
| 394 | 0.00 | 80 | 80 | S1 | 367775 | Cryptococcus gattii WM276 |
| 395 | 0.00 | 24 | 0 | G1 | 1897064 | Cryptococcus neoformans species complex |
| 396 | 0.00 | 24 | 0 | S | 5207 | Cryptococcus neoformans |
| 397 | 0.00 | 24 | 0 | S1 | 40410 | Cryptococcus neoformans var. neoformans |
| 398 | 0.00 | 24 | 24 | S2 | 214684 | Cryptococcus neoformans var. neoformans JEC21 |
| 399 | 0.00 | 383 | 0 | D1 | 2611352 | Discoba |
| 400 | 0.00 | 383 | 0 | P | 33682 | Euglenozoa |
| 401 | 0.00 | 383 | 0 | C | 5653 | Kinetoplastea |
| 402 | 0.00 | 383 | 0 | C1 | 2704647 | Metakinetoplastina |
| 403 | 0.00 | 383 | 0 | O | 2704949 | Trypanosomatida |
| 404 | 0.00 | 383 | 0 | F | 5654 | Trypanosomatidae |
| 405 | 0.00 | 383 | 26 | G | 5690 | Trypanosoma |
| 406 | 0.00 | 284 | 0 | G1 | 47570 | Schizotrypanum |
| 407 | 0.00 | 284 | 284 | S | 5693 | Trypanosoma cruzi |
| 408 | 0.00 | 73 | 0 | G1 | 39700 | Trypanozoon |
| 409 | 0.00 | 73 | 0 | S | 5691 | Trypanosoma brucei |
| 410 | 0.00 | 73 | 0 | S1 | 5702 | Trypanosoma brucei brucei |
| 411 | 0.00 | 73 | 73 | S2 | 185431 | Trypanosoma brucei brucei TREU927 |
| 412 | 0.00 | 41 | 0 | D | 2157 | Archaea |
| 413 | 0.00 | 41 | 0 | P | 28890 | Euryarchaeota |
| 414 | 0.00 | 41 | 0 | P1 | 2283794 | Methanomada group |
| 415 | 0.00 | 41 | 0 | C | 183925 | Methanobacteria |
| 416 | 0.00 | 41 | 0 | O | 2158 | Methanobacteriales |
| 417 | 0.00 | 41 | 0 | F | 2159 | Methanobacteriaceae |
| 418 | 0.00 | 41 | 0 | G | 2172 | Methanobrevibacter |
| 419 | 0.00 | 41 | 41 | S | 66851 | Methanobrevibacter oralis |
| 420 | 0.00 | 3662 | 0 | D | 10239 | Viruses |
| 421 | 0.00 | 3660 | 0 | D1 | 2731341 | Duplodnaviria |
| 422 | 0.00 | 3660 | 0 | K | 2731360 | Heunggongvirae |
| 423 | 0.00 | 3660 | 0 | P | 2731361 | Peploviricota |
| 424 | 0.00 | 3660 | 0 | C | 2731363 | Herviviricetes |
| 425 | 0.00 | 3660 | 0 | O | 548681 | Herpesvirales |
| 426 | 0.00 | 3660 | 0 | F | 10292 | Herpesviridae |
| 427 | 0.00 | 3660 | 0 | F1 | 10293 | Alphaherpesvirinae |
| 428 | 0.00 | 3660 | 9 | G | 10319 | Varicellovirus |
| 429 | 0.00 | 3649 | 0 | S | 3050278 | Varicellovirus equidalpha4 |
| 430 | 0.00 | 3649 | 3649 | S1 | 10331 | Equid alphaherpesvirus 4 |
| 431 | 0.00 | 2 | 0 | S | 3050279 | Varicellovirus equidalpha8 |
| 432 | 0.00 | 2 | 2 | S1 | 39637 | Equid alphaherpesvirus 8 |
| 433 | 0.00 | 2 | 0 | D1 | 2732004 | Varidnaviria |
| 434 | 0.00 | 2 | 0 | K | 2732005 | Bamfordvirae |
| 435 | 0.00 | 2 | 0 | P | 2732007 | Nucleocytoviricota |
| 436 | 0.00 | 2 | 0 | C | 2732525 | Pokkesviricetes |
| 437 | 0.00 | 2 | 0 | O | 2732527 | Chitovirales |
| 438 | 0.00 | 2 | 0 | F | 10240 | Poxviridae |
| 439 | 0.00 | 2 | 0 | F1 | 10241 | Chordopoxvirinae |
| 440 | 0.00 | 2 | 0 | G | 10242 | Orthopoxvirus |
| 441 | 0.00 | 2 | 2 | S | 10243 | Cowpox virus |

| | Genbank Accession Number | Sample ID | Region/Location Specific | Country | Year of isolation | Years BP* | Reference |
|---|---|---|---|---|---|---|---|
| *Modern - EHV-1* | AY665713 | strain Ab4p | N/A | Britain | 1991** | 26** | Telford *et al.*, 1992 [1] |
| *Modern - EHV-4* | GCF_000846345/AF030027 | strain NS80567 | Irish Equine Centre, Johnstown, Naas, County Kildare | Ireland | 1998** | 19** | Telford *et al.*, 1998 [2] |
| | KT324735 | 1546-99 | Toolern Vale, Victoria | Australia | 1999 | 18 | Vaz *et al.*, 2016 [3] |
| | KT324736 | RT4-70 | Oakey, Queensland | Australia | 1970 | 47 | Vaz *et al.*, 2016 [3] |
| | KT324737 | 3409-77 | Gosnells, Western Australia | Australia | 1977 | 40 | Vaz et al., 2016 [3] |
| | KT324738 | 157-69 | Yarra Glen, Victoria | Australia | 1969 | 48 | Vaz et al., 2016 [3] |
| | KT324739 | 3407-77 | N/A | Australia | 1977 | 40 | Vaz et al., 2016 [3] |
| | KT324740 | 405-76 | Flemington, Victoria | Australia | 1976 | 41 | Vaz et al., 2016 [3] |
| | KT324741 | 960-90 | Diggers Rest, Victoria | Australia | 1990 | 27 | Vaz et al., 2016 [3] |
| | KT324742 | 2387-75 | Esperance, Western Australia | Australia | 1975 | 42 | Vaz et al., 2016 [3] |
| | KT324743 | 306-74 | Flemington, Victoria | Australia | 1974 | 43 | Vaz et al., 2016 [3] |
| | KT324744 | 1532-99 | Flemington, Victoria | Australia | 1999 | 18 | Vaz et al., 2016 [3] |
| | KT324745 | R19-68 | Toowoomba, Queensland | Australia | 1968 | 49 | Vaz *et al.*, 2016 [3]; Bagust & Pascoe 1968 [4] |
| | KT324746 | 3056-07 | Tasmania | Australia | 2007 | 10 | Vaz et al., 2016 [3] |
| | KT324747 | 475-78 | Mount Pleasant, Tasmania | Australia | 1978 | 39 | Vaz et al., 2016 [3] |
| | KT324748 | ER39-67 | South Australia | Australia | 1967 | 50 | Vaz et al., 2016 [3] |
| | LC063142 | TH20p | Aomori Prefecture | Japan | 1962 | 55 | Izume *et al.*, 2017 [5]; Maeda *et al.*, 2005 [6] |
| | LC075582 | 83-MB | N/A | Japan | 1983 | 34 | Izume *et al.*, 2017 [5] |
| | LC075583 | 91c1 | Saga Prefecture | Japan | 1991 | 26 | Izume *et al.*, 2017 [5]; Maeda *et al.*, 2005 [6] |
| | LC075584 | 01-10-2001 | N/A | Japan | 2001 | 16 | Izume *et al.*, 2017 [5] |
| | LC075585 | 03-VR | N/A | Japan | 2003 | 14 | Izume *et al.*, 2017 [5] |
| | LC075586 | 05-I-202 | N/A | Japan | 2005 | 12 | Izume *et al.*, 2017 [5] |
| | LC075587 | 11-10 | N/A | Japan | 2011 | 6 | Izume *et al.*, 2017 [5] |
| | LC075588 | 12-I-203 | N/A | Japan | 2012 | 5 | Izume *et al.*, 2017 [5] |
| | MW892435 | DE17_1 | Northern Germany | Germany | 2017 | 0 | Pavulraj *et al.*, 2021 [7] |
| | MW892436 | DE17_2 | Northern Germany | Germany | 2017 | 0 | Pavulraj et al., 2021 [7] |
| | MW892437 | DE17_3 | Northern Germany | Germany | 2017 | 0 | Pavulraj et al., 2021 [7] |
| | MW892438 | DE17_4 | Northern Germany | Germany | 2017 | 0 | Pavulraj et al., 2021 [7] |
| *Ancient - EHV-4* | SAMEA9533417 (BioSample) | UR17x29 | Kamenyi Ambar 5 | Russia | 1853 BC | 3870 | This study, Librado et al., 2021 [8] |

* 'BP - Before Present' here takes 2017 as its BP reference based on the most recently sampled isolates. Dates used for the BEAST analyses.

** The year of collection for these samples is unknown. The date is based upon when the work was first presented.

**Table S4:** EHV-1 and EHV-4 sequences used in this study with corresponding Genbank accession number, location and sampling date of isolates. For UR17x29, site location and sample age.

| | EHV1_AY665713_26 | Ural-2017-29_3869 | LC075588_Japan_5 | LC075587_Japan_6 | LC075584_Japan_16 | KT324741_Australia_27 | KT324742_Australia_42 | GCF000846345_Ireland_19 | LC075583_Japan_26 | KT324740_Australia_41 | KT324743_Australia_43 | KT324737_Australia_40 | KT324739_NA_40 | KT324738_Australia_48 | LC075585_Japan_14 | MW892437_Germany_0 | MW892436_Germany_0 | MW892438_Germany_0 | MW892435_Germany_0 | KT324746_Australia_10 | LC075582_Japan_34 | KT324736_Australia_47 | KT324745_Australia_49 | KT324735_Australia_18 | KT324748_Australia_50 | KT324744_Australia_18 | KT324747_Australia_39 | LC063142_Japan_55 | LC075586_Japan_12 |
|---|---|---|---|---|---|---|---|---|---|---|---|---|---|---|---|---|---|---|---|---|---|---|---|---|---|---|---|---|---|
| **EHV1_AY665713_26** | 0.00000 | | | | | | | | | | | | | | | | | | | | | | | | | | | | |
| **Ural-2017-29_3869** | 0.32739 | 0.00000 | | | | | | | | | | | | | | | | | | | | | | | | | | | |
| **LC075588_Japan_5** | 0.32882 | 0.00316 | 0.00000 | | | | | | | | | | | | | | | | | | | | | | | | | | |
| **LC075587_Japan_6** | 0.32890 | 0.00324 | 0.00064 | 0.00000 | | | | | | | | | | | | | | | | | | | | | | | | | |
| **LC075584_Japan_16** | 0.32885 | 0.00319 | 0.00078 | 0.00086 | 0.00000 | | | | | | | | | | | | | | | | | | | | | | | | |
| **KT324741_Australia_27** | 0.32891 | 0.00325 | 0.00084 | 0.00092 | 0.00029 | 0.00000 | | | | | | | | | | | | | | | | | | | | | | | |
| **KT324742_Australia_42** | 0.32882 | 0.00315 | 0.00075 | 0.00083 | 0.00019 | 0.00010 | 0.00000 | | | | | | | | | | | | | | | | | | | | | | |
| **GCF000846345_Ireland_19** | 0.32891 | 0.00325 | 0.00084 | 0.00092 | 0.00081 | 0.00087 | 0.00078 | 0.00000 | | | | | | | | | | | | | | | | | | | | | |
| **LC075583_Japan_26** | 0.32896 | 0.00330 | 0.00089 | 0.00097 | 0.00086 | 0.00092 | 0.00082 | 0.00090 | 0.00000 | | | | | | | | | | | | | | | | | | | | |
| **KT324740_Australia_41** | 0.32878 | 0.00312 | 0.00071 | 0.00079 | 0.00068 | 0.00074 | 0.00064 | 0.00072 | 0.00076 | 0.00000 | | | | | | | | | | | | | | | | | | | |
| **KT324743_Australia_43** | 0.32883 | 0.00316 | 0.00076 | 0.00084 | 0.00073 | 0.00078 | 0.00069 | 0.00077 | 0.00081 | 0.00020 | 0.00000 | | | | | | | | | | | | | | | | | | |
| **KT324737_Australia_40** | 0.32883 | 0.00316 | 0.00076 | 0.00084 | 0.00073 | 0.00078 | 0.00069 | 0.00078 | 0.00083 | 0.00065 | 0.00070 | 0.00000 | | | | | | | | | | | | | | | | | |
| **KT324739_NA_40** | 0.32883 | 0.00316 | 0.00076 | 0.00084 | 0.00073 | 0.00078 | 0.00069 | 0.00078 | 0.00083 | 0.00065 | 0.00070 | 0.00000 | 0.00000 | | | | | | | | | | | | | | | | |
| **KT324738_Australia_48** | 0.32881 | 0.00314 | 0.00074 | 0.00082 | 0.00071 | 0.00076 | 0.00067 | 0.00076 | 0.00081 | 0.00063 | 0.00068 | 0.00010 | 0.00010 | 0.00000 | | | | | | | | | | | | | | | |
| **LC075585_Japan_14** | 0.32885 | 0.00319 | 0.00078 | 0.00087 | 0.00075 | 0.00081 | 0.00072 | 0.00081 | 0.00086 | 0.00068 | 0.00073 | 0.00015 | 0.00015 | 0.00007 | 0.00000 | | | | | | | | | | | | | | |
| **MW892437_Germany_0** | 0.32902 | 0.00336 | 0.00164 | 0.00172 | 0.00167 | 0.00173 | 0.00163 | 0.00173 | 0.00178 | 0.00160 | 0.00165 | 0.00165 | 0.00165 | 0.00163 | 0.00167 | 0.00000 | | | | | | | | | | | | | |
| **MW892436_Germany_0** | 0.32903 | 0.00337 | 0.00165 | 0.00173 | 0.00168 | 0.00174 | 0.00164 | 0.00174 | 0.00179 | 0.00161 | 0.00165 | 0.00165 | 0.00165 | 0.00163 | 0.00168 | 0.00001 | 0.00000 | | | | | | | | | | | | |
| **MW892438_Germany_0** | 0.32904 | 0.00338 | 0.00166 | 0.00174 | 0.00169 | 0.00175 | 0.00165 | 0.00175 | 0.00180 | 0.00161 | 0.00166 | 0.00166 | 0.00166 | 0.00164 | 0.00169 | 0.00002 | 0.00003 | 0.00000 | | | | | | | | | | | |
| **MW892435_Germany_0** | 0.32903 | 0.00337 | 0.00165 | 0.00173 | 0.00168 | 0.00174 | 0.00164 | 0.00174 | 0.00179 | 0.00161 | 0.00165 | 0.00165 | 0.00165 | 0.00163 | 0.00168 | 0.00001 | 0.00002 | 0.00001 | 0.00000 | | | | | | | | | | |
| **KT324746_Australia_10** | 0.32908 | 0.00342 | 0.00170 | 0.00178 | 0.00173 | 0.00179 | 0.00169 | 0.00179 | 0.00184 | 0.00166 | 0.00170 | 0.00170 | 0.00170 | 0.00168 | 0.00173 | 0.00149 | 0.00150 | 0.00151 | 0.00150 | 0.00000 | | | | | | | | | |
| **LC075582_Japan_34** | 0.32917 | 0.00351 | 0.00179 | 0.00187 | 0.00182 | 0.00188 | 0.00178 | 0.00188 | 0.00193 | 0.00174 | 0.00179 | 0.00179 | 0.00179 | 0.00177 | 0.00182 | 0.00158 | 0.00159 | 0.00160 | 0.00159 | 0.00061 | 0.00000 | | | | | | | | |
| **KT324736_Australia_47** | 0.32911 | 0.00344 | 0.00172 | 0.00180 | 0.00175 | 0.00181 | 0.00172 | 0.00181 | 0.00186 | 0.00168 | 0.00173 | 0.00173 | 0.00173 | 0.00171 | 0.00175 | 0.00152 | 0.00153 | 0.00153 | 0.00153 | 0.00054 | 0.00019 | 0.00000 | | | | | | | |
| **KT324745_Australia_49** | 0.32910 | 0.00343 | 0.00171 | 0.00179 | 0.00174 | 0.00180 | 0.00171 | 0.00180 | 0.00185 | 0.00167 | 0.00172 | 0.00172 | 0.00172 | 0.00170 | 0.00175 | 0.00151 | 0.00152 | 0.00152 | 0.00152 | 0.00053 | 0.00018 | 0.00011 | 0.00000 | | | | | | |
| **KT324735_Australia_18** | 0.32911 | 0.00345 | 0.00173 | 0.00181 | 0.00176 | 0.00182 | 0.00173 | 0.00182 | 0.00187 | 0.00169 | 0.00174 | 0.00174 | 0.00174 | 0.00172 | 0.00176 | 0.00153 | 0.00154 | 0.00154 | 0.00154 | 0.00055 | 0.00019 | 0.00012 | 0.00010 | 0.00000 | | | | | |
| **KT324748_Australia_50** | 0.32912 | 0.00346 | 0.00174 | 0.00182 | 0.00177 | 0.00183 | 0.00173 | 0.00183 | 0.00188 | 0.00170 | 0.00175 | 0.00175 | 0.00175 | 0.00173 | 0.00177 | 0.00154 | 0.00154 | 0.00155 | 0.00154 | 0.00056 | 0.00020 | 0.00014 | 0.00013 | 0.00015 | 0.00000 | | | | |
| **KT324744_Australia_18** | 0.32911 | 0.00345 | 0.00173 | 0.00181 | 0.00176 | 0.00182 | 0.00173 | 0.00182 | 0.00187 | 0.00169 | 0.00174 | 0.00174 | 0.00174 | 0.00172 | 0.00176 | 0.00153 | 0.00153 | 0.00154 | 0.00153 | 0.00055 | 0.00019 | 0.00013 | 0.00012 | 0.00014 | 0.00007 | 0.00000 | | | |
| **KT324747_Australia_39** | 0.32936 | 0.00369 | 0.00197 | 0.00205 | 0.00200 | 0.00206 | 0.00197 | 0.00206 | 0.00211 | 0.00193 | 0.00198 | 0.00198 | 0.00198 | 0.00196 | 0.00201 | 0.00177 | 0.00178 | 0.00178 | 0.00178 | 0.00079 | 0.00044 | 0.00037 | 0.00036 | 0.00038 | 0.00037 | 0.00036 | 0.00000 | | |
| **LC063142_Japan_55** | 0.32917 | 0.00351 | 0.00179 | 0.00187 | 0.00182 | 0.00188 | 0.00178 | 0.00188 | 0.00193 | 0.00174 | 0.00179 | 0.00179 | 0.00179 | 0.00177 | 0.00182 | 0.00158 | 0.00159 | 0.00160 | 0.00159 | 0.00061 | 0.00025 | 0.00018 | 0.00017 | 0.00019 | 0.00018 | 0.00017 | 0.00040 | 0.00000 | |
| **LC075586_Japan_12** | 0.32916 | 0.00350 | 0.00178 | 0.00186 | 0.00181 | 0.00187 | 0.00177 | 0.00187 | 0.00192 | 0.00174 | 0.00178 | 0.00178 | 0.00178 | 0.00176 | 0.00181 | 0.00157 | 0.00158 | 0.00159 | 0.00158 | 0.00060 | 0.00024 | 0.00017 | 0.00016 | 0.00018 | 0.00017 | 0.00016 | 0.00039 | 0.00016 | 0.00000 |

**Table S5:** Pairwise genetic distances computed from the maximum likelihood phylogenetic tree using its branch lengths through the cophenetic.phylo function in the R package ape.

| Recombination Event No. | Recombinant | Beginning of breakpoint | Ending of breakpoint | RDP (*p*-value) | GENECONV (*p*-value) | BootScan (*p*-value) | MaxChi (*p*-value) | Chimaera (*p*-value) | SiScan (*p*-value) | 3 Seq (*p*-value) | Recombination score |
|---|---|---|---|---|---|---|---|---|---|---|---|
| 1 | KT324743 | 1 | 4253 | - | 4.55E-08 | - | - | - | 3.22E-02 | 2.87E-05 | 0.725 |
| 2 | KT324747 | 90129 | 112590 | - | 7.76E-03 | - | 1.01E-06 | 1.72E-02 | 1.34E-22 | 1.61E-04 | 0.589 |
| 3 | LC075588 | 97230 | 144956 | 2.32E-02 | 9.04E-04 | - | 1.38E-06 | 7.38E-05 | 9.39E-07 | - | 0.459 |
| 4 | KT324737 | 122386 | 135664 | 4.56E-06 | 2.31E-03 | - | 6.28E-04 | 1.43E-02 | 1.45E-18 | 1.76E-03 | 0.633 |
| 5 | KT324741 | 126598 | 127411 | - | 2.75E-05 | - | - | - | - | - | 0.637 |
| 6 | KT324735 | 112500 | 112590 | - | 2.87E-04 | - | - | - | - | - | 0.596 |
| 7 | LC075583 | 48 | 2324 | 5.85E-03 | - | 4.22E-02 | - | - | - | - | 0.714 |
| 8 | KT324744 | 135347 | 135969 | - | 1.78E-02 | - | - | - | - | - | 0.661 |
| 9 | KT324744 | 122083 | 122951 | - | 9.47E-04 | 9.49E-04 | - | - | - | - | 0.618 |
| 10 | LC075586 | 122032 | 136018 | - | - | - | 2.51E-02 | 1.38E-02 | - | 1.86E-01 | 0.545 |
| 11* | KT324737 | 127007 | Undetermined - 135426 | - | - | - | 1.89E-03 | 5.58E-03 | - | 3.15E-05 | 0.698 |
| 12* | LC075586 | 123344 | 135590 | - | - | - | - | - | - | 9.29E-03 | 0.539 |

* Following rescanning run

**Table S6:** Whole genome recombination events and detection of recombination breakpoints using the seven recombination detection methods (RDP, GENECONV, Chimaera, MaxChi, BootScan, SiScan and 3Seq) implemented in RDP5 [9]. Nucleotide positions based on strain NS80567 (AF030027) [2] and excluding gaps.
